# Helical Twists and β-Turns in Structures at Serine–Proline Sequences: Stabilization of cis-Proline and type VI β-turns via C–H/O interactions

**DOI:** 10.1101/2024.03.14.585129

**Authors:** Harrison C. Oven, Glenn P. A. Yap, Neal J. Zondlo

**Affiliations:** Department of Chemistry and Biochemistry, University of Delaware Newark, DE 19716 United States

## Abstract

Structures at serine-proline sites in proteins were analyzed using a combination of peptide synthesis with structural methods and bioinformatics analysis of the PDB. Dipeptides were synthesized with the proline derivative (2*S*,4*S*)-(4-iodophenyl)hydroxyproline [hyp(4-I-Ph)]. The crystal structure of Boc-Ser-hyp(4-I-Ph)-OMe had two molecules in the unit cell. One molecule exhibited *cis*-proline and a type VIa2 β-turn (BcisD). The *cis*-proline conformation was stabilized by a C–H/O interaction between Pro C–H_α_ and the Ser side-chain oxygen. NMR data were consistent with stabilization of *cis*-proline by a C–H/O interaction in solution. The other crystallographically observed molecule had *trans*-Pro and both residues in the PPII conformation. Two conformations were observed in the crystal structure of Ac-Ser-hyp(4-I-Ph)-OMe, with Ser adopting PPII in one and the β conformation in the other, each with Pro in the δ conformation and *trans*-Pro. Structures at Ser-Pro sequences were further examined via bioinformatics analysis of the PDB and via DFT calculations. Ser–Pro *versus* Ala-Pro sequences were compared to identify bases for Ser stabilization of local structures. C–H/O interactions between the Ser side-chain O_γ_ and Pro C–H_α_ were observed in 45% of structures with Ser-*cis*- Pro in the PDB, with nearly all Ser-*cis*-Pro structures adopting a type VI β-turn. 53% of Ser- *trans*-Pro sequences exhibited main-chain C=O*_i_*•••H–N*_i_*_+3_ or C=O*_i_*•••H–N*_i_*_+4_ hydrogen bonds, with Ser as the *i* residue and Pro as the *i*+1 residue. These structures were overwhelmingly either type I β-turns or N-terminal capping motifs on α-helices or a 3_10_-helices. These results indicate that Ser-Pro sequences are particularly potent in favoring these structures. In each, Ser is in either the PPII or β conformation, with the Ser O_γ_ capable of engaging in a hydrogen bond with the amide N–H of the *i*+2 (type I β-turn or 3 -helix; Ser *χ_1_ t*) or *i*+3 (α-helix; Ser *χ_1_ g^+^*) residue. Non-proline *cis* amide bonds can also be stabilized by C–H/O interactions.

Graphical Table of Contents

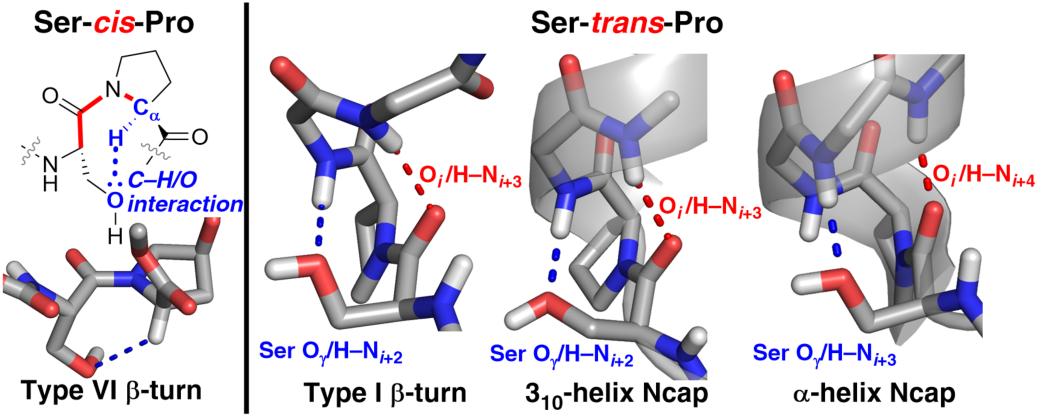

## Introduction

Serine–proline (Ser–Pro, SP) sequences are pervasive in protein structures. There are over 68,000 SP sequences in the human proteome, with SP sequences present in 76% of all human proteins.^1^ Proline residues are inherently sites of structural modulation in proteins, capable of inducing disorder (via disruption of hydrogen-bonded α-helical and β-sheet secondary structures) or order (via stabilization of turns, loops, or the PPII conformation).^2–4^ Ser-Pro sequences are important both in globular proteins and in intrinsically disordered regions of proteins (IDPs).^5,6^ Of particular note, the human RNA Polymerase II C-terminal domain contains 52 imperfect repeats of the YSPTSPS sequence, with two SP sequences in each consensus repeat.^7–9^ In addition, Ser–Pro sequences represent 25% of all phosphorylation sites *in vivo*.^10^ A better understanding of the structural preferences at Ser–Pro sequences would provide insights into structures in IDPs, into inherent questions in protein structure and folding, and into how protein phosphorylation at SP sites can change protein structure.^11^

In addition to main-chain conformational preferences at the *φ* and *ψ* torsion angles, Pro sequences exhibit *cis*-*trans* isomerism about the X–Pro amide bond (*ω* torsion angle) (Figure 1). The identity of the residue before Pro significantly impacts the frequency of *cis*-Pro and the structures observed, in part via local interactions of the X residue with Pro or with adjacent residues.^2,12–15^ We recently proposed that *cis*-Pro is stabilized in phosphoserine–proline (pSer-Pro) and Glu–Pro sequences via a C–H/O interaction between the side-chain anionic oxygen and Pro C–H_α_.^16^ More broadly, Pro residue C–H bonds are particularly likely to interact with oxygens via C–H/O interactions, due to the presence of multiple polarized C–H bonds on the proline ring.^17^ C–H/O interactions are also observed with neutral oxygens of water molecules, alcohols, and carbonyls.^18–24^ Thus, we hypothesized that the Ser side-chain oxygen might modulate structure via C–H/O interactions with the proline ring C–H bonds.

**Figure 1.**
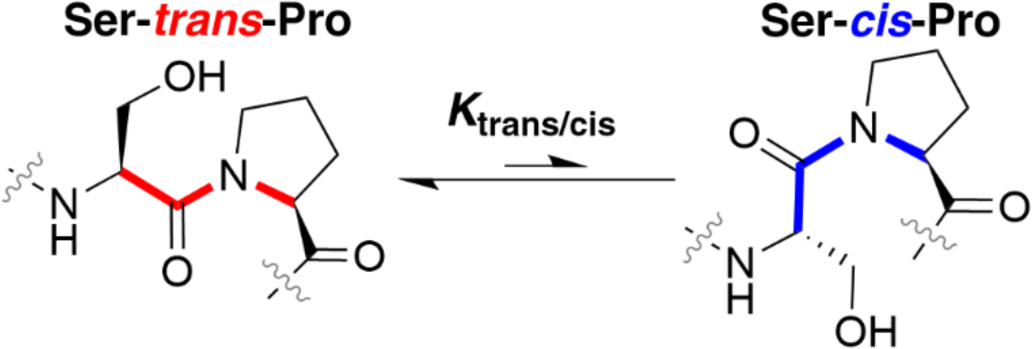
*trans*-Proline and *cis*-proline in Ser-Pro sequences.

Herein, we conduct a comprehensive analysis of protein structure at Ser–Pro sequences, including potential structural stabilization mediated by the Ser side-chain hydroxyl.^25–28^ This work was performed using a combination of small-molecule synthesis and structural analysis in the solid and solution states; via bioinformatics analysis of Ser–Pro sequences in the PDB; and via DFT calculations on structures observed in Ser–Pro motifs.

## Results and Discussion

### Synthesis of Boc-Ser-hyp(4-I-Ph)-OMe and Ac-Ser-hyp(4-I-Ph)

In order to examine Ser– Pro structures via small-molecule X-ray crystallography, we synthesized dipeptides with the unnatural proline derivative (2*S*,4*S*)-(4-iodophenyl)hydroxyproline [hyp(4-I-Ph)] (Scheme 1).^17,29,30^ This amino acid promotes crystallization via the aryl iodide. The aryl iodide has also been employed for further amino acid modification via Suzuki or Sonogashira cross-coupling reactions, which can be conducted on peptides in water.^29,30^ In addition, hyp(4-I-Ph) increases the likelihood of a *cis*-proline amide bond via the 4*S*-substitution of the aryloxy group, which leads to a greater preference for the proline *endo* ring pucker, the predominant ring pucker observed in *cis*-proline.^31–33^

**Scheme 1.**
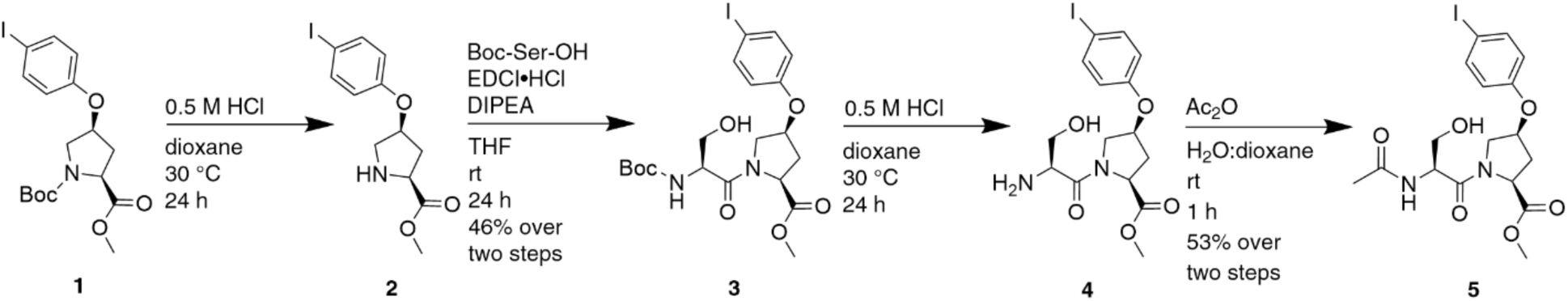
Synthesis of Ac-Ser-hyp(4-I-Ph)-OMe.

Ser–Pro dipeptides were synthesized in solution from Boc-hyp(4-I-Ph)-OMe (**1**), whose synthesis was described previously.^30^ Boc deprotection and amide coupling with Boc-Ser-OH and EDCI generated Boc-Ser-hyp(4-I-Ph)-OMe (**3**) (Scheme 1). In addition, **3** was converted into Ac-Ser-hyp(4-I-Ph) (**5**) in two steps via Boc deprotection and acetylation. In order to understand the specific effects of the Ser hydroxyl on structure, the Ala derivatives Boc-Ala-hyp(4-I-Ph)-OMe (**6**) and Ac-Ala-hyp(4-I-Ph)-OMe (**8**) were synthesized as controls (Scheme S2).

### X-ray crystallography of Ser-Pro dipeptides

Boc-Ser-hyp(4-I-Ph)-OMe crystallized from acetone and the crystal structure was determined. Interestingly, two molecules with distinct conformations were present in the unit cell, with one molecule exhibiting a *cis*-proline amide bond and the other molecule exhibiting a *trans*-proline amide bond (Figure 2). Crystal assembly was mediated by intermolecular hydrogen bonds between Ser hydroxyls; by C–H/O interactions at Pro C–H_δ_ and at both diastereotopic Ser C–H_β_; and by a particularly close (2.94 Å) C=O•••I distance consistent with a strong halogen bond^34^ between a Boc carbonyl and an aryl iodide.

**Figure 2.**
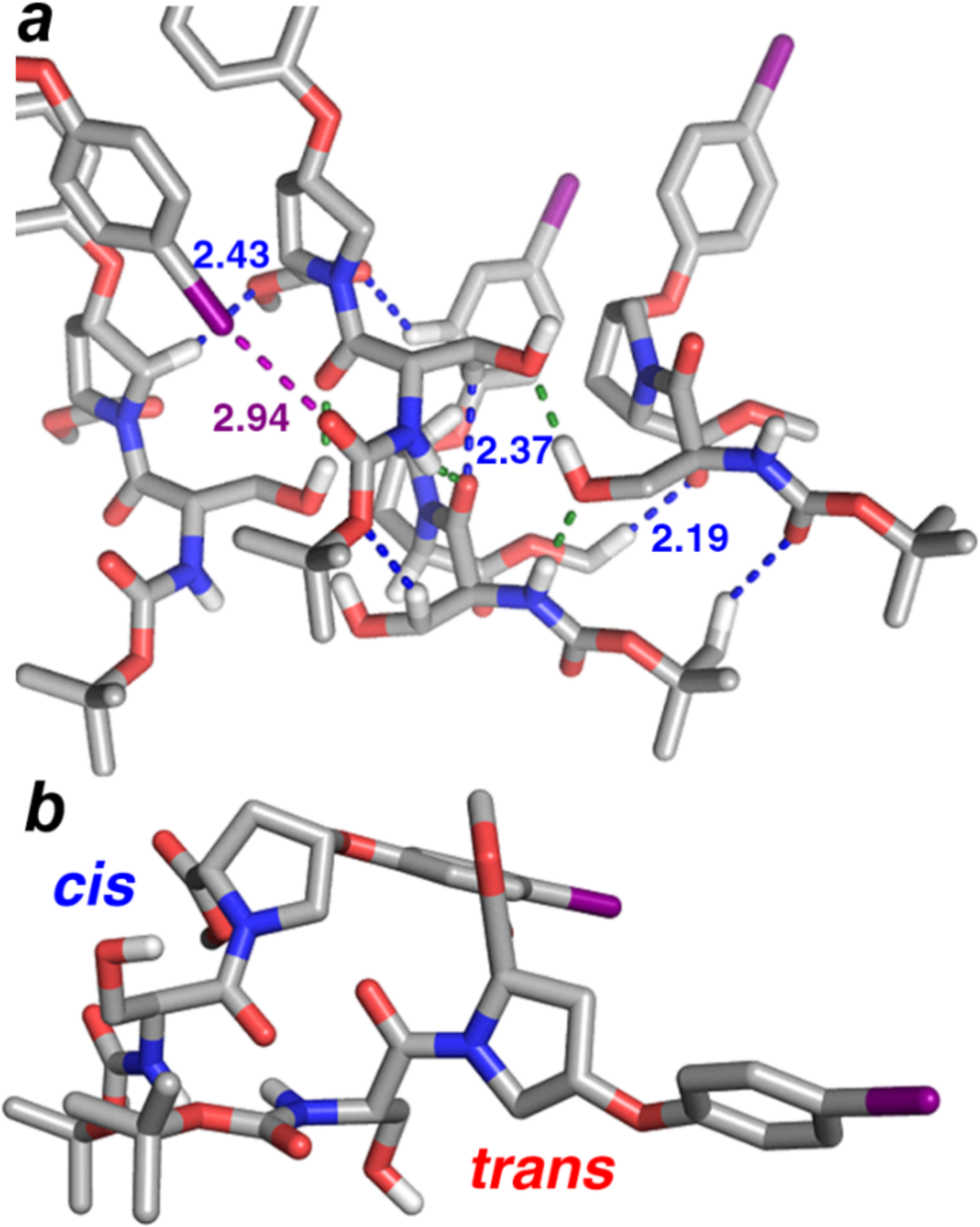
X-ray crystal structure of Boc-Ser-hyp(4-I-Ph)-OMe with two molecules in the unit cell, one with *trans*-proline and one with *cis*-proline. (a) Crystal assembly is mediated by the intermolecular interactions including C–H/O interactions, blue; N–H•••O and O–H•••O hydrogen bonds, green; and a C–I•••O halogen bond, purple. Interatomic distances of the halogen bond and of close C–H/O interactions are indicated (Å). (b) The unit cell contains one molecule with the *cis*-proline conformation and one molecule with the *trans*-proline conformation.

In the molecule with *cis*-Pro (*ω* = –12°), Ser was in the β conformation and Pro was in the δ conformation (Figure 3). The overall structure was a type VIa2 β-turn (BcisD in the Dunbrack β-turn nomenclature^35^).^25,26,36–38^ Type VIa2 β-turns are not mediated by hydrogen bonds between the *i* and *i*+3 residues of the turn, but are defined by the torsion angles and proximity of these termini. In this structure, the *cis*-Pro conformation appears to be stabilized by a C–H/O interaction between the Ser O_γ_ lone pair(s) and the Pro C–H_α_, mediated via the Ser *t χ*_1_ rotamer. C–H/O interactions are observed in diverse contexts in protein structures, including in recognition of α-helical GXXXG motifs via interaction of a main-chain carbonyl with Gly C–H_α_.^23,39–43^ C–H/O interactions are also observed at proline kinks in α-helices, where interaction with Pro C–H_α_ replaces a typical hydrogen bonding interaction with an amide hydrogen.^22,44^ C– H/O interactions also stabilize the *ζ* (pre-proline) conformation [(*φ*,*ψ*) ∼ (–130°, +60°)], where the carbonyl oxygen of the residue *i*–2 to proline interacts with the Pro C–H_δ_.^35,45,46^

**Figure 3.**
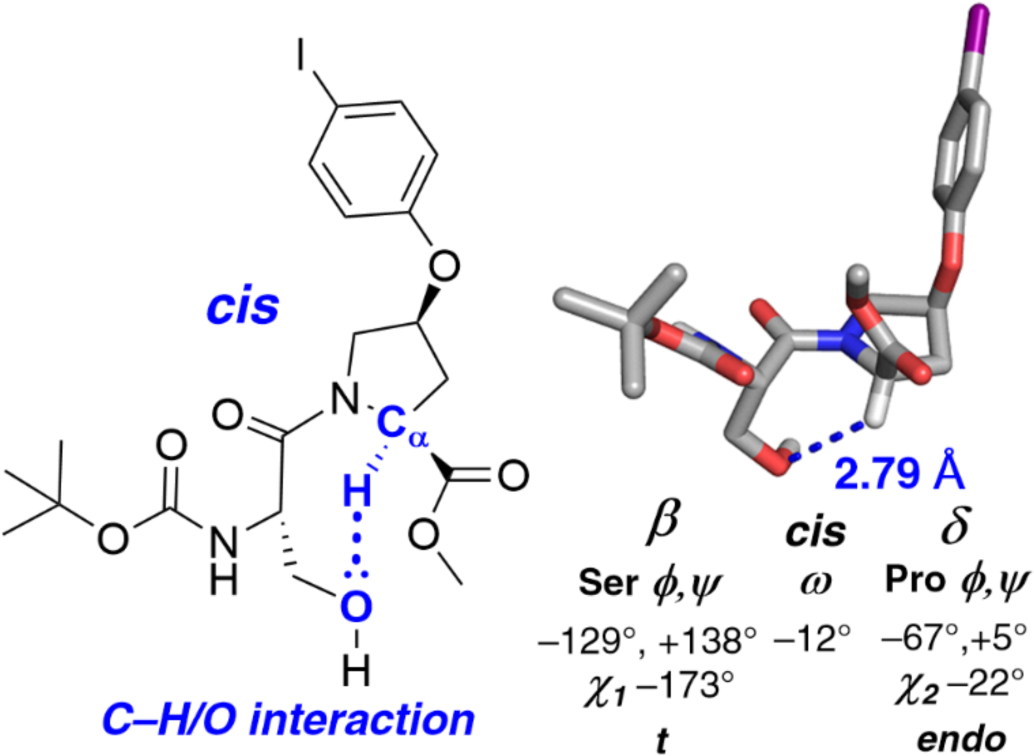
Molecule from the X-ray structure of Boc-Ser-hyp(4-I-Ph)-OMe with Ser-*cis*-Pro and an intramolecular proline-serine C–H/O interaction. One molecule in the unit cell exhibited the *cis*-proline amide conformation, with an *endo* ring pucker, the β conformation at Ser, and the δ conformation at Pro. This conformation represents a type VIa2 (BcisD) β-turn. The O_Boc_**•••**O_OMe_ distance, which is equivalent to a C_α, *i*_•••C_α,_*_i+_*_3_ distance in a protein context, is 4.67 Å. This molecule has a C–H/O interaction between Ser O_g_ and Pro H_α_ (C_α_–H•••O distance 2.79 Å, blue). The positions of the hydrogens, which are not definitively determined in the crystal structure, were optimized using the M06-2X DFT functional with the Def2TZVP basis set and implicit H_2_O solvation, while keeping the crystallographically determined positions of the heavy atoms fixed.

The molecule of Boc-Ser-hyp(4-I-Ph)-OMe with *trans*-Pro exhibited both Ser and Pro in the PPII conformation (Figure 4). The *n* →π* interaction across Ser was relatively weak, with a 3.14 Å O •••C=O distance, consistent with the low PPII propensity of Ser.^47–49^ The Ser side chain was in the *g*^−^ *χ_1_* conformation. Pro exhibited an *exo* ring pucker, consistent with the relatively modest *endo/exo* ring pucker preference of aryloxy groups.^17,30^

**Figure 4.**
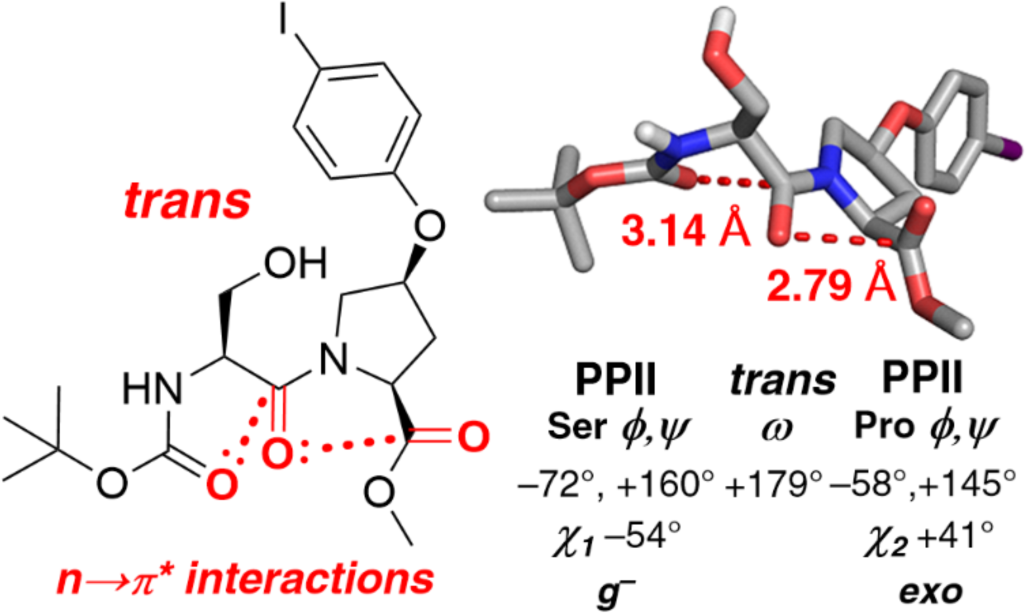
Molecule from the X-ray structure of Boc-Ser-hyp(4-I-Ph)-OMe with Ser-*trans*- Pro. One molecule in the unit cell exhibited the *trans-*proline amide conformation, with the PPII conformation at both Ser and Pro, with PPII stabilized by consecutive n→*π\** interactions.

Ac-Ser-hyp(4-I-Ph)-OMe crystallized with two different conformations for the molecule in the unit cell (Figure 5). Crystal packing (Figure 5a) was again mediated by hydrogen bonds, C–H/O interactions, and halogen bonds. Both conformations had a *trans*-proline amide bond with Pro in the δ conformation. In one conformation (Figure 5b), Ser was in the PPII conformation, with an intraresidue side chain-amide hydrogen bond similar to (but significantly longer than) that observed for phosphothreonine and phosphoserine.^11,50,51^ As had been observed for Boc-Ser-hyp(4-I-Ph)-OMe, the *n* →p* interaction across Ser was relatively weak. This conformation also exhibited Ser in the *g*^+^ *χ_1_* conformation. Thus, across all structures, all three Ser *χ*_1_ rotamers were represented, similar to what is observed in the PDB, where a relatively even distribution of Ser *χ_1_* rotamers is present.^52,53^

**Figure 5.**
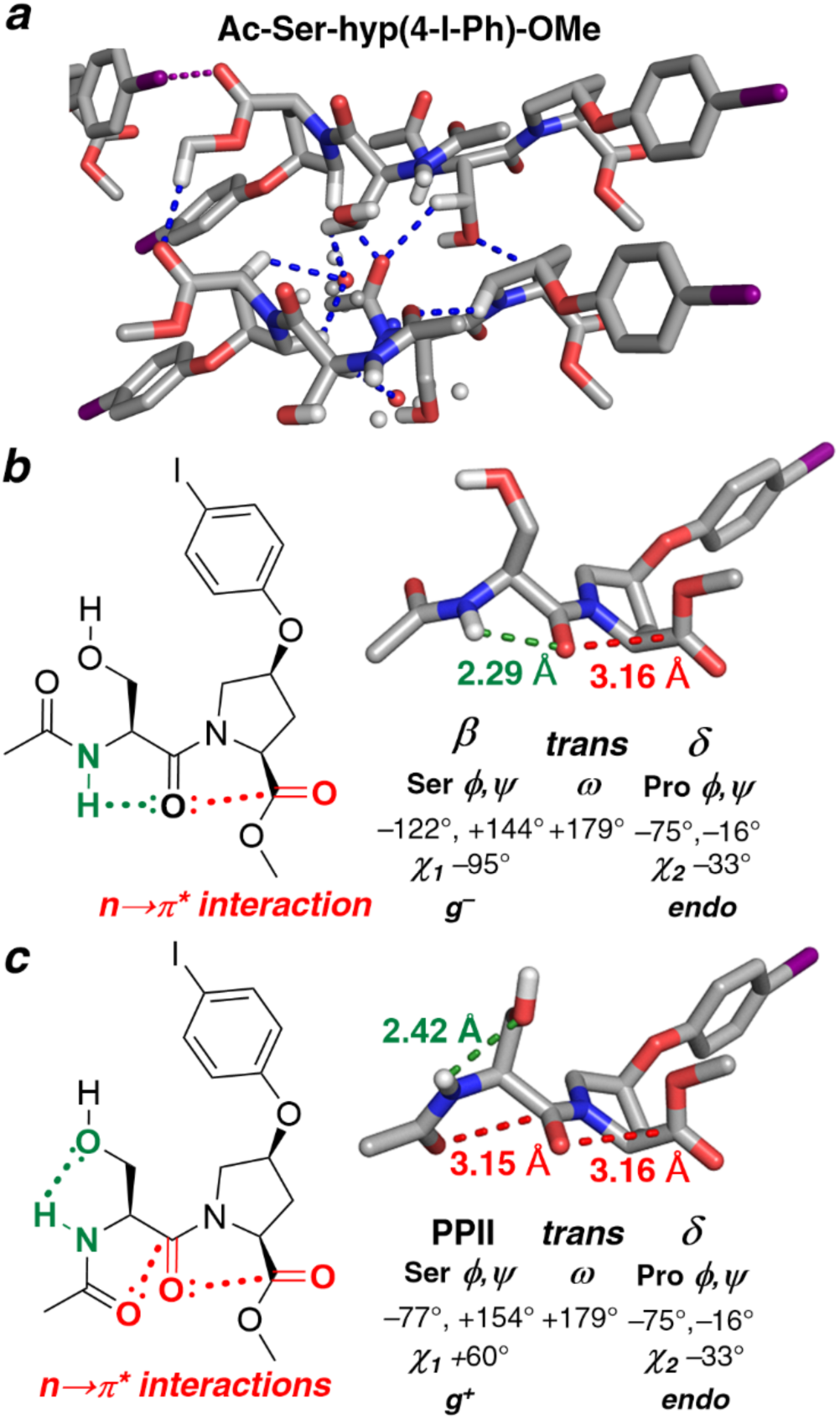
Crystal structure of Ac-Ser-hyp(4-I-Ph)-OMe. =(a). Crystal packing, with intermolecular C–H/O interactions (blue) and C–I•••O halogen bonds (purple) highlighted. In the unit cell, two distinct conformations were observed at Ser (β or PPII). Pro was in the δ conformation with a *trans*-Pro amide for both conformations at Ser. (b) In one conformation, Ser is in the β conformation, with the amide hydrogen engaged in an intraresidue backbone C5 hydrogen bond (N–H•••O distance 2.29 Å, green). The Ser side chain is not involved in any intramolecular hydrogen bonds. (c) In the other conformation, Ser is in the PPII conformation. The Ser side chain O_γ_ is engaged in an intraresidue hydrogen bond with its amide hydrogen (N– H•••O distance 2.42 Å, green).

In the other conformation in the unit cell (Figure 5c), Ser was in the β conformation. The β structure was stabilized by an intraresidue C5 hydrogen bond.^54^ A relatively long (weak) *n* →p* interaction was observed across Pro. This *n* →p* interaction is potentially weakened by the C5 hydrogen bond reducing electron density at the Ser carbonyl, making it a weaker electron donor for *n* →p* interactions.^55^

In order to further understand the structures observed crystallographically, all structures were subjected to full geometry optimization via DFT computational methods (Figure S1). These calculations can identify whether the structures observed represent local energy minima, or alternatively whether interactions identified might be artefacts resulting from crystal packing. All four geometry-optimized structures exhibited conformations very similar to those observed crystallographically. In particular, the C–H/O interaction in Ser-*cis*-Pro was confirmed to be present in the local energy minimum identified via these calculations. In all cases, the computationally optimized and X-ray structures were very similar in all torsion angles and in their intramolecular noncovalent interactions. Geometry optimization did result in closer *n* →p* interactions across Ser than were observed in the solid state. The calculations, by allowing full optimization of the position of the Ser hydroxyl hydrogen, also identified a potential O–H•••N interaction with the Pro amide nitrogen (analogous to N–H/N interactions^56,57^) that could help stabilize the δ structure at Pro in Ac-Ser-hyp(4-I-Ph)-OMe.

### Analysis of dipeptides by NMR spectroscopy in solution

We sought to identify whether the C–H/O interaction observed at *cis*-proline in the solid state was present in solution. Therefore, we examined the NMR spectra of Ac-Ser-hyp(4-I-Ph)-OMe and Boc-Ser-hyp(4-I-Ph)-OMe in methanol, comparing the NMR spectra to those of the dipeptides with Ala, which lacks the Ser hydroxyl (Table 1). Similar results were obtained either with an acetyl (Ac) or Boc N-terminal acyl group. Peptides with Ser had a significantly higher population of *cis*-proline (lower *K*_trans/cis_) than those with Ala. *cis*-Pro was relatively more stable than *trans*-Pro in the Ser peptides compared to the Ala peptides by ΔΔ*G* = 0.4–0.5 kcal mol^−1^. These results indicate either that the Ser hydroxyl stabilizes *cis*-proline, or that the Ser hydroxyl destabilizes *trans*-proline. Based on the crystallographic and computational data, we propose that Ser-*cis*-Pro is stabilized in part by a C–H/O interaction between the Ser oxygen and the Pro C–H_α_. Consistent with a C– H/O interaction,^11,16^ the chemical shift of Pro H_α_ is downfield by 0.26–0.29 ppm in Ser-*cis*-Pro compared to Ser-*trans*-Pro. In contrast, for Ala–Pro peptides, the chemical shifts of Pro H_α_ are similar for Ala-*cis*-Pro and Ala-*trans*-Pro, which were also similar to those in Ser-*trans*-Pro. Collectively, these results suggest that the C–H/O interaction observed crystallographically and computationally contributes to stabilizing the Ser-*cis*-Pro conformation, even in solution with competitive hydrogen-bonding groups.

**Table 1.** Summary of ^1^H NMR data on Ser-hyp(4-I-Ph) and Ala-hyp(4-I-Ph) dipeptides in d -MeOH.*^a^*^,*b*^ δ Pro H_α_, ppm.

| | $K_{\text{trans/cis}}$ | $\delta \text{ Pro H}_\alpha$, ppm | | |
| --- | --- | --- | --- | --- |
| | | $\delta_{\text{cis}}$ | $\delta_{\text{trans}}$ | $\Delta\delta_{\text{cis-trans}}$ |
| **Boc-Ser-hyp(4-I-Ph)-OMe** | 2.6 | 5.02 | 4.76 | +0.26 |
| **Boc-Ala-hyp(4-I-Ph)-OMe** | 5.9 | 4.78 | 4.74 | +0.04 |
| **Ac-Ser-hyp(4-I-Ph)-OMe** | 2.3 | 5.03 | 4.74 | +0.29 |
| **Ac-Ala-hyp(4-I-Ph)-OMe** | 4.2 | 4.78 | 4.74 | +0.04 |

<sup>a</sup> $K_{\text{trans/cis}}$ is the equilibrium constant between *trans*-Pro and *cis*-Pro, calculated as the ratio of the volumes of peaks with *trans*-Pro versus *cis*-Pro.
<sup>b</sup> $\Delta\delta_{\text{cis-trans}} = \delta_{\text{cis}}(\text{Pro H}_\alpha) - \delta_{\text{trans}}(\text{Pro H}_\alpha)$

### Bioinformatics analysis of Ser-Pro structures in the PDB

In order to more broadly understand the structures present at Ser-Pro sequences, we conducted a bioinformatics analysis of high-resolution protein structures in the PDB. The bioinformatics analysis included quantifying distances between relevant heavy atoms that would be consistent with noncovalent interactions, as well as determining key torsion angles associated with different conformations.

In order to identify the presence and quantify the frequency of C–H/O interactions between the Ser side-chain oxygen and proline C–H bonds, the distances between Ser O_γ_ and Pro C_α_ were measured as a function of proline amide conformation (Figure 6). Potential C–H/O interactions were defined as C•••O distances < 3.8 Å (∼ 1.09 Å C–H bond length + 1.20 Å van der Waals radius of H + 1.52 Å van der Waals radius of O). 45% of Ser-*cis*-Pro structures exhibited a Pro/Ser C_α_–H/O_γ_ interaction using this definition of C–H/O interactions. These results, in combination with other data above, strongly suggest that C–H/O interactions directly stabilize the *cis*-proline conformation at Ser–Pro sequences in proteins in water. In contrast, for Ser-*trans*-Pro sequences, essentially no Pro/Ser C_α_–H/O_γ_ interactions were observed, although 28% of all sequences exhibited the possibility of a C–H/O interaction between Ser O_γ_ and Pro C– H_δ_ (Figure S10).

**Figure 6.**
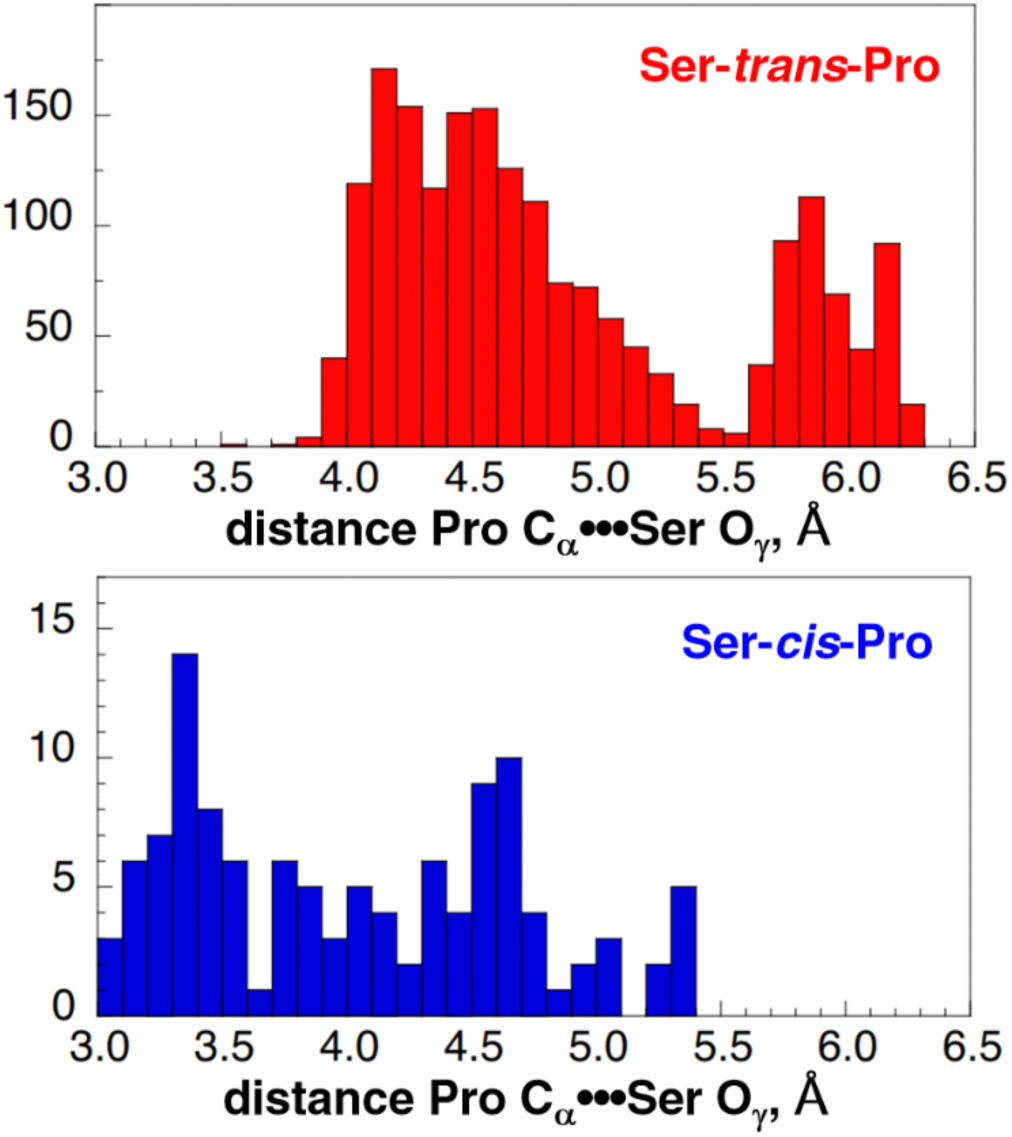
Potential C–H/O interactions in Ser-Pro sequences in the PDB. Ser-Pro sequences were analyzed for the distance between Pro C_α_ and Ser O_γ_, in both Ser-*trans*-Pro (top) and Ser- *cis*-Pro (bottom). The Pro C_α_•••Ser O_γ_ distance was used to identify C_α_–H/O interactions, as few PDB structures have accurate hydrogen positions. A Pro C_α_•••Ser O_γ_ distance ≤ 3.8 Å is consistent with a C_α_–H/O interaction. Using these parameters, 45% of structures containing *cis*- proline have a C_α_–H/O interaction (52 total structures). Only 2 structures containing *trans*-proline were observed with Pro C_α_•••Ser O_γ_ distances < 3.8 Å; both had Ser O_γ_–H•••Pro O hydrogen bonds. The data set includes all crystal structures containing a SP sequence solved with a resolution ≤ 2.0 Å and sequence similarity < 30%. This search resulted in a total of 2045 structures: 1929 structures with *trans*-proline and 116 structures with *cis*-proline.

Analysis of the Ser *χ*_1_ conformations (Figure 7, Table 2) revealed that the *t* rotamer is substantially more populated in Ser-*cis*-Pro structures (60%). The *g*^+^ *χ_1_* conformation was relatively underpopulated for Ser-*cis*-Pro. In contrast, in Ser-*trans*-Pro structures, the three *χ*_1_ rotamers are relatively equally populated, as has been observed previously for Ser.^52,53^ Notably, the *t* rotamer was exclusively observed in structures with C–H/O interactions at Ser-*cis*-Pro, as had also been observed by small-molecule X-ray crystallography (Figure 3).

**Figure 7.**
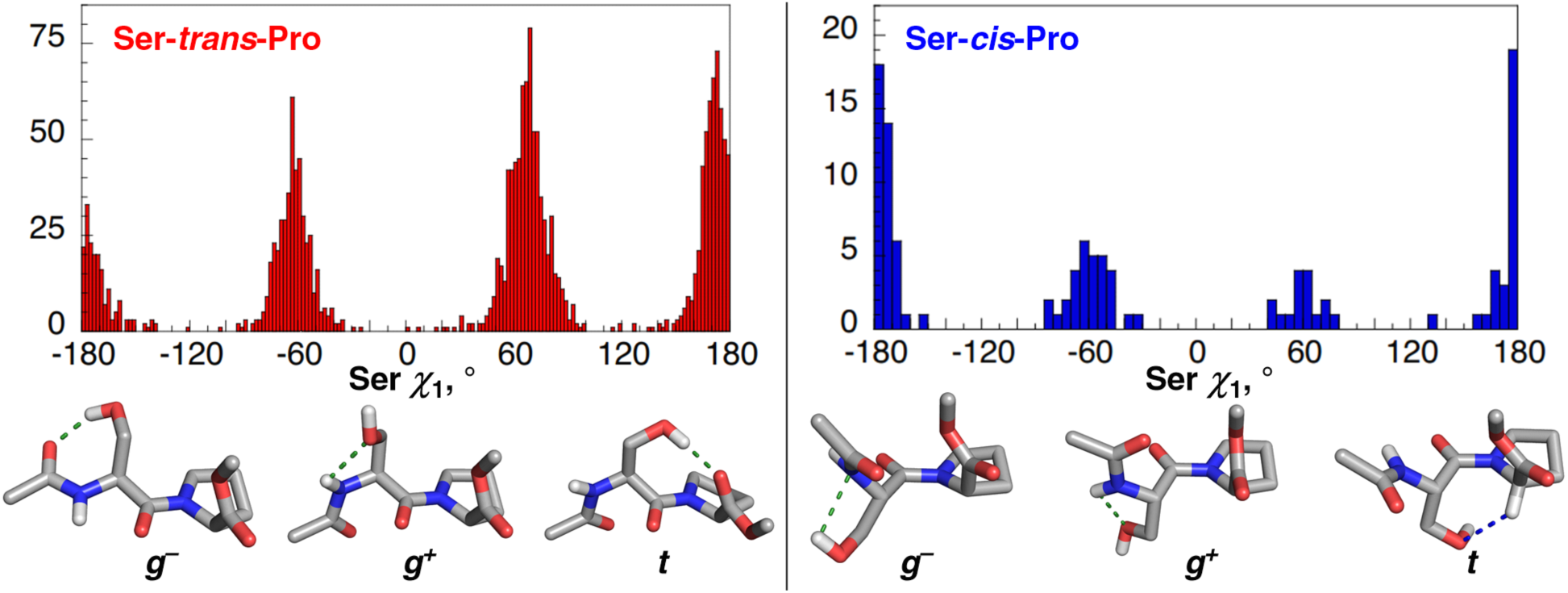
Ser *χ*_1_ rotamers at Ser-Pro sequences in the PDB. Analysis of Ser *χ*_1_ rotamers at (left) Ser-*trans*-Pro and (right) Ser-*cis*-Pro structures in the PDB. In structures containing *trans*- proline, 37% adopt the Ser *χ*_1_ *trans* (*t*) rotamer, 24% adopt the Ser *χ*_1_ *gauche*^−^ (*g^−^*) rotamer and 39% adopt the Ser *χ*_1_ *gauche*^+^ (*g^+^*) rotamer. In structures containing *cis*-proline, 60% adopt the Ser *χ*_1_ *t* rotamer, 28% adopt the Ser *χ*_1_ *g^−^* rotamer, and 12% adopt the Ser *χ*_1_ *g^+^* rotamer. Models of Ac-SP-OMe peptides with examples of interactions observed at each Ser *χ*_1_ rotamer are shown.

**Table 2.** Ser *χ*_1_ rotamer populations at Ser-*trans*-Pro and Ser-*cis*-Pro sequences in the PDB. . Populations of each Ser *χ*_1_ rotamer are represented as a percent of structures in each Pro amide conformation.

| | rotamer | $\chi_1$ | %<sub>total</sub> |
| --- | --- | --- | --- |
| **trans** | $g^-$ | -60 | 24.0 |
| | $g^+$ | +60 | 39.0 |
| | $t$ | $\pm 180$ | 37.0 |
| **cis** | $g^-$ | -60 | 28 |
| | $g^+$ | +60 | 12 |
| | $t$ | $\pm 180$ | 60 |

The Ramachandran plots for Ser-Pro and Ala-Pro sequences were analyzed as a function of proline amide conformation (Figure S9, Table 3). The data on structures with *trans*-proline indicated that, overall, both Ser and Ala populated the PPII (53% *versus* 50%), ζ (8% versus 10%), and α_L_ (1.1% *versus* 1.4%) conformations with similar frequencies. In contrast, Ala was more likely to adopt the α_R_ (14% *versus* 5%) conformation, while Ser was more likely in the β conformation (32% *versus* 23%). At proline, SP sequences were far more likely than AP sequences to have Pro in the α_R_ or δ conformation (65% for SP *versus* 37% for AP). In contrast, AP sequences were much more likely to have Pro in the PPII conformation (60% for AP *versus* 34% for SP).^58–60^

**Table 3.** Conformations of individual residues in Ser-Pro and Ala-Pro sequences in the PDB. . Populations of conformations observed at individual residues in Ser-Pro and Ala-Pro sequences in the PDB. Populations are represented as a percent of structures in each Pro amide isomer. The torsion angle ranges used to define the regions of the Ramachandran plot are in Figure S8 and Table S2.

|  |  | % X at X-Pro |  | % Pro |  |
| --- | --- | --- | --- | --- | --- |
|  | conformation | Ser | Ala | at SP | at AP |
| **trans** | $\alpha_R$ | 4.9 | 14.1 | 52.3 | 32.1 |
| | $\delta$ | 0.1 | 0.1 | 12.3 | 5.1 |
| | $\beta$ | 31.7 | 23.4 | 0.1 | 0.2 |
|  | PPII | 52.6 | 49.6 | 34.1 | 60.0 |
| | $\zeta$ | 8.1 | 9.8 | 0.0 | 0.0 |
| | $\alpha_L$ | 1.1 | 1.4 | n.a.<sup>a</sup> | n.a. |
|  | undefined | 1.4 | 1.5 | 1.2 | 2.6 |
| **cis** | $\alpha_R$ | 0 | 1 | 1 | 2 |
| | $\delta$ | 0 | 1 | 27 | 36 |
| | $\beta$ | 66 | 33 | 1 | 2 |
|  | PPII | 30 | 60 | 67 | 55 |
| | $\zeta$ | 3 | 4 | 1 | 5 |
| | $\alpha_L$ | 1 | 0 | n.a. | n.a. |
|  | undefined | 0 | 1 | 3 | 0 |

<sup>a</sup> n.a. = not applicable. Pro is covalently restrained to negative values of $\phi$ and therefore cannot adopt the $\alpha_L$ conformation.

Analysis of the combinations of conformations of Ser and Pro at SP sites in proteins, compared to those in AP sequences, revealed substantial differences in structures of the dipeptides (Table 4). In structures with *trans*-Pro, AP sequences were significantly more likely to have both residues in the same secondary structure (both α-helix or both PPII). Overall, 47% of AP sequences had the same secondary structure at both residues, while only 20% of SP sequences did. These results are consistent with the high propensities of Ala for both α-helix and PPII.^47–49,61,62^ The observation that 80% of SP sequences had different conformations at Ser and Pro is consistent with SP sequences being particularly prominent in turns, loops, bends, and disordered structures.

**Table 4.** Combinations of conformations observed at Ser-Pro and Ala-Pro sequences. .*^a^* Populations of the most frequent combinations of conformations observed at Ser-Pro and Ala- Pro sequences in the PDB. Populations are represented as a percent of structures in each Pro amide conformation. See Table S4 for complete analysis of all combinations of conformations.

|  | conformation |  | SP | AP |
| --- | --- | --- | --- | --- |
|  | X | Pro | %<sub>total</sub> | %<sub>total</sub> |
| **trans** | $\alpha_R$ | $\alpha_R$ | 4.2 | 12.8 |
| | $\beta$ | $\alpha_R$ | 13.8 | 3.2 |
| | $\beta$ | $\delta$ | 4.8 | 1.0 |
| | $\beta$ | PPII | 12.2 | 18.0 |
| | PPII | $\alpha_R$ | 30.2 | 11.5 |
| | PPII | $\delta$ | 6.1 | 2.2 |
|  | PPII | PPII | 15.9 | 34.3 |
| **cis** | $\beta$ | $\delta$ | 9 | 3 |
| | $\beta$ | PPII | 55 | 28 |
| | PPII | $\delta$ | 17 | 32 |
|  | PPII | PPII | 9 | 23 |

<sup>a</sup> Populations of combinations of conformations that are either present at > 5% of the total population or that are different between SP and AP by greater than a factor of 1.5 are shown.

In structures with *cis*-Pro (Table 3), the β conformation was substantially more likely for Ser than Ala (66% *versus* 33%), while Ala was far more likely to adopt PPII (60% *versus* 30%). The geometric constraints of *cis*-proline inherently cause the vast majority of structures with *cis*- proline to be β-turns, with Cα•••Cα *i*/*i*+3 distances ≤ 7.0 Å (92% of Ser-*cis*-Pro, 78% of Ala-*cis*- Pro).^15,35^ Type VI β-turns include both backbone-hydrogen-bonded (type VIa1, PcisD) and non-backbone-hydrogen-bonded types (others). SP sequences appear particularly likely to adopt type VIb (BcisP) β-turns (55% of SP *versus* 28% of AP).

We examined the role of C–H/O interactions as a function of type VI β-turn subtype in Ser-*cis*-Pro structures (Table 5, Figure 8). The majority of structures in type VIa1 (PcisD), VIa2 (BcisD), and PcisP (one type of VIb) β-turns exhibited C–H/O interactions crystallographically between the Ser O_γ_ and Pro C–H_α_. C–H/O interactions were mediated exclusively via the *t χ*_1_ rotamer (Figure 8a), but could be observed in different secondary structures of both Ser and Pro (Table 5). In contrast, BcisP type VIb β-turns, while the most commonly observed, exhibited the lowest frequency of C–H/O interactions (21% of structures).

**Figure 8.**
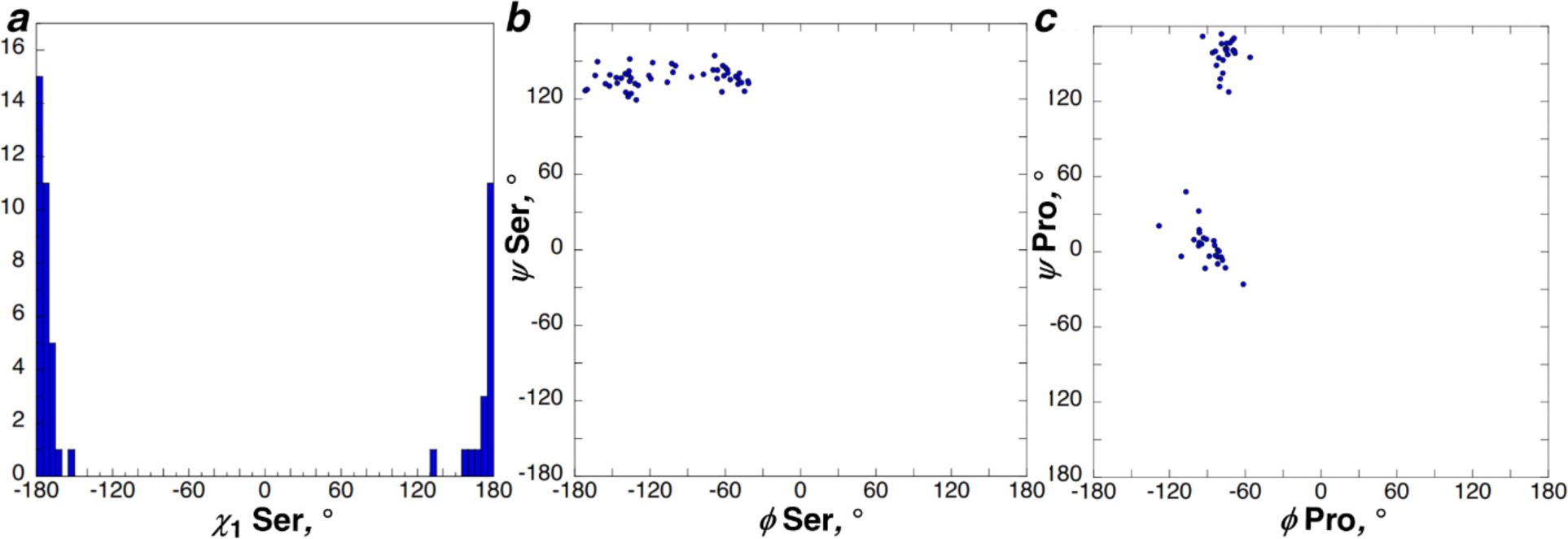
Structures at Ser-*cis*-Pro sequences with C–H/O interactions. (a) Ser *χ*_1_ rotamer distribution and (b, c) Ramachandran plots of (b) Ser and (c) Pro in structures with a C_α_–H•••O interaction.

**Table 5.** Frequencies of C–H/O interactions at Ser-*cis*-Pro conformations in different type VI b-turns. . Frequencies of C–H/O interactions observed at the most common conformations observed at Ser-*cis*-Pro structures. See Table S5 for complete analysis.

| **cis** | conformation | | β-turn type | | %<sub>total</sub> | % Cα–H/O |
| --- | --- | --- | --- | --- | --- | --- |
|  | Ser | Pro | classical | Dunbrack |  |  |
| | PPII | $\delta$ | Vla1 | PcisD | 17 | 65 |
| | $\beta$ | $\delta$ | Vla2 | BcisD | 9 | 70 |
|  | PPII | PPII | Vlb | PcisP | 9 | 55 |
| | $\beta$ | PPII | Vlb | BcisP | 55 | 31 |

The presence of hydrogen-bonded β-turns was examined for *trans*-Pro and *cis*-Pro structures, via quantifying the O*_i_*•••H–N*_i_*_+3_ distance, with distances ≤ 2.7 Å indicating possible hydrogen-bonded β-turns (Table 6, Figure S12, Figure S14). This hydrogen bond pattern is also present in 3_10_-helices and at the N-terminus of α-helices. Potential β-turns were examined with two different registers: with Ser or Ala as the *i* residue of the β-turn (Pro–X central residues of the β-turn; Ser*_i_*/Ala*_i_*–Pro-X–X*_i_*_+3_ register), or with Ser or Ala as the *i*+1 residue of the β-turn (Ser-Pro or Ala-Pro central residues of the β-turn; X*_i_*–Ser*_i_*_+1_/Ala*_i_*_+1_–Pro–X*_i_*_+3_ register). In registers with Ser–Pro or Ala–Pro as the central residues, 14% of *trans*-Pro structures with Ala as the *i*+1 residue were hydrogen-bonded β-turns or related hydrogen-bonded structures, while only 5% of structures were with Ser. For *cis*-Pro structures with this register, 24% and 37% of structures were hydrogen-bonded for Ser and Ala, respectively.

**Table 6.** Frequencies of main-chain *i*/*i*+3 C=O*_i_*•••H–N*_i_*_+3_ and *i*/*i*+4 C=O*_i_*•••H–N*_i_*_+4_ hydrogen bonds at Ser-Pro and Ala-Pro sequences, with Ser/Ala as the *i*+1 residue (X*_i_*–(Ser/Ala)-Pro– X*_i_*_+3_ register) or as the *i* residue (Ser*_i_*/Ala*_i_*–Pro-X–X*_i_*_+3_ register). Frequencies of C=O*_i_*•••H–N*_i_*_+3_ and C=O*_i_*•••H–N*_i_*_+4_ backbone hydrogen bonds in each register for Ser and Ala.

| register | %total<br>$\text{C}=\text{O}_i\cdots\text{H}-\text{N}_{i+3}$ | %total<br>$\text{C}=\text{O}_i\cdots\text{H}-\text{N}_{i+4}$ | %total<br>both<sup>a</sup> | %total<br>any<sup>b</sup> |
| --- | --- | --- | --- | --- |
| **X-S-*trans*-P-X** | 3.6 | 3.1 | 2.0 | 4.7 |
| **X-A-*trans*-P-X** | 10.5 | 9.4 | 6.1 | 13.8 |
| **X-S-*cis*-P-X** | 18 | 8 | 2 | 24 |
| **X-A-*cis*-P-X** | 27 | 16 | 6 | 37 |
| **S-*trans*-P-X-X** | 42.9 | 25.2 | 16.9 | 51.2 |
| **A-*trans*-P-X-X** | 27.1 | 14.3 | 7.5 | 33.9 |
| **S-*cis*-P-X-X** | 0 | 0 | 0 | 0 |
| **A-*cis*-P-X-X** | 0 | 0 | 0 | 0 |

<sup>a</sup> Percent of structures with both $\text{C}=\text{O}_i\cdots\text{H}-\text{N}_{i+3}$ and $\text{C}=\text{O}_i\cdots\text{H}-\text{N}_{i+4}$ hydrogen bonds.
<sup>b</sup> Percent of structures with any $\text{C}=\text{O}_i\cdots\text{H}-\text{N}_{i+3}$ or $\text{C}=\text{O}_i\cdots\text{H}-\text{N}_{i+4}$ hydrogen bond, including structures with both hydrogen bonds.

In contrast, in the register with Pro–X as the central residues, 51% of structures with Ser as the *i* residue had *i*/*i*+3 and/or *i*/*i*+4 hydrogen bonding patterns, while only 34% of those with Ala were. Among these hydrogen-bonded structures at SP, 43% are at the N-terminus of or within α-helices or 3_10_-helices (22% of all Ser-*trans*-Pro sequences), with the other 49% representing β-turns (25% of all Ser-*trans*-Pro sequences) (Table 7). In contrast, no structures with *cis*-Pro in this register exhibited hydrogen-bonded β-turns. Collectively, these data suggest that Ser–Pro sequences most frequently adopt β-turns with Ser as the *i* residue and with Pro–X as the central residues, and are significantly less likely to adopt β-turns with Ser-Pro as the central residues.

**Table 7.** Most frequent hydrogen-bonded secondary structures present at Ser-*trans*-Pro and Ala-*trans*-Pro sequences in the PDB. . Helix initiating structures were identified by a sequence of five or more consecutive residues adopting the α_R_/δ conformation following the first residue of the sequence.

| secondary structure | %total **S-*trans*-P-X-X** | %total **A-*trans*-P-X-X** |
| --- | --- | --- |
| β-turn | 25.2 | 18.0 |
| $3_{10}$-helix or α-helix initiation | 22.1 | 13.6 |
| α-turn | 3.9 | 2.3 |

β-Turn types that can accommodate Pro as the *i*+1 residue include types I (AD), II (Pd or Pa), and VIII (AB1, AZ, AG). Further analysis of these data indicated that 86% of all β-turns in Ser-*trans*-Pro sequences with the Pro–X register were type I (AD) β-turns (Table 8). In type I β-turns, the Ser O_γ_ lone pair can hydrogen bond to the *i*+2 amide hydrogen (O_γ_•••H–N*_i_*_+2_), which is mediated by the *t χ_1_* rotamer.^25–28^ This hydrogen-bonding pattern was observed frequently in the PDB (Table 9), and results in intramolecular solvation of all amide hydrogens of the β-turn (Figure S5).

**Table 8.**
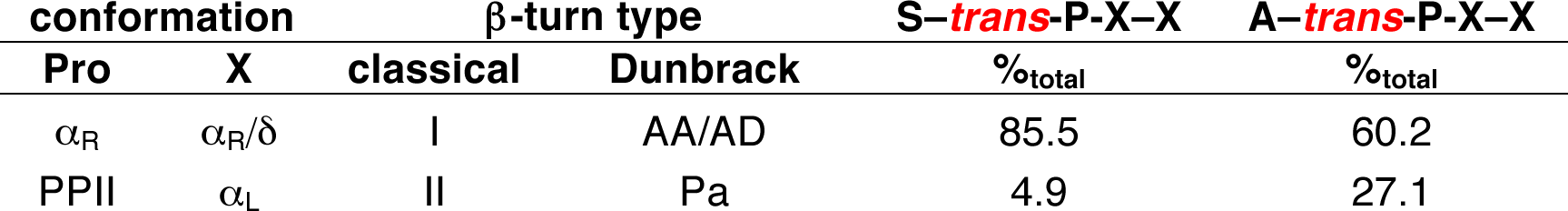
Most frequent β-turn types at S–*trans*-P-X–X and A–*trans*-P-X–X sequences. . 36.8% of S–*trans*-P-X–X and 18.2% of A–*trans*-P-X–X sequences in the PDB were found in to be in β-turns (Table 7). Among the structures in the β-turn conformation, populations of the most common types of β-turn observed at P-*trans*-X sequences are indicated. See Table S10 for full analysis.

**Table 9.** Frequencies of Ser*_i_*O_γ_•••H–N*_i_*_+2_ and Ser*_i_*O_γ_•••H–N*_i_*_+3_ side chain-main chain hydrogen bonds with Ser as the *i* residue in type I β-turns and as the N–capping residue of 3_10_-helices and α-helices. Main-chain hydrogen bonded secondary structures in the PDB with Ser as the *i* residue and Pro as the *i*+1 residue that have Ser*_i_*O_γ_•••H–N*_i_*_+2_ and/or Ser*_i_* O_γ_•••H–N*_i_*_+3_ hydrogen bonds.

| S-*trans*-P-X-X | Ser<sub>i</sub> O<sub>γ</sub> hydrogen bonds |  |  |  |
| --- | --- | --- | --- | --- |
|  | % Ser<sub>i</sub> O<sub>γ</sub>...H-N<sub>i+2</sub> | % Ser<sub>i</sub> O<sub>γ</sub>...H-N<sub>i+3</sub> | % both | % any |
| type I β-turn | 50.7 | 6.3 | 2.6 | 54.4 |
| 3<sub>10</sub>-helix Ncap | 39.8 | 31.4 | 3.4 | 67.8 |
| α-helix Ncap | 9.7 | 76.5 | 6.7 | 79.5 |

In SP structures in 3_10_-helices and α-helices, the Ser residue commonly functions as the helix capping residue (Ncap). The Ncap residue (*i*) is not in the 3_10_- or α-helical conformation, but provides the carbonyl that is hydrogen-bonded to the *i*+3 (3_10_-helix) or *i*+4 (α-helix) amide N–H, which represents the first main-chain hydrogen bond of the helix (Figure S5).^63–65^ 3 - Helices and α-helices without Pro have 2 or 3, respectively, unsatisfied amide N–H hydrogen-bond donors at the N-terminus of the helix. The high frequency of Ser residues at the Ncap position (Table 7) is due to the ability of the Ser O_γ_ to function as a hydrogen-bond acceptor to the *i*+2 or *i*+3 amide N–H (Table 9).^59,66–69^ Therefore, because Pro lacks an amide hydrogen, 3 - helices and α-helices with an SP sequence at the N-terminus, and with the Ser O_γ_ functioning as a hydrogen-bonding amide capping group, only have 0 or 1 (respectively) solvent-exposed amide hydrogen at the helical N-termini. When functioning as a helix cap, O_γ_•••H–N*_i_*_+2_ hydrogen bonds are associated with Ser in the *t χ*_1_ rotamer, while O_γ_•••H–N*_i_*_+3_ hydrogen bonds are associated with Ser in the *g*^+^ *χ_1_* rotamer.

### DFT calculations on combinations of conformations (conformational poses) observed at Ser–Pro sequences

In the bioinformatics work, we identified the most frequent sets of conformations observed at these residues, with a particular focus on structures that involve interactions with the Ser side-chain hydroxyl. Geometry optimization calculations were conducted using DFT methods on minimal peptides in order to better understand the roles of Ser noncovalent interactions in stabilizing specific conformations.

For structures with *cis*-Pro, C–H/O interactions were found to be local energy minima in all subtypes of type VI β-turns (Figure 9). For type VI β-turns with Ser in either the PPII conformation (type VIa1 PcisD, type VIb PcisP) or in the β conformation (type VIa2 BcisD, type VIb BcisP), C–H/O interactions were observed with a *t χ*_1_ conformation at Ser. Notably, C–H/O interactions were energy minima with Pro in either the PPII or α_R_/δ conformation, which are the two predominant energy minima of Pro in Ramachandran space. Overall, 4 different combinations of main-chain (Ser at PPII and β, Pro at PPII and α_R_/δ) and side-chain (*t χ*_1_) conformations exhibited close C–H/O interactions, consistent with bioinformatics data.

**Figure 9.**
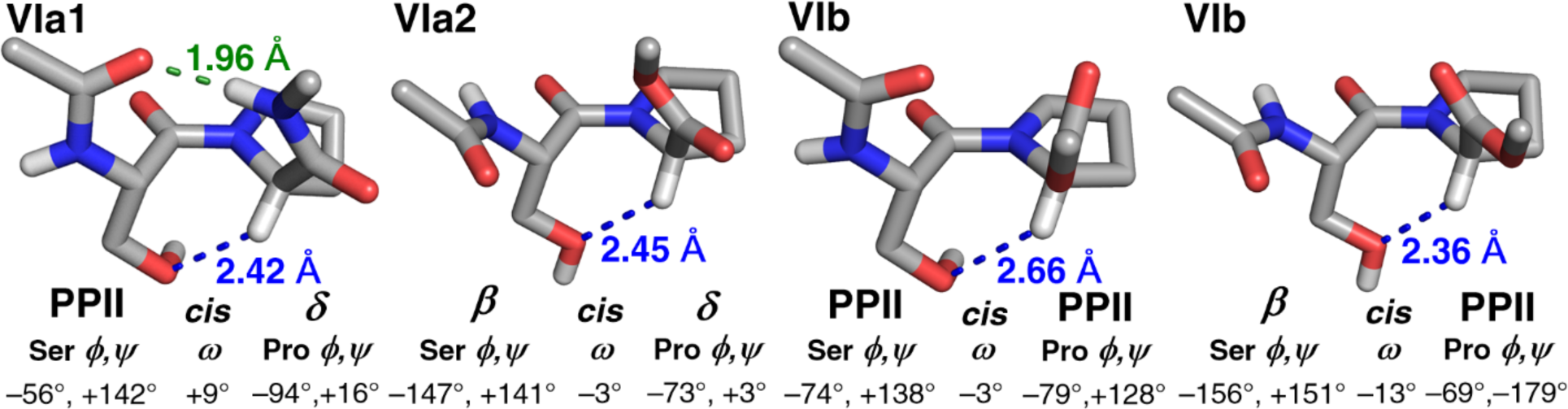
Structures of type VI β-turns stabilized via C–H/O interactions. Structures of (type VIa1) Ac-Ser-Pro-NHMe and (others) Ac-Ser-Pro-OMe type VI β-turns with C–H/O interactions (blue) between Ser O_γ_ and Pro C–H_α_ that stabilize the *cis*-Pro conformation in Ser-*cis*-Pro. These structures were derived from those identified via bioinformatics, then subjected geometry optimization by DFT methods. All C–H/O interactions are mediated via the Ser *t χ*_1_ rotamer. Type VIa1 β-turns are mediated via *i*/*i*+3 main-chain O*_i_*•••H–N*_i_*_+3_ hydrogen bonds (green), while other type VI β-turn subtypes are not.

Hydrogen bonds with the Ser hydroxyl were observed in other structures, in both *trans*- Pro and *cis*-Pro (Figure S4). In the *g^−^*and *g^+^ χ_1_* conformations, the Ser hydroxyl can serve as a hydrogen bond donor with the carbonyl of the prior residue or of Ser, respectively. In the *g^−^* or *g^+^χ*_1_ conformations, Ser O_γ_ can also function as a hydrogen-bond acceptor to the Ser amide N–H. O–H/N interactions with the Ser amide N are also possible in the *g^+^* conformation. In structures with *trans*-Pro, hydrogen bonds of the Ser O–H with the Pro carbonyl were observed in the *t χ*_1_ conformation.

Structures of SP sequences in type I β-turns (Ac-S–PA–NHMe), with Ser as the *i* residue, demonstrated the specific stabilization provided by Ser O_γ_ serving as a hydrogen-bond acceptor to the *i*+2 residue amide N–H (Figure 10). With Ser as the Ncap residue of a 3_10_-helix, and with Ser in either the PPII or β conformation, a hydrogen bond between Ser O_γ_ and the *i*+2 residue amide N–H resulted in there being no solvent-exposed amide N–H hydrogens at the N-terminus of the 3_10_-helix (Figure 11). This hydrogen bond is mediated via the *t χ*_1_ rotamer. In contrast, in an α-helix stabilized by a Ser*_i_* O_γ_•••H–N*_i_*_+3_ hydrogen bond, this capping interaction occurs via the *g^+^χ_1_* rotamer (Figure 12). Collectively, these computational results demonstrate a multitude of ways that the Ser hydroxyl can stabilize local structures in SP sequences.

**Figure 10.**
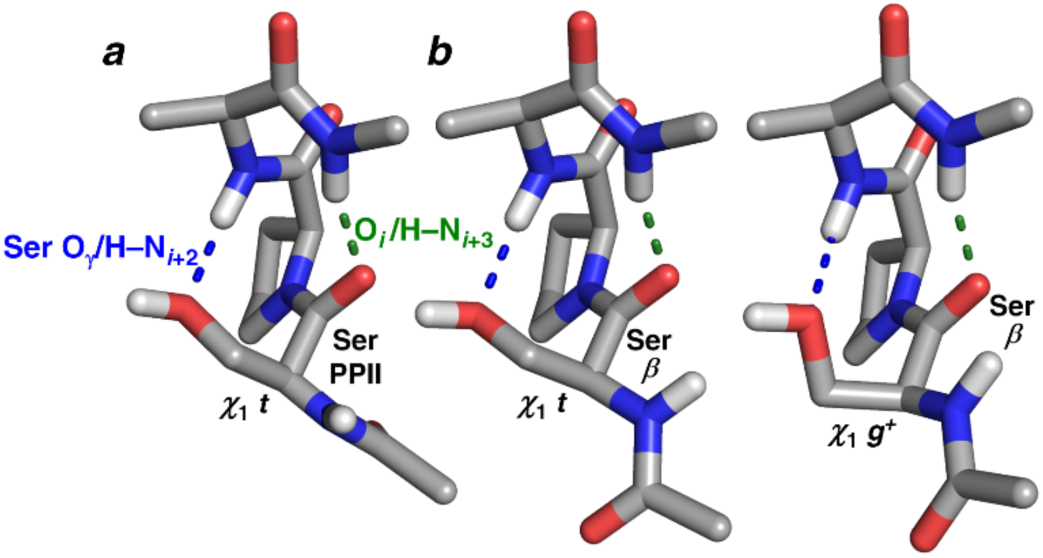
Stabilization of type I β-turns by Ser*_i_*O_γ_•••H–N*_i_*_+2_ hydrogen bonds. (a,b) Geometry-optimized structures of Ac-Ser*_i_*–Pro*_i_*_+1_-Ala*_i_*_+2_–N*_i_*_+3_HMe type I β-turns (mediated by a main-chain *i*/*i*+3 C*_i_*=O••••H–N*_i_*_+3_ hydrogen bond, green) in which the Ser O_γ_ engages in a hydrogen bond (blue) with the amide N–H of the *i*+2 residue of the turn (here, Ala). The combination of Pro as the *i*+1 residue and the Ser hydrogen bond to the *i*+2 residue amide results in no solvent-exposed amide hydrogens within the β-turn. Ser is the *i* residue of the turn, with Ser in the (a) PPII or (b) β conformation. The side chain-main chain hydrogen bond is mediated (a) via the *t χ*_1_ rotamer when Ser adopts the PPII conformation or (b) via either the (left) *t* or (right) *g*^+^ *χ*_1_ rotamer when Ser is in the β conformation. The type I β-turn requires a *trans*-proline amide (Ser-*trans*-Pro).

**Figure 11.**
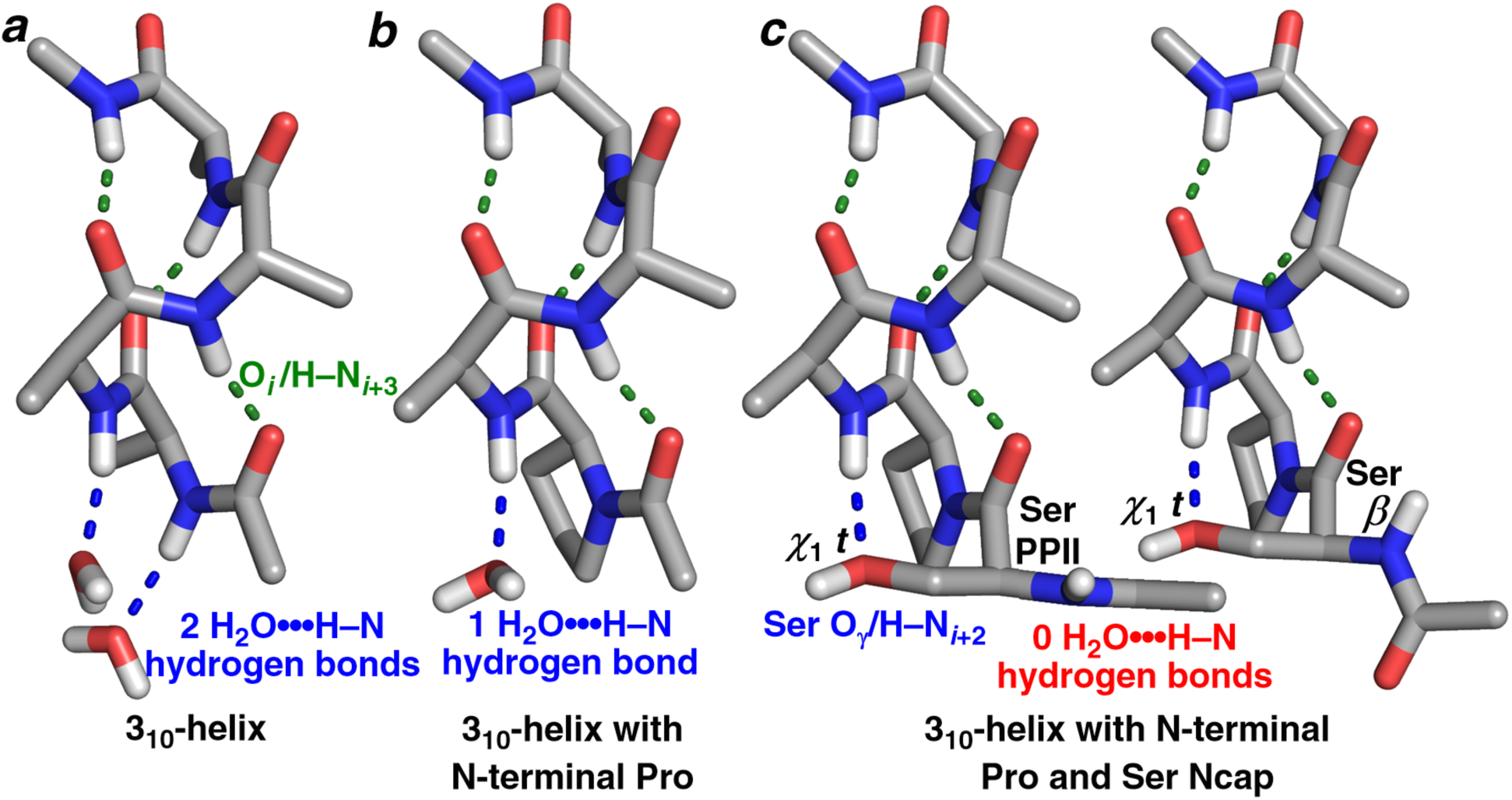
Solvation at the N-termini of 3_10_-helices, including 3_10_-helix capping by a Ser O_γ_•••H–N*_i_*_+2_ hydrogen bond. Geometry-optimized structures of (a) Ac-Ala_4_-NHMe, (b) Ac-Pro- Ala_3_-NHMe, and (c) Ac-Ser-Pro-Ala_3_-NHMe, with Pro and Ala residues in the 3_10_-helix conformation. In (c), Ser functions as the Ncap residue (*i*) of the 3_10_-helix, with a Ser O_γ_ hydrogen bond with the amide N–H of the *i*+2 residue, and with Ser in either the (left) PPII or (right) β conformation. This Ser O_γ_•••H–N*_i_*_+2_ side chain-main chain hydrogen bond is mediated via Ser in the *t χ*_1_ rotamer. In (a), the N-terminus of the 3_10_-helix has 2 unsatisfied amide hydrogen-bond donors which must be solvated. When (b) the N-terminal residue of the 3_10_-helix is Pro, only one unsatisfied amide hydrogen-bond donor is present. In 3_10_-helices with Ser as the Ncap (*i*) residue and Pro as the first (*i*+1) residue, there are no N-terminal amide hydrogens that must be solvated, as a result of the helix-capping Ser O_γ_•••H–N*_i_*_+2_ hydrogen bond. A similar example of a capped 3_10_-helix stabilized via a SP sequence is in Figure S5. Geometry optimization was conducted using the M06-2X DFT functional, the 6-311++G(d,p) basis set, and implicit water solvation.

**Figure 12.**
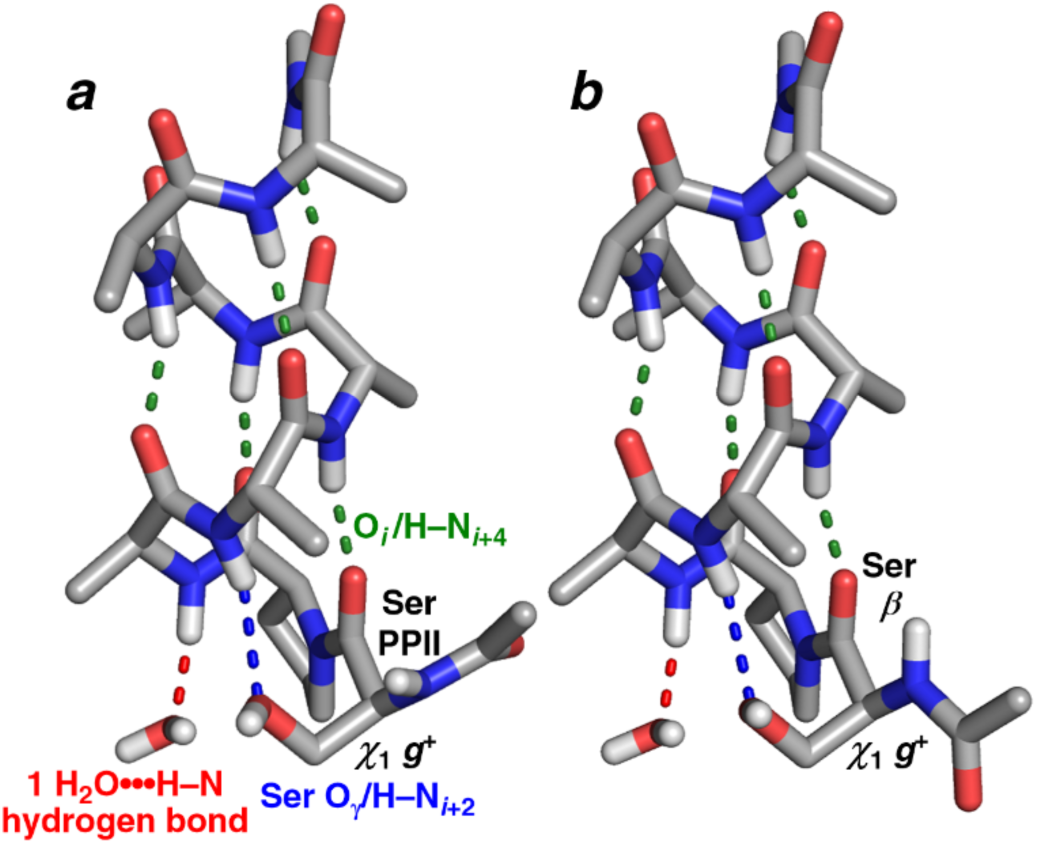
N-Capping of α-helices in Ser*_i_*-*trans*-Pro*_i_*_+1_ sequences via a Ser O_γ_•••H–N*_i_*_+3_ hydrogen bond. Geometry-optimized structures of Ac-Ser-*trans*-Pro-Ala_6_-NHMe, with Pro and Ala residues in the α-helix conformation, with the N-terminal Ser in (a) the PPII conformation or (b) the β conformation. In both structures, a Ser O_γ_•••H–N*_i_*_+3_ hydrogen bond, mediated via the Ser *g*^+^ *χ_1_* rotamer, results in only one unsatisfied N-terminal amide hydrogen-bond donor (at N2). In contrast, an uncapped α-helix without an N-terminal Pro has three unsatisfied amide N–H hydrogen-bond donors.

## Discussion

Herein, we conducted a comprehensive analysis of structures stabilized at Ser–Pro sequences in proteins, using a combination of small-molecule X-ray crystallography, solution NMR spectroscopy, bioinformatics analysis of proteins in the PDB, and DFT calculations. SP sequences are particularly prone to adopt local structures stabilized by hydrogen bonds or by C– H/O interactions.

### Ser-trans-Pro sequences stabilize type I β-turns

51% of SP sequences in the PDB with *trans*-Pro exhibit a hydrogen bond between the Ser (*i* residue) C=O and the amide N–H of the *i*+3 and/or *i*+4 residue. These hydrogen-bonding patterns are present in β-turns, 3_10_-helices, and α-helices. β-turns were observed at 25% of all Ser-*trans*-Pro sequences with Ser as the *i* residue and Pro as the *i*+1 residue, with the vast majority type I β-turns. Notably, the first turn of 3_10_-helices and α-helices has the same conformation at the *i*+1 and *i*+2 residues (α_R_/δ) as a type I β-turn.

The key interaction of Ser that specifically stabilizes a type I β-turn is a hydrogen bond between the Ser Oγ and the amide N–H of the *i*+2 residue (Figure 10). This interaction has previously been identified to stabilize type I β-turns.^25,27,28^ More broadly, residues with the hydrogen bond acceptors Asp, Asn, and Ser are substantially overrepresented at the *i* position of type I β-turns, and Pro is by far the most frequent residue at the *i*+1 position of type I β-turns.^25,26^ A hydrogen bond between an oxygen of the *i* residue side chain and the amide N–H of the *i*+2 residue inherently stabilizes the type I β-turn conformation, via intra-turn solvation of that amide, which prevents competing hydrogen-bonding interactions at that amide that would otherwise lead to different structures. While the specific hydrogen-bonding motifs and residue frequencies at individual positions in type I β-turns have been previously identified, the bioinformatics data herein represent a far larger data set, with a singular focus on SP sequences. These data strongly suggest that type I β-turns should be a major conformation at SP sequences in proteins, including in intrinsically disordered proteins.

### Ser-Pro sequences stabilize the N-termini of 3_10_-helices and α-helices

Similarly, in this analysis, we found that 22% of all Ser-*trans*-Pro sequences in the PDB represent helical capping motifs. The roles of both residues at helical N-termini have previously been extensively studied. Ser is well-recognized to function as a common hydrogen-bond acceptor in α-helical N-terminal capping motifs.^65,67,68,70^ Proline is also a common residue at the N-termini of α-helices, due to Pro not having an amide hydrogen. Amide hydrogens are solvent-exposed in the first 3 residues of the α-helix and thus does not participate in intrahelical hydrogen bonds.^2,64,70–72^ The combination of Ser at the Ncap position of an α-helix, with the Ser O_γ_ hydrogen-bonded to the N3 amide N–H, and Pro at the N1 position, results in only one solvent-exposed amide N–H at the α-helical N-terminus (at residue N2) (Figure 12). Hydrogen bonding of a side-chain hydrogen-bond acceptor to the N3 amide N–H might be particularly favorable, due to the relatively weak nature of amide-water N–H•••OH hydrogen bonds at the N3 amide.^68,73,74^

3_10_-Helices are common in short helical structures, due to the *i*/*i*+3 register of hydrogen bonds yielding a larger total number of intrahelical hydrogen bonds in a short sequence versus those with an α-helix.^63,71,75^ Mechanistic studies of protein folding also suggest that 3 -helices are intermediates in the formation of α-helices, as they require the organization of fewer residues to form a helical turn.^76–78^ The data herein suggest that SP sequences might promote the 3 -helix to α-helix conversion in protein folding. 3_10_-Helices are stabilized in SP sequences by a Ser O_γ_•••H–N*_i_*_+2_ hydrogen bond, which is mediated by Ser in the *t χ*_1_ rotamer (Figure 11). Rotation of Ser to the *g^+^* rotamer promotes instead a hydrogen bond with the N3 amide N–H (Figure 12), as is observed in α-helical capping motifs. Thus, simple rotation about Ser *χ*_1_ at SP sequences might favor a change in structure from 3_10_-helix to α-helix, thereby promoting adoption of native protein structure at α-helices.

Both in β-turns and in helical capping motifs, Ser O_γ_ can stabilize structures via hydrogen bonds, to either the *i*+2 or *i*+3 amide hydrogen, when Ser is in either the PPII or β main-chain conformation. The ability to stabilize secondary structures via either PPII or β is consistent with the high frequencies of SP sequences with Ser in either the PPII or β conformation and with Pro in the α_R_ or δ conformation (Table 4). Even without an intramolecular hydrogen bond, the crystal structure of Ac-hyp(4-I-Ph)-OMe (Figure 5) exhibited the same combinations of conformations (Ser in PPII or β, Pro in α_R_/δ) as are present in β-turn and helical capping motifs, consistent with these structures being strongly favored in SP sequences.

### Changes in structure at SP sequences due to proline cis-trans isomerism

The vast majority of SP sequences with *cis*-Pro adopt a type VI β-turn structure. β-Turns at Ser-*cis*-Pro are centered on SP (Ser at the *i*+1 residue, Pro at the *i*+2 residue). In contrast, β-turns at Ser- *trans*-Pro are predominantly centered on Pro-X (Ser at the *i* residue, Pro at the *i*+1 residue). Proline *cis*-*trans* isomerization inherently generates large changes in protein structure.^3,4,79–81^ At SP sequences, the Ser-*trans*-Pro to Ser-*cis*-Pro amide conformational change both induces a β-turn and switches the register of any β-turn present, and thus can change protein structure even if a β-turn is present with *trans*-Pro. In addition, we recently proposed that pSer-Pro sequences can promote β-turns with pSer the *i*+1 residue and Pro the *i*+2 residue,^16^ and thus Ser phosphorylation at SP sequences could also change structure in part via changing both the presence and the register of β-turns in proteins.

The RNA Polymerase II C-terminal domain in eukaryotes consists of consensus repeats of the YSPTSPS sequence, with the number of repeats and identities of non-consensus residues differing between organisms.^7,82^ Humans have 52 such repeats, with each consensus repeat containing two SP sequences (one SPT and one SPS). Resides 5-6 of the repeats are nearly perfectly conserved in humans as SP sequences (51 of 52), while at residues 2-3, 10 SP sequences are replaced by TP, one by EP, and one by SL. Bioinformatics analysis of proteins (by others and by us) and DFT calculations found that the SPS and SPT sequences are particularly prone to stabilization of type I β-turns, with the Ser O_γ_•••H–N*_i_*_+2_ hydrogen bond further stabilized by a hydrogen bond between Ser O –H and Ser/Thr O (Figure 13).^6,25–28,83–85^ This pattern of hydrogen bonds is geometrically favorable with Ser*_i_* in either the PPII or β conformation, and with Pro in either the *endo* or *exo* ring pucker. The Pol II CTD has previously been identified to be conformationally heterogeneous, with a significant propensity to adopt β-turn conformations.^9,83,86–89^ The data herein further confirm the favorability of type I β-turns within the Pol II CTD. Moreover, phosphorylation of Ser2, Thr4, and/or Ser5 of the Pol II CTD is central to changes in transcriptional elongation and termination, providing a “phosphorylation code” for transcription.^7,8^ The data herein provide further context for considering the nature of the transcriptional code of structural changes in the Pol II CTD.

**Figure 13.**
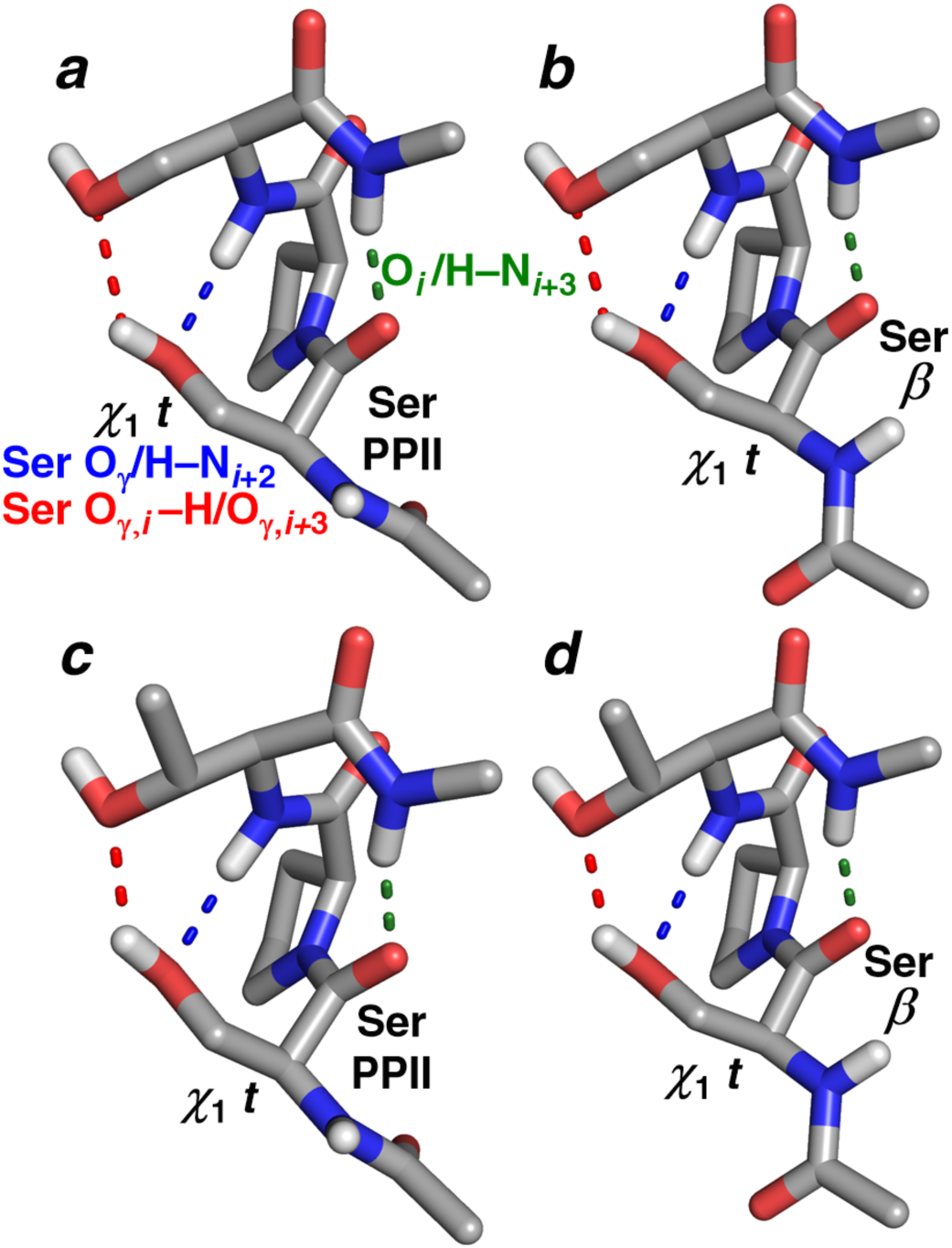
Type I β-turns in SPS and SPT sequences stabilized via simultaneous Ser*_i_* O_γ_•••H–N*_i_*_+2_ and Ser*_i_* O_γ_–H•••O_γ_ Ser*_i_*_+2_ hydrogen bonds. (a–d) Optimized structures of (a,b) Ac-Ser*_i_*–Pro*_i_*_+1_-Ser*_i_*_+2_–N*_i_*_+3_HMe or (c,d) Ac-Ser*_i_*–Pro*_i_*_+1_-Thr*_i_*_+2_–N*_i_*_+3_HMe in a type I β-turn mediated by a main-chain *i*/*i*+3 hydrogen bond (green). Ser*_i_* O_γ_ simultaneously acts as a hydrogen-bond acceptor to the main-chain amide H–N*_i_*_+2_ (blue) and as a hydrogen-bond donor to the Ser*_i_*_+2_ or Thr*_i_*_+2_ side chain O_γ_ (red). This dual hydrogen-bonding interaction of the Ser*_i_*side chain is mediated by Ser in the *t χ*_1_ rotamer, with Ser in either (a,c) the PPII conformation or (b,d) the β conformation.

### The cis-Proline conformation is stabilized by C–H/O interactions in Ser-cis-Pro sequences

Herein, we identified that *cis*-Pro can be stabilized at SP sequences via a C–H/O interaction between Ser Oγ and Pro C–H_α_. We observed evidence of this interaction in the solid state by small-molecule X-ray crystallography; in solution via an increase in the population of *cis*-Pro in SP dipeptides versus AP dipeptides and via a downfield change in the Pro Hα chemical shift; via bioinformatics analysis of Ser-*cis*-Pro structures in the PDB; and via DFT calculations. C–H/O interactions are inherently favorable at the polarized C–H bonds of proline.^17^ We observed that C–H/O interactions can stabilize all type VI β-turn subtypes. Notably, the crystal structure of the marine natural product cyclic peptide stylopeptide 1 (CSD ZOHMIS) exhibits Ser-*cis*-Pro in a type VIa1 β-turn (PcisD) conformation, with a close (2.57 Å) Pro C–H_α_•••O_γ_ Ser C–H/O interaction that appears to stabilize the *cis*-proline conformation, which is also present in solution.^90^

Thus, while C–H/O interactions mediated by a neutral Ser O_γ_ lone pair are inherently weaker^20,91^ than those of anionic Glu-*cis*-Pro and pSer-*cis*-Pro, the data indicate that they are still significant in peptides and proteins in water. While C–H/O interactions are typically described in electrostatic terms (interaction of the δ^+^ on a polarized C–H bond with the δ^−^ on O), they are predominantly stereoelectronic (molecular orbital-based) in nature, and exhibit only a weak dependence of interaction strength on solvent dielectric constant.^17^ The stereoelectronic nature of C–H/O interactions can explain the common observation of C–H/O interactions stabilizing diverse protein structures.^22–24,92^

### C–H/O interactions can stabilize cis-non-proline amide bonds

Herein, we identified that C–H/O interactions between a Ser side-chain O and Pro C–H_α_ are commonly observed and stabilizing to the *cis*-Pro conformation. *cis* amide bonds are also observed at 0.03% of non-proline residues (versus at 5% of proline residues).^13,15,79,93^ While *cis* amide bonds are much less likely at non-proline residues than at Pro for any given amide bond, over 95% of all amide bonds in proteins are at non-proline residues. Thus, while *cis*-Pro represents approximately 0.25% of all amide bonds in proteins (5% × 5%), *cis*-non-Pro represents approximately 0.03% of all amide bonds in proteins (95% × 0.03%), or 1 out of every ∼3000 amide bonds. As such, the overall frequency of *cis*-non-Pro amide bonds is substantial, representing ∼10% of all *cis* amide bonds in proteins. Stating the overall frequency of *cis*-non-Pro amide bonds differently, approximately 10% of all 300 amino-acid proteins or protein domains would be expected to contain at least one *cis*-non-Pro amide bond. Because *cis*-non-Pro amide bonds are substantially higher in energy than *trans*-non-Pro amide bonds (the 0.03% frequency suggests that *cis*-non-Pro is approximately 5 kcal mol^−1^ higher in energy than *trans*-non-Pro), any observed *cis*-non-Pro amide bond needs to be substantially stabilized by other interactions within the protein.^94,95^

In addition, *cis-trans* isomerization from *trans*-non-Pro to *cis*-non-Pro is characterized by extremely slow kinetics (*t*_1/2_ ∼ 1000 s), with rapid reversion from *cis*-non-Pro to *trans*-non-Pro (*t* < 1 s) in simple peptides.^96,97^ These kinetics and thermodynamics suggest a significant role for local structures in stabilizing observed *cis*-non-Pro amide bonds.^97–100^ The rate of *cis*-*trans* amide interconversion at non-*cis*-Pro amide bonds is accelerated by molecular chaperones such as the *E. coli* Hsp70 DnaK, which functions as a non-prolyl amide isomerase.^101^ Notably, *cis*- non-Pro amide bonds are structurally highly conserved, and are usually present at or near functional sites in proteins.^93,102–107^ Therefore, we examined protein structures to see whether C–H/O interactions between a Ser side-chain O and C–H_α_ of the subsequent residue might stabilize the *cis* amide bond at Ser-*cis*-non-Pro structures.

Xanthine-guanine phosphoribosyl transferase (XGPRT) is a critical enzyme for numerous bacteria, functioning in purine salvage. X-ray crystallography has demonstrated that a conserved Ser-*cis*-Arg in XGPRT^108,109^ is present (or implied by sequence conservation) in divergent bacteria, including *E. coli*, *Salmonella*, *Shigella*, and *Yersinia pestis*. This non-prolyl *cis* amide bond amide bond is part of the phosphoribose binding site, with the *cis*-Arg amide N–H hydrogen bonded to the ribosyl phosphate (Figure 14). Structures from *E. coli* and *Yersinia pestis* exhibit a close C–H/O interaction between the Ser O_γ_ and the Arg C–H_α_, which helps stabilize the *cis*-Arg amide bond. Geometry optimization of the local structures confirms the presence of a C–H/O interaction that appears to stabilize the *cis*-Arg conformation (Figure 14b). These crystallographic data suggest that C–H/O interactions might be a potentially general method to stabilize *cis* amide bonds in proteins.

**Figure 14.**
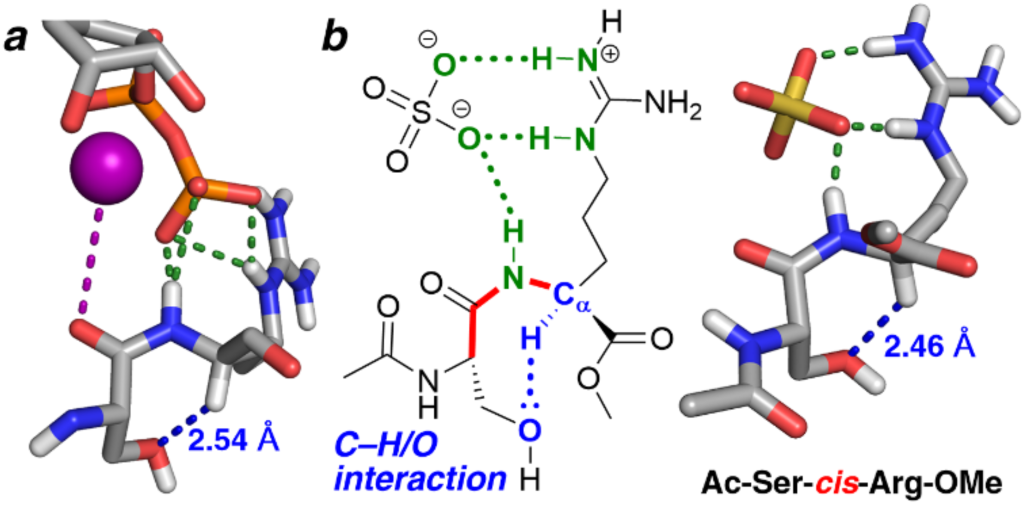
Ser-*cis*-nonPro structures with C–H/O interactions. (a) X-ray crystal structure of *E. coli* Xanthine-guanine phosphoribosyltransferase (XGPRT) bound to a 5-phospho-alpha-D-ribose 1-diphosphate (PRPP) analog (PDB 1a95, 2.00 Å resolution). The Ser-*cis*-Arg motif binds the anionic phosphate via N–H•••O hydrogen bonds (green) as well as the magnesium cofactor (Mg^2+^•••O, purple). Ser stabilizes the *cis*-Arg conformation via a C–H/O interaction (C_α_–H•••O distance 2.54 Å, blue). The Ser-*cis*-Arg motif is conserved in purine scavenging enzymes across multiple pathogenic bacteria. (b) ChemDraw representation of the structure of XGPRT from *Yersinia pestis* (PDB 6kp5, 1.10 Å resolution), with a sulfate anion in the active site and an equivalent C–H/O interaction (C_α_–H•••O distance 2.57 Å). The core Ser-*cis*-Arg•SO_4_ ^2-^ structure from this protein was truncated to Ac-Ser-*cis*-Arg-OMe•SO_4_ ^2-^ and subjected to full geometry optimization (right), which demonstrated a close C–H/O interaction (C_α_–H•••O distance 2.46 Å, blue) stabilizing the *cis*-Arg conformation.

## Methods

### Peptide synthesis

Dipeptides were synthesized by solution-phase synthesis. Synthesis and characterization details are in the Supporting Information.

### NMR spectroscopy

Compounds were characterized at 298 K in CDCl_3_ or CD_3_OD using either a Bruker 400 MHz NMR spectrometer equipped with a cryogenic QNP probe or a Bruker 600 MHz spectrometer equipped with a 5-mm Bruker SMART probe. 1-D NMR spectra were obtained using an excitation sculpting pulse sequence. Peaks were assigned using COSY and ^1^H-^13^C HSQC spectra. NMR spectra and additional experimental details are in the Supporting Information.

### X-ray crystallography

Boc-Ser-hyp(4-I-Ph)-OMe and Ac-Ser-hyp(4-I-Ph)-OMe were allowed to dissolve in acetone and subjected to crystallization via evaporation at 25 °C.

Crystallization of both compounds occurred over one week. Additional details are in the Supporting Information.

### Computational chemistry

Calculations were conducted with Gaussian 09.^110^ For small-molecule crystal structures, hydrogen positions were determined using a restrained geometry optimization, in which the positions of the heavy atoms were fixed based on those observed crystallographically, while the positions of the hydrogens were optimized using the M06-2X DFT functional and the Def2TZVP basis set in implicit water (IEFPCM).^111–113^ Full, unrestrained geometry optimization of molecules derived from crystal structures were determined using the same methods. Structures of all other molecules were determined via iterative geometry optimization, with final geometry optimization using the M06-2X functional with the 6-311++G(d,p) basis set in implicit water.^114^ Additional details of calculations and coordinates of all geometry-optimized structures are in the Supporting Information.

### Bioinformatics

On June 14, 2023, a search of the Protein Data Bank (PDB) was conducted for structures containing Ser-Pro sequences with a resolution ≤ 2.0 Å and a sequence similarity less than 30%. A search of the PDB for structures containing Ala-Pro sequences was conducted on July 17, 2023, using the same search parameters. Perl scripts were written to extract and calculate information used in analyses such as residue numbers, residue identities, dihedral angles, Ramachandran plots, interatomic distances, and amide hydrogen position estimations. Structures used in analyses were manually filtered to remove structures with nearby broken backbone amide bonds. The final data sets contained 2045 structures with SP sequences (1929 structures with *trans*-proline and 116 structures with *cis*-proline) and 1832 structures with AP sequences (1729 with *trans-*proline and 103 with *cis-*proline). Additional details are in the Supporting Information.

## Acknowledgements

We thank Himal Ganguly for preliminary analysis of C–H/O interactions in Ser-Pro sequences. We thank NSF (CHE-2004110) for funding. Instrumentation support was provided by NIH (GM110758, S10 OD026896A) and NSF (CHE-1229234).

## Supporting Information Available

Details of peptide synthesis and characterization; additional analysis of structures determined by DFT calculations; additional analysis of bioinformatics data; details of structures solved by X-ray crystallography; and coordinates of all geometry-optimized structures. This material is available free of charge via the journal web site.

