## Supporting Information for "Helical Twists and β-Turns in Structures at Serine–Proline Sequences: Stabilization of cis-Proline and type VI β-turns via C–H/O interactions"

**Contents**

S2. Materials
S2. Mass spectrometry
S2. NMR spectroscopy
S3. Synthesis of Boc- and Ac-L-serine-(2*S*,4*S*)-4-iodophenyl-hydroxyproline methyl ester (**3** and **5**)
S6. Synthesis of Boc- and Ac-L-alanine-(2*S*,4*S*)-4-iodophenyl-hydroxyproline methyl ester (**6** and **8**)
S8. NMR analysis of Boc-Ser-hyp(4-I-Ph)-OMe and Ac-Ser-hyp(4-I-Ph)-OMe
S9. Computational and bioinformatics methods and analysis
  S9. Computational geometry optimization of small molecule crystal structures and models generated from the PDB
  S14. Bioinformatics analysis of conformations at Ser-Pro sequences in the PDB
  S18. Analysis of type VI β-turns and noncovalent interactions at Ser-*cis*-Pro and Ala-*cis*-Pro
  S19. Analysis of noncovalent interactions at Ser-*trans*-Pro
  S20. Analysis of main-chain hydrogen-bonded secondary structures at Ser-Pro and Ala-Pro sequences
S27. Crystal structures, crystallographic details, and refinement parameters
  S28. Crystallographic details for Boc-L-serine-(2*S*,4*S*)-4-iodophenyl-hydroxyproline methyl ester (**3**)
  S41. Crystallographic details for Ac-L-serine-(2*S*,4*S*)-4-iodophenyl-hydroxyproline methyl ester (**5**)
S50. References
S51. Coordinates of structures obtained via geometry optimization calculations
S146. $^{1}$H and $^{13}$C NMR spectra of compounds **2**, **3**, **4**, **5**, **6**, **7**, and **8**

**Materials**

Boc-(2*S*,4*R*)-4-Hydroxyproline (Boc-Hyp-OH), Boc-L-serine (Boc-Ser-OH), Boc-L-alanine (Boc-Ala-OH), and 1-ethyl-3-(3-dimethylaminopropyl) carbodiimide hydrochloride (EDCI•HCl) were purchased from Chem-Impex (Wood Dale, IL). All solvents, acetic anhydride ($Ac_2O$), triphenylphosphine ($Ph_3P$), dimethyl sulfate, potassium carbonate ($K_2CO_3$), sodium hydroxide (NaOH), hydrochloric acid (HCl), and chloroform-d ($CDCl_3$) were purchased from Fisher Scientific. 4-Iodophenol and N,N-diisopropylethylamine (DIPEA) were purchased from Acros. Diisopropylazodicarboxylate (DIAD) was purchased from Sigma-Aldrich. Methylene chloride ($CH_2Cl_2$), tetrahydrofuran (THF), and toluene were dried using sodium sulfate and stored over 4 Å molecular sieves. Thin layer chromatography was conducted using Sorbent glass-backed plates (silica gel, 250 μm, 60 Å, F254). Flash chromatography was performed using 230-400 mesh (40-63 μm, 60 Å) silica gel from Sorbent.

**Mass spectrometry**

High-resolution mass spectrometry was performed on a Thermo Q-Exactive Orbitrap using conventional heated electrospray ionization (HESI).

**NMR spectroscopy**

Compounds were characterized via NMR spectroscopy recorded at 298 K on either on a Bruker 400 MHz NMR spectrometer equipped with a cryogenic QNP probe or on a Bruker 600 MHz spectrometer equipped with a 5-mm Bruker SMART probe, using a standard 30° flip pulse sequence, 65,536 data points, and a relaxation delay of 2.6 s. $^{13}C$ NMR spectra were acquired on the same instrumentation using a standard 30° flip pulse sequence, collecting 32,768 data points, and a relaxation delay of 5.0 s. $^{1}H$ and $^{13}C$ NMR spectra were internally referenced with deuterated chloroform, methanol, or trimethylsilylpropanoic acid, depending on sample solvent.

**Synthesis of Boc-L-Serine-(2*S*,4*S*)-4-iodophenyl-hydroxyproline methyl ester (3) and Ac-L-Serine-(2*S*,4*S*)-4-iodophenyl-hydroxyproline methyl ester (5)**

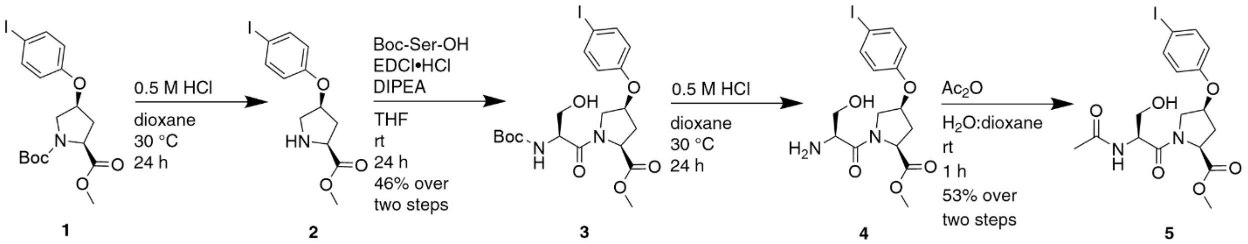

1 2 3 4 5

**Scheme S1**. Synthesis of Boc- and Ac-L-Serine-(2*S*,4*S*)-4-iodophenyl-hydroxyproline methyl ester (**3** and **5**).

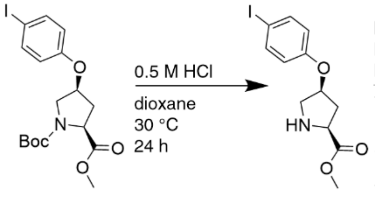

1 2

**(2*S*,4*S*)-4-Iodophenyl-hydroxyproline methyl ester (2)**. Boc-hyp(4-I-Ph)-OMe (**1**) was prepared via the synthesis reported by Daniecki *et al*. The NMR data corresponded to the literature values.[1] **1** (23.3 g, 2.24 52.0 mmol) was allowed to dissolve in 1,4-dioxane (260 mL). 0.5 M HCl (260 mL) was added to the solution and stirred for 24 hours at 30 °C. The solvent was removed under reduced pressure and the yellow oil was resuspended in water before being acidified to pH 3. The aqueous solution was washed with ethyl acetate (3 × 100 mL). The aqueous solution was concentrated under reduced pressure to obtain **2** (16.1 g crude) as a white solid. This compound was used in the next step without purification. $^1$H NMR (600 MHz, $CD_3OD$) δ 7.76–7.52 (m, 2H), 6.92–6.66 (m, 2H), 5.20 (m, 1H), 4.69 (dd, $J$ = 9.4, 3.8 Hz, 1H), 3.82 (s, 3H), 3.77–3.52 (m, 2H), 2.84–2.50 (m, 2H). $^{13}$C NMR (151 MHz, $CD_3OD$) δ 168.9, 155.7, 139.1, 117.7, 83.7, 74.4, 59.1, 52.7, 51.0, 34.0. HRMS (HESI) *m/z*: $[M + H]^+$ calcd for $C_{12}H_{14}INO_3$ 347.0019, found 347.0087.

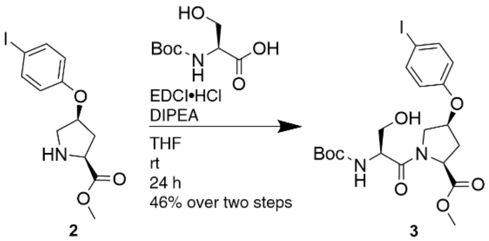

2 3

**Boc-L-Serine-(2*S*,4*S*)-4-iodophenyl-hydroxyproline methyl ester (3)**. **2** (16.1 g crude), Boc-L-serine (4.87 g, 46.3 mmol), and DIPEA (17.7 mL, 102 mmol) were allowed to dissolve in THF (309 mL). EDCI•HCl (10.7 g, 55.6 mmol) was added, and the solution was placed under nitrogen and allowed to stir for 24 hours at room temperature. The solvent was removed under reduced pressure and resuspended in ethyl acetate (100 mL). The solution was washed with distilled water, then acidified to pH 3 with 1 M HCl (3 × 100 mL). The organic layer was dried over $Na_2SO_4$ and filtered. The crude product was purified via silica column chromatography (0 to 5% MeOH in $CH_2Cl_2$ v/v) to yield the product **3** (12.9 g, 24.1 mmol) as a white solid in 46% yield over two steps. NMR spectra reflected a mixture of *trans* and *cis* proline rotamers ($K_{trans/cis}$ = 6.6). $^1$H NMR (600 MHz, $CDCl_3$) δ 7.66–7.43 (m, 2H), 6.74–6.43 (m, 2H), 4.96 (m, 0.9H), 4.91–4.76 (m, 1.1H), 4.56 (m, 1H), 4.21–3.96 (m, 1.7H), 3.94–3.77 (m, 1.4H), 3.77–3.63 (m, 4H), 2.82–2.23 (m, 2H), 1.43 (s, 9H). $^{13}$C NMR (151 MHz, $CDCl_3$) δ 171.8, 171.1, 156.4, 155.9, 139.6, 118.8, 84.7, 84.4, 80.5, 76.0, 74.1, 64.6, 57.7, 53.5, 53.2, 52.6, 37.2, 35.5, 28.7. HRMS (HESI) *m/z*: $[M + H]^+$ calcd for $C_{20}H_{27}IN_2O_7$ 535.0941, found 535.0934.

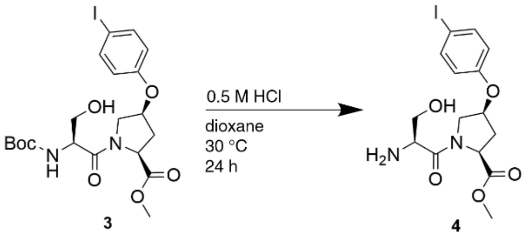

3 4

**L-Serine-(2*S*,4*S*)-4-iodophenyl-hydroxyproline methyl ester (4)**. **3** (2.00 g, 3.74 mmol) was allowed to dissolve in 1,4-dioxane (18.7 mL). 0.5 M HCl (18.7 mL) was added to the solution, which was then stirred for 24 hours at 30 °C. The solvent was removed under reduced pressure and the yellow oil was resuspended in water before being acidified to pH 3. The aqueous solution was washed with ethyl acetate (3 × 10 mL). The aqueous solution was concentrated under reduced pressure to obtain **4** (1.56 g crude) as a white solid. This crude compound was used in the subsequent reaction without further purification. NMR spectra reflected a mixture of *trans* and *cis* proline rotamers ($K_{trans/cis}$ = 1.9). $^1$H NMR (600 MHz, $CD_3OD$) δ 7.58 (m, 2H), 6.98–6.34 (m, 2H), 5.24–5.00 (m, 1H), 4.95 (d, *J* = 8.6 Hz, 0.6H), 4.78 (d, *J* = 9.3 Hz, 0.4H), 4.65–4.22 (m, 1H), 4.15–3.86 (m, 2H), 3.87–3.50 (m, 5H), 2.94–2.31 (m, 2H). $^{13}$C NMR (151 MHz, $CD_3OD$) δ 173.1, 172.6, 168.3, 167.4, 157.6, 139.7, 119.1, 118.5, 77.2, 75.04, 61.4, 60.8, 59.2, 59.0, 55.5, 55.1, 53.6, 53.2, 36.1, 34.9. HRMS (HESI) *m/z*: $[M + H]^+$ calcd for $C_{15}H_{20}IN_2O_5^+$ 435.0417, found 435.0421.

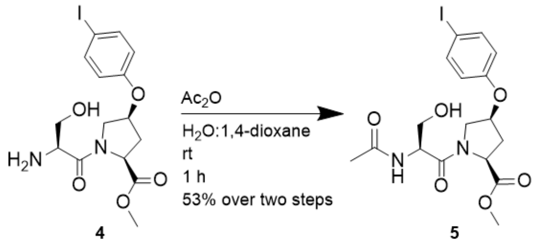

**4** **5**

**Ac-L-Serine-(2*S*,4*S*)-4-iodophenyl-hydroxyproline methyl ester (5)**. **4** (1.56 g crude) was allowed to dissolve in 1,4-dioxane (18.0 mL) and distilled water (18.0 mL). $Ac_2O$ (0.551 g, 5.40) and DIPEA (1.88 mL, 10.8 mmol) were added, and the solution was allowed to stir for 1 hour. The solvent was removed under reduced pressure and the crude product was resuspended in ethyl acetate. The solution was washed with distilled water acidified to pH 3 with 1 M HCl (3 × 10 mL). The organic layer was dried over $Na_2SO_4$ and filtered. After the solvent was removed under reduced pressure, the colorless oil was purified via silica column chromatography (50% to 100% 2-propanol in hexanes v/v) to yield the product **4** (1.16 g, 2.45 mmol) as a white solid in 53% yield over two steps. NMR spectra reflected a mixture of *trans* and *cis* proline rotamers ($K_{trans/cis}$ = 4.9). $^{1}H$ NMR (600 MHz, $CDCl_3$) δ 7.69–7.45 (m, 2H), 6.73–6.46 (m, 3H), 5.01–4.88 (m, 1H), 4.88–4.71 (m, 2H), 4.19–3.98 (m, 2H), 3.95–3.77 (m, 1H), 3.78–3.61 (m, 4H), 3.11–2.32 (m, 2H), 2.01 (s, 3H). $^{13}C$ NMR (151 MHz, $CDCl_3$) δ 171.4, 170.3, 155.6, 139.3, 118.0, 84.4, 84.0, 75.6, 73.7, 64.6, 63.8, 58.1, 57.4, 53.0, 52.8, 52.4, 52.3, 36.6, 34.2, 23.6, 23.0. HRMS (HESI) *m*/*z*: $[M + H]^+$ calcd for $C_{17}H_{21}IN_2O_6$ 476.0444, found 477.0517.

**Synthesis of Boc-L-Alanine-(2*S*,4*S*)-4-iodophenyl-hydroxyproline methyl ester (6) and Ac-L-Alanine-(2*S*,4*S*)-p-iodophenyl-4-hydroxyproline methyl ester (8)**

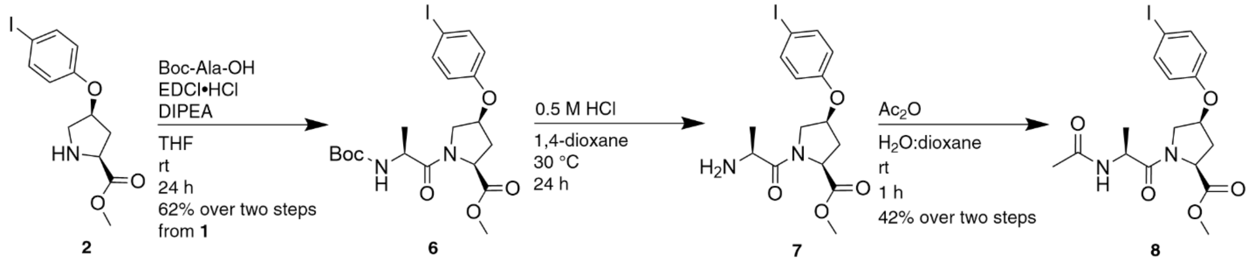

**Scheme S2**. Synthesis of Boc- and Ac-L-Alanine-(2*S*,4*S*)-4-iodophenyl-hydroxyproline methyl ester (**6** and **8**).

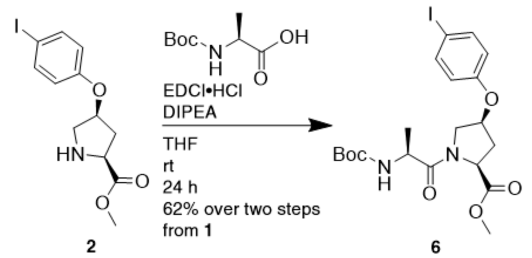

**Boc-L-Alanine-(2*S*,4*S*)-4-iodophenyl-hydroxyproline methyl ester (6)**. **2** (5 g crude), Boc-L-alanine (1.83 g, 9.64 mmol), and DIPEA (3.69 mL, 21.2 mmol) were allowed to dissolve in THF (309 mL). EDCI•HCl (1.80 g, 11.6 mmol) was added, and the solution was placed under nitrogen and allowed to stir for 24 hours at room temperature. The solvent was removed under reduced pressure and resuspended in ethyl acetate (20 mL). The solution was washed with distilled water, then acidified to pH 3 with 1 M HCl (3 × 20 mL). The organic layer was dried over $Na_2SO_4$ and filtered. The crude product was purified via silica column chromatography (0 to 5% MeOH in $CH_2Cl_2$ v/v) to yield the product **2** (3.10 g, 5.98 mmol) as a white solid in 62% yield over two steps. NMR spectra reflected a mixture of *trans* and *cis* proline rotamers ($K_{trans/cis}$ = 6.7). $^1$H NMR (400 MHz, $CDCl_3$) δ 7.67–7.41 (m, 2H), 6.72–6.42 (m, 2H), 5.43 (m, 1H), 5.07–4.86 (m, 1H), 4.82 (m, 1H), 4.10 (m, 1H), 3.92–3.61 (m, 4H), 2.89–2.25 (m, 2H), 1.42 (m, 9H), 1.32 (m, 3H).$^{13}$C NMR (101 MHz, $CDCl_3$) δ 172.6, 170.8, 156.7, 155.5, 154.9, 138.6, 118.6, 84.3, 84.0, 80.0, 75.8, 73.6, 58.6, 56.5, 52.9, 52.4, 52.2, 52.0, 51.5, 48.2, 36.8, 34.2, 30.9, 28.3, 19.5, 18.0. HRMS (HESI) *m/z*: $[M + H]^+$ calcd for $C_{20}H_{28}IN_2O_6^+$ 519.0992, found 519.0985.

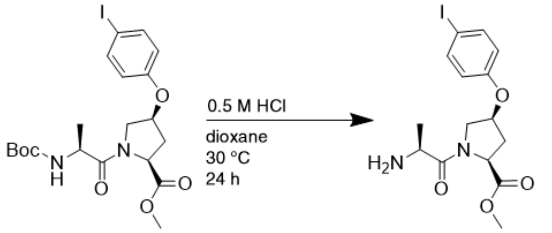

6 7

**L-Alanine-(2*S*,4*S*)-4-iodophenyl-hydroxyproline methyl ester (7)**. **6** (3.10 g, 5.98 mmol) was allowed to dissolve in 1,4-dioxane (29.9 mL). 0.5 M HCl (29.9 mL) was added to the solution and the resultant mixture stirred for 24 hours at 30 °C. The solvent was removed under reduced pressure and the yellow oil was resuspended in water before being acidified to pH 3. The aqueous solution was washed with ethyl acetate (3 × 100 mL). The aqueous solution was concentrated under reduced pressure to obtain **7** (2.00 g crude) as a white solid. This crude compound was used in the subsequent reaction without further purification. NMR spectra reflected a mixture of *trans* and *cis* proline rotamers ($K_{trans/cis}$ = 7.7). $^{1}$H NMR (600 MHz, $CDCl_3$) δ 7.55 (m, 2H), 6.57 (m, 2H), 4.92 (m, 1H), 4.65 (m, 1H), 4.20 (m, 1H), 4.05–3.47 (m, 5H), 3.18–2.11 (m, 2H), 1.49 (s, 3H). $^{13}$C NMR (151 MHz, $CDCl_3$) δ 170.6, 155.9, 155.7, 138.7, 117.9, 117.2, 84.5, 84.3, 75.5, 73.2, 58.0, 57.4, 53.4, 52.7, 52.6, 51.8, 48.5, 48.3, 36.0, 33.8, 16.4, 15.7. HRMS (HESI) *m/z*: $[M + H]^+$ calcd for $C_{15}H_{20}IN_2O_4^+$ 419.0467, found 419.0449.

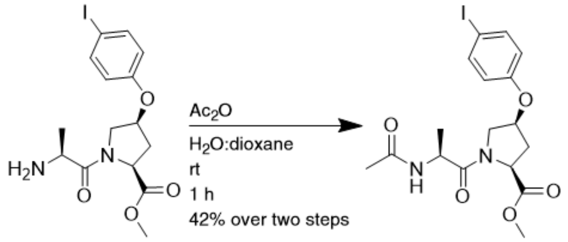

7 8

**Ac-L-Alanine-(2*S*,4*S*)-4-iodophenyl-hydroxyproline methyl ester (8)**. **7** (2.00 g crude) was allowed to dissolve in 1,4-dioxane (23.9 mL) and distilled water (23.9 mL). $Ac_2O$ (0.678 mL, 7.17 mmol) and DIPEA (2.50 mL, 14.3 mmol) were added, and the solution was allowed to stir for 1 hour. The solvent was removed under reduced pressure and the crude product was resuspended in ethyl acetate. The solution was washed with distilled water acidified to pH 3 with 1 M HCl (3 × 10 mL). The organic layer was dried over $Na_2SO_4$ and filtered. After the solvent was removed under reduced pressure, the colorless oil was purified via silica column chromatography (50% to 100% 2-propanol in hexanes v/v) to yield the product **8** (1.17 g, 2.53 mmol) as a white solid in 42% yield over two steps. NMR spectra reflected a mixture of *trans* and *cis* proline rotamers ($K_{trans/cis}$ = 4.9). $^{1}$H NMR (400 MHz, $CDCl_3$) δ $^{1}$H NMR (400 MHz, $CDCl_3$) δ 7.67–7.45 (m, 2H), 6.72–6.44 (m, 3H), 5.05–4.87 (m, 1H), 4.81 (m, 0.8H), 4.68 (m, 1H), 4.54 (m, 0.2H), 4.10 (m, 1H), 3.93–3.65 (m, 4H), 2.88–2.30 (m, 2H), 2.01 (d, *J* = 2.5 Hz, 3H), 1.35 (m, 3H). $^{13}$C NMR (101 MHz, $CDCl_3$) δ 171.8, 170.6, 170.4, 156.1, 156.0, 139.4, 118.5, 84.3, 58.0, 57.2, 53.0, 52.4, 52.2, 52.0, 47.1, 46.8, 36.7, 34.3, 30.4, 22.9, 19.1, 17.9. HRMS (HESI) *m/z*: $[M + H]^+$ calcd for $C_{17}H_{21}IN_2O_5$ 461.0573, found 461.0559.

### NMR analysis of Boc-Ser-hyp(4-I-Ph)-OMe and Ac-Ser-hyp(4-I-Ph)-OMe

**Table S1. $^1$H NMR analysis of Pro H$_\alpha$ and $K_{trans/cis}$ in $D_2O$, $CD_3OD$, and $CDCl_3$.**[a,b] $^1$H NMR spectra of Boc-Ser-hyp(4-I-Ph)-OMe and Ac-Ser-hyp(4-I-Ph)-OMe were collected in $CD_3OD$ and $CDCl_3$. Only Ac-Ser-hyp(4-I-Ph)-OMe was sufficiently soluble in $D_2O$ to achieve an adequate measurement. The Pro H$_\alpha$ chemical shifts of Ac-Ser-hyp(4-I-Ph)-OMe in $D_2O$ were determined using TOCSY experiment crosspeaks. Boc-Ala-hyp(4-I-Ph)-OMe and Ac-Ala-hyp(4-I-Ph)-OMe were analyzed as controls.

| | | δ Pro H$_\alpha$, ppm | | |
|---|---|---|---|---|
| **$D_2O$** | **$K_{trans/cis}$** | **$\delta_{cis}$** | **$\delta_{trans}$** | **$\Delta\delta_{cis–trans}$** |
| **Ac-Ser-hyp(4-I-Ph)-OMe** | 2.3 | 5.05 | 4.79 | +0.26 |
| **$CD_3OD$** | **$K_{trans/cis}$** | **$\delta_{cis}$** | **$\delta_{trans}$** | **$\Delta\delta_{cis–trans}$** |
| **Boc-Ser-hyp(4-I-Ph)-OMe** | 2.6 | 5.02 | 4.76 | +0.26 |
| **Boc-Ala-hyp(4-I-Ph)-OMe** | 5.9 | 4.78 | 4.74 | +0.04 |
| **Ac-Ser-hyp(4-I-Ph)-OMe** | 2.3 | 5.03 | 4.74 | +0.29 |
| **Ac-Ala-hyp(4-I-Ph)-OMe** | 4.2 | 4.78 | 4.74 | +0.04 |
| **$CDCl_3$** | **$K_{trans/cis}$** | **$\delta_{cis}$** | **$\delta_{trans}$** | **$\Delta\delta_{cis–trans}$** |
| **Boc-Ser-hyp(4-I-Ph)-OMe** | 6.6 | 4.81 | 4.88 | −0.07 |
| **Boc-Ala-hyp(4-I-Ph)-OMe** | 6.7 | 4.54 | 4.83 | −0.29 |
| **Ac-Ser-hyp(4-I-Ph)-OMe** | 4.9 | 4.93 | 4.86 | +0.13 |
| **Ac-Ala-hyp(4-I-Ph)-OMe** | 4.9 | 4.54 | 4.81 | −0.27 |

[a] $K_{trans/cis}$ is determined from the integration of resonances of *trans*-proline and *cis-proline*.

[b] $\Delta\delta_{cis–trans} = \delta_{cis}$ Pro H$_\alpha$ – $\delta_{trans}$ Pro H$_\alpha$

### Computational geometry optimization of small molecule crystal structures and models generated from the PDB

#### Calculations

All calculations were conducted with Gaussian09 or Gaussian16. Visualization was conducted with PyMOL.

#### Hydrogen position optimization of crystal structures

Hydrogen positions were optimized computationally from the solved crystal structures since the X-ray radiation is only weakly scattered by hydrogen and therefore the positions of hydrogens are not fully accurate when mapping hydrogen atom location. Hydrogen position optimization was performed using the atomic coordinates provided by X-ray crystal structures. The positions of the heavy atoms were fixed. Initial restrained geometry optimization was conducted on each molecule with the M06-2X DFT functional and the Def2SVP basis set, with subsequent optimization using the Def2TZVP basis set, in both cases with implicit water solvation (IEFPCM continuum polarization model).

#### Full geometry optimization of 3 and 5

Full geometry optimization of **3** and **5** was conducted using the M06-2X DFT functional, initially optimized with the Def2SVP basis set and further optimized with the Def2TZVP basis set using implicit water solvation (IEFPCM continuum polarization model). Models were generated using the X-ray crystal structures as initial structures.

#### Full geometry optimization of minimal model peptides

Structures of all minimal peptides generated from small molecules and structures from the Protein Data Bank were iteratively optimized, with final geometry optimization using the M06-2X DFT functional, the 6-311++G(d,p) basis set, and implicit water solvation (IEFPCM continuum polarization model).

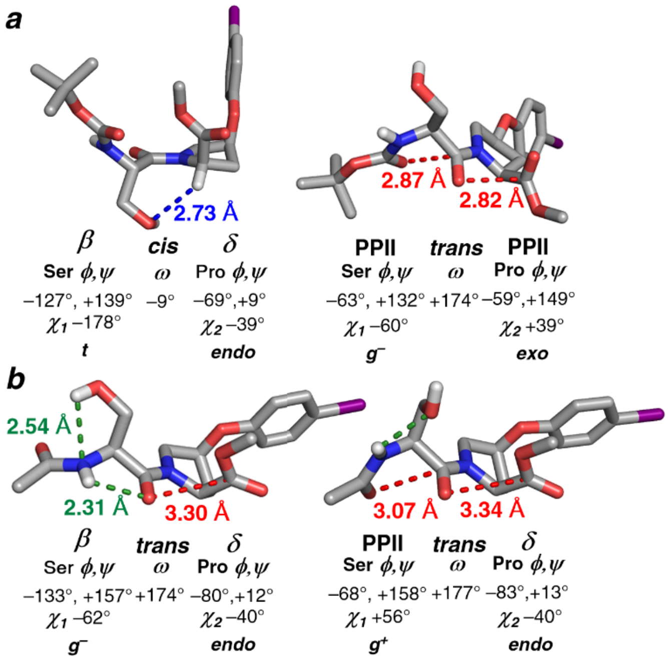

**Figure S1. Geometry-optimized structures of Boc-Ser-hyp(4-I-Ph)-OMe and Ac-Ser-hyp(4-I-Ph)-OMe in the conformations observed crystallographically**. (a) Boc-Ser-hyp(4-I-Ph)-OMe and (b) Ac-Ser-hyp(4-I-Ph)-OMe structures were subjected to full geometry optimization using the M06-2X DFT functional with the Def2TZVP basis set and implicit $H_2O$ solvation. Boc-Ser-hyp(4-I-Ph)-OMe with *cis*-proline (left) showed a closer $C_\alpha$–H•••O interaction ($C_\alpha$–H•••O distance 2.73 Å, blue) while maintaining the same overall conformation as in the crystal structure. Boc-Ser-hyp(4-I-Ph)-OMe with *trans*-proline (right) maintained the same overall conformation as in the crystal structure, with closer $n \rightarrow \pi^*$ interactions (red). In the structure of Ac-Ser-hyp(4-I-Ph)-OMe with Ser in the β conformation (left), a new intraresidue interaction was observed between the Ser side chain and the amide N (O–H•••N distance 2.54 Å, green). The structure with Ser in the PPII conformation (right) maintained the same overall conformation as in the crystal structure.

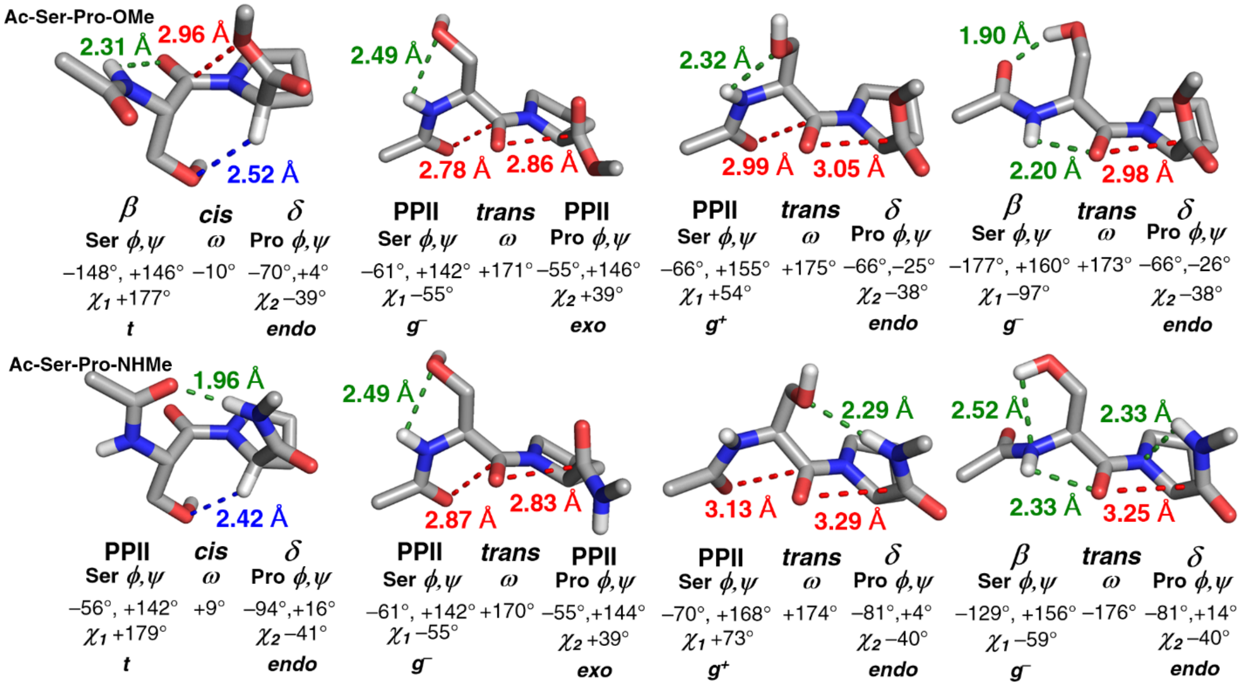

**Figure S2. Ac-SP-OMe and Ac-SP-NHMe optimized from crystallographically observed structures**. Ac-SP-OMe and Ac-SP-NHMe structures were Geometry-optimized from structures observed crystallographically in Boc-Ser-hyp(4-I-Ph)-OMe and Ac-Ser-hyp(4-I-Ph)-OMe. From the crystal structures, the aryloxy group was changed to a hydrogen. When applicable, Boc groups were truncated to Ac, and the C-terminal methyl ester was changed to a methyl amide. These truncated models confirmed the relevance of the crystallographically observed local structures and interactions with unmodified proline. All structures were Geometry-optimized using the M06-2X DFT functional, the 6-311++G(d,p) basis set, and implicit water solvation (IEFPCM continuum polarization model).

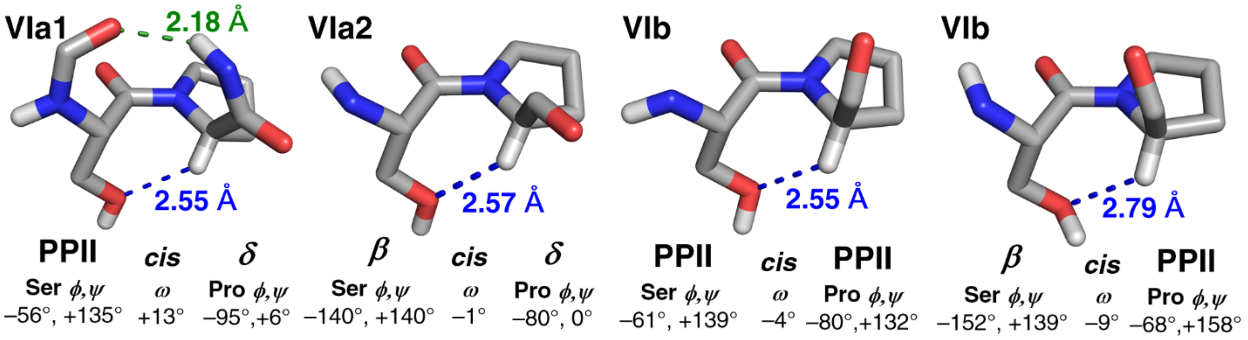

**Figure S3. Type VI β-turns at Ser-Pro stabilized via C–H/O interactions in the PDB**. Structures of the hyaluronan binding module of the *S. pneumoniae* hyaluronate lyase (pdb 4d0q: type VIa1, PcisD), angiostatin (pdb 2doh: type VIa2, BcisD), *P. falciparum* FVO apical membrane antigen 1 (pdb 6n87: type VIb, BcisP), and *M. tuberculosis* dUTPase (pdb 3loj: type VIb, PcisP), which contain different type VI β-turns at Ser-Pro with $C_{\alpha}$–H•••O interactions (blue). Type VIa1 β-turns inherently have $O_i$•••H–$N_{i+3}$ hydrogen bonds (green), while other type VI β-turn subtypes are not hydrogen-bonded via the main chain.

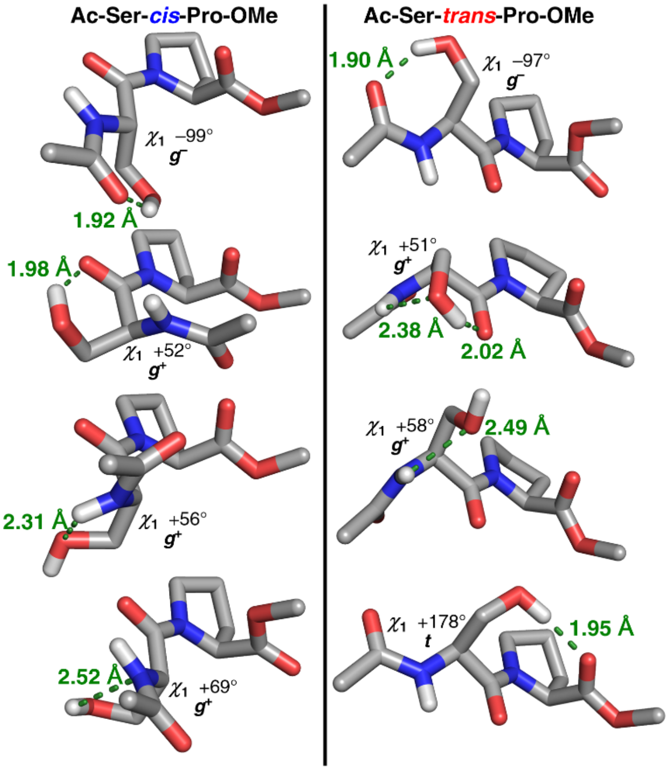

**Figure S4. Local Ser hydroxyl side chain-main chain hydrogen bonds at Ser-Pro sequences observed via DFT calculations**. Ac-Ser-*cis*-Pro-OMe (left) and Ac-Ser-*trans*-Pro-OMe (right) structures were examined for local hydrogen bonding patterns by rotating Ser $\chi_1$ to the $g^-$, $g^+$, and *t* rotamers, and followed by conducting geometry optimization calculations to generate the resultant structures.

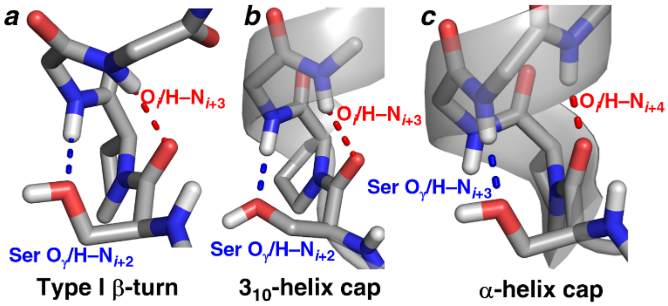

**Figure S5. Most frequent hydrogen-bonded secondary structures at Ser-*trans*-Pro in the PDB**. (a) Structure of fungal serine ammonia-lyase from *R. miehei* (pdb 5c3u) containing a type I β-turn in the Ser-*trans*-Pro-X-X register. The turn is stabilized via a Ser $O_\gamma$/H–$N_{i+2}$ hydrogen bond. (b) Structure of Golgi alpha-mannosidase II D341N mutant complex with 2-F-mannosyl-F (pdb 1qx1) with a Ser-*trans*-Pro sequence initiating a $3_{10}$-helix. With SP as the helix N-capping sequence and Ser engaged in an $O_\gamma$/H–$N_{i+2}$ hydrogen bond, there are no solvent-exposed amide hydrogens at the N-terminus of the $3_{10}$-helix. (c) Structure of the minimal scaffold domain of Ste5 (pdb 3fze) containing an α-helix where Ser is the Ncap residue and Pro is the first residue of the helix in the α-helical conformation. With Pro as the helix-initiating residue and the Ncap Ser engaged in an $O_\gamma$/H–$N_{i+3}$ hydrogen bond, there is only one solvent-exposed amide hydrogen at the N-terminus of the α-helix.

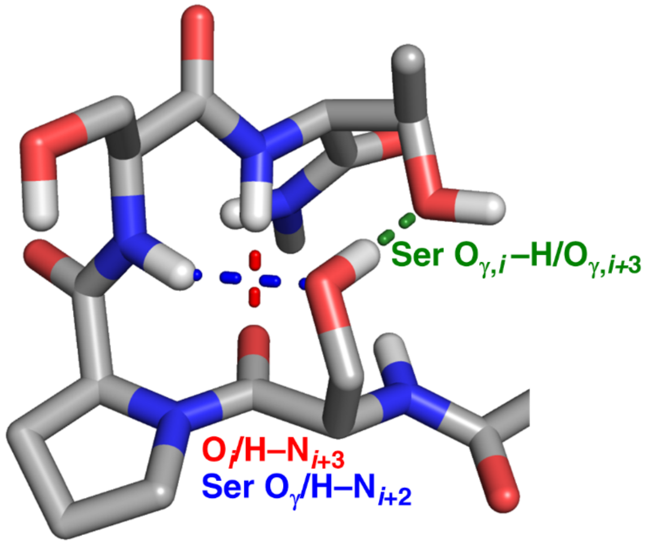

**Figure S6. S-*trans*-P-S-T type I β-turn stabilization via simultaneous Ser $O_\gamma$•••H–$N_{i+2}$ and Ser $O_\gamma$–H•••$O_{\gamma,i+2}$ hydrogen bonds in the PDB**. Structure of colicin M imported by *E. coli* (pdb 3da4) containing a Ser-*trans*-Pro-S-T type I β-turn. In the structure, Ser $O_{\gamma,i}$ is observed simultaneously acting as a hydrogen bond acceptor to H–$N_{i+2}$ (blue) and as a hydrogen bond donor to $O_{\gamma,i+3}$ (green).

### Bioinformatics analysis of conformations at Ser-Pro sequences in the PDB

#### Structure data set collection, refinement, and analysis

Structures containing Ser-Pro and Ala-Pro sequences with a resolution better than 2.0 Å and a sequence similarity of less than 30% were downloaded from the RCSB Protein Data Bank. Perl scripts were used to generate text files containing the PDB code, resolution, chain, residue number, secondary structures, dihedral angles, Ramachandran regions, specified interatomic distances, and neighboring residue identities of each SP and AP structure. The data set was manually refined on Microsoft Excel via removal of structures with broken Ser and Ala $\phi$ angles, removal of structures with a positive Pro $\phi$, and removal of all structures with broken $\psi$ angles on a case-by-case basis when the required analysis extended to residues past Pro. The data search after refinement yielded 2045 structures with SP sequences and 1832 structures with AP sequences.

#### Estimation of main-chain amide hydrogen positions for bioinformatics

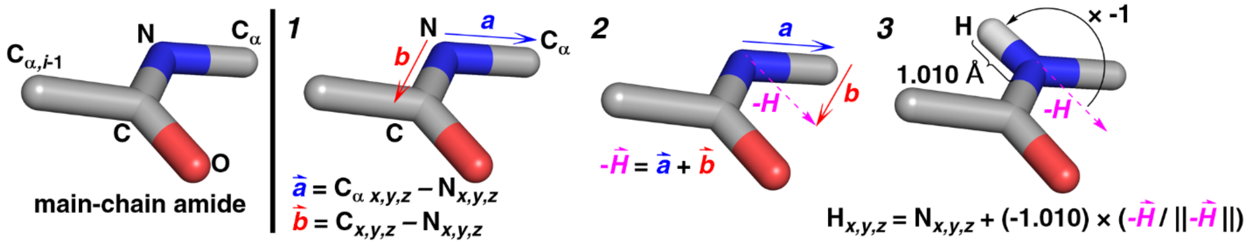

**Figure S7. General formula for estimating main-chain amide hydrogen positions from pdb files**. Amide hydrogen positions were estimated by (1) generating two vectors from the position of the amide nitrogen to the neighboring $C_\alpha$ and from the position of the amide nitrogen to its amide carbon, (2) summing these vectors to generate a new vector in the same plane as the amide in the assumed opposite direction of the hydrogen, and (3) inversing the new vector, scaling it to 1.010 Å, and adding it to the position of the amide nitrogen to generate an estimated position for H.

### Definition and analysis of Ramachandran space occupied by Ser-Pro and Ala-Pro

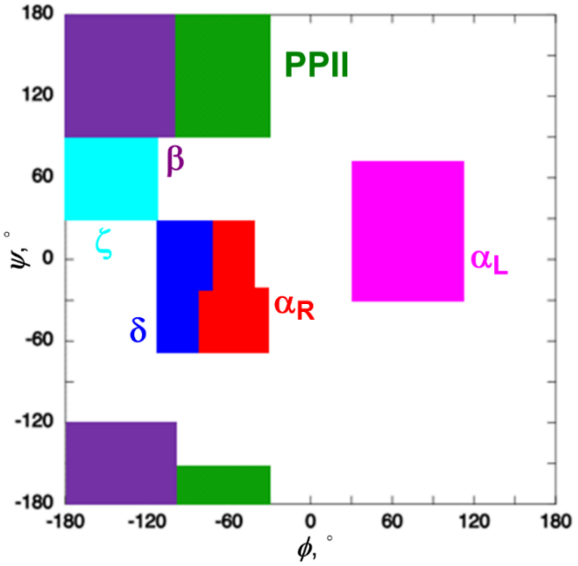

**Figure S8. Definition of secondary structures by their position on the Ramachandran plot**. Nomenclature and definitions of the regions occupied by residues in the Ramachandran plot. The $\phi$, $\psi$ ranges corresponding to the designated regions are indicated in Table S2.

**Table S2. Definitions of conformations as employed in bioinformatics analysis**. 10 nonoverlapping rectangular regions were used to define secondary structures via ranges of combinations of $\phi$ and $\psi$ torsion angles on the Ramachandran plot. The values in parentheses represent the maximum range of the $\phi$ and $\psi$ values of each region on the Ramachandran plot.

| conformation | $\phi$, $\psi$ range |
|---|---|
| $\alpha_L$ | +70° (±40°), +15° (±45°) |
| $\alpha_R$ | œ55° (±15°), +5° (±25°) |
| $\alpha_R$ | œ55° (±25°), œ40° (±20°) |
| β | œ140° (±40°), œ150° (±30°) |
| β | œ140° (±40°), +135° (±45°) |
| δ | œ90° (±20°), +5° (±25°) |
| δ | œ95° (±15°), œ40° (±20°) |
| PPII | œ65° (±35°), +135° (±45°) |
| PPII | œ65° (±35°), œ165° (±15°) |
| ζ | œ145° (±35°), +60° (±30°) |

**Table S3. Frequency of Pro-amide isomers observed at Ser-Pro and Ala-Pro sequences in the PDB**. PDB structures containing Ser-Pro and Ala-Pro sequences were analyzed and separated by Pro-amide isomer.

| | number of structures | | | % of structures | |
|---|---|---|---|---|---|
| **X** | **total** | **X-*trans*-Pro** | **X-*cis*-Pro** | **X-*trans*-Pro** | **X-*cis*-Pro** |
| **Ser** | 2045 | 1929 | 116 | 94.3 | 5.7 |
| **Ala** | 1832 | 1729 | 103 | 94.4 | 5.6 |

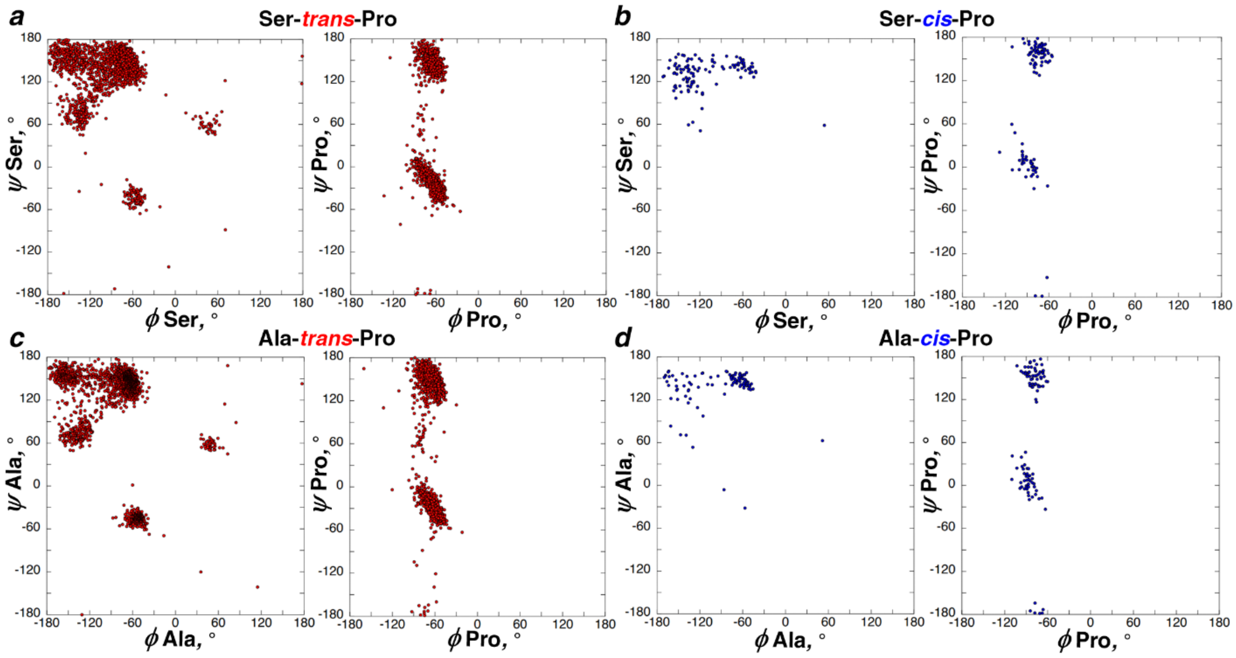

**Figure S9. Ramachandran plots of Ser-Pro and Ala-Pro sequences in the PDB**. Ramachandran plots of (a) Ser-*trans*-Pro and (b) Ser-*cis*-Pro sequences in the PDB compared to the Ramachandran plots of (c) Ala-*trans*-Pro and (d) Ala-*cis*-Pro sequences in the PDB. Data sets were compiled from all crystal structures containing SP sequences and AP sequences solved with resolution ≤ 2.0 Å and sequence similarity less than 30%. The search yielded 2045 SP structures, 1929 containing *trans*-proline and 116 containing *cis*-proline; and 1832 AP structures, 1729 containing *trans*-proline and 103 containing *cis*-proline.

**Table S4. Frequency of conformations observed at Ser-Pro and Ala-Pro sequences in the PDB**. Populations of combinations of conformations observed at Ser-Pro and Ala-Pro sequences in the PDB. Populations are represented as a percent of structures among those at each proline amide isomer.

| conformation | | *trans* | | *cis* | |
|---|---|---|---|---|---|
| | | SP | AP | SP | AP |
| X | Pro | $\%_{total}$ | $\%_{total}$ | $\%_{total}$ | $\%_{total}$ |
| $\alpha_R$ | $\alpha_R$ | 4.2 | 12.8 | 0 | 0 |
| $\alpha_R$ | β | 0.0 | 0.0 | 0 | 0 |
| $\alpha_R$ | δ | 0.4 | 0.8 | 0 | 0 |
| $\alpha_R$ | PPII | 0.3 | 0.5 | 0 | 1 |
| $\alpha_R$ | ζ | 0.0 | 0.0 | 0 | 0 |
| $\alpha_R$ | undefined | 0.0 | 0.0 | 0 | 0 |
| $\alpha_L$ | $\alpha_R$ | 0.1 | 0.2 | 0 | 0 |
| $\alpha_L$ | β | 0.0 | 0.0 | 0 | 0 |
| $\alpha_L$ | δ | 0.2 | 0.1 | 0 | 0 |
| $\alpha_L$ | PPII | 0.9 | 1.2 | 1 | 0 |
| $\alpha_L$ | ζ | 0.0 | 0.0 | 0 | 0 |
| $\alpha_L$ | undefined | 0.0 | 0.1 | 0 | 0 |
| β | $\alpha_R$ | 13.8 | 3.2 | 1 | 2 |
| β | β | 0.0 | 0.1 | 0 | 0 |
| β | δ | 4.8 | 1.0 | 9 | 3 |
| β | PPII | 12.2 | 18.0 | 55 | 28 |
| β | ζ | 0.0 | 0.0 | 0 | 0 |
| β | undefined | 0.8 | 0.9 | 1 | 0 |
| δ | $\alpha_R$ | 0.0 | 0.1 | 0 | 0 |
| δ | β | 0.0 | 0.0 | 0 | 0 |
| δ | δ | 0.1 | 0.0 | 0 | 1 |
| δ | PPII | 0.0 | 0.0 | 0 | 0 |
| δ | ζ | 0.0 | 0.0 | 0 | 0 |
| δ | undefined | 0.0 | 0.0 | 0 | 0 |
| PPII | $\alpha_R$ | 30.2 | 11.5 | 0 | 0 |
| PPII | β | 0.1 | 0.1 | 0 | 1 |
| PPII | δ | 6.1 | 2.2 | 17 | 32 |
| PPII | PPII | 15.9 | 34.3 | 9 | 23 |
| PPII | ζ | 0.0 | 0.0 | 1 | 4 |
| PPII | undefined | 0.4 | 1.5 | 3 | 0 |
| ζ | $\alpha_R$ | 3.4 | 4.0 | 0 | 0 |
| ζ | β | 0.0 | 0.0 | 1 | 1 |
| ζ | δ | 0.6 | 1.0 | 1 | 0 |
| ζ | PPII | 4.1 | 4.9 | 2 | 3 |
| ζ | ζ | 0.0 | 0.0 | 0 | 0 |
| ζ | undefined | 0.0 | 0.1 | 0 | 0 |
| undefined | $\alpha_R$ | 0.6 | 0.2 | 0 | 0 |
| undefined | β | 0.0 | 0.0 | 0 | 0 |
| undefined | δ | 0.1 | 0.0 | 0 | 0 |
| undefined | ζ | 0.7 | 1.2 | 0 | 0 |
| undefined | undefined | 0.0 | 0.0 | 0 | 1 |

**Analysis of type VI β-turns and noncovalent interactions at Ser-*cis*-Pro and Ala-*cis*-Pro**

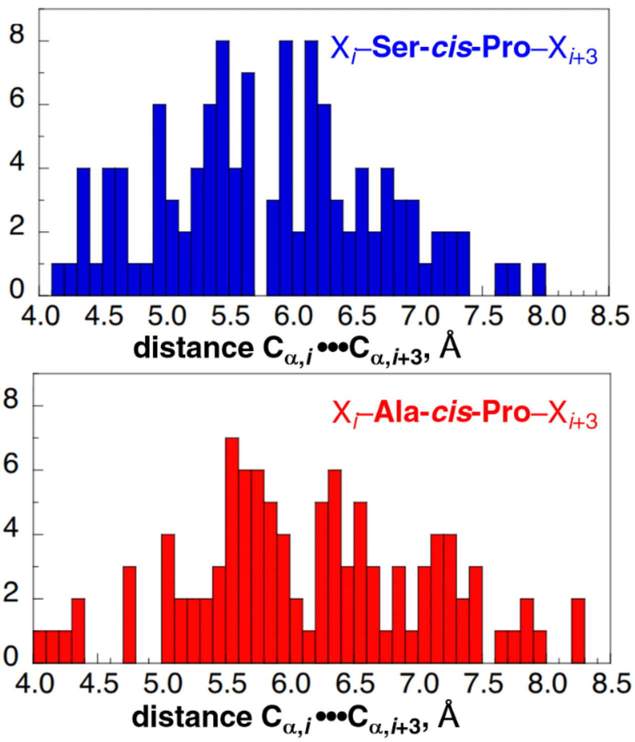

**Figure S10. Quantification of type VI β-turns at X$_i$–S-*cis*-P–X$_{i+3}$ and X$_i$–A-*cis*-P–X$_{i+3}$.** $C_{\alpha,i}$ •••$C_{\alpha,i+3}$ distances were examined at Ser-*cis*-Pro and Ala-*cis*-Pro as a determinant of type VI β-turns centered on these sequences. Distances below 7.0 Å were considered to be β-turns. By these standards, Ser-*cis*-Pro was a β-turn in 92% of structures and Ala-*cis*-Pro was a β-turn in 78% of structures.

**Table S5. Frequency of conformations observed at Ser-*cis*-Pro structures containing C–H/O interactions**. PDB structures containing Ser-*cis*-Pro sequences with C–H/O interactions analyzed by conformations at each residue. The data set of 116 Ser-*cis*-Pro structures was limited to a Pro $C_\alpha$•••Ser $O_\gamma$ distance shorter than 3.8 Å. The search resulted in 52 total structures. Populations of combinations of conformations are represented as a percent of structures containing Ser-*cis*-Pro sequences with C–H/O interactions (%$_{total}$).

| conformation | | |
|---|---|---|
| Ser - *cis* | - Pro | %$_{total}$ |
| β | $\alpha_R$ | 2 |
| β | δ | 16 |
| β | PPII | 39 |
| PPII | δ | 25 |
| PPII | PPII | 12 |
| PPII | undefined | 6 |

### Analysis of noncovalent interactions at Ser-*trans*-Pro

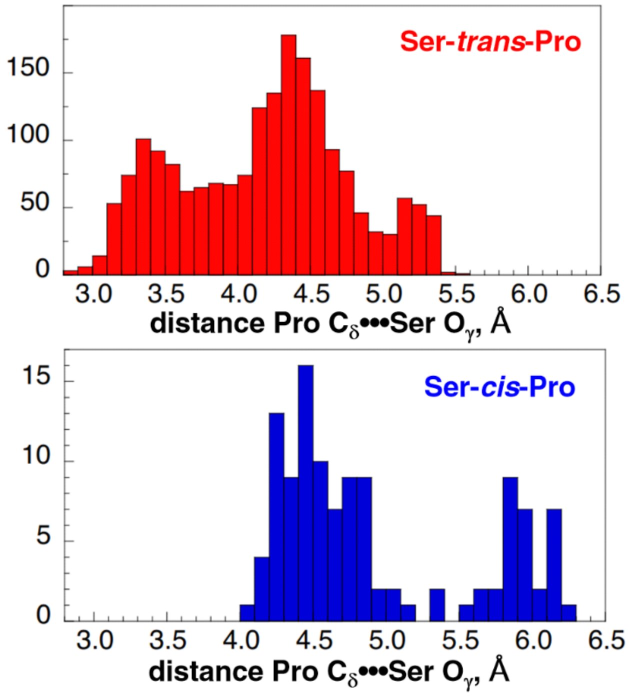

**Figure S11. Quantifying potential C–H/O interactions with Pro $H_\delta$ at Ser-*trans*-Pro in the PDB**. Structures in the PDB containing Ser-Pro sequences were analyzed by Pro $C_\delta$•••Ser $O_\gamma$ distance as a metric of potential $C_\delta$–H/O interactions. Pro $C_\delta$•••Ser $O_\gamma$ distances were used to identify a potential $C_\delta$–H/O interaction at both Ser-*trans*-Pro and Ser-*cis*-Pro structures. Potential C–H/O interactions were limited to distances below 3.8 Å, the sum of the van der Waals radii of oxygen and hydrogen and the average C–H bond length. Using these parameters, 28% of structures containing *trans*-proline are potentially in $C_\delta$–H/O interactions. No structures containing *cis*-proline were observed with Pro $C_\delta$•••Ser $O_\gamma$ distances lower than 3.8 Å.

**Analysis of main-chain hydrogen-bonded secondary structures at Ser-Pro and Ala-Pro sequences**

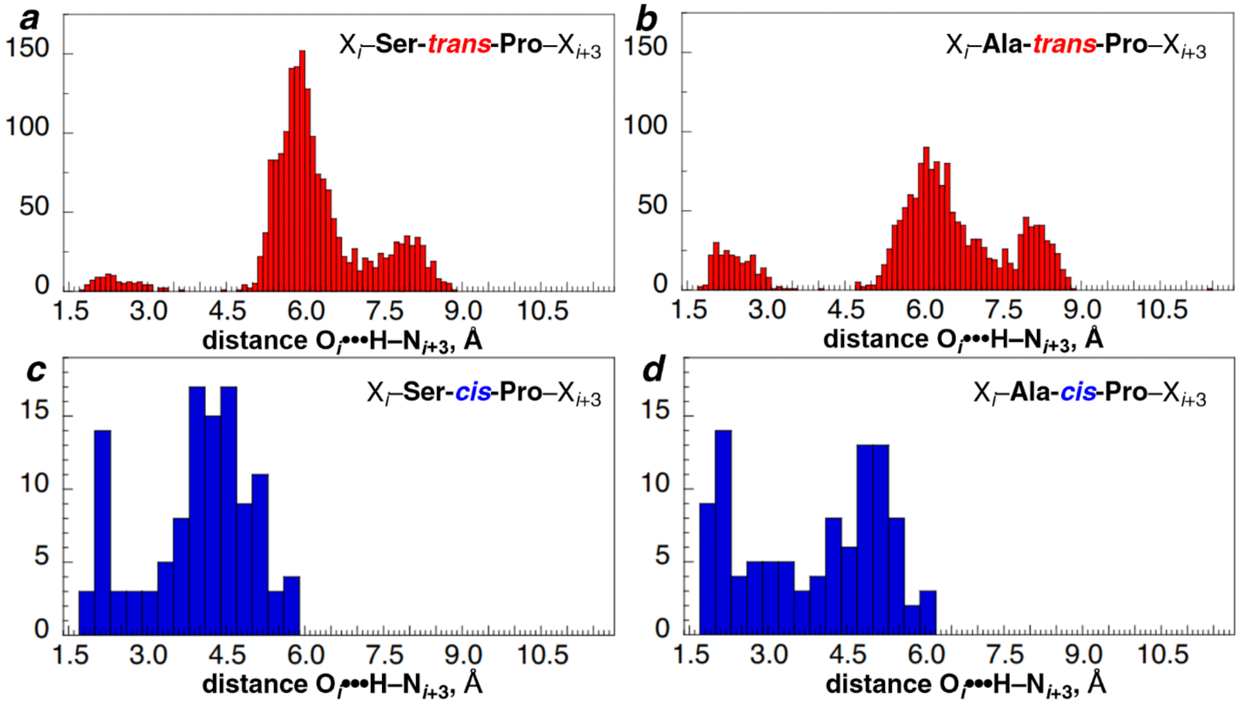

**Figure S12. Quantification of main-chain *i*/*i*+3 $O_i$•••H–$N_{i+3}$ hydrogen bonds at Ser-Pro and Ala-Pro sequences in the PDB, with Ser/Ala as the *i*+1 residue ($X_i$–(Ser/Ala)-Pro–$X_{i+3}$ register)**. Structures in the PDB containing Ser-Pro and Ala-Pro sequences were analyzed for $O_i$•••H–$N_{i+3}$ hydrogen bonds at (a) $X_i$–S-*trans*-P–$X_{i+3}$, (b) $X_i$–A-*trans*-P–$X_{i+3}$, (c) $X_i$–S-*cis*-P–$X_{i+3}$, and (d) $X_i$–A-*trans*-P–$X_{i+3}$ to identify hydrogen-bonded β-turns and $3_{10}$-helices with Ser/Ala as the *i*+1 residue and Pro as the *i*+2 residue. X is used to indicate any amino acid. A hydrogen bond is defined as an $O_i$•••H–$N_{i+3}$ distance below 2.7 Å, the sum of the van der Waals radii of H and O. Amide hydrogen positions were estimated during data refinement. $X_i$–S-*trans*-P–$X_{i+3}$ sequences have an $O_i$•••H–$N_{i+3}$ hydrogen bond in 3.6% of structures. $X_i$–A-*trans*-P–$X_{i+3}$ sequences have an $O_i$•••H–$N_{i+3}$ hydrogen bond in 10.5% of structures. $X_i$–S-*cis*-P–$X_{i+3}$ sequences have an $O_i$•••H–$N_{i+3}$ hydrogen bond in 18% of structures. $X_i$–A-*cis*-P–$X_{i+3}$ sequences have an $O_i$•••H–$N_{i+3}$ hydrogen bond in 27% of structures.

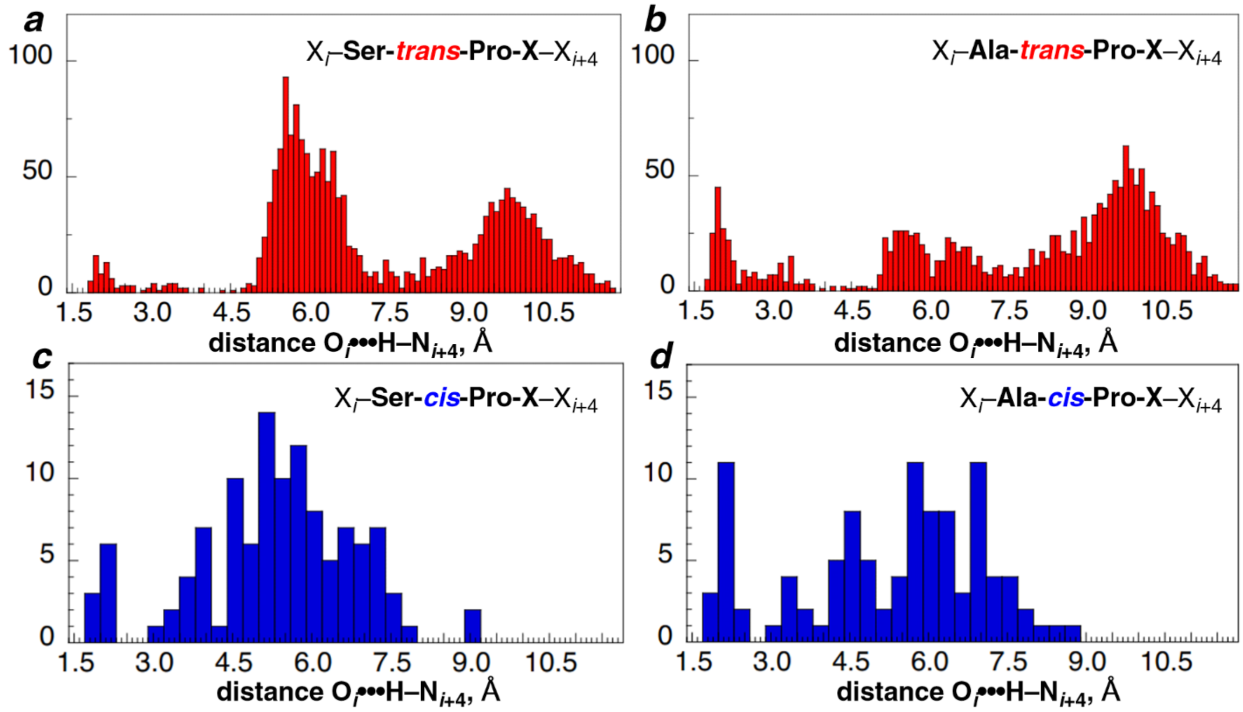

**Figure S13. Quantification of main-chain *i*/*i*+4 $O_i$•••H–$N_{i+4}$ hydrogen bonds at Ser-Pro and Ala-Pro sequences in the PDB, with Ser/Ala as the *i*+1 residue ($X_i$–(Ser/Ala)-Pro-X–$X_{i+4}$ register).** PDB structures containing Ser-Pro and Ala-Pro sequences were analyzed for $O_i$•••H–$N_{i+4}$ hydrogen bonds at (a) $X_i$–S-*trans*-P-X–$X_{i+4}$, (b) $X_i$–A-*trans*-P-X–$X_{i+4}$, (c) $X_i$–S-*cis*-P-X–$X_{i+4}$, and (d) $X_i$–A-*trans*-P-X–$X_{i+4}$ to identify α-helices and α-turns with Ser/Ala as the *i*+1 residue and Pro as the *i*+2 residue. $X_i$–S-*trans*-P-X–$X_{i+4}$ sequences have an $O_i$•••H–$N_{i+4}$ hydrogen bond in 3.1% of structures. $X_i$–A-*trans*-P-X–$X_{i+4}$ sequences have an $O_i$•••H–$N_{i+4}$ hydrogen bond in 9.4% of structures. $X_i$–S-*cis*-P-X–$X_{i+4}$ sequences have an $O_i$•••H–$N_{i+4}$ hydrogen bond in 8% of structures. $X_i$–A-*cis*-P-X–$X_{i+4}$ sequences have an $O_i$•••H–$N_{i+4}$ hydrogen bond in 16% of structures.

**Table S6. Frequencies of main-chain *i*/*i*+3 $O_i$•••H–$N_{i+3}$ and *i*/*i*+4 $O_i$•••H–$N_{i+4}$ hydrogen bonds at Ser-Pro and Ala-Pro sequences, with Ser/Ala as the *i*+1 residue ($X_i$–(Ser/Ala)-Pro–$X_{i+3}$ register).** Percentages of Ser-Pro and Ala-Pro sequences in the PDB containing $O_i$•••H–$N_{i+3}$ and $O_i$•••H–$N_{i+4}$ backbone hydrogen bonds with Ser/Ala as the *i*+1 residue and Pro as the *i*+2 residue. Populations of structures in this register containing each hydrogen bond ($\%_{total}$ $O_i$•••H–$N_{i+3}$, $\%_{total}$ $O_i$•••H–$N_{i+4}$), both hydrogen bonds ($\%_{total}$ both), and any combination of those hydrogen bonds ($\%_{total}$ any) are reported. Structures are separated by Pro amide conformation.

| Register | $\%_{total}$ $O_i$•••H–$N_{i+3}$ | $\%_{total}$ $O_i$•••H–$N_{i+4}$ | $\%_{total}$ both | $\%_{total}$ any |
|---|---|---|---|---|
| **XS-*trans*-PX** | 3.6 | 3.1 | 2.0 | 4.7 |
| **XA-*trans*-PX** | 10.5 | 9.4 | 6.1 | 13.8 |
| **XS-*cis*-PX** | 18 | 8 | 2 | 24 |
| **XA-*cis*-PX** | 27 | 16 | 6 | 37 |

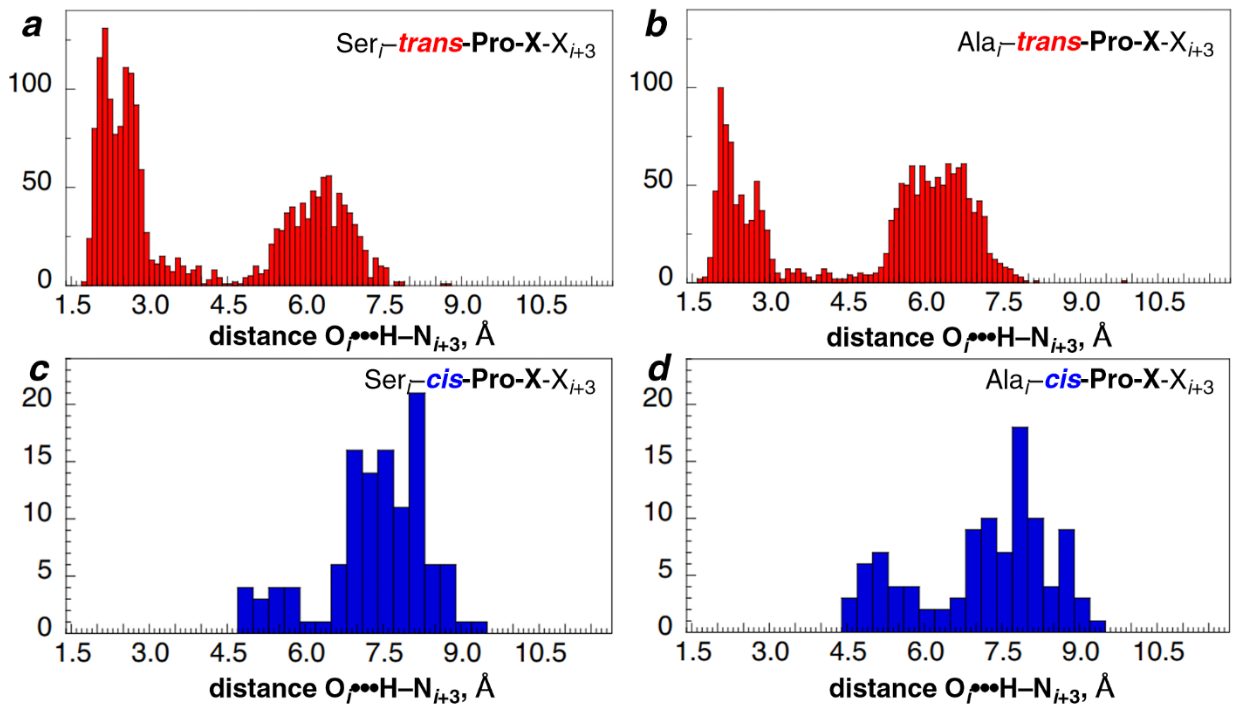

**Figure S14. Quantification of main-chain *i*/*i*+3 O$_i$•••H–N$_{i+3}$ hydrogen bonds at Ser-Pro and Ala-Pro sequences in the PDB, with Ser/Ala as the *i* residue (Ser$_i$/Ala$_i$–Pro-X–X$_{i+3}$ register)**. PDB structures containing Ser-Pro and Ala-Pro sequences were analyzed for O$_i$•••H–N$_{i+3}$ hydrogen bonds at (a) S$_i$–*trans*-P-X–X$_{i+3}$, (b) A$_i$–*trans*-P-X–X$_{i+3}$, (c) S$_i$–*cis*-P-X–X$_{i+3}$, and (d) A$_i$–*trans*-P-X–X$_{i+3}$ to identify hydrogen-bonded β-turns and $3_{10}$-helices with Ser/Ala as the *i* residue and Pro as the *i*+1 residue. S$_i$–*trans*-P-X–X$_{i+3}$ sequences have an O$_i$•••H–N$_{i+3}$ hydrogen bond in 42.9% of structures. X$_i$–A-*trans*-P–X$_{i+3}$ sequences have an O$_i$•••H–N$_{i+3}$ hydrogen bond in 27.1% of structures. S$_i$–*cis*-P-X–X$_{i+3}$ sequences and A$_i$–*cis*-P-X–X$_{i+3}$ sequences do not have an O$_i$•••H–N$_{i+3}$ hydrogen bond in any structures.

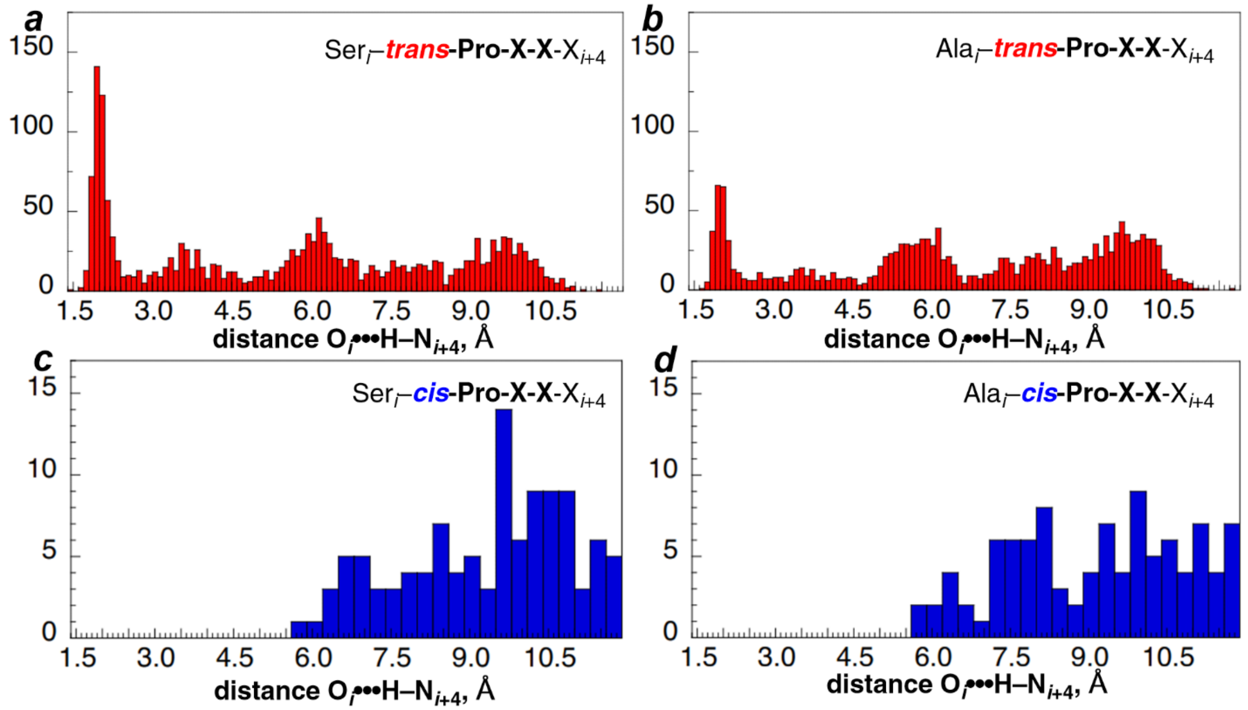

**Figure S15. Quantification of main-chain *i*/*i*+4 $O_i$•••H–$N_{i+4}$ hydrogen bonds at Ser-Pro and Ala-Pro sequences in the PDB, with Ser/Ala as the *i* residue (Ser$_i$/Ala$_i$–Pro-X-X–X$_{i+4}$ register)**. PDB structures containing Ser-Pro and Ala-Pro sequences were analyzed for $O_i$•••H–$N_{i+4}$ hydrogen bonds at (a) S$_i$–*trans*-P-X-X–X$_{i+4}$, (b) S$_i$–*trans*-P-X-X–X$_{i+4}$, (c) S$_i$–*cis*-P-X-X–X$_{i+4}$, and (d) A$_i$–*trans*-P-X-X–X$_{i+4}$ to identify α-helices and α-turns with Ser/Ala as the *i*+1 residue and Pro as the *i*+2 residue. S$_i$–*trans*-P-X-X–X$_{i+4}$ sequences have an $O_i$•••H–$N_{i+4}$ hydrogen bond in 25.2% of structures. A$_i$–*trans*-P-X-X–X$_{i+4}$ sequences have an $O_i$•••H–$N_{i+4}$ hydrogen bond in 14.3% of structures. S$_i$–*cis*-P-X-X–X$_{i+4}$ sequences and A$_i$–*cis*-P-X-X–X$_{i+4}$ sequences have no $O_i$•••H–$N_{i+4}$ hydrogen bonds in any structures.

**Table S7. Frequencies of main-chain *i*/*i*+3 $O_i$•••H–$N_{i+3}$ and *i*/*i*+4 $O_i$•••H–$N_{i+4}$ hydrogen bonds at Ser-Pro and Ala-Pro sequences, with Ser/Ala as the *i* residue (Ser$_i$/Ala$_i$–Pro-X–X$_{i+3}$ register)**. Percentages of Ser-Pro and Ala-Pro sequences in the PDB containing $O_i$•••H–$N_{i+3}$ and $O_i$•••H–$N_{i+4}$ backbone hydrogen bonds with Ser/Ala as the *i* residue and Pro as the *i*+1 residue. Populations of structures in this register containing each hydrogen bond (%$_{total}$ $O_i$•••H–$N_{i+3}$, %$_{total}$ $O_i$•••H–$N_{i+4}$), both hydrogen bonds (%$_{total}$ both), and any combination of those hydrogen bonds (%$_{total}$ any) are reported. Percentages are separated by Pro-amide isomer.

| Register | %$_{total}$ $O_i$•••H–$N_{i+3}$ | %$_{total}$ $O_i$•••H–$N_{i+4}$ | %$_{total}$ both | %$_{total}$ any |
|---|---|---|---|---|
| S-*trans*-PXX | 42.9 | 25.2 | 16.9 | 51.2 |
| A-*trans*-PXX | 27.1 | 14.3 | 7.5 | 33.9 |
| S-*cis*-PXX | 0 | 0 | 0 | 0 |
| A-*cis*-PXX | 0 | 0 | 0 | 0 |

**Table S8. Helix initiation and β-turns at $S_i$–*trans*-P-X–$X_{i+3}$ and $A_i$–*trans*-P-X–$X_{i+3}$ sequences with *i*/*i*+3 $O_i$•••H–$N_{i+3}$ hydrogen bonds**. $O_i$•••H–$N_{i+3}$ hydrogen-bonded Ser-*trans*-Pro structures in the PDB were analyzed for the presence of secondary structures and compared to Ala-*trans*-Pro. $O_i$•••H–$N_{i+3}$ hydrogen bonding can induce either a turn structure or initiate helix formation. Helix initiating structures were identified by a sequence of five or more consecutive residues adopting an $\alpha_R/\delta$ conformation following the first residue in the turn.

| $O_i$•••H–$N_{i+3}$ | $\%_{total}$ $3_{10}$ or α–helix initiating | $\%_{total}$ β-turn |
|---|---|---|
| **S-*trans*-PXX** | 17.5 | 25.2 |
| **A-*trans*-PXX** | 9.1 | 18.0 |

**Table S9. Helix initiation and β-turns at $S_i$–*trans*-P-X-X–$X_{i+4}$ and $A_i$–*trans*-P-X–$X_{i+4}$ sequences with *i*/*i*+4 $O_i$•••H–$N_{i+4}$ hydrogen bonds**. Consecutive $O_i$•••H–$N_{i+4}$ hydrogen bonding is associated with helix propagation, but a single $O_i$•••H–$N_{i+4}$ hydrogen bond is considered an α-turn. Helix initiating structures were identified by a sequence of five or more consecutive residues adopting an $\alpha_R/\delta$ conformation following the first residue in the turn.

| $O_i$•••H–$N_{i+4}$ | $\%_{total}$ α-helix initiating | $\%_{total}$ α-turn |
|---|---|---|
| **S-*trans*-PXX** | 21.6 | 3.9 |
| **A-*trans*-PXX** | 12.0 | 2.3 |

**Table S10. β-turn type populations at S-*trans*-PXX and A-*trans*-PXX**. 36.8% of S-*trans*-PXX and 18.2% of A-*trans*-PXX sequences in the PDB were found in to be in β-turns. Populations of each type of β-turn centered on *trans*-P-X are reported as a percent of S-*trans*-PXX or A-*trans*-PXX structures in any type of β-turn, respectively.

| conformation | | β-turn type | | S-*trans*-PXX | A-*trans*-PXX |
|---|---|---|---|---|---|
| Pro | X | classical | Dunbrack | % of turns | % of turns |
| $\alpha_R$ | $\alpha_R$ | I | AA | 48.2 | 17.8 |
| $\alpha_R$ | $\alpha_L$ | II | Aa | 0.0 | 0.0 |
| $\alpha_R$ | β | VIII | AB | 0.0 | 0.0 |
| $\alpha_R$ | δ | I | AD | 37.3 | 42.4 |
| $\alpha_R$ | PPII | VIII | AP | 0.0 | 0.0 |
| $\alpha_R$ | undefined | N/A | N/A | 3.7 | 4.8 |
| $\alpha_R$ | ζ | VIII | AZ | 0.8 | 1.0 |
| β | $\alpha_R$ | IV | BA | 0.0 | 0.0 |
| β | $\alpha_L$ | II | Ba | 0.0 | 0.0 |
| β | β | IV | BB | 0.0 | 0.0 |
| β | δ | IV | Bd | 0.0 | 0.0 |
| β | PPII | IV | BP | 0.0 | 0.0 |
| β | undefined | IV | N/A | 0.0 | 0.0 |
| β | ζ | IV | BZ | 0.0 | 0.0 |
| δ | $\alpha_R$ | I | DA | 0.8 | 0.0 |
| δ | $\alpha_L$ | II | Da | 0.0 | 0.0 |
| δ | β | VIII | DB | 0.0 | 0.0 |
| δ | δ | I | DD | 3.2 | 4.5 |
| δ | PPII | VIII | DP | 0.0 | 0.0 |
| δ | undefined | IV | N/A | 0.4 | 0.3 |
| δ | ζ | VIII | DZ | 0.0 | 0.0 |
| PPII | $\alpha_R$ | IV | PA | 0.0 | 0.0 |
| PPII | $\alpha_L$ | II | Pa | 4.9 | 27.1 |
| PPII | β | IV | PB | 0.1 | 0.3 |
| PPII | δ | IV | PD | 0.4 | 0.3 |
| PPII | PPII | IV | PP | 0.0 | 0.0 |
| PPII | undefined | IV | N/A | 0.0 | 1.3 |
| PPII | ζ | IV | PZ | 0.0 | 0.3 |
| undefined | $\alpha_R$ | IV | N/A | 0.0 | 0.0 |
| undefined | $\alpha_L$ | IV | N/A | 0.0 | 0.0 |
| undefined | β | IV | N/A | 0.0 | 0.0 |
| undefined | δ | IV | N/A | 0.0 | 0.0 |
| undefined | PPII | IV | N/A | 0.0 | 0.0 |
| undefined | ζ | IV | N/A | 0.0 | 0.0 |

**Table S11. Conformations of Ser and Ala in S-*trans*-PXX and A-*trans*-PXX type I β-turns and $O_i$•••H–$N_{i+4}$ hydrogen bonded helix N-termini**. The most common secondary structures at Ser-*trans*-Pro are SPXX type I β-turns (28.9%) and N-terminal α-helix caps (21.3%). Populations of conformations of Ser in both structures are reported alongside conformations of Ala in the same structures at A-*trans*-PXX.

| | type I β-turn | | α-helix capping | |
|---|---|---|---|---|
| | Ser | Ala | Ser | Ala |
| conformation | % | % | % | % |
| $\alpha_R$ | 4.6 | 27.2 | 11.9 | 62.7 |
| $\alpha_L$ | 0.0 | 1.5 | 0.2 | 0.0 |
| β | 26.8 | 10.7 | 18.9 | 2.0 |
| δ | 0.0 | 0.0 | 0.0 | 0.0 |
| PPII | 57.3 | 37.4 | 63.8 | 26.0 |
| ζ | 10.0 | 21.8 | 4.1 | 7.8 |
| undefined | 1.4 | 1.5 | 1.0 | 1.5 |

**X-ray crystallography**

X-ray structural analysis for **3** and **5**: Crystals were mounted using viscous oil onto a plastic mesh and cooled to the data collection temperature. Data were collected on a D8 Venture Photon III diffractometer with Cu-Kα radiation (λ = 1.54178 Å) focused with Goebel mirrors. Unit cell parameters were obtained from fast scan data frames, 1°/s ω, of an Ewald hemisphere. The unit-cell dimensions, equivalent reflections and systematic absences in the diffraction data are consistent with $P2_1$ and $P2_1/m$. The noncentrosymmetric space group, $P2_1$, is consistent with the enantiomerically pure compounds and the anomalous displacement parameter refined to nil in each case indicating the true hand was determined. The data were treated with multi-scan absorption corrections.[3] Structures were solved using intrinsic phasing methods[4] and refined with full-matrix, least-squares procedures on $F^2$.[5]

Two compound molecules were located in the asymmetric unit of **3**. A severely disordered molecule of acetone solvent was located in the asymmetric unit of **3** that treated as a diffused contribution using Squeeze.[6] The phenyl rings in **3** were treated as idealized hexagonal rigid groups. The pendent group on the pyrrolidine N in **5** was located disordered in two positions with refined site occupancies of 58/42. Intermolecular hydrogen bonding interactions of types OH•••OH, OH•••OC(N) and NH•••OC(N) were observed in **3** but only OH•••OC(N) and NH•••OC(N) were observed in **5**. The hydrogen atoms involved in hydrogen bonding were located in **3** and their positions were allowed to refined. The disordered, donor hydrogen atoms could not be located in **5**. In any case, no close contacts between hydroxyl oxygen atoms were found in **5**. Excepting the hydrogen bond donors in **3**, all other hydrogen atoms were treated as idealized contributions with geometrically calculated positions and with $U_{iso}$ equal to 1.2 $U_{eq}$ (1.5 $U_{eq}$ for methyl) of the attached atom.

Atomic scattering factors are contained in the SHELXTL program library.[2] The structures have been deposited at the Cambridge Structural Database under the following CCDC depositary numbers: Boc-L-serine-(2*S*,4*S*)-4-iodophenyl-hydroxyproline methyl ester CCDC 2330290 Ac-L-serine-(2*S*,4*S*)-4-iodophenyl-hydroxyproline methyl ester CCDC 2330291.

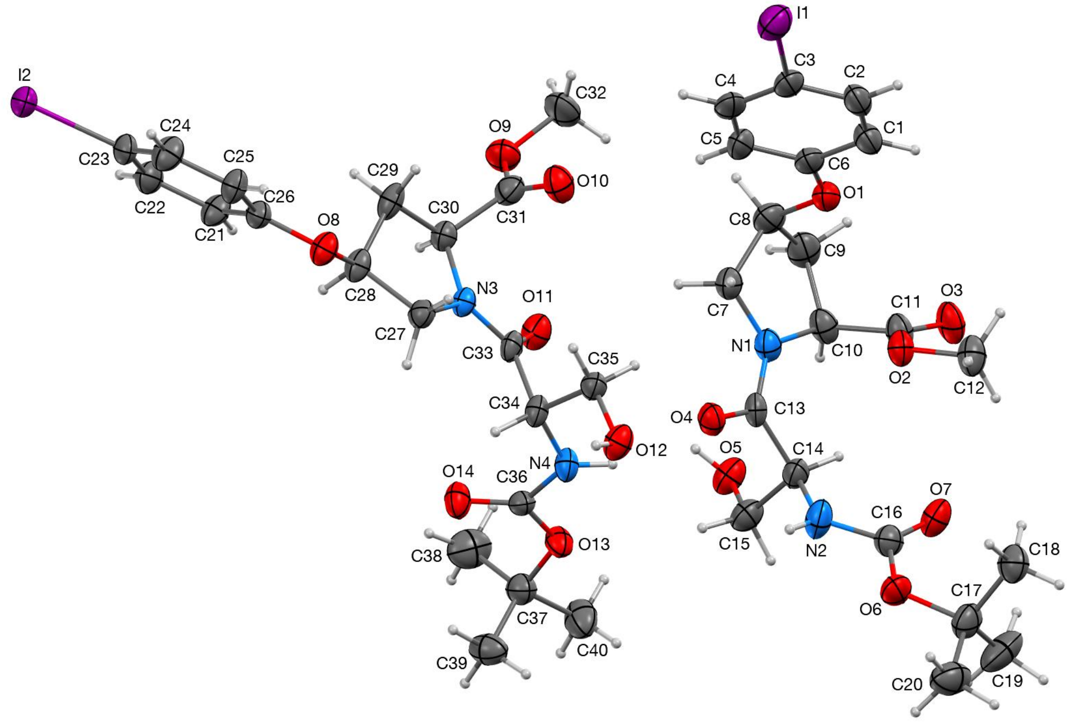

**Figure S16.** Molecular diagram of Boc-L-serine-(2*S*,4*S*)-4-iodophenyl-hydroxyproline methyl ester (**3**) showing two symmetry unique compound molecules with 50% probability ellipsoids. H-atoms depicted at arbitrary radius. Diffractable crystals were obtained by slow evaporation at room temperature from a solution of **3** in acetone. This structure has CCDC number 2330290.

**Table S12.** Crystallographic data and refinement details for **3**.

| | |
|---|---|
| Empirical formula | $C_{21.5}H_{30}IN_2O_{7.5}$ |
| Formula weight | 563.37 |
| Temperature/K | 100.00 |
| Crystal system | monoclinic |
| Space group | $P2_1$ |
| a/Å | 15.5879(11) |
| b/Å | 5.8627(4) |
| c/Å | 26.5512(18) |
| α/° | 90 |
| β/° | 90.058(3) |
| γ/° | 90 |
| Volume/Å$^3$ | 2426.4(3) |
| Z | 4 |

| | |
|---|---|
| $\varrho_{calc}$ g/cm$^3$ | 1.542 |
| $\mu$/mm$^{-1}$ | 10.761 |
| F(000) | 1144.0 |
| Crystal size/mm$^3$ | 0.194 × 0.038 × 0.02 |
| Radiation | CuK$\alpha$ ($\lambda$ = 1.54178) |
| 2$\Theta$ range for data collection/° | 3.328 to 140.758 |
| Index ranges | -19 ≤ h ≤ 19, -7 ≤ k ≤ 7, -32 ≤ [illegible] |
| Reflections collected | 33599 |
| Independent reflections | 8925 [$R_{int}$ = 0.0630, $R_{sigma}$ = 0.[illegible] |
| Data/restraints/parameters | 8925/4/536 |
| Goodness-of-fit on $F^2$ | 1.066 |
| Final R indexes [I>=2σ (I)] | $R_1$ = 0.0554, $wR_2$ = 0.1393 |
| Final R indexes [all data] | $R_1$ = 0.0674, $wR_2$ = 0.1465 |
| Largest diff. peak/hole / e Å$^{-3}$ | 0.81/-0.68 |
| Flack parameter | -0.010(6) |

**Table S13.** Fractional Atomic Coordinates (×10$^4$) and Equivalent Isotropic Displacement Parameters (Å$^2$×10$^3$) for **3**. $U_{eq}$ is defined as 1/3 of the trace of the orthogonalised $U_{IJ}$ tensor.

| **Atom** | ***x*** | ***y*** | ***z*** | **U(eq)** |
|---|---|---|---|---|
| I1 | 697.2(5) | -315(2) | 9609.6(3) | 50.3(2) |
| O1 | 3333(5) | 7464(14) | 9173(3) | 38.1(18) |
| O2 | 5321(4) | 6401(15) | 9127(2) | 35.4(15) |
| O3 | 5552(6) | 9842(15) | 9456(3) | 47(2) |
| O4 | 4897(5) | 5141(13) | 7739(3) | 36.0(17) |
| O5 | 5958(5) | 10804(13) | 7542(3) | 42.1(19) |
| O6 | 7665(5) | 3065(15) | 8324(3) | 41.3(18) |
| O7 | 7324(5) | 6260(17) | 8758(3) | 43.1(18) |
| N1 | 4609(6) | 7922(16) | 8289(3) | 33.4(19) |
| N2 | 6591(7) | 5209(17) | 8052(4) | 44(2) |
| C1 | 2784(4) | 4621(13) | 9685(2) | 40(2) |
| C2 | 2204(5) | 2883(13) | 9789(2) | 41(3) |
| C3 | 1589(4) | 2278(11) | 9434(3) | 38(3) |
| C4 | 1554(4) | 3410(13) | 8974(2) | 44(3) |
| C5 | 2134(5) | 5148(13) | 8870(2) | 42(3) |
| C6 | 2749(4) | 5753(12) | 9226(3) | 38(3) |
| C7 | 3691(7) | 7420(20) | 8273(4) | 37(2) |
| C8 | 3318(8) | 8790(20) | 8719(5) | 42(3) |
| C9 | 3971(8) | 10700(20) | 8787(5) | 45(3) |
| C10 | 4837(7) | 9630(20) | 8667(4) | 37(2) |
| C11 | 5278(8) | 8642(19) | 9133(4) | 36(2) |
| C12 | 5753(8) | 5380(20) | 9559(4) | 44(3) |
| C13 | 5152(6) | 6695(19) | 8014(3) | 31(2) |
| C14 | 6101(7) | 7295(19) | 8029(4) | 34(2) |
| C15 | 6345(8) | 8630(20) | 7554(5) | 42(3) |
| C16 | 7214(7) | 4940(20) | 8414(4) | 36(2) |
| C17 | 8368(8) | 2400(20) | 8667(5) | 43(3) |
| C18 | 7983(8) | 1690(20) | 9166(4) | 46(3) |
| C19 | 9012(8) | 4270(30) | 8739(7) | 65(4) |
| C20 | 8763(9) | 400(30) | 8405(6) | 57(4) |
| I2 | -2722.7(4) | 1415.21 | 4673.0(2) | 36.12(19) |
| O8 | 559(5) | 872(12) | 6024(3) | 32.8(16) |
| O9 | 1083(5) | 8616(14) | 7300(3) | 45(2) |
| O10 | 1505(6) | 5236(13) | 7611(3) | 43.3(19) |
| O11 | 3052(5) | 6874(12) | 6927(3) | 36.2(17) |
| O12 | 4283(5) | 12(13) | 7184(3) | 37.7(17) |
| O13 | 5417(4) | 6725(14) | 6342(2) | 34.5(16) |
| O14 | 4298(5) | 5331(14) | 5889(3) | 39.6(19) |

| | | | | |
|---|---|---|---|---|
| N3 | 2151(5) | 4241(13) | 6634(3) | 25.1(17) |
| N4 | 4493(5) | 4371(16) | 6702(3) | 32.2(19) |
| C21 | -342(4) | 3112(10) | 5446(3) | 33(2) |
| C22 | -1074(4) | 3194(10) | 5147(2) | 37(2) |
| C23 | -1631(3) | 1343(12) | 5134(2) | 32(2) |
| C24 | -1456(4) | -588(10) | 5419(3) | 40(3) |
| C25 | -724(4) | -670(9) | 5718(3) | 32(2) |
| C26 | -167(3) | 1181(11) | 5731(2) | 29(2) |
| C27 | 1902(6) | 2129(17) | 6374(4) | 30(2) |
| C28 | 1055(6) | 2832(17) | 6136(4) | 30(2) |
| C29 | 660(7) | 4380(19) | 6532(4) | 35(2) |
| C30 | 1424(7) | 5762(17) | 6712(4) | 30(2) |
| C31 | 1361(6) | 6460(20) | 7263(4) | 36(2) |
| C32 | 990(9) | 9460(30) | 7810(4) | 56(3) |
| C33 | 2945(6) | 4956(18) | 6762(4) | 28(2) |
| C34 | 3676(6) | 3247(18) | 6726(4) | 28(2) |
| C35 | 3644(6) | 1700(20) | 7194(4) | 30(2) |
| C36 | 4710(6) | 5491(18) | 6271(4) | 28(2) |
| C37 | 5781(7) | 8020(20) | 5920(4) | 38(2) |
| C38 | 5151(9) | 9660(30) | 5706(6) | 59(3) |
| C39 | 6129(7) | 6320(30) | 5542(4) | 46(3) |
| C40 | 6525(8) | 9310(30) | 6159(5) | 53(3) |

**Table S14.** Anisotropic Displacement Parameters ($Å^2 \times 10^3$) for **3**. The Anisotropic displacement factor exponent takes the form: $-2\pi^2[h^2a^{*2}U_{11}+2hka^*b^*U_{12}+\ldots]$.

| **Atom** | **$U_{11}$** | **$U_{22}$** | **$U_{33}$** | **$U_{23}$** | **$U_{13}$** | **$U_{12}$** |
|---|---|---|---|---|---|---|
| I1 | 40.4(4) | 54.2(5) | 56.4(5) | 0.0(4) | -7.9(3) | 6.9(4) |
| O1 | 35(4) | 51(5) | 29(4) | 1(3) | -1(3) | 10(4) |
| O2 | 42(4) | 34(4) | 30(3) | 0(4) | -9(3) | -4(4) |
| O3 | 68(5) | 39(4) | 33(4) | -15(4) | -12(4) | 4(4) |
| O4 | 42(4) | 36(4) | 30(4) | -7(3) | -6(3) | 4(3) |
| O5 | 36(4) | 36(4) | 54(5) | 3(4) | -11(4) | 1(3) |
| O6 | 38(4) | 47(5) | 39(4) | -2(4) | -4(3) | 16(4) |
| O7 | 37(4) | 41(4) | 51(4) | 3(5) | -15(3) | 2(4) |
| N1 | 40(5) | 32(5) | 29(4) | 0(4) | -6(4) | 4(4) |
| N2 | 46(6) | 43(6) | 43(6) | -4(4) | -20(5) | 4(5) |
| C1 | 45(6) | 51(7) | 26(5) | -3(5) | 0(4) | 6(6) |
| C2 | 49(7) | 48(7) | 25(5) | -3(5) | 0(5) | 16(6) |
| C3 | 31(5) | 46(6) | 37(6) | 1(5) | -2(5) | 16(5) |
| C4 | 31(6) | 61(8) | 41(6) | -14(6) | -3(5) | 20(6) |
| C5 | 35(6) | 58(8) | 32(6) | 3(5) | -6(5) | 15(6) |
| C6 | 31(5) | 50(7) | 33(5) | 0(5) | 1(4) | 15(5) |
| C7 | 35(6) | 46(6) | 30(5) | 6(5) | -5(4) | 4(5) |
| C8 | 37(6) | 45(7) | 42(6) | 4(5) | 0(5) | 14(5) |
| C9 | 55(7) | 38(6) | 43(6) | -2(5) | -5(5) | 18(5) |
| C10 | 51(6) | 29(5) | 31(5) | -2(5) | -3(5) | 5(5) |
| C11 | 49(7) | 31(6) | 28(5) | -4(4) | -6(5) | 1(5) |
| C12 | 53(7) | 42(7) | 36(6) | 8(5) | -11(5) | -7(5) |
| C13 | 37(5) | 34(6) | 22(4) | 1(5) | -10(4) | -1(5) |
| C14 | 32(5) | 40(6) | 30(5) | -3(4) | -7(4) | 8(4) |
| C15 | 34(6) | 44(7) | 48(7) | 3(5) | -4(5) | 7(5) |
| C16 | 37(6) | 37(6) | 33(5) | 6(5) | -1(4) | 4(5) |
| C17 | 39(6) | 47(7) | 43(7) | 10(5) | -9(5) | 8(5) |
| C18 | 46(6) | 47(7) | 44(6) | -7(6) | -9(5) | -8(6) |
| C19 | 32(6) | 71(10) | 94(11) | 18(9) | -14(7) | 0(7) |
| C20 | 47(7) | 63(9) | 61(8) | 15(7) | 9(6) | 17(6) |
| I2 | 29.3(3) | 48.9(4) | 30.2(3) | -2.4(3) | -6.1(2) | 1.3(3) |
| O8 | 31(4) | 26(4) | 41(4) | 0(3) | -10(3) | -5(3) |
| O9 | 46(5) | 45(5) | 43(5) | -8(4) | 1(4) | 11(4) |
| O10 | 57(5) | 39(5) | 34(4) | 2(3) | -5(4) | 5(4) |
| O11 | 39(4) | 25(4) | 45(4) | -5(3) | -10(3) | -1(3) |
| O12 | 37(4) | 30(4) | 46(4) | 4(3) | -11(3) | 0(3) |
| O13 | 35(4) | 43(4) | 26(3) | 4(3) | 0(3) | -11(3) |

| | | | | | | |
|---|---|---|---|---|---|---|
| O14 | 43(4) | 47(5) | 29(4) | 10(3) | -8(3) | -7(3) |
| N3 | 26(4) | 22(4) | 27(4) | 2(3) | -5(3) | -3(3) |
| N4 | 34(4) | 35(5) | 28(4) | 1(4) | -9(4) | -5(4) |
| C21 | 28(5) | 31(5) | 42(6) | 3(5) | -5(4) | -6(4) |
| C22 | 39(6) | 39(6) | 33(6) | 6(5) | -4(5) | 5(5) |
| C23 | 29(5) | 39(5) | 27(4) | -5(5) | -7(4) | -3(5) |
| C24 | 35(6) | 34(6) | 50(7) | 2(5) | -6(5) | -9(5) |
| C25 | 34(5) | 27(5) | 37(6) | 1(4) | -11(4) | -2(4) |
| C26 | 26(4) | 34(5) | 28(4) | -3(5) | -3(4) | -3(5) |
| C27 | 26(5) | 33(5) | 32(5) | -4(4) | -3(4) | -2(4) |
| C28 | 30(5) | 22(5) | 36(5) | 2(4) | -9(4) | -7(4) |
| C29 | 31(5) | 28(5) | 45(6) | 4(5) | -15(5) | -5(4) |
| C30 | 29(5) | 31(5) | 29(5) | 2(4) | -4(4) | 0(4) |
| C31 | 28(5) | 39(5) | 42(6) | -3(6) | -6(4) | -1(5) |
| C32 | 64(8) | 65(9) | 38(7) | -15(6) | 4(6) | 23(7) |
| C33 | 28(5) | 29(5) | 26(5) | 9(4) | -8(4) | -1(4) |
| C34 | 27(5) | 28(5) | 28(5) | 4(4) | -8(4) | -4(4) |
| C35 | 28(5) | 35(6) | 29(5) | -4(5) | -3(4) | 4(5) |
| C36 | 23(5) | 35(5) | 28(5) | -3(4) | 1(4) | 0(4) |
| C37 | 35(6) | 43(6) | 35(6) | -5(5) | 0(4) | -1(5) |
| C38 | 58(8) | 47(7) | 71(9) | 21(8) | 9(7) | 4(7) |
| C39 | 42(6) | 46(7) | 50(6) | -15(7) | 9(5) | -7(7) |
| C40 | 48(7) | 65(9) | 45(7) | 0(6) | -6(6) | -23(7) |

**Table S15.** Bond lengths for **3**.

| **Atom** | **Atom** | **Length, Å** | **Atom** | **Atom** | **Length, Å** |
|---|---|---|---|---|---|
| I1 | C3 | 2.112(6) | I2 | C23 | 2.095(4) |
| O1 | C6 | 1.362(10) | O8 | C26 | 1.385(8) |
| O1 | C8 | 1.435(14) | O8 | C28 | 1.415(12) |
| O2 | C11 | 1.316(14) | O9 | C31 | 1.338(16) |
| O2 | C12 | 1.460(12) | O9 | C32 | 1.448(14) |
| O3 | C11 | 1.189(13) | O10 | C31 | 1.193(14) |
| O4 | C13 | 1.234(13) | O11 | C33 | 1.218(13) |
| O5 | C15 | 1.408(14) | O12 | C35 | 1.406(13) |
| O6 | C16 | 1.327(14) | O13 | C36 | 1.332(12) |
| O6 | C17 | 1.476(13) | O13 | C37 | 1.468(14) |
| O7 | C16 | 1.210(14) | O14 | C36 | 1.203(12) |
| N1 | C7 | 1.461(14) | N3 | C27 | 1.469(12) |
| N1 | C10 | 1.464(14) | N3 | C30 | 1.458(13) |
| N1 | C13 | 1.329(14) | N3 | C33 | 1.350(12) |
| N2 | C14 | 1.443(14) | N4 | C34 | 1.436(13) |
| N2 | C16 | 1.374(14) | N4 | C36 | 1.362(13) |
| C1 | C2 | 1.390 | C21 | C22 | 1.390 |
| C1 | C6 | 1.390 | C21 | C26 | 1.390 |
| C2 | C3 | 1.390 | C22 | C23 | 1.390 |
| C3 | C4 | 1.390 | C23 | C24 | 1.390 |
| C4 | C5 | 1.390 | C24 | C25 | 1.390 |
| C5 | C6 | 1.390 | C25 | C26 | 1.390 |
| C7 | C8 | 1.546(16) | C27 | C28 | 1.521(13) |
| C8 | C9 | 1.523(18) | C28 | C29 | 1.520(16) |
| C9 | C10 | 1.520(16) | C29 | C30 | 1.517(14) |
| C10 | C11 | 1.528(14) | C30 | C31 | 1.524(14) |
| C13 | C14 | 1.521(15) | C33 | C34 | 1.520(14) |
| C14 | C15 | 1.532(17) | C34 | C35 | 1.538(14) |
| C17 | C18 | 1.513(17) | C37 | C38 | 1.485(18) |
| C17 | C19 | 1.495(19) | C37 | C39 | 1.516(17) |
| C17 | C20 | 1.501(19) | C37 | C40 | 1.522(16) |

**Table S16.** Bond angles for **3**.

| Atom | Atom | Atom | Angle, ° | Atom | Atom | Atom | Angle, ° |
|---|---|---|---|---|---|---|---|
| C6 | O1 | C8 | 118.4(8) | C26 | O8 | C28 | 117.2(7) |
| C11 | O2 | C12 | 115.2(8) | C31 | O9 | C32 | 115.0(10) |
| C16 | O6 | C17 | 120.0(9) | C36 | O13 | C37 | 119.6(8) |
| C7 | N1 | C10 | 113.3(9) | C30 | N3 | C27 | 112.1(7) |
| C13 | N1 | C7 | 120.0(9) | C33 | N3 | C27 | 128.5(8) |
| C13 | N1 | C10 | 126.4(9) | C33 | N3 | C30 | 119.2(8) |
| C16 | N2 | C14 | 120.1(10) | C36 | N4 | C34 | 118.7(8) |
| C2 | C1 | C6 | 120.0 | C22 | C21 | C26 | 120.0 |
| C1 | C2 | C3 | 120.0 | C23 | C22 | C21 | 120.0 |
| C2 | C3 | I1 | 119.2(4) | C22 | C23 | I2 | 120.4(4) |
| C4 | C3 | I1 | 120.8(4) | C22 | C23 | C24 | 120.0 |
| C4 | C3 | C2 | 120.0 | C24 | C23 | I2 | 119.6(3) |
| C5 | C4 | C3 | 120.0 | C23 | C24 | C25 | 120.0 |
| C6 | C5 | C4 | 120.0 | C26 | C25 | C24 | 120.0 |
| O1 | C6 | C1 | 114.6(5) | O8 | C26 | C21 | 125.0(5) |
| O1 | C6 | C5 | 125.4(5) | O8 | C26 | C25 | 115.0(5) |
| C5 | C6 | C1 | 120.0 | C25 | C26 | C21 | 120.0 |
| N1 | C7 | C8 | 104.1(9) | N3 | C27 | C28 | 101.4(8) |
| O1 | C8 | C7 | 110.7(9) | O8 | C28 | C27 | 109.9(8) |
| O1 | C8 | C9 | 106.8(10) | O8 | C28 | C29 | 114.1(9) |
| C9 | C8 | C7 | 102.7(10) | C29 | C28 | C27 | 103.0(8) |
| C10 | C9 | C8 | 105.6(10) | C30 | C29 | C28 | 102.6(8) |
| N1 | C10 | C9 | 102.1(9) | N3 | C30 | C29 | 103.8(8) |
| N1 | C10 | C11 | 113.7(9) | N3 | C30 | C31 | 110.6(8) |
| C9 | C10 | C11 | 112.7(10) | C29 | C30 | C31 | 113.3(9) |
| O2 | C11 | C10 | 113.2(9) | O9 | C31 | C30 | 110.3(9) |
| O3 | C11 | O2 | 125.5(10) | O10 | C31 | O9 | 124.9(10) |
| O3 | C11 | C10 | 121.3(10) | O10 | C31 | C30 | 124.7(11) |
| O4 | C13 | N1 | 121.3(10) | O11 | C33 | N3 | 120.1(10) |
| O4 | C13 | C14 | 119.9(10) | O11 | C33 | C34 | 122.0(9) |
| N1 | C13 | C14 | 118.8(9) | N3 | C33 | C34 | 117.8(9) |
| N2 | C14 | C13 | 108.7(9) | N4 | C34 | C33 | 111.4(9) |
| N2 | C14 | C15 | 109.8(9) | N4 | C34 | C35 | 109.6(8) |
| C13 | C14 | C15 | 109.9(9) | C33 | C34 | C35 | 108.2(8) |
| O5 | C15 | C14 | 112.1(10) | O12 | C35 | C34 | 112.1(8) |
| O6 | C16 | N2 | 110.1(10) | O13 | C36 | N4 | 110.4(8) |
| O7 | C16 | O6 | 126.2(10) | O14 | C36 | O13 | 127.1(10) |
| O7 | C16 | N2 | 123.7(11) | O14 | C36 | N4 | 122.5(9) |

| | | | | | | | |
|---|---|---|---|---|---|---|---|
| O6 | C17 | C18 | 108.5(10) | O13 | C37 | C38 | 111.7(9) |
| O6 | C17 | C19 | 112.6(10) | O13 | C37 | C39 | 107.6(10) |
| O6 | C17 | C20 | 102.9(10) | O13 | C37 | C40 | 103.5(9) |
| C19 | C17 | C18 | 111.0(11) | C38 | C37 | C39 | 114.1(11) |
| C19 | C17 | C20 | 110.8(12) | C38 | C37 | C40 | 110.0(12) |
| C20 | C17 | C18 | 110.7(11) | C39 | C37 | C40 | 109.2(10) |

**Table S17.** Torsion angles for **3**.

| A | B | C | D | Angle, ° | A | B | C | D | Angle, ° |
|---|---|---|---|---|---|---|---|---|---|
| I1 | C3 | C4 | C5 | 178.5(5) | I2 | C23 | C24 | C25 | 178.7(5) |
| O1 | C8 | C9 | C10 | 81.3(11) | O8 | C28 | C29 | C30 | 160.1(8) |
| O4 | C13 | C14 | N2 | -44.2(13) | O11 | C33 | C34 | N4 | -24.0(13) |
| O4 | C13 | C14 | C15 | 75.9(12) | O11 | C33 | C34 | C35 | 96.6(11) |
| N1 | C7 | C8 | O1 | -89.0(11) | N3 | C27 | C28 | O8 | -158.7(8) |
| N1 | C7 | C8 | C9 | 24.7(11) | N3 | C27 | C28 | C29 | -36.8(10) |
| N1 | C10 | C11 | O2 | -4.5(14) | N3 | C30 | C31 | O9 | 144.8(9) |
| N1 | C10 | C11 | O3 | 174.9(11) | N3 | C30 | C31 | O10 | -38.1(14) |
| N1 | C13 | C14 | N2 | 138.1(10) | N3 | C33 | C34 | N4 | 160.1(8) |
| N1 | C13 | C14 | C15 | -101.8(11) | N3 | C33 | C34 | C35 | -79.4(10) |
| N2 | C14 | C15 | O5 | -173.2(9) | N4 | C34 | C35 | O12 | -60.3(11) |
| C1 | C2 | C3 | I1 | -178.5(5) | C21 | C22 | C23 | I2 | -178.7(5) |
| C1 | C2 | C3 | C4 | 0.0 | C21 | C22 | C23 | C24 | 0.0 |
| C2 | C1 | C6 | O1 | 178.2(7) | C22 | C21 | C26 | O8 | 177.0(7) |
| C2 | C1 | C6 | C5 | 0.0 | C22 | C21 | C26 | C25 | 0.0 |
| C2 | C3 | C4 | C5 | 0.0 | C22 | C23 | C24 | C25 | 0.0 |
| C3 | C4 | C5 | C6 | 0.0 | C23 | C24 | C25 | C26 | 0.0 |
| C4 | C5 | C6 | O1 | -178.0(8) | C24 | C25 | C26 | O8 | -177.3(7) |
| C4 | C5 | C6 | C1 | 0.0 | C24 | C25 | C26 | C21 | 0.0 |
| C6 | O1 | C8 | C7 | -74.5(11) | C26 | O8 | C28 | C27 | -169.7(8) |
| C6 | O1 | C8 | C9 | 174.5(8) | C26 | O8 | C28 | C29 | 75.3(10) |
| C6 | C1 | C2 | C3 | 0.0 | C26 | C21 | C22 | C23 | 0.0 |
| C7 | N1 | C10 | C9 | -16.0(12) | C27 | N3 | C30 | C29 | 5.7(11) |
| C7 | N1 | C10 | C11 | 105.7(11) | C27 | N3 | C30 | C31 | 127.6(9) |
| C7 | N1 | C13 | O4 | 0.8(15) | C27 | N3 | C33 | O11 | 171.8(9) |
| C7 | N1 | C13 | C14 | 178.5(9) | C27 | N3 | C33 | C34 | -12.2(14) |
| C7 | C8 | C9 | C10 | -35.2(11) | C27 | C28 | C29 | C30 | 41.1(10) |
| C8 | O1 | C6 | C1 | -177.2(7) | C28 | O8 | C26 | C21 | 15.5(11) |
| C8 | O1 | C6 | C5 | 0.9(11) | C28 | O8 | C26 | C25 | -167.3(7) |
| C8 | C9 | C10 | N1 | 31.5(12) | C28 | C29 | C30 | N3 | -28.7(10) |
| C8 | C9 | C10 | C11 | -90.9(11) | C28 | C29 | C30 | C31 | -148.7(9) |
| C9 | C10 | C11 | O2 | 111.2(12) | C29 | C30 | C31 | O9 | -99.1(11) |
| C9 | C10 | C11 | O3 | -69.5(15) | C29 | C30 | C31 | O1 | 078.0(13) |
| C10 | N1 | C7 | C8 | -5.5(12) | C30 | N3 | C27 | C28 | 19.6(11) |
| C10 | N1 | C13 | O4 | 173.1(10) | C30 | N3 | C33 | O11 | -2.1(14) |
| C10 | N1 | C13 | C14 | -9.2(15) | C30 | N3 | C33 | C34 | 173.9(8) |
| C12 | O2 | C11 | O3 | -0.5(18) | C32 | O9 | C31 | O1 | 02.2(16) |
| C12 | O2 | C11 | C10 | 178.8(9) | C32 | O9 | C31 | C30 | 179.3(10) |

| | | | | | | | | | |
|---|---|---|---|---|---|---|---|---|---|
| C13 | N1 | C7 | C8 | 167.7(9) | C33 | N3 | C27 | C28 | -154.6(9) |
| C13 | N1 | C10 | C9 | 171.2(10) | C33 | N3 | C30 | C29 | -179.5(9) |
| C13 | N1 | C10 | C11 | -67.0(14) | C33 | N3 | C30 | C31 | -57.6(12) |
| C13 | C14 | C15 | O5 | 67.3(12) | C33 | C34 | C35 | O12 | 178.0(8) |
| C14 | N2 | C16 | O6 | -171.0(10) | C34 | N4 | C36 | O13 | 170.0(9) |
| C14 | N2 | C16 | O7 | 8.5(18) | C34 | N4 | C36 | O14 | -9.9(16) |
| C16 | O6 | C17 | C18 | 68.7(13) | C36 | O13 | C37 | C38 | 57.6(14) |
| C16 | O6 | C17 | C19 | -54.5(15) | C36 | O13 | C37 | C39 | -68.4(12) |
| C16 | O6 | C17 | C20 | -173.9(10) | C36 | O13 | C37 | C40 | 176.0(10) |
| C16 | N2 | C14 | C13 | -129.1(11) | C36 | N4 | C34 | C33 | -71.9(12) |
| C16 | N2 | C14 | C15 | 110.7(12) | C36 | N4 | C34 | C35 | 168.4(9) |
| C17 | O6 | C16 | O7 | 1.1(18) | C37 | O13 | C36 | O14 | -1.9(16) |
| C17 | O6 | C16 | N2 | -179.4(10) | C37 | O13 | C36 | N4 | 178.2(9) |

**Table S18.** Hydrogen Atom Coordinates (Å×10$^4$) and Isotropic Displacement Parameters (Å$^2$×10$^3$) for **3**.

| **Atom** | **x** | **y** | **z** | **U(eq)** |
|---|---|---|---|---|
| H5 | 5460(40) | 10600(300) | 7430(50) | 63 |
| H2 | 6450(90) | 4030(150) | 7860(40) | 53 |
| H1 | 3203.72 | 5034.01 | 9927.9 | 48 |
| H2A | 2227.5 | 2108.49 | 10103.3 | 49 |
| H4 | 1134.03 | 2996.94 | 8731.63 | 53 |
| H5A | 2110.25 | 5922.46 | 8556.28 | 50 |
| H7A | 3585.12 | 5767.14 | 8316.76 | 44 |
| H7B | 3436.23 | 7922.13 | 7950.16 | 44 |
| H8 | 2731.36 | 9397.54 | 8643.58 | 50 |
| H9A | 3848.15 | 11978 | 8554.54 | 54 |
| H9B | 3961.54 | 11279.8 | 9137.07 | 54 |
| H10 | 5217.55 | 10809.5 | 8510.72 | 44 |
| H12A | 6363.33 | 5778.88 | 9552.4 | 65 |
| H12B | 5692.59 | 3712.64 | 9543.93 | 65 |
| H12C | 5494.18 | 5944.85 | 9870.58 | 65 |
| H14 | 6223.81 | 8240.87 | 8333.6 | 41 |
| H15A | 6975.89 | 8814.02 | 7542.29 | 50 |
| H15B | 6166.31 | 7759.43 | 7253.13 | 50 |
| H18A | 7703.21 | 3008.07 | 9323.72 | 69 |
| H18B | 8437.08 | 1121.34 | 9388.19 | 69 |
| H18C | 7558.21 | 485.71 | 9109.52 | 69 |
| H19A | 9181.48 | 4872.37 | 8409.32 | 98 |
| H19B | 9517.93 | 3658.69 | 8911.97 | 98 |
| H19C | 8758.71 | 5491.29 | 8941.05 | 98 |
| H20A | 8341.28 | -838.41 | 8381.14 | 85 |
| H20B | 9263.29 | -130.91 | 8595.88 | 85 |
| H20C | 8942.85 | 847.93 | 8065.42 | 85 |
| H12 | 4090(90) | -1210(130) | 7070(50) | 56 |
| H4A | 4770(60) | 4700(200) | 6980(20) | 39 |
| H21 | 39.05 | 4377.08 | 5454.6 | 40 |
| H22 | -1193.3 | 4514.1 | 4951.48 | 44 |
| H24 | -1836.6 | -1853 | 5410.34 | 48 |
| H25 | -604.33 | -1990.1 | 5913.47 | 39 |
| H27A | 1824.21 | 854.16 | 6614.51 | 36 |
| H27B | 2329.02 | 1688.73 | 6116.52 | 36 |
| H28 | 1164.76 | 3715.92 | 5820.02 | 36 |
| H29A | 212.96 | 5374.7 | 6383.44 | 42 |

| | | | | |
|---|---|---|---|---|
| H29B | 406.17 | 3482.9 | 6810.07 | 42 |
| H30 | 1489.94 | 7150.07 | 6496.47 | 36 |
| H32A | 739.43 | 10989.6 | 7802.54 | 83 |
| H32B | 1554.78 | 9519.15 | 7972.53 | 83 |
| H32C | 614.19 | 8432.68 | 8000.28 | 83 |
| H34 | 3596.4 | 2292.15 | 6417.46 | 33 |
| H35A | 3719.02 | 2649.41 | 7499.82 | 37 |
| H35B | 3074.17 | 964.52 | 7213.53 | 37 |
| H38A | 4699.01 | 8815.5 | 5529.35 | 89 |
| H38B | 5442.16 | 10679 | 5469.61 | 89 |
| H38C | 4896.33 | 10556.3 | 5979.06 | 89 |
| H39A | 6549.72 | 5327.72 | 5707.33 | 69 |
| H39B | 6405.2 | 7138.05 | 5264.19 | 69 |
| H39C | 5656.57 | 5391.37 | 5409.92 | 69 |
| H40A | 6304.81 | 10360 | 6415.04 | 79 |
| H40B | 6829.79 | 10176.1 | 5898.97 | 79 |
| H40C | 6920.21 | 8221.55 | 6316.41 | 79 |

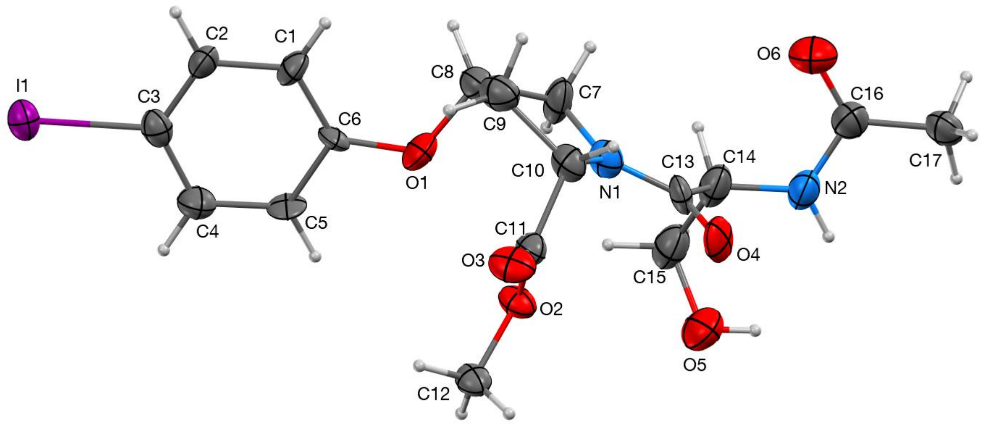

**Figure S17.** Molecular diagram of Ac-L-serine-(2*S*,4*S*)-4-iodophenyl-hydroxyproline methyl ester (**5**) showing two symmetry unique compound molecules with 50% probability ellipsoids. H-atoms depicted at arbitrary radius. Minor disordered contribution omitted for clarity. Diffractable crystals were obtained by slow evaporation at room temperature from a solution of **5** in acetone. This structure has CCDC number 2330290.

**Table S19.** Crystallographic data and refinement details for **5**.

| | |
|---|---|
| Empirical formula | $C_{17}H_{21}IN_2O_6$ |
| Formula weight | 476.26 |
| Temperature/K | 100.00 |
| Crystal system | monoclinic |
| Space group | $P2_1$ |
| a/Å | 8.4990(6) |
| b/Å | 6.0043(4) |
| c/Å | 18.3908(12) |
| α/° | 90 |
| β/° | 92.6460(10) |
| γ/° | 90 |
| Volume/Å$^3$ | 937.49(11) |
| Z | 2 |
| $\rho_{calc}$g/cm$^3$ | 1.687 |
| μ/mm$^{-1}$ | 13.741 |
| F(000) | 476.0 |
| Crystal size/mm$^3$ | 0.5 × 0.169 × 0.086 |
| Radiation | CuKα (λ = 1.54178) |
| 2Θ range for data collection/° | 4.81 to 133.228 |
| Index ranges | -10 ≤ h ≤ 10, -7 ≤ k ≤ 7, -21 ≤ l ≤ 21 |
| Reflections collected | 10031 |

| | |
|---|---|
| Independent reflections | 3213 [$R_{int}$ = 0.0755, $R_{sigma}$ = 0.0946] |
| Data/restraints/parameters | 3213/154/277 |
| Goodness-of-fit on $F^2$ | 1.142 |
| Final R indexes [I>=2σ (I)] | $R_1$ = 0.0859, $wR_2$ = 0.2347 |
| Final R indexes [all data] | $R_1$ = 0.0862, $wR_2$ = 0.2359 |
| Largest diff. peak/hole / e Å$^{-3}$ | 2.55/-1.36 |
| Flack parameter | -0.003(17) |

**Table S20.** Fractional Atomic Coordinates ($\times 10^4$) and Equivalent Isotropic Displacement Parameters ($Å^2 \times 10^3$) for **5**. $U_{eq}$ is defined as 1/3 of the trace of the orthogonalised $U_{IJ}$ tensor.

| Atom | x | y | z | U(eq) |
|---|---|---|---|---|
| I1 | 8227.4(7) | 4369(4) | -504.5(3) | 35.2(5) |
| O1 | 4654(13) | 5700(20) | 2346(7) | 48(3) |
| O2 | 1069(10) | 5073(13) | 2467(5) | 31.3(19) |
| O3 | 191(12) | 8304(15) | 2017(6) | 40(2) |
| O4 | 430(11) | 6864(19) | 4125(6) | 43(2) |
| N1 | 2747(13) | 7060(20) | 3592(6) | 33(2) |
| C1 | 6343(13) | 7330(20) | 1440(7) | 29(2) |
| C2 | 7153(13) | 6980(20) | 801(7) | 33(3) |
| C3 | 6981(17) | 5000(30) | 434(7) | 31(3) |
| C4 | 5976(17) | 3350(30) | 672(8) | 41(3) |
| C5 | 5210(19) | 3700(20) | 1330(9) | 35(3) |
| C6 | 5423(14) | 5680(20) | 1712(8) | 28(3) |
| C7 | 4462(17) | 6800(30) | 3557(8) | 42(4) |
| C8 | 4830(17) | 7640(30) | 2818(8) | 39(3) |
| C9 | 3579(17) | 9330(40) | 2637(8) | 41(3) |
| C10 | 2090(14) | 8480(20) | 2995(7) | 31(2) |
| C11 | 994(14) | 7280(20) | 2445(7) | 30(2) |
| C12 | 138(16) | 3980(20) | 1890(8) | 32(3) |
| C13 | 1872(16) | 6430(20) | 4137(7) | 31(3) |
| O5 | 1320(20) | 1620(40) | 4499(12) | 60(5) |
| O6 | 3310(30) | 8290(40) | 5627(14) | 66(5) |
| N2 | 2070(40) | 5040(40) | 5440(14) | 40(3) |
| C14 | 2675(15) | 4980(30) | 4739(8) | 42(2) |
| C15 | 2840(30) | 2570(40) | 4500(16) | 46(3) |
| C16 | 2430(30) | 6820(50) | 5859(15) | 43(3) |
| C17 | 1720(40) | 6860(60) | 6587(18) | 38(5) |
| O5A | 3420(30) | 1040(40) | 4793(18) | 57(5) |
| O6A | 4180(30) | 6530(50) | 6008(16) | 52(5) |
| N2A | 1960(50) | 5770(80) | 5390(20) | 40(3) |
| C14A | 2675(15) | 4980(30) | 4739(8) | 42(2) |
| C15A | 2270(40) | 2570(40) | 4520(20) | 46(3) |
| C16A | 2700(30) | 6630(50) | 5983(18) | 43(3) |
| C17A | 1930(50) | 7640(90) | 6620(20) | 45(7) |

**Table S21.** Anisotropic Displacement Parameters ($Å^2 \times 10^3$) for **5**. The Anisotropic displacement factor exponent takes the form: $-2\pi^2[h^2a^{*2}U_{11}+2hka^*b^*U_{12}+\ldots]$.

| **Atom** | **$U_{11}$** | **$U_{22}$** | **$U_{33}$** | **$U_{23}$** | **$U_{13}$** | **$U_{12}$** |
|---|---|---|---|---|---|---|
| I1 | 39.5(6) | 40.1(7) | 25.9(6) | -2.0(4) | 2.6(4) | 4.0(4) |
| O1 | 47(6) | 58(7) | 40(6) | 2(5) | 18(5) | -23(5) |
| O2 | 42(4) | 22(4) | 29(4) | 0(3) | -5(4) | -2(3) |
| O3 | 52(5) | 24(4) | 43(5) | 10(4) | -10(4) | -3(4) |
| O4 | 31(4) | 57(6) | 41(6) | -15(5) | 14(4) | 13(4) |
| N1 | 34(5) | 41(6) | 22(5) | 4(5) | -5(4) | 3(5) |
| C1 | 27(5) | 37(7) | 25(6) | 11(5) | 7(4) | -4(5) |
| C2 | 25(5) | 44(7) | 29(7) | 0(5) | 8(5) | -7(5) |
| C3 | 33(6) | 37(9) | 23(6) | -2(5) | -2(5) | 4(5) |
| C4 | 49(8) | 34(7) | 39(8) | -6(6) | 5(6) | -13(6) |
| C5 | 48(7) | 13(6) | 44(8) | 1(5) | 10(6) | -17(5) |
| C6 | 38(6) | 21(7) | 26(6) | -12(5) | 3(5) | -10(5) |
| C7 | 32(6) | 74(11) | 21(7) | 7(7) | 6(5) | 2(7) |
| C8 | 46(7) | 42(7) | 29(7) | -5(6) | 3(6) | -14(6) |
| C9 | 48(7) | 42(7) | 31(7) | 2(8) | -7(5) | -9(8) |
| C10 | 39(6) | 31(5) | 23(6) | 5(5) | -5(5) | 2(5) |
| C11 | 37(6) | 28(6) | 23(6) | 3(5) | 3(5) | -5(5) |
| C12 | 45(7) | 22(8) | 30(7) | 1(5) | 2(5) | -6(5) |
| C13 | 38(6) | 40(7) | 14(6) | -8(5) | 3(5) | 5(5) |
| O5 | 65(9) | 68(10) | 47(9) | 5(8) | 14(8) | -25(8) |
| O6 | 69(9) | 60(10) | 72(10) | -18(8) | 15(9) | -29(8) |
| N2 | 32(4) | 58(6) | 31(5) | 1(5) | 8(3) | -4(5) |
| C14 | 33(4) | 62(6) | 31(5) | 6(4) | 9(4) | -1(4) |
| C15 | 39(6) | 63(7) | 36(6) | 10(5) | 8(5) | -1(5) |
| C16 | 35(5) | 56(6) | 37(5) | -5(5) | 4(4) | -7(5) |
| C17 | 36(10) | 45(12) | 32(9) | -7(10) | -5(8) | -3(10) |
| O5A | 57(10) | 65(11) | 48(12) | 8(10) | -7(9) | 0(9) |
| O6A | 39(7) | 68(12) | 50(11) | -13(10) | 0(8) | 1(8) |
| N2A | 32(4) | 58(6) | 31(5) | 1(5) | 8(3) | -4(5) |
| C14A | 33(4) | 62(6) | 31(5) | 6(4) | 9(4) | -1(4) |
| C15A | 39(6) | 63(7) | 36(6) | 10(5) | 8(5) | -1(5) |
| C16A | 35(5) | 56(6) | 37(5) | -5(5) | 4(4) | -7(5) |
| C17A | 37(12) | 61(14) | 35(11) | -6(11) | 0(10) | -6(12) |

**Table S22.** Bond lengths for **5**.

| Atom | Atom | Length, Å | Atom | Atom | Length, Å |
|---|---|---|---|---|---|
| I1 | C3 | 2.101(15) | C8 | C9 | 1.50(3) |
| O1 | C6 | 1.362(17) | C9 | C10 | 1.541(19) |
| O1 | C8 | 1.456(19) | C10 | C11 | 1.522(17) |
| O2 | C11 | 1.329(16) | C13 | C14 | 1.55(2) |
| O2 | C12 | 1.452(17) | C13 | C14 | A1.55(2) |
| O3 | C11 | 1.188(17) | O5 | C15 | 1.413(15) |
| O4 | C13 | 1.252(17) | O6 | C16 | 1.24(3) |
| N1 | C7 | 1.471(18) | N2 | C14 | 1.41(3) |
| N1 | C10 | 1.479(16) | N2 | C16 | 1.35(3) |
| N1 | C13 | 1.329(18) | C14 | C15 | 1.52(3) |
| C1 | C2 | 1.405(18) | C16 | C17 | 1.50(3) |
| C1 | C6 | 1.370(17) | O5A | C15A | 1.418(15) |
| C2 | C3 | 1.38(2) | O6A | C16A | 1.26(3) |
| C3 | C4 | 1.39(2) | N2A | C14A | 1.45(5) |
| C4 | C5 | 1.42(2) | N2A | C16A | 1.34(3) |
| C5 | C6 | 1.391(18) | C14A | C15A | 1.53(3) |
| C7 | C8 | 1.50(2) | C16A | C17A | 1.49(3) |

**Table S23.** Bond angles for **5**.

| Atom | Atom | Atom | Angle, ° | Atom | Atom | Atom | Angle, ° |
|---|---|---|---|---|---|---|---|
| C6 | O1 | C8 | 118.4(10) | O3 | C11 | O2 | 124.1(12) |
| C11 | O2 | C12 | 114.0(9) | O3 | C11 | C10 | 120.8(12) |
| C7 | N1 | C10 | 111.7(11) | O4 | C13 | N1 | 120.7(13) |
| C13 | N1 | C7 | 126.4(12) | O4 | C13 | C14 | 121.9(12) |
| C13 | N1 | C10 | 121.1(11) | O4 | C13 | C14A | 121.9(12) |
| C6 | C1 | C2 | 120.6(13) | N1 | C13 | C14 | 117.2(11) |
| C3 | C2 | C1 | 119.6(13) | N1 | C13 | C14A | 117.2(11) |
| C2 | C3 | I1 | 120.8(11) | C16 | N2 | C14 | 117(3) |
| C2 | C3 | C4 | 121.1(14) | N2 | C14 | C13 | 118.0(15) |
| C4 | C3 | I1 | 118.1(11) | N2 | C14 | C15 | 109.7(18) |
| C3 | C4 | C5 | 118.4(14) | C15 | C14 | C13 | 112.1(16) |
| C6 | C5 | C4 | 120.3(12) | O5 | C15 | C14 | 106.6(18) |
| O1 | C6 | C1 | 127.7(12) | O6 | C16 | N2 | 120(3) |
| O1 | C6 | C5 | 112.5(12) | O6 | C16 | C17 | 125(3) |
| C1 | C6 | C5 | 119.8(12) | N2 | C16 | C17 | 115(3) |
| N1 | C7 | C8 | 104.7(12) | C16A | N2A | C14A | 127(4) |
| O1 | C8 | C7 | 104.6(12) | N2A | C14A | C13 | 102.6(19) |
| O1 | C8 | C9 | 111.0(12) | N2A | C14A | C15A | 115(2) |
| C7 | C8 | C9 | 104.7(13) | C15A | C14A | C13 | 105.0(15) |
| C8 | C9 | C10 | 105.5(15) | O5A | C15A | C14A | 112(2) |
| N1 | C10 | C9 | 102.7(11) | O6A | C16A | N2A | 116(4) |
| N1 | C10 | C11 | 114.9(11) | O6A | C16A | C17A | 118(3) |
| C11 | C10 | C9 | 111.2(10) | N2A | C16A | C17A | 126(4) |
| O2 | C11 | C10 | 115.0(11) | | | | |

**Table S24.** Torsion angles for **5**.

| **A** | **B** | **C** | **D** | **Angle, °** | **A** | **B** | **C** | **D** | **Angle, °** |
|---|---|---|---|---|---|---|---|---|---|
| I1 | C3 | C4 | C5 | 175.9(12) | C7 | N1 | C13 | C14A | -10(2) |
| O1 | C8 | C9 | C10 | 78.9(13) | C7 | C8 | C9 | C10 | -33.4(16) |
| O4 | C13 | C14 | N2 | -32(2) | C8 | O1 | C6 | C1 | -2(2) |
| O4 | C13 | C14 | C15 | 97.2(15) | C8 | O1 | C6 | C5 | 177.4(13) |
| O4 | C13 | C14A | N2A | -42(2) | C8 | C9 | C10 | N1 | 24.6(14) |
| O4 | C13 | C14A | C15A | 79(2) | C8 | C9 | C10 | C11 | -98.8(13) |
| N1 | C7 | C8 | O1 | -88.3(15) | C9 | C10 | C11 | O2 | 100.5(15) |
| N1 | C7 | C8 | C9 | 28.5(17) | C9 | C10 | C11 | O3 | -76.8(16) |
| N1 | C10 | C11 | O2 | -15.6(15) | C10 | N1 | C7 | C8 | -13.3(17) |
| N1 | C10 | C11 | O3 | 167.1(12) | C10 | N1 | C13 | O4 | 6(2) |
| N1 | C13 | C14 | N2 | 154.2(17) | C10 | N1 | C13 | C14 | -179.3(12) |
| N1 | C13 | C14 | C15 | -77.0(16) | C10 | N1 | C13 | C14A | -179.3(12) |
| N1 | C13 | C14A | N2A | 144(2) | C12 | O2 | C11 | O3 | 3.2(17) |
| N1 | C13 | C14A | C15A | -95(2) | C12 | O2 | C11 | C10 | -174.0(10) |
| C1 | C2 | C3 | I1 | -178.0(9) | C13 | N1 | C7 | C8 | 176.7(14) |
| C1 | C2 | C3 | C4 | 2(2) | C13 | N1 | C10 | C9 | 163.6(14) |
| C2 | C1 | C6 | O1 | 175.2(13) | C13 | N1 | C10 | C11 | -75.4(15) |
| C2 | C1 | C6 | C5 | -4(2) | C13 | C14 | C15 | O5 | -74(2) |
| C2 | C3 | C4 | C5 | -4(2) | C13 | C14A | C15A | O5A | 153(3) |
| C3 | C4 | C5 | C6 | 2(2) | N2 | C14 | C15 | O5 | 60(3) |
| C4 | C5 | C6 | O1 | -177.5(15) | C14 | N2 | C16 | O6 | -2(3) |
| C4 | C5 | C6 | C1 | 2(2) | C14 | N2 | C16 | C17 | 178(2) |
| C6 | O1 | C8 | C7 | -157.0(12) | C16 | N2 | C14 | C13 | -77(3) |
| C6 | O1 | C8 | C9 | 90.6(15) | C16 | N2 | C14 | C15 | 153(2) |
| C6 | C1 | C2 | C3 | 2.0(19) | N2A | C14A | C15A | O5A | -95(4) |
| C7 | N1 | C10 | C9 | -7.0(16) | C14A | N2A | C16A | O6A | -8(5) |
| C7 | N1 | C10 | C11 | 114.0(13) | C14A | N2A | C16A | C17A | 174(4) |
| C7 | N1 | C13 | O4 | 175.6(15) | C16A | N2A | C14A | C13 | -122(3) |
| C7 | N1 | C13 | C14 | -10(2) | C16A | N2A | C14A | C15A | 125(3) |

**Table S25.** Hydrogen Atom Coordinates (Å×$10^4$) and Isotropic Displacement Parameters (Å$^2$×$10^3$) for **5**.

| Atom | x | y | z | U(eq) |
|---|---|---|---|---|
| H1 | 6433.71 | 8718.5 | 1686.02 | 35 |
| H2 | 7814.88 | 8116.08 | 623.1 | 39 |
| H4 | 5807.05 | 2018.08 | 399.13 | 49 |
| H5 | 4548.28 | 2571.52 | 1511.87 | 42 |
| H7A | 4771.11 | 5215.28 | 3614.32 | 51 |
| H7B | 5021.51 | 7684.04 | 3942.17 | 51 |
| H8 | 5912.12 | 8291.17 | 2810.44 | 47 |
| H9A | 3392.1 | 9462.58 | 2103.4 | 49 |
| H9B | 3889.15 | 10810.6 | 2834.87 | 49 |
| H10 | 1519.91 | 9763.04 | 3206.19 | 37 |
| H12A | 449.56 | 4533.96 | 1416.89 | 48 |
| H12B | -981.48 | 4291.05 | 1947.16 | 48 |
| H12C | 315.96 | 2365.88 | 1916.4 | 48 |
| H5A | 999.26 | 1635.36 | 4923.59 | 89 |
| H2A | 1468.46 | 3965.1 | 5596.78 | 48 |
| H14 | 3775.26 | 5555.86 | 4802.23 | 50 |
| H15A | 3261.18 | 2499.83 | 4006.42 | 55 |
| H15B | 3574.16 | 1758.55 | 4840.82 | 55 |
| H17A | 2515.43 | 7309.74 | 6959.46 | 57 |
| H17B | 1328.74 | 5366.85 | 6700.17 | 57 |
| H17C | 840.35 | 7918.13 | 6577.62 | 57 |
| H5AA | 2998.21 | 115.41 | 5066.81 | 86 |
| H2AA | 929.24 | 5674.77 | 5396.31 | 48 |
| H14A | 3840.27 | 5214.32 | 4768.34 | 50 |
| H15C | 1233.55 | 2173.7 | 4710.79 | 55 |
| H15D | 2178.82 | 2464.51 | 3983.84 | 55 |
| H17D | 2242.17 | 6816.01 | 7061.97 | 67 |
| H17E | 781.06 | 7563.37 | 6538.13 | 67 |
| H17F | 2253.24 | 9198.12 | 6671.9 | 67 |

**Table S26.** Atomic occupancy of 5**.**

| **Atom** | **Occupancy** | **Atom** | **Occupancy** | **Atom** | **Occupancy** |
|---|---|---|---|---|---|
| O5 | 0.579(18) | H5A | 0.579(18) | O6 | 0.579(18) |
| N2 | 0.579(18) | H2A | 0.579(18) | C14 | 0.579(18) |
| H14 | 0.579(18) | C15 | 0.579(18) | H15A | 0.579(18) |
| H15B | 0.579(18) | C16 | 0.579(18) | C17 | 0.579(18) |
| H17A | 0.579(18) | H17B | 0.579(18) | H17C | 0.579(18) |
| O5A | 0.421(18) | H5AA | 0.421(18) | O6A | 0.421(18) |
| N2A | 0.421(18) | H2AA | 0.421(18) | C14A | 0.421(18) |
| H14A | 0.421(18) | C15A | 0.421(18) | H15C | 0.421(18) |
| H15D | 0.421(18) | C16A | 0.421(18) | C17A | 0.421(18) |
| H17D | 0.421(18) | H17E | 0.421(18) | H17F | 0.421(18) |

### Coordinates of structures obtained via geometry optimization calculations

### Boc-Ser-hyp(4-I-Ph)-OMe and Ac-Ser-hyp(4-I-Ph)-OMe structures subjected hydrogen position optimization with positions of heavy atoms fixed to those observed crystallographically

#### Structure of Boc-Ser-*cis*-hyp(4-I-Ph)-OMe subjected to hydrogen position optimization with heavy atom positions fixed to those observed crystallographically

optimized M06-2X/Def2TZVP/H2O

0 1

| | | | | |
|---|---|---|---|---|
| I | -1 | 6.60978700 | 1.39379300 | -0.45967400 |
| O | -1 | 1.38852300 | -1.63844500 | 1.14569600 |
| O | -1 | -1.38653300 | -0.11993000 | 1.24349700 |
| O | -1 | -1.95013300 | -1.42801100 | 2.95584400 |
| O | -1 | -1.62624300 | -1.06610400 | -2.44532500 |
| O | -1 | -4.09221900 | -3.59518900 | -1.19935200 |
| H | 0 | -3.34601600 | -3.96028800 | -1.68650800 |
| O | -1 | -4.92440500 | 1.99095400 | -0.93687000 |
| O | -1 | -4.54704800 | 0.61828200 | 0.82144600 |
| N | -1 | -1.18281200 | -2.08644500 | -0.50783700 |
| N | -1 | -3.86228700 | 0.07959100 | -1.28445600 |
| H | 0 | -3.80676500 | 0.37757000 | -2.24513800 |
| C | -1 | 2.95687000 | 0.03015500 | 1.48733100 |
| H | 0 | 2.39715500 | 0.29422300 | 2.37449300 |
| C | -1 | 4.11526000 | 0.71866500 | 1.14602000 |
| H | 0 | 4.45727200 | 1.52802600 | 1.77672700 |
| C | -1 | 4.82789200 | 0.36021600 | 0.00796700 |
| C | -1 | 4.38155200 | -0.68718900 | -0.79134100 |
| H | 0 | 4.92905900 | -0.97767000 | -1.67732200 |
| C | -1 | 3.22312000 | -1.37554100 | -0.44997000 |
| H | 0 | 2.90092400 | -2.18888500 | -1.08446600 |
| C | -1 | 2.51121200 | -1.01612600 | 0.69038600 |
| C | -1 | 0.20777200 | -2.31800000 | -0.89296800 |
| H | 0 | 0.64298300 | -1.40260500 | -1.29152100 |
| H | 0 | 0.26642100 | -3.09102800 | -1.66146300 |
| C | -1 | 0.89012900 | -2.76927900 | 0.41277900 |
| H | 0 | 1.68913800 | -3.49152900 | 0.25361000 |
| C | -1 | -0.26958100 | -3.33714500 | 1.22261100 |
| H | 0 | -0.46298700 | -4.35518100 | 0.88656400 |
| H | 0 | -0.05791000 | -3.35767300 | 2.28934100 |
| C | -1 | -1.46849600 | -2.46560100 | 0.87427700 |
| H | 0 | -2.39159800 | -3.04057000 | 0.94420500 |
| C | -1 | -1.62765600 | -1.27828200 | 1.82021800 |
| C | -1 | -1.55107700 | 1.04702800 | 2.10076200 |
| H | 0 | -2.57604200 | 1.08255600 | 2.46059200 |
| H | 0 | -1.33263900 | 1.89959700 | 1.46694800 |

| | | | | |
|---|---|---|---|---|
| H | 0 | -0.85136300 | 0.98586900 | 2.93111300 |
| C | -1 | -2.00615000 | -1.45404400 | -1.33870900 |
| C | -1 | -3.45517300 | -1.25589600 | -0.92118400 |
| H | 0 | -3.59939700 | -1.35043700 | 0.15062600 |
| C | -1 | -4.34712400 | -2.28333300 | -1.63355000 |
| H | 0 | -5.38199700 | -2.05113700 | -1.38111500 |
| H | 0 | -4.22420000 | -2.18006700 | -2.71560700 |
| C | -1 | -4.46104300 | 0.89373000 | -0.35239900 |
| C | -1 | -5.59537500 | 3.02788300 | -0.12954500 |
| C | -1 | -4.57189200 | 3.68396200 | 0.76624700 |
| H | 0 | -4.19030600 | 2.98145300 | 1.50482800 |
| H | 0 | -5.03657000 | 4.51917300 | 1.29189500 |
| H | 0 | -3.73899600 | 4.07167400 | 0.17760800 |
| C | -1 | -6.73917100 | 2.48026500 | 0.67277200 |
| H | 0 | -7.39193900 | 1.86818400 | 0.04679900 |
| H | 0 | -7.32808400 | 3.31933100 | 1.04601500 |
| H | 0 | -6.40355200 | 1.88642300 | 1.51712500 |
| C | -1 | -6.08986300 | 3.99651100 | -1.16888300 |
| H | 0 | -5.26334500 | 4.35832400 | -1.78106600 |
| H | 0 | -6.56039300 | 4.85023700 | -0.68184800 |
| H | 0 | -6.82462400 | 3.52106700 | -1.82010800 |

1 17 1.0
2 22 1.0 26 1.0
3 33 1.5 34 1.0
4 33 2.0
5 38 2.0
6 7 1.0 41 1.0
7
8 44 1.5 45 1.0
9 44 2.0
10 23 1.0 31 1.0 38 1.5
11 12 1.0 39 1.0 44 1.5
12
13 14 1.0 15 1.5 22 1.5
14
15 16 1.0 17 1.5
16
17 18 1.5
18 19 1.0 20 1.5
19
20 21 1.0 22 1.5
21
22
23 24 1.0 25 1.0 26 1.0
24
25

26 27 1.0 28 1.0
27
28 29 1.0 30 1.0 31 1.0
29
30
31 32 1.0 33 1.0
32
33
34 35 1.0 36 1.0 37 1.0
35
36
37
38 39 1.0
39 40 1.0 41 1.0
40
41 42 1.0 43 1.0
42
43
44
45 46 1.0 50 1.0 54 1.0
46 47 1.0 48 1.0 49 1.0
47
48
49
50 51 1.0 52 1.0 53 1.0
51
52
53
54 55 1.0 56 1.0 57 1.0
55
56
57

**Structure of Boc-Ser-*trans*-hyp(4-I-Ph)-OMe subjected to hydrogen position optimization with heavy atom positions fixed to those observed crystallographically**
optimized M06-2X/Def2TZVP/H2O

0 1

| | | | | |
|---|---|---|---|---|
| I | -1 | -7.48753700 | -1.09022300 | -0.42519900 |
| O | -1 | -1.69825000 | 0.31739900 | 1.47276800 |
| O | -1 | 1.30897800 | 3.48977000 | -2.22116600 |
| O | -1 | 1.70449900 | 3.86502900 | -0.04334400 |
| O | -1 | 3.50555600 | 1.36141700 | -0.77388500 |
| O | -1 | 4.49774000 | 0.98656000 | 3.60898700 |
| H | 0 | 3.92019500 | 0.36763300 | 4.06863000 |
| O | -1 | 6.29724000 | -1.44775700 | -0.19312900 |
| O | -1 | 4.10427500 | -1.97736900 | 0.05820700 |

| | | | | |
|---|---|---|---|---|
| N | -1 | 1.58867700 | 1.04609900 | 0.31524700 |
| N | -1 | 4.98068500 | -0.13141000 | 1.00248400 |
| H | 0 | 5.75306900 | 0.51462100 | 0.95888800 |
| C | -1 | -3.17653300 | -0.48787000 | -0.32002400 |
| H | 0 | -2.37166400 | -0.65615500 | -1.01989800 |
| C | -1 | -4.47142000 | -0.79778500 | -0.72013200 |
| H | 0 | -4.63751200 | -1.20183000 | -1.70906700 |
| C | -1 | -5.53562100 | -0.58877400 | 0.14946200 |
| C | -1 | -5.30570800 | -0.07223700 | 1.41775100 |
| H | 0 | -6.12669800 | 0.09211800 | 2.10221900 |
| C | -1 | -4.01084600 | 0.23767100 | 1.81795500 |
| H | 0 | -3.81780500 | 0.63640500 | 2.80464100 |
| C | -1 | -2.94676600 | 0.02862300 | 0.94884500 |
| C | -1 | 0.70068600 | 0.46044900 | 1.32691600 |
| H | 0 | 0.58935600 | 1.11127100 | 2.19846900 |
| H | 0 | 1.04398600 | -0.52055000 | 1.64497500 |
| C | -1 | -0.61694500 | 0.39007500 | 0.56220900 |
| H | 0 | -0.59376100 | -0.49597500 | -0.07583800 |
| C | -1 | -0.59393100 | 1.66134400 | -0.27221600 |
| H | 0 | -1.29744600 | 1.66249600 | -1.10134200 |
| H | 0 | -0.81925600 | 2.50265500 | 0.38783600 |
| C | -1 | 0.84911900 | 1.72303000 | -0.74184000 |
| H | 0 | 0.98393600 | 1.19538700 | -1.68735200 |
| C | -1 | 1.36007700 | 3.14498100 | -0.92741500 |
| C | -1 | 1.76125000 | 4.83452100 | -2.51597700 |
| H | 0 | 1.66266200 | 4.94615100 | -3.59008600 |
| H | 0 | 2.79876400 | 4.94601800 | -2.20850400 |
| H | 0 | 1.13941900 | 5.55751000 | -1.99272600 |
| C | -1 | 2.92997100 | 0.93406400 | 0.21165300 |
| C | -1 | 3.68622600 | 0.34994600 | 1.39473700 |
| H | 0 | 3.12818100 | -0.48079000 | 1.82594500 |
| C | -1 | 3.84473000 | 1.45268700 | 2.45628300 |
| H | 0 | 4.46017600 | 2.25494500 | 2.04412800 |
| H | 0 | 2.86135900 | 1.87030500 | 2.68833400 |
| C | -1 | 5.05644600 | -1.26829800 | 0.25568100 |
| C | -1 | 6.61803000 | -2.64149900 | -0.98730500 |
| C | -1 | 5.76080100 | -2.73068900 | -2.20606000 |
| H | 0 | 4.73515200 | -3.00161700 | -1.97345300 |
| H | 0 | 6.17626800 | -3.48702300 | -2.87371900 |
| H | 0 | 5.76239400 | -1.77815400 | -2.73932700 |
| C | -1 | 6.51843400 | -3.84326900 | -0.07474900 |
| H | 0 | 7.15321600 | -3.70062600 | 0.80023800 |
| H | 0 | 6.86111700 | -4.73304600 | -0.60509600 |
| H | 0 | 5.49320700 | -4.01036500 | 0.24958600 |
| C | -1 | 8.06536700 | -2.42675700 | -1.38902800 |
| H | 0 | 8.16122000 | -1.54611200 | -2.02545800 |
| H | 0 | 8.42153900 | -3.29582600 | -1.94118900 |

H 0 8.69234600 -2.29212500 -0.50737500

1 17 1.0
2 22 1.0 26 1.0
3 33 1.5 34 1.0
4 33 2.0
5 38 2.0
6 7 1.0 41 1.0
7
8 44 1.5 45 1.0
9 44 2.0
10 23 1.0 31 1.0 38 1.5
11 12 1.0 39 1.0 44 1.5
12
13 14 1.0 15 1.5 22 1.5
14
15 16 1.0 17 1.5
16
17 18 1.5
18 19 1.0 20 1.5
19
20 21 1.0 22 1.5
21
22
23 24 1.0 25 1.0 26 1.0
24
25
26 27 1.0 28 1.0
27
28 29 1.0 30 1.0 31 1.0
29
30
31 32 1.0 33 1.0
32
33
34 35 1.0 36 1.0 37 1.0
35
36
37
38 39 1.0
39 40 1.0 41 1.0
40
41 42 1.0 43 1.0
42
43
44
45 46 1.0 50 1.0 54 1.0

46 47 1.0 48 1.0 49 1.0
47
48
49
50 51 1.0 52 1.0 53 1.0
51
52
53
54 55 1.0 56 1.0 57 1.0
55
56
57

**Structure of Ac-Ser-*trans*-hyp(4-I-Ph)-OMe with Ser in the β conformation subjected to hydrogen position optimization with heavy atom positions fixed to those observed crystallographically**
optimized M06-2X/Def2TZVP/H2O

0 1

| | | | | |
|---|---|---|---|---|
| I | -1 | -6.45245000 | -0.47765500 | 0.26442200 |
| O | -1 | -0.23470500 | -0.25720600 | -0.05660400 |
| O | -1 | 1.45210000 | 2.28161100 | 0.44144800 |
| O | -1 | 1.24071400 | 3.30034900 | -1.52454000 |
| O | -1 | 4.50648400 | 1.34743100 | -0.39744000 |
| N | -1 | 2.66594100 | 0.09157900 | -0.65144000 |
| C | -1 | -2.34971100 | -0.73894200 | -1.20012600 |
| H | 0 | -1.89549800 | -1.08053800 | -2.12011800 |
| C | -1 | -3.75023700 | -0.79453500 | -1.10004800 |
| H | 0 | -4.32194300 | -1.18605400 | -1.92946500 |
| C | -1 | -4.36680600 | -0.35389500 | 0.04724100 |
| C | -1 | -3.62995100 | 0.18878800 | 1.09565500 |
| H | 0 | -4.10782600 | 0.56097600 | 1.99025900 |
| C | -1 | -2.21703000 | 0.19552600 | 0.99390800 |
| H | 0 | -1.61188500 | 0.57573600 | 1.80573800 |
| C | -1 | -1.59311200 | -0.28811700 | -0.15095000 |
| C | -1 | 1.86943900 | -1.14092400 | -0.55218300 |
| H | 0 | 1.73928200 | -1.46627400 | 0.47659100 |
| H | 0 | 2.34959100 | -1.94291000 | -1.11854400 |
| C | -1 | 0.54854300 | -0.78933600 | -1.16208300 |
| H | 0 | 0.05146300 | -1.66192800 | -1.57991100 |
| C | -1 | 0.86317800 | 0.28097600 | -2.16215300 |
| H | 0 | 0.00550900 | 0.91302500 | -2.38831900 |
| H | 0 | 1.19170900 | -0.18280300 | -3.09232200 |
| C | -1 | 2.03709500 | 1.07665700 | -1.55810000 |
| H | 0 | 2.74864300 | 1.39236200 | -2.31817200 |
| C | -1 | 1.55019000 | 2.35037300 | -0.88169000 |
| C | -1 | 0.85630800 | 3.45687800 | 1.05123900 |
| H | 0 | -0.14603200 | 3.60407000 | 0.65544100 |

| | | | | |
|---|---|---|---|---|
| H | 0 | 1.46933200 | 4.33115100 | 0.84576900 |
| H | 0 | 0.82651700 | 3.24640900 | 2.11470300 |
| C | -1 | 3.89485900 | 0.27674900 | -0.18181100 |
| O | -1 | 4.49401400 | 0.38609300 | 2.76415900 |
| H | 0 | 4.24664500 | 0.41197600 | 3.69370700 |
| O | -1 | 5.79920400 | -2.04156500 | -1.14418400 |
| N | -1 | 5.87039100 | -0.96243900 | 0.81201300 |
| H | 0 | 6.38784400 | -0.46724000 | 1.52146400 |
| C | -1 | 4.46826100 | -0.81935600 | 0.74530900 |
| H | 0 | 4.09542300 | -1.76170800 | 0.33828700 |
| C | -1 | 3.89501600 | -0.73589800 | 2.14924600 |
| H | 0 | 2.80802200 | -0.62808000 | 2.11668400 |
| H | 0 | 4.14624300 | -1.65789000 | 2.67905500 |
| C | -1 | 6.47980400 | -1.58997100 | -0.21056400 |
| C | -1 | 7.96959600 | -1.67891500 | -0.11488100 |
| H | 0 | 8.29126000 | -2.67705200 | -0.40591600 |
| H | 0 | 8.33035000 | -1.45218300 | 0.88520000 |
| H | 0 | 8.40572000 | -0.96935500 | -0.81927700 |

1 11 1.0
2 16 1.0 20 1.0
3 27 1.5 28 1.0
4 27 2.0
5 32 2.0
6 17 1.0 25 1.0 32 1.5
7 8 1.0 9 1.5 16 2.0
8
9 10 1.0 11 2.0
10
11 12 1.5
12 13 1.0 14 1.5
13
14 15 1.0 16 1.5
15
16
17 18 1.0 19 1.0 20 1.0
18
19
20 21 1.0 22 1.0
21
22 23 1.0 24 1.0 25 1.0
23
24
25 26 1.0 27 1.0
26
27
28 29 1.0 30 1.0 31 1.0

29
30
31
32 38 1.0
33 34 1.0 40 1.0
34
35 43 2.0
36 37 1.0 38 1.0 43 1.5
37
38 39 1.0 40 1.0
39
40 41 1.0 42 1.0
41
42
43 44 1.0
44 45 1.0 46 1.0 47 1.0
45
46
47

**Structure of Ac-Ser-*trans*-hyp(4-I-Ph)-OMe with Ser in the PPII conformation subjected to hydrogen position optimization with heavy atom positions fixed to those observed crystallographically**
optimized M06-2X/Def2TZVP/H2O

0 1

| | | | | |
|---|---|---|---|---|
| I | -1 | -6.44655200 | -0.60120900 | 0.13307700 |
| O | -1 | -0.23695900 | -0.15710000 | -0.10292300 |
| O | -1 | 1.38191400 | 2.26671500 | 0.91851900 |
| O | -1 | 1.14295700 | 3.66448700 | -0.79517800 |
| O | -1 | 4.45982900 | 1.60634300 | -0.07961200 |
| N | -1 | 2.65321200 | 0.38275000 | -0.59753900 |
| C | -1 | -2.33874100 | -0.44664900 | -1.33240300 |
| H | 0 | -1.87580400 | -0.57904600 | -2.30061000 |
| C | -1 | -3.73726600 | -0.55823700 | -1.25330800 |
| H | 0 | -4.29828300 | -0.78627700 | -2.14860300 |
| C | -1 | -4.36499700 | -0.38085800 | -0.04290200 |
| C | -1 | -3.64252400 | -0.04793200 | 1.09895000 |
| H | 0 | -4.13020700 | 0.11949900 | 2.04832500 |
| C | -1 | -2.23031100 | 0.01654500 | 1.00817500 |
| H | 0 | -1.63518000 | 0.23613500 | 1.88418100 |
| C | -1 | -1.59409000 | -0.20320500 | -0.20871200 |
| C | -1 | 1.88979300 | -0.86394300 | -0.75966400 |
| H | 0 | 1.77345900 | -1.38819700 | 0.18726300 |
| H | 0 | 2.39108200 | -1.52164600 | -1.47293900 |
| C | -1 | 0.55984400 | -0.42826400 | -1.29048500 |
| H | 0 | 0.08697000 | -1.20817800 | -1.88294500 |

| | | | | |
|---|---|---|---|---|
| C | -1 | 0.84562100 | 0.83370800 | -2.04576700 |
| H | 0 | -0.02885200 | 1.47588200 | -2.14133500 |
| H | 0 | 1.18637200 | 0.58149200 | -3.04995300 |
| C | -1 | 1.99815400 | 1.51743000 | -1.28401600 |
| H | 0 | 2.70072600 | 2.00211100 | -1.95883600 |
| C | -1 | 1.47775900 | 2.61041500 | -0.36122300 |
| C | -1 | 0.75527000 | 3.27436400 | 1.75519800 |
| H | 0 | -0.25114300 | 3.47272600 | 1.39364700 |
| H | 0 | 1.34407200 | 4.18829600 | 1.73723600 |
| H | 0 | 0.73235500 | 2.84802700 | 2.75209800 |
| C | -1 | 3.87694700 | 0.49877000 | -0.09319600 |
| O | -1 | 4.08509800 | -1.81727300 | 2.75746300 |
| H | 0 | 4.96372600 | -2.21143100 | 2.79795900 |
| O | -1 | 6.03854900 | -2.67737500 | -0.64774900 |
| N | -1 | 5.87843900 | -0.66279800 | 0.22962700 |
| H | 0 | 6.26748700 | 0.26472400 | 0.34554200 |
| C | -1 | 4.47949000 | -0.75039300 | 0.58957000 |
| H | 0 | 4.07399900 | -1.68247700 | 0.21110200 |
| C | -1 | 4.17313100 | -0.57426200 | 2.08090100 |
| H | 0 | 4.91564100 | 0.08944400 | 2.53123700 |
| H | 0 | 3.19085200 | -0.11090400 | 2.18606300 |
| C | -1 | 6.59888200 | -1.57331000 | -0.43447200 |
| C | -1 | 8.00584600 | -1.40355400 | -0.90881200 |
| H | 0 | 8.60577700 | -2.17753000 | -0.43467700 |
| H | 0 | 8.43218800 | -0.42591900 | -0.68993200 |
| H | 0 | 8.01251900 | -1.57248000 | -1.98365300 |

1 11 1.0
2 16 1.0 20 1.0
3 27 1.5 28 1.0
4 27 2.0
5 32 2.0
6 17 1.0 25 1.0 32 1.5
7 8 1.0 9 1.5 16 2.0
8
9 10 1.0 11 2.0
10
11 12 1.5
12 13 1.0 14 1.5
13
14 15 1.0 16 1.5
15
16
17 18 1.0 19 1.0 20 1.0
18
19
20 21 1.0 22 1.0

21
22 23 1.0 24 1.0 25 1.0
23
24
25 26 1.0 27 1.0
26
27
28 29 1.0 30 1.0 31 1.0
29
30
31
32 38 1.0
33 34 1.0 40 1.0
34
35 43 2.0
36 37 1.0 38 1.0 43 1.5
37
38 39 1.0 40 1.0
39
40 41 1.0 42 1.0
41
42
43 44 1.0
44 45 1.0 46 1.0 47 1.0
45
46
47

### Structures of Boc-Ser-hyp(4-I-Ph)-OMe and Ac-Ser-hyp(4-I-Ph)-OMe subjected to full geometry optimization in the conformations observed crystallographically

#### Structure of Boc-Ser-*cis*-hyp(4-I-Ph)-OMe subjected to full geometry optimization in the conformation observed crystallographically

optimized M06-2X/Def2TZVP/H2O

0 1

| | | | |
|---|---|---|---|
| I | 6.29409600 | 1.43675200 | -0.44298300 |
| O | 1.25635700 | -1.88054100 | 1.08239000 |
| O | -1.49065700 | -0.25403000 | 1.20714800 |
| O | -1.60763000 | -1.61402400 | 2.97966500 |
| O | -1.72056800 | -1.11860600 | -2.48211700 |
| O | -4.15820100 | -3.54136500 | -1.21431600 |
| H | -3.46074900 | -3.88480000 | -1.78265900 |
| O | -4.37078400 | 2.23760200 | -0.82808100 |
| O | -4.59107800 | 0.70426800 | 0.83504900 |
| N | -1.25228800 | -2.21252600 | -0.57413300 |
| N | -3.82356200 | 0.13767800 | -1.22933900 |
| H | -3.49513600 | 0.49913700 | -2.11247300 |
| C | 2.70109300 | -0.08733500 | 1.44264400 |
| H | 2.07934100 | 0.17459800 | 2.28857600 |
| C | 3.81848800 | 0.66313800 | 1.12845300 |
| H | 4.07291400 | 1.52417800 | 1.73109200 |
| C | 4.60337400 | 0.30189300 | 0.03890200 |
| C | 4.27218900 | -0.80053200 | -0.72849400 |
| H | 4.87860500 | -1.08408900 | -1.57775800 |
| C | 3.14775000 | -1.55726200 | -0.41425200 |
| H | 2.90838200 | -2.41126700 | -1.03134900 |
| C | 2.36110000 | -1.20138100 | 0.67631100 |
| C | 0.12697300 | -2.49398500 | -0.98257800 |
| H | 0.59349800 | -1.59346000 | -1.38012900 |
| H | 0.14745900 | -3.26507700 | -1.75471000 |
| C | 0.78792300 | -2.97930100 | 0.30632000 |
| H | 1.59341500 | -3.69170200 | 0.13199400 |
| C | -0.36729500 | -3.57798400 | 1.09279700 |
| H | -0.61204000 | -4.55906600 | 0.68886700 |
| H | -0.15190100 | -3.66544900 | 2.15413700 |
| C | -1.51564200 | -2.60791100 | 0.80143200 |
| H | -2.48101400 | -3.10623900 | 0.89346800 |
| C | -1.52787000 | -1.44245300 | 1.79099500 |
| C | -1.57649700 | 0.87839800 | 2.07991900 |
| H | -2.52105700 | 0.85609500 | 2.61913900 |
| H | -1.53008400 | 1.74971500 | 1.43481900 |
| H | -0.74332400 | 0.87157500 | 2.78003200 |
| C | -2.06622200 | -1.50261200 | -1.37090900 |
| C | -3.48882600 | -1.22248900 | -0.89375300 |

| | | | |
|---|---|---|---|
| H | -3.59507600 | -1.32505800 | 0.18187200 |
| C | -4.44760200 | -2.20516900 | -1.57535800 |
| H | -5.45866100 | -1.98441500 | -1.23363200 |
| H | -4.40722300 | -2.06075100 | -2.65835200 |
| C | -4.28784400 | 1.01093100 | -0.30054900 |
| C | -4.92897400 | 3.35322900 | -0.06924500 |
| C | -4.05518700 | 3.65688600 | 1.13878500 |
| H | -4.08231000 | 2.84386500 | 1.86037300 |
| H | -4.41779300 | 4.56655700 | 1.61904200 |
| H | -3.02460100 | 3.82557700 | 0.82257500 |
| C | -6.37088600 | 3.05897100 | 0.31909300 |
| H | -6.94281900 | 2.76353800 | -0.56184000 |
| H | -6.81791700 | 3.96548500 | 0.72876300 |
| H | -6.42953700 | 2.27101700 | 1.06560600 |
| C | -4.87252200 | 4.50031100 | -1.06610100 |
| H | -3.84435500 | 4.67809500 | -1.38226700 |
| H | -5.25809500 | 5.40825500 | -0.60287900 |
| H | -5.47672000 | 4.27126200 | -1.94428700 |

1 17 1.0
2 22 1.0 26 1.0
3 33 1.5 34 1.0
4 33 2.0
5 38 2.0
6 7 1.0 41 1.0
7
8 44 1.5 45 1.0
9 44 2.0
10 23 1.0 31 1.0 38 1.5
11 12 1.0 39 1.0 44 1.5
12
13 14 1.0 15 2.0 22 1.5
14
15 16 1.0 17 1.5
16
17 18 2.0
18 19 1.0 20 1.5
19
20 21 1.0 22 1.5
21
22
23 24 1.0 25 1.0 26 1.0
24
25
26 27 1.0 28 1.0
27
28 29 1.0 30 1.0 31 1.0

29
30
31 32 1.0 33 1.0
32
33
34 35 1.0 36 1.0 37 1.0
35
36
37
38 39 1.0
39 40 1.0 41 1.0
40
41 42 1.0 43 1.0
42
43
44
45 46 1.0 50 1.0 54 1.0
46 47 1.0 48 1.0 49 1.0
47
48
49
50 51 1.0 52 1.0 53 1.0
51
52
53
54 55 1.0 56 1.0 57 1.0
55
56
57

**Structure of Boc-Ser-*trans*-hyp(4-I-Ph)-OMe subjected to full geometry optimization in the conformation observed crystallographically**
optimized M06-2X/Def2TZVP/H2O

0 1

| | | | |
|---|---|---|---|
| I | -7.25772100 | -0.97731200 | -0.24351600 |
| O | -1.37975900 | 0.33284700 | 1.32318700 |
| O | 1.40343800 | 3.90973000 | -2.09151300 |
| O | 1.83850000 | 4.08662200 | 0.09663400 |
| O | 3.78086400 | 1.76533800 | -0.78258700 |
| O | 5.53189100 | 1.22863700 | 3.04340100 |
| H | 4.99847800 | 0.87207300 | 3.76232000 |
| O | 5.57849500 | -2.24410100 | -0.13424500 |
| O | 3.39333500 | -1.61182800 | -0.22854100 |
| N | 1.84660100 | 1.25674200 | 0.22654200 |
| N | 4.94745700 | -0.33683400 | 0.79936000 |
| H | 5.87230200 | -0.31682100 | 1.20088900 |

| | | | |
|---|---|---|---|
| C | -3.00060400 | -0.13685000 | -0.43283000 |
| H | -2.27889800 | -0.06198100 | -1.23265300 |
| C | -4.32210100 | -0.43261000 | -0.75250000 |
| H | -4.59432600 | -0.57882000 | -1.78866900 |
| C | -5.27225100 | -0.53420700 | 0.24758300 |
| C | -4.91697400 | -0.34202300 | 1.57823400 |
| H | -5.65615300 | -0.41892700 | 2.36375300 |
| C | -3.60445000 | -0.05025600 | 1.89817400 |
| H | -3.30686600 | 0.10025400 | 2.92744800 |
| C | -2.63983800 | 0.05261400 | 0.89644200 |
| C | 0.98457200 | 0.49151600 | 1.12244500 |
| H | 0.85591600 | 0.98997300 | 2.08639200 |
| H | 1.37224200 | -0.51311300 | 1.27268900 |
| C | -0.34639500 | 0.48190900 | 0.37409800 |
| H | -0.36069600 | -0.34928400 | -0.33602400 |
| C | -0.35564500 | 1.82605800 | -0.35419400 |
| H | -1.04661900 | 1.88403400 | -1.19061500 |
| H | -0.60597500 | 2.60216700 | 0.37266700 |
| C | 1.10251300 | 1.97201200 | -0.80010200 |
| H | 1.27314100 | 1.52459600 | -1.78060400 |
| C | 1.51387200 | 3.43246700 | -0.85845800 |
| C | 1.69506200 | 5.30278300 | -2.25152100 |
| H | 1.54763500 | 5.51724300 | -3.30448300 |
| H | 2.72444700 | 5.50203600 | -1.96010200 |
| H | 1.02000900 | 5.89851600 | -1.64016400 |
| C | 3.18829300 | 1.22466100 | 0.14150400 |
| C | 3.96351700 | 0.59073600 | 1.29580000 |
| H | 3.28371400 | 0.08865400 | 1.98730100 |
| C | 4.67862400 | 1.72315000 | 2.03400700 |
| H | 5.30921300 | 2.25919600 | 1.32509800 |
| H | 3.93285000 | 2.41517600 | 2.43371500 |
| C | 4.54776900 | -1.43268700 | 0.11280500 |
| C | 5.40524900 | -3.48441400 | -0.88638300 |
| C | 4.92535500 | -3.17900500 | -2.29774600 |
| H | 3.90964300 | -2.79166700 | -2.29604200 |
| H | 4.95092300 | -4.09685700 | -2.88628400 |
| H | 5.58743300 | -2.45130500 | -2.76927300 |
| C | 4.46391000 | -4.42205800 | -0.14437500 |
| H | 4.80309000 | -4.55673200 | 0.88379500 |
| H | 4.47232600 | -5.39475100 | -0.63762000 |
| H | 3.44618900 | -4.04006900 | -0.13932700 |
| C | 6.81269500 | -4.05910400 | -0.91662000 |
| H | 7.49467100 | -3.36801100 | -1.41264700 |
| H | 6.81208400 | -5.00237300 | -1.46232200 |
| H | 7.17091100 | -4.24113000 | 0.09694200 |

1 17 1.0

2 22 1.0 26 1.0
3 33 1.5 34 1.0
4 33 2.0
5 38 2.0
6 7 1.0 41 1.0
7
8 44 1.5 45 1.0
9 44 2.0
10 23 1.0 31 1.0 38 1.5
11 12 1.0 39 1.0 44 1.5
12
13 14 1.0 15 1.5 22 1.5
14
15 16 1.0 17 2.0
16
17 18 1.5
18 19 1.0 20 2.0
19
20 21 1.0 22 1.5
21
22
23 24 1.0 25 1.0 26 1.0
24
25
26 27 1.0 28 1.0
27
28 29 1.0 30 1.0 31 1.0
29
30
31 32 1.0 33 1.0
32
33
34 35 1.0 36 1.0 37 1.0
35
36
37
38 39 1.0
39 40 1.0 41 1.0
40
41 42 1.0 43 1.0
42
43
44
45 46 1.0 50 1.0 54 1.0
46 47 1.0 48 1.0 49 1.0
47
48

49
50 51 1.0 52 1.0 53 1.0
51
52
53
54 55 1.0 56 1.0 57 1.0
55
56
57

**Structure of Ac-Ser-*trans*-hyp(4-I-Ph)-OMe with Ser in the β conformation subjected to full geometry optimization in the conformation observed crystallographically**
optimized M06-2X/Def2TZVP/H2O

0 1

| | | | |
|---|---|---|---|
| I | 6.36653800 | -0.04937000 | -0.08588600 |
| O | 0.19386700 | -0.46321700 | -0.75223400 |
| O | -1.46252200 | 1.89093300 | 0.69816500 |
| O | -0.05368300 | 1.15854100 | 2.27021500 |
| O | -4.27189900 | 0.56900000 | 1.29388100 |
| N | -2.56749000 | -0.49343500 | 0.27524400 |
| C | 2.27786700 | -1.51404100 | -0.05715900 |
| H | 1.80580700 | -2.43971600 | 0.23822800 |
| C | 3.65763000 | -1.39478200 | 0.07438000 |
| H | 4.22480000 | -2.22875200 | 0.46424600 |
| C | 4.29089800 | -0.21952200 | -0.28918500 |
| C | 3.55699300 | 0.85050100 | -0.78874600 |
| H | 4.04719100 | 1.77127900 | -1.07338400 |
| C | 2.18597600 | 0.73355500 | -0.92178100 |
| H | 1.59661700 | 1.55216800 | -1.31371200 |
| C | 1.53994300 | -0.44769600 | -0.55973600 |
| C | -1.98881200 | -1.34747900 | -0.76901400 |
| H | -1.98564200 | -0.86906000 | -1.74682100 |
| H | -2.52229900 | -2.29779400 | -0.83255700 |
| C | -0.56469600 | -1.57614700 | -0.28479600 |
| H | -0.15291700 | -2.50828700 | -0.67022400 |
| C | -0.69830600 | -1.52301700 | 1.23474600 |
| H | 0.24204300 | -1.34250700 | 1.74898300 |
| H | -1.13214400 | -2.45389800 | 1.59780000 |
| C | -1.68980900 | -0.37467100 | 1.42600600 |
| H | -2.26118400 | -0.46764800 | 2.34994500 |
| C | -0.96822300 | 0.96846200 | 1.50824100 |
| C | -0.84581800 | 3.18055100 | 0.76888900 |
| H | 0.21246700 | 3.10261600 | 0.52723800 |
| H | -0.96103800 | 3.59494400 | 1.76866100 |
| H | -1.36153400 | 3.79478000 | 0.03836800 |
| C | -3.81776300 | 0.01293700 | 0.30789600 |

| | | | |
|---|---|---|---|
| O | -4.65939100 | 2.22895900 | -1.39741200 |
| H | -4.55211900 | 2.92691900 | -2.05090300 |
| O | -6.06175800 | -2.08184800 | 0.21244000 |
| N | -6.03336200 | -0.04006500 | -0.72686000 |
| H | -6.46249900 | 0.87214700 | -0.76624900 |
| C | -4.62198600 | -0.10900800 | -0.99285700 |
| H | -4.40373900 | -1.06845100 | -1.46202400 |
| C | -4.23668700 | 1.00199500 | -1.96196600 |
| H | -3.15511100 | 0.99620600 | -2.12018500 |
| H | -4.73429000 | 0.81869300 | -2.91740000 |
| C | -6.64519400 | -1.03876000 | -0.05286500 |
| C | -8.07611500 | -0.79669000 | 0.34728700 |
| H | -8.66161100 | -1.68376500 | 0.11459000 |
| H | -8.50912900 | 0.07109200 | -0.14474900 |
| H | -8.10652800 | -0.64593400 | 1.42693800 |

1 11 1.0
2 16 1.0 20 1.0
3 27 1.5 28 1.0
4 27 2.0
5 32 2.0
6 17 1.0 25 1.0 32 1.5
7 8 1.0 9 1.5 16 1.5
8
9 10 1.0 11 2.0
10
11 12 1.5
12 13 1.0 14 2.0
13
14 15 1.0 16 1.5
15
16
17 18 1.0 19 1.0 20 1.0
18
19
20 21 1.0 22 1.0
21
22 23 1.0 24 1.0 25 1.0
23
24
25 26 1.0 27 1.0
26
27
28 29 1.0 30 1.0 31 1.0
29
30
31

32 38 1.0
33 34 1.0 40 1.0
34
35 43 2.0
36 37 1.0 38 1.0 43 1.5
37
38 39 1.0 40 1.0
39
40 41 1.0 42 1.0
41
42
43 44 1.0
44 45 1.0 46 1.0 47 1.0
45
46
47

**Structure of Ac-Ser-*trans*-hyp(4-I-Ph)-OMe with Ser in the PPII conformation subjected to full geometry optimization in the conformation observed crystallographically**
optimized M06-2X/Def2TZVP/H2O

0 1

| | | | |
|---|---|---|---|
| I | -6.38781600 | -0.20043200 | 0.23857000 |
| O | -0.18760900 | -0.71994300 | 0.18611900 |
| O | 1.34545500 | 2.07170100 | 0.26295500 |
| O | -0.13596800 | 2.35544800 | -1.38611500 |
| O | 4.10091500 | 1.50856600 | -1.06995700 |
| N | 2.49319100 | -0.05581500 | -0.83958300 |
| C | -2.31994000 | -1.24982700 | -0.86481000 |
| H | -1.87471700 | -1.82142000 | -1.66616000 |
| C | -3.70458700 | -1.12110000 | -0.82381000 |
| H | -4.30151300 | -1.59525600 | -1.59064500 |
| C | -4.30461000 | -0.39173300 | 0.18728100 |
| C | -3.53239700 | 0.21662400 | 1.17050700 |
| H | -3.99665700 | 0.78614800 | 1.96376100 |
| C | -2.15635200 | 0.09001800 | 1.13170100 |
| H | -1.53665300 | 0.55039800 | 1.89006900 |
| C | -1.54377600 | -0.64346700 | 0.11680500 |
| C | 1.98735000 | -1.37272500 | -0.43262900 |
| H | 2.05532400 | -1.52846800 | 0.64259100 |
| H | 2.52353700 | -2.16934000 | -0.95029000 |
| C | 0.53233000 | -1.33877600 | -0.87662700 |
| H | 0.14633700 | -2.33921400 | -1.06916600 |
| C | 0.55997100 | -0.42598100 | -2.10034200 |
| H | -0.41353800 | -0.02154800 | -2.36476500 |
| H | 0.96577000 | -0.96821400 | -2.95343200 |
| C | 1.53739300 | 0.66567200 | -1.66210100 |

| | | | |
|---|---|---|---|
| H | 2.03987900 | 1.13850300 | -2.50602600 |
| C | 0.81450800 | 1.78420200 | -0.91450500 |
| C | 0.71991100 | 3.13727700 | 0.98612300 |
| H | -0.32742500 | 2.90207500 | 1.16573100 |
| H | 0.79304100 | 4.06439800 | 0.42073700 |
| H | 1.26125300 | 3.21635800 | 1.92286900 |
| C | 3.71336400 | 0.44903800 | -0.59547100 |
| O | 4.93053500 | -0.54265100 | 2.79175000 |
| H | 5.86241300 | -0.33334900 | 2.65627500 |
| O | 6.73020500 | -2.12795600 | 0.31190200 |
| N | 5.97005100 | -0.00538800 | 0.13805000 |
| H | 6.15992200 | 0.93087700 | -0.18996200 |
| C | 4.58395000 | -0.31529300 | 0.39281800 |
| H | 4.46174200 | -1.39410200 | 0.30439700 |
| C | 4.18310100 | 0.13032400 | 1.80765600 |
| H | 4.31346100 | 1.21584300 | 1.87918100 |
| H | 3.13286000 | -0.09765400 | 1.99050600 |
| C | 6.93795100 | -0.95017200 | 0.05877800 |
| C | 8.29980400 | -0.45743600 | -0.35795800 |
| H | 9.03071300 | -0.79868400 | 0.37308500 |
| H | 8.35212100 | 0.62478400 | -0.44895800 |
| H | 8.54758800 | -0.91198700 | -1.31701500 |

1 11 1.0
2 16 1.0 20 1.0
3 27 1.5 28 1.0
4 27 2.0
5 32 2.0
6 17 1.0 25 1.0 32 1.5
7 8 1.0 9 1.5 16 1.5
8
9 10 1.0 11 2.0
10
11 12 1.5
12 13 1.0 14 2.0
13
14 15 1.0 16 1.5
15
16
17 18 1.0 19 1.0 20 1.0
18
19
20 21 1.0 22 1.0
21
22 23 1.0 24 1.0 25 1.0
23
24

25 26 1.0 27 1.0
26
27
28 29 1.0 30 1.0 31 1.0
29
30
31
32 38 1.0
33 34 1.0 40 1.0
34
35 43 2.0
36 37 1.0 38 1.0 43 1.5
37
38 39 1.0 40 1.0
39
40 41 1.0 42 1.0
41
42
43 44 1.0
44 45 1.0 46 1.0 47 1.0
45
46
47

### Ac-SP-OMe and Ac-SP-NHMe structures Geometry-optimized from conformations observed in the crystal structures of Boc-Ser-hyp(4-I-Ph)-OMe and Ac-Ser-hyp(4-I-Ph)-OMe

#### Geometry-optimized structure of Ac-S-*cis*-P-OMe (β-*cis*-δ)

optimized M06-2X/6-311++G(d,p)/H2O

0 1

| | | | |
|---|---|---|---|
| O | -0.18900000 | -2.43100000 | 1.86700000 |
| C | -1.32000000 | -2.08100000 | 1.09200000 |
| C | -1.07100000 | -0.82500000 | 0.23800000 |
| H | -2.13300000 | -1.85900000 | 1.78500000 |
| N | -2.27000000 | -0.46600000 | -0.48500000 |
| C | 0.02600000 | -1.11100000 | -0.78400000 |
| H | -0.82000000 | 0.00300000 | 0.89800000 |
| C | -3.17100000 | 0.41400000 | 0.00900000 |
| N | 1.29100000 | -0.79600000 | -0.45900000 |
| O | -0.26400000 | -1.66700000 | -1.84000000 |
| O | -3.03800000 | 0.93300000 | 1.11200000 |
| C | 2.39100000 | -1.04600000 | -1.40800000 |
| C | 1.72500000 | -0.01900000 | 0.69900000 |
| C | 3.55000000 | -0.23000000 | -0.83400000 |
| H | 2.61300000 | -2.11600000 | -1.43000000 |
| H | 2.09400000 | -0.73500000 | -2.40900000 |
| C | 3.25000000 | -0.19300000 | 0.66700000 |
| C | 1.34800000 | 1.46100000 | 0.62400000 |
| H | 1.29600000 | -0.40900000 | 1.62300000 |
| H | 4.51800000 | -0.67300000 | -1.06200000 |
| H | 3.50100000 | -1.14800000 | 1.13200000 |
| H | 3.75900000 | 0.60500000 | 1.20400000 |
| O | 1.68400000 | 2.25300000 | 1.46800000 |
| C | 0.18400000 | 3.13800000 | -0.53200000 |
| H | -0.41200000 | 3.19500000 | -1.43800000 |
| H | -0.41400000 | 3.40000000 | 0.34000000 |
| H | 1.05100000 | 3.79500000 | -0.59800000 |
| H | -2.38800000 | -0.86800000 | -1.40600000 |
| C | -4.35000000 | 0.72500000 | -0.88000000 |
| H | -5.26600000 | 0.51400000 | -0.32800000 |
| H | -4.33400000 | 1.79100000 | -1.11100000 |
| H | -4.34700000 | 0.15500000 | -1.80800000 |
| H | 3.53200000 | 0.78400000 | -1.24400000 |
| H | 0.42200000 | -2.94300000 | 1.32800000 |
| H | -1.63100000 | -2.90300000 | 0.43900000 |
| O | 0.61100000 | 1.77000000 | -0.43700000 |

1 2 1.0 34 1.0

2 3 1.0 4 1.0 35 1.0
3 5 1.0 6 1.0 7 1.0
4
5 8 1.5 28 1.0
6 9 1.5 10 2.0
7
8 11 2.0 29 1.0
9 12 1.0 13 1.0
10
11
12 14 1.0 15 1.0 16 1.0
13 17 1.0 18 1.0 19 1.0
14 17 1.0 20 1.0 33 1.0
15
16
17 21 1.0 22 1.0
18 23 2.0 36 1.5
19
20
21
22
23
24 25 1.0 26 1.0 27 1.0 36 1.0
25
26
27
28
29 30 1.0 31 1.0 32 1.0
30
31
32
33
34
35
36

**Geometry-optimized structure of Ac-S-*trans*-P-OMe (PPII-*trans*-PPII)**
optimized M06-2X/6-311++G(d,p)/H2O

0 1

| | | | |
|---|---|---|---|
| O | -2.45500000 | 1.20500000 | -1.49000000 |
| O | -0.10800000 | -1.04600000 | -0.87800000 |
| O | 2.44200000 | -1.15500000 | 0.91700000 |
| O | -2.65900000 | -1.36600000 | 2.58800000 |
| N | -2.77500000 | -0.52700000 | -0.09900000 |
| H | -3.35800000 | -1.29400000 | 0.20100000 |
| N | 0.53500000 | 0.82800000 | 0.18500000 |

| | | | |
|---|---|---|---|
| C | -3.11300000 | 0.21800000 | -1.17400000 |
| C | -1.65200000 | -0.15200000 | 0.72700000 |
| H | -1.83700000 | 0.83000000 | 1.16900000 |
| C | -1.47100000 | -1.18600000 | 1.84200000 |
| H | -1.23800000 | -2.15300000 | 1.39300000 |
| C | -0.35700000 | -0.14100000 | -0.08800000 |
| C | 0.33600000 | 2.05200000 | 0.97900000 |
| H | 0.39700000 | 1.83000000 | 2.04900000 |
| H | -0.63200000 | 2.50000000 | 0.75900000 |
| C | 1.49800000 | 2.93600000 | 0.52500000 |
| H | 1.77600000 | 3.66900000 | 1.28100000 |
| H | 1.22200000 | 3.46500000 | -0.39000000 |
| C | 2.61200000 | 1.92900000 | 0.23500000 |
| H | 3.05500000 | 1.58400000 | 1.17300000 |
| H | 3.40100000 | 2.31800000 | -0.40600000 |
| C | 1.85700000 | 0.76400000 | -0.42800000 |
| H | 1.78700000 | 0.88900000 | -1.51200000 |
| C | 2.52900000 | -0.56500000 | -0.13000000 |
| C | -4.31400000 | -0.23500000 | -1.96300000 |
| H | -4.99200000 | 0.61000000 | -2.08200000 |
| H | -3.97800000 | -0.54000000 | -2.95500000 |
| H | -4.84100000 | -1.06400000 | -1.49400000 |
| C | 4.02500000 | -2.17400000 | -0.94800000 |
| H | 3.34200000 | -2.99600000 | -0.73900000 |
| H | 4.56200000 | -2.34800000 | -1.87600000 |
| H | 4.72200000 | -2.05400000 | -0.11900000 |
| H | -0.63300000 | -0.88500000 | 2.47800000 |
| H | -2.83100000 | -0.57000000 | 3.10000000 |
| O | 3.28500000 | -0.96200000 | -1.15000000 |

1 8 2.0
2 13 2.0
3 25 2.0
4 11 1.0 35 1.0
5 6 1.0 8 1.5 9 1.0
6
7 13 1.5 14 1.0 23 1.0
8 26 1.0
9 10 1.0 11 1.0 13 1.0
10
11 12 1.0 34 1.0
12
13
14 15 1.0 16 1.0 17 1.0
15
16
17 18 1.0 19 1.0 20 1.0

18
19
20 21 1.0 22 1.0 23 1.0
21
22
23 24 1.0 25 1.0
24
25 36 1.5
26 27 1.0 28 1.0 29 1.0
27
28
29
30 31 1.0 32 1.0 33 1.0 36 1.0
31
32
33
34
35
36

**Geometry-optimized structure of Ac-S-*trans*-P-OMe (PPII-*trans*-δ)**
optimized M06-2X/6-311++G(d,p)/H2O

0 1

| | | | |
|---|---|---|---|
| O | 1.19790000 | 2.49550000 | 0.94150000 |
| C | 1.16080000 | 1.33180000 | 1.75140000 |
| C | 1.54690000 | 0.14540000 | 0.87080000 |
| H | 0.16020000 | 1.17860000 | 2.16590000 |
| N | 2.81480000 | 0.42060000 | 0.24140000 |
| C | 0.46060000 | -0.11690000 | -0.18260000 |
| H | 1.65470000 | -0.73930000 | 1.50140000 |
| C | 3.44310000 | -0.54710000 | -0.46990000 |
| N | -0.58650000 | -0.86940000 | 0.21160000 |
| O | 0.52220000 | 0.36350000 | -1.30740000 |
| O | 3.04550000 | -1.70630000 | -0.46210000 |
| C | -0.79520000 | -1.53300000 | 1.50990000 |
| C | -1.70130000 | -1.06200000 | -0.70460000 |
| C | -2.29560000 | -1.82380000 | 1.50320000 |
| H | -0.20870000 | -2.45570000 | 1.55470000 |
| H | -0.50550000 | -0.89230000 | 2.34130000 |
| C | -2.59390000 | -2.08340000 | 0.02410000 |
| C | -2.45200000 | 0.23190000 | -0.98920000 |
| H | -1.35640000 | -1.43030000 | -1.67110000 |
| H | -2.55290000 | -2.66760000 | 2.14160000 |
| H | -2.27440000 | -3.08830000 | -0.25820000 |
| H | -3.64590000 | -1.97210000 | -0.23690000 |
| O | -3.12200000 | 0.39540000 | -1.97870000 |

| | | | |
|---|---|---|---|
| C | -3.04060000 | 2.36290000 | -0.20780000 |
| H | -2.80740000 | 2.97320000 | 0.65970000 |
| H | -2.69950000 | 2.84860000 | -1.12160000 |
| H | -4.11210000 | 2.17450000 | -0.26840000 |
| H | 3.03110000 | 1.39070000 | 0.05730000 |
| C | 4.65000000 | -0.11470000 | -1.26350000 |
| H | 5.44090000 | -0.85220000 | -1.13280000 |
| H | 4.37450000 | -0.09640000 | -2.32020000 |
| H | 5.01220000 | 0.87080000 | -0.97410000 |
| H | -2.84100000 | -0.94360000 | 1.84980000 |
| H | 1.07170000 | 3.26670000 | 1.50110000 |
| H | 1.88000000 | 1.40620000 | 2.57240000 |
| O | -2.33320000 | 1.12960000 | -0.01600000 |

1 2 1.0 34 1.0
2 3 1.0 4 1.0 35 1.0
3 5 1.0 6 1.0 7 1.0
4
5 8 1.5 28 1.0
6 9 1.5 10 2.0
7
8 11 2.0 29 1.0
9 12 1.0 13 1.0
10
11
12 14 1.0 15 1.0 16 1.0
13 17 1.0 18 1.0 19 1.0
14 17 1.0 20 1.0 33 1.0
15
16
17 21 1.0 22 1.0
18 23 2.0 36 1.5
19
20
21
22
23
24 25 1.0 26 1.0 27 1.0 36 1.0
25
26
27
28
29 30 1.0 31 1.0 32 1.0
30
31
32
33

34
35
36

**Geometry-optimized structure of Ac-S-*trans*-P-OMe (β-*trans*-δ)**
optimized M06-2X/6-311++G(d,p)/H2O

0 1

| | | | |
|---|---|---|---|
| O | -2.01090000 | 1.48760000 | 1.93740000 |
| C | -1.34920000 | 0.35640000 | 1.41800000 |
| C | -1.44790000 | 0.28060000 | -0.13050000 |
| H | -1.75620000 | -0.57580000 | 1.82530000 |
| N | -2.51870000 | -0.58120000 | -0.58960000 |
| C | -0.14430000 | -0.26590000 | -0.69800000 |
| H | -1.61740000 | 1.29580000 | -0.50270000 |
| C | -3.80820000 | -0.32840000 | -0.30770000 |
| N | 0.88810000 | 0.59020000 | -0.74750000 |
| O | -0.04270000 | -1.44070000 | -1.04140000 |
| O | -4.14330000 | 0.64320000 | 0.37550000 |
| C | 0.89880000 | 2.00860000 | -0.34760000 |
| C | 2.20570000 | 0.09770000 | -1.12550000 |
| C | 2.38420000 | 2.27800000 | -0.10930000 |
| H | 0.50670000 | 2.62440000 | -1.16240000 |
| H | 0.29270000 | 2.17680000 | 0.54230000 |
| C | 3.07200000 | 1.37180000 | -1.13480000 |
| C | 2.74360000 | -0.93320000 | -0.14160000 |
| H | 2.17890000 | -0.39320000 | -2.09880000 |
| H | 2.63440000 | 3.32960000 | -0.24090000 |
| H | 3.02200000 | 1.81800000 | -2.12970000 |
| H | 4.11580000 | 1.16060000 | -0.90450000 |
| O | 3.57550000 | -1.75220000 | -0.44180000 |
| C | 2.72300000 | -1.71800000 | 2.06920000 |
| H | 2.19460000 | -1.46950000 | 2.98480000 |
| H | 2.50070000 | -2.73980000 | 1.76310000 |
| H | 3.79790000 | -1.59740000 | 2.20050000 |
| H | -2.24590000 | -1.42050000 | -1.08480000 |
| C | -4.82870000 | -1.27850000 | -0.87570000 |
| H | -5.45790000 | -1.64010000 | -0.06220000 |
| H | -5.46220000 | -0.72390000 | -1.56960000 |
| H | -4.37850000 | -2.12240000 | -1.39540000 |
| H | 2.65500000 | 1.97860000 | 0.90520000 |
| H | -2.91100000 | 1.45380000 | 1.57550000 |
| H | -0.30400000 | 0.42750000 | 1.72650000 |
| O | 2.24080000 | -0.79560000 | 1.08130000 |

1 2 1.0 34 1.0
2 3 1.0 4 1.0 35 1.0

3 5 1.0 6 1.0 7 1.0
4
5 8 1.5 28 1.0
6 9 1.5 10 2.0
7
8 11 2.0 29 1.0
9 12 1.0 13 1.0
10
11
12 14 1.0 15 1.0 16 1.0
13 17 1.0 18 1.0 19 1.0
14 17 1.0 20 1.0 33 1.0
15
16
17 21 1.0 22 1.0
18 23 2.0 36 1.5
19
20
21
22
23
24 25 1.0 26 1.0 27 1.0 36 1.0
25
26
27
28
29 30 1.0 31 1.0 32 1.0
30
31
32
33
34
35
36

**Geometry-optimized structure of Ac-S-*cis*-P-NHMe (PPII-*cis*-δ)**
optimized M06-2X/6-311++G(d,p)/H2O

0 1

| | | | |
|---|---|---|---|
| O | 0.14100000 | -2.67700000 | -1.73600000 |
| C | 1.40500000 | -2.11200000 | -1.45300000 |
| C | 1.27500000 | -0.76900000 | -0.72100000 |
| H | 1.90800000 | -1.94500000 | -2.40700000 |
| N | 2.60400000 | -0.25600000 | -0.46700000 |
| C | 0.53500000 | -0.98500000 | 0.60800000 |
| H | 0.73500000 | -0.06000000 | -1.35300000 |
| C | 2.78800000 | 0.87500000 | 0.24700000 |

| | | | |
|---|---|---|---|
| N | -0.77900000 | -0.68700000 | 0.65800000 |
| O | 1.13500000 | -1.43900000 | 1.57400000 |
| O | 1.83300000 | 1.55300000 | 0.62300000 |
| C | -1.52800000 | -0.98500000 | 1.89400000 |
| C | -1.67900000 | -0.34400000 | -0.45100000 |
| C | -2.97800000 | -0.67700000 | 1.52200000 |
| H | -1.38500000 | -2.03900000 | 2.14700000 |
| H | -1.14900000 | -0.38100000 | 2.72000000 |
| C | -3.01200000 | -0.94100000 | 0.01600000 |
| C | -1.82500000 | 1.16300000 | -0.71200000 |
| H | -1.34400000 | -0.80400000 | -1.38200000 |
| H | -3.68100000 | -1.29000000 | 2.08300000 |
| H | -3.00800000 | -2.01500000 | -0.18400000 |
| H | -3.85500000 | -0.48500000 | -0.49800000 |
| N | -0.91400000 | 1.97400000 | -0.16400000 |
| O | -2.74300000 | 1.56500000 | -1.42200000 |
| C | -0.92100000 | 3.39800000 | -0.45100000 |
| H | -0.05200000 | 3.84700000 | 0.02500000 |
| H | -0.87800000 | 3.57700000 | -1.52700000 |
| H | -1.82700000 | 3.86400000 | -0.06000000 |
| H | -0.10400000 | 1.59200000 | 0.31500000 |
| H | 3.40300000 | -0.84200000 | -0.66300000 |
| C | 4.20900000 | 1.25500000 | 0.56600000 |
| H | 4.34300000 | 2.31600000 | 0.36000000 |
| H | 4.37200000 | 1.09300000 | 1.63400000 |
| H | 4.93800000 | 0.67500000 | 0.00200000 |
| H | -3.20500000 | 0.37400000 | 1.72200000 |
| H | -0.15900000 | -3.18900000 | -0.97800000 |
| H | 2.01700000 | -2.78900000 | -0.84800000 |

1 2 1.0 36 1.0
2 3 1.0 4 1.0 37 1.0
3 5 1.0 6 1.0 7 1.0
4
5 8 1.5 30 1.0
6 9 1.5 10 2.0
7
8 11 2.0 31 1.0
9 12 1.0 13 1.0
10
11
12 14 1.0 15 1.0 16 1.0
13 17 1.0 18 1.0 19 1.0
14 17 1.0 20 1.0 35 1.0
15
16
17 21 1.0 22 1.0

18 23 1.5 24 2.0
19
20
21
22
23 25 1.0 29 1.0
24
25 26 1.0 27 1.0 28 1.0
26
27
28
29
30
31 32 1.0 33 1.0 34 1.0
32
33
34
35
36
37

**Geometry-optimized structure of Ac-S-*trans*-P-NHMe (PPII-*trans*-PPII)**
optimized M06-2X/6-311++G(d,p)/H2O

0 1

| | | | |
|---|---|---|---|
| O | -2.49500000 | 1.18100000 | -1.49200000 |
| O | -0.11900000 | -1.04000000 | -0.89200000 |
| O | 2.46000000 | -1.05400000 | 0.99100000 |
| O | -2.63300000 | -1.37200000 | 2.60000000 |
| N | -2.77800000 | -0.54700000 | -0.08800000 |
| H | -3.34400000 | -1.32600000 | 0.21500000 |
| N | 0.52300000 | 0.84200000 | 0.16300000 |
| N | 3.32000000 | -1.03100000 | -1.09500000 |
| C | -3.13300000 | 0.18500000 | -1.16600000 |
| C | -1.65200000 | -0.15500000 | 0.72600000 |
| H | -1.84500000 | 0.82700000 | 1.16400000 |
| C | -1.45200000 | -1.18000000 | 1.84600000 |
| H | -1.20700000 | -2.14700000 | 1.40100000 |
| C | -0.36200000 | -0.13600000 | -0.10000000 |
| C | 0.31300000 | 2.06200000 | 0.95900000 |
| H | 0.38600000 | 1.84100000 | 2.02800000 |
| H | -0.66400000 | 2.49600000 | 0.74800000 |
| C | 1.45800000 | 2.96300000 | 0.49700000 |
| H | 1.72600000 | 3.70400000 | 1.24900000 |
| H | 1.17000000 | 3.48300000 | -0.41900000 |
| C | 2.58400000 | 1.96900000 | 0.21000000 |
| H | 3.03600000 | 1.63700000 | 1.14800000 |

| | | | |
|---|---|---|---|
| H | 3.36600000 | 2.36700000 | -0.43600000 |
| C | 1.85100000 | 0.78500000 | -0.44000000 |
| H | 1.77200000 | 0.90900000 | -1.52500000 |
| C | 2.54900000 | -0.53500000 | -0.11400000 |
| C | -4.33100000 | -0.29400000 | -1.94600000 |
| H | -5.02100000 | 0.54000000 | -2.07400000 |
| H | -3.99300000 | -0.60700000 | -2.93500000 |
| H | -4.84600000 | -1.12500000 | -1.46500000 |
| C | 4.09600000 | -2.24500000 | -0.90700000 |
| H | 3.44200000 | -3.08600000 | -0.66900000 |
| H | 4.63700000 | -2.45800000 | -1.82500000 |
| H | 4.81000000 | -2.11800000 | -0.09100000 |
| H | -0.61500000 | -0.86400000 | 2.47600000 |
| H | 3.30800000 | -0.58900000 | -2.00100000 |
| H | -2.81400000 | -0.57400000 | 3.10700000 |

1 9 2.0
2 14 2.0
3 26 2.0
4 12 1.0 37 1.0
5 6 1.0 9 1.5 10 1.0
6
7 14 1.5 15 1.0 24 1.0
8 26 1.5 31 1.0 36 1.0
9 27 1.0
10 11 1.0 12 1.0 14 1.0
11
12 13 1.0 35 1.0
13
14
15 16 1.0 17 1.0 18 1.0
16
17
18 19 1.0 20 1.0 21 1.0
19
20
21 22 1.0 23 1.0 24 1.0
22
23
24 25 1.0 26 1.0
25
26
27 28 1.0 29 1.0 30 1.0
28
29
30
31 32 1.0 33 1.0 34 1.0

32
33
34
35
36
37

**Geometry-optimized structure of Ac-S-*trans*-P-NHMe (PPII-*trans*-δ)**
optimized M06-2X/6-311++G(d,p)/H2O

0 1

| | | | |
|---|---|---|---|
| O | -3.90268900 | 0.27460200 | -1.50969000 |
| O | 0.15828700 | 0.58496100 | -1.57290800 |
| N | -0.76665800 | -0.68063900 | 0.05526700 |
| C | -0.68678600 | -1.55202100 | 1.24667300 |
| H | -0.32627300 | -1.00188100 | 2.11563300 |
| H | -0.01267200 | -2.39075000 | 1.05037000 |
| C | -2.61174200 | -2.14200100 | -0.04401200 |
| H | -3.69580700 | -2.15589200 | -0.14622800 |
| H | -2.20788700 | -3.04424700 | -0.50715900 |
| C | -1.98899400 | -0.90844100 | -0.72098400 |
| H | -1.73931000 | -1.09149100 | -1.76664600 |
| C | -2.96224800 | 0.27142200 | -0.72217100 |
| C | -3.64656800 | 2.39170900 | 0.26343900 |
| H | -4.66989500 | 2.06895500 | 0.45938200 |
| H | -3.63219800 | 2.95599100 | -0.67061800 |
| H | -3.30933200 | 3.03157400 | 1.07523800 |
| C | 0.25366400 | 0.01397400 | -0.49323600 |
| O | 2.51292700 | 1.50260200 | 2.10424900 |
| H | 3.23559300 | 1.72733600 | 1.50750500 |
| O | 4.02811500 | -1.14808800 | 0.35222400 |
| N | 2.64229200 | 0.40278300 | -0.54826600 |
| H | 2.44163000 | 0.97452500 | -1.35872200 |
| C | 1.52871700 | 0.14365900 | 0.33826000 |
| H | 1.75002000 | -0.77007700 | 0.89051900 |
| C | 1.35346800 | 1.30988900 | 1.32690000 |
| H | 1.09891900 | 2.21533100 | 0.76299300 |
| H | 0.54037800 | 1.09585300 | 2.02239500 |
| C | 3.79498100 | -0.31884500 | -0.51628000 |
| C | 4.77846400 | -0.02614400 | -1.62272300 |
| H | 5.75812500 | 0.15128200 | -1.17972700 |
| H | 4.49236100 | 0.82910400 | -2.23266300 |
| H | 4.85098800 | -0.91138100 | -2.25689200 |
| C | -2.12839200 | -2.03047000 | 1.40443000 |
| H | -2.71237700 | -1.28100000 | 1.94420200 |
| H | -2.18717900 | -2.97172100 | 1.94839000 |
| N | -2.76175200 | 1.24215700 | 0.18018900 |

H -1.95276600 1.18938400 0.77891400

1 12 2.0
2 17 2.0
3 4 1.0 10 1.0 17 1.5
4 5 1.0 6 1.0 33 1.0
5
6
7 8 1.0 9 1.0 10 1.0 33 1.0
8
9
10 11 1.0 12 1.0
11
12 36 1.5
13 14 1.0 15 1.0 16 1.0 36 1.0
14
15
16
17 23 1.0
18 19 1.0 25 1.0
19
20 28 2.0
21 22 1.0 23 1.0 28 1.5
22
23 24 1.0 25 1.0
24
25 26 1.0 27 1.0
26
27
28 29 1.0
29 30 1.0 31 1.0 32 1.0
30
31
32
33 34 1.0 35 1.0
34
35
36 37 1.0
37

**Geometry-optimized structure of Ac-S-*trans*-P-NHMe (β-*trans*-δ)**
optimized M06-2X/6-311++G(d,p)/H2O

0 1
O -3.60448500 0.25997100 -1.53288500
O 0.66354400 0.32501900 -1.32243900
N -0.58148900 -0.82879800 0.16626800

| | | | |
|---|---|---|---|
| C | -0.72690900 | -1.71990700 | 1.33590800 |
| H | -0.47435100 | -1.21607700 | 2.26648700 |
| H | -0.07589800 | -2.59143100 | 1.21795500 |
| C | -2.47478300 | -2.17368400 | -0.23111200 |
| H | -3.53055500 | -2.12293200 | -0.49019400 |
| H | -2.05233200 | -3.08525000 | -0.65852200 |
| C | -1.70224000 | -0.96043800 | -0.76648700 |
| H | -1.33173600 | -1.12212500 | -1.78022600 |
| C | -2.59218400 | 0.28745000 | -0.83777700 |
| C | -2.98434100 | 2.60153800 | -0.21519500 |
| H | -4.01374300 | 2.43660600 | 0.10538000 |
| H | -2.99735000 | 2.98381900 | -1.23811100 |
| H | -2.51915900 | 3.33712200 | 0.43750200 |
| C | 0.53875300 | -0.17095900 | -0.21326800 |
| O | 0.59424300 | 2.17388400 | 1.13789000 |
| H | 0.43889700 | 2.91083900 | 1.73667000 |
| O | 3.33083500 | -1.54562300 | -0.53438500 |
| N | 2.83865000 | 0.49598900 | 0.28527700 |
| H | 2.92676700 | 1.49049300 | 0.12537600 |
| C | 1.60588400 | 0.03669400 | 0.87736000 |
| H | 1.79825800 | -0.90436400 | 1.39643300 |
| C | 1.09947900 | 1.07032600 | 1.88126300 |
| H | 0.31195000 | 0.64501900 | 2.50606500 |
| H | 1.93008200 | 1.37371700 | 2.52320200 |
| C | 3.57543400 | -0.34713400 | -0.48362500 |
| C | 4.70660300 | 0.27884000 | -1.25911300 |
| H | 5.57449200 | -0.37721100 | -1.21120700 |
| H | 4.97000500 | 1.26923100 | -0.89065100 |
| H | 4.39827400 | 0.35988000 | -2.30371500 |
| N | -2.22440500 | 1.36627900 | -0.13212300 |
| H | -1.33354700 | 1.38200200 | 0.34773300 |
| C | -2.19950000 | -2.11947900 | 1.27303200 |
| H | -2.38770200 | -3.06430500 | 1.78047100 |
| H | -2.81191500 | -1.34482400 | 1.74202100 |

1 12 2.0
2 17 2.0
3 4 1.0 10 1.0 17 1.5
4 5 1.0 6 1.0 35 1.0
5
6
7 8 1.0 9 1.0 10 1.0 35 1.0
8
9
10 11 1.0 12 1.0
11
12 33 1.5

13 14 1.0 15 1.0 16 1.0 33 1.0
14
15
16
17 23 1.0
18 19 1.0 25 1.0
19
20 28 2.0
21 22 1.0 23 1.0 28 1.5
22
23 24 1.0 25 1.0
24
25 26 1.0 27 1.0
26
27
28 29 1.0
29 30 1.0 31 1.0 32 1.0
30
31
32
33 34 1.0
34
35 36 1.0 37 1.0
36
37

### Geometry-optimized structures of minimal Ser-*cis*-Pro type VI β-turns with C–H/O interactions

#### Geometry-optimized structure of Ac-Ser-*cis*-Pro-NHMe in a type VIa1 (PcisD) β-turn with a C–H/O interaction

optimized M06-2X/6-311++G(d,p)/H2O

0 1

| | | | |
|---|---|---|---|
| O | 0.14144900 | -2.67698800 | -1.73638700 |
| C | 1.40522800 | -2.11199900 | -1.45291900 |
| C | 1.27508000 | -0.76855700 | -0.72140400 |
| H | 1.90773900 | -1.94491100 | -2.40684600 |
| N | 2.60449200 | -0.25636400 | -0.46652900 |
| C | 0.53525500 | -0.98476700 | 0.60811000 |
| H | 0.73506700 | -0.05968900 | -1.35337800 |
| C | 2.78752600 | 0.87524000 | 0.24669200 |
| N | -0.77882100 | -0.68683600 | 0.65839100 |
| O | 1.13522500 | -1.43929600 | 1.57443300 |
| O | 1.83314700 | 1.55298800 | 0.62257800 |
| C | -1.52766800 | -0.98471200 | 1.89437600 |
| C | -1.67910100 | -0.34442300 | -0.45146400 |
| C | -2.97801900 | -0.67665400 | 1.52246400 |
| H | -1.38505400 | -2.03865900 | 2.14736100 |
| H | -1.14916500 | -0.38056500 | 2.71961700 |
| C | -3.01175400 | -0.94120800 | 0.01570600 |
| C | -1.82478900 | 1.16257100 | -0.71187800 |
| H | -1.34358400 | -0.80395000 | -1.38215000 |
| H | -3.68145000 | -1.29024000 | 2.08323300 |
| H | -3.00804700 | -2.01543300 | -0.18395100 |
| H | -3.85524200 | -0.48511300 | -0.49803500 |
| N | -0.91382600 | 1.97420700 | -0.16386100 |
| O | -2.74281700 | 1.56536500 | -1.42241700 |
| C | -0.92127600 | 3.39804100 | -0.45089800 |
| H | -0.05220900 | 3.84678200 | 0.02540300 |
| H | -0.87832800 | 3.57657800 | -1.52727800 |
| H | -1.82738400 | 3.86391000 | -0.05992500 |
| H | -0.10447400 | 1.59206600 | 0.31506900 |
| H | 3.40299800 | -0.84241900 | -0.66296700 |
| C | 4.20867800 | 1.25453000 | 0.56625500 |
| H | 4.34275600 | 2.31581500 | 0.36048600 |
| H | 4.37192400 | 1.09254000 | 1.63350500 |
| H | 4.93756100 | 0.67494700 | 0.00211300 |
| H | -3.20535000 | 0.37408900 | 1.72206100 |
| H | -0.15862400 | -3.18865100 | -0.97843400 |
| H | 2.01696600 | -2.78906200 | -0.84779500 |

1 2 1.0 36 1.0
2 3 1.0 4 1.0 37 1.0
3 5 1.0 6 1.0 7 1.0
4

5 8 1.5 30 1.0
6 9 1.5 10 2.0
7
8 11 2.0 31 1.0
9 12 1.0 13 1.0
10
11
12 14 1.0 15 1.0 16 1.0
13 17 1.0 18 1.0 19 1.0
14 17 1.0 20 1.0 35 1.0
15
16
17 21 1.0 22 1.0
18 23 1.5 24 2.0
19
20
21
22
23 25 1.0 29 1.0
24
25 26 1.0 27 1.0 28 1.0
26
27
28
29
30
31 32 1.0 33 1.0 34 1.0
32
33
34
35
36
37

**Geometry-optimized structure of Ac-Ser-*cis*-Pro-OMe in a type VIa2 (BcisD) β-turn with a C–H/O interaction**
optimized M06-2X/6-311++G(d,p)/H2O

0 1

| | | | |
|---|---|---|---|
| O | -0.15372500 | -2.35521100 | 1.85619400 |
| C | -1.33512800 | -2.08710400 | 1.12012400 |
| C | -1.05931800 | -0.85900400 | 0.25219000 |
| H | -2.17702900 | -1.85840500 | 1.78067900 |
| N | -2.24212500 | -0.49068700 | -0.49450000 |
| C | 0.04503400 | -1.17529800 | -0.75745400 |
| H | -0.80534800 | -0.02199200 | 0.90064300 |
| C | -3.12460900 | 0.42748900 | -0.03667000 |
| N | 1.29601000 | -0.77969000 | -0.48209100 |
| O | -0.24094300 | -1.79599500 | -1.77890700 |
| O | -2.99117400 | 0.97250400 | 1.05361500 |

| | | | |
|---|---|---|---|
| C | 2.39120400 | -1.03982700 | -1.43035500 |
| C | 1.73211500 | -0.00957900 | 0.67711500 |
| C | 3.53657300 | -0.18511500 | -0.88750100 |
| H | 2.63472900 | -2.10586900 | -1.42093900 |
| H | 2.08268100 | -0.76520900 | -2.43906100 |
| C | 3.26185200 | -0.14730300 | 0.61773300 |
| C | 1.32017500 | 1.46182800 | 0.63397800 |
| H | 1.32931900 | -0.42583400 | 1.60166500 |
| H | 4.51258700 | -0.60168800 | -1.13096900 |
| H | 3.54458400 | -1.09417500 | 1.08170600 |
| H | 3.76161200 | 0.66456600 | 1.14336000 |
| O | 1.64148900 | 2.24465800 | 1.49243100 |
| C | 0.11631100 | 3.13671900 | -0.48338000 |
| H | -0.48694100 | 3.19948700 | -1.38411100 |
| H | -0.48095100 | 3.36998900 | 0.39751500 |
| H | 0.96870000 | 3.81251600 | -0.54192900 |
| H | -2.36146900 | -0.91667200 | -1.40355000 |
| C | -4.28212500 | 0.74729100 | -0.94971300 |
| H | -5.21108300 | 0.57183900 | -0.40650700 |
| H | -4.23633400 | 1.80746700 | -1.20307100 |
| H | -4.28046600 | 0.15713700 | -1.86476400 |
| H | 3.47971800 | 0.82366800 | -1.30505700 |
| H | -0.28316000 | -3.15234900 | 2.37748500 |
| H | -1.60442100 | -2.92960000 | 0.47418700 |
| O | 0.57230000 | 1.77619000 | -0.41802600 |

1 2 1.0 34 1.0
2 3 1.0 4 1.0 35 1.0
3 5 1.0 6 1.0 7 1.0
4
5 8 1.5 28 1.0
6 9 1.5 10 2.0
7
8 11 2.0 29 1.0
9 12 1.0 13 1.0
10
11
12 14 1.0 15 1.0 16 1.0
13 17 1.0 18 1.0 19 1.0
14 17 1.0 20 1.0 33 1.0
15
16
17 21 1.0 22 1.0
18 23 2.0 36 1.5
19
20
21
22
23
24 25 1.0 26 1.0 27 1.0 36 1.0
25

26
27
28
29 30 1.0 31 1.0 32 1.0
30
31
32
33
34
35
36

**Geometry-optimized structure of Ac-Ser-*cis*-Pro-OMe in a type VIb (PcisP) β-turn with a C–H/O interaction**

optimized M06-2X/6-311++G(d,p)/H2O

0 1

| | | | |
|---|---|---|---|
| O | -0.46383100 | -1.48159400 | 2.39555100 |
| C | -1.63895700 | -1.31859800 | 1.61991800 |
| C | -1.26010000 | -0.53818200 | 0.35828000 |
| H | -2.39929900 | -0.74555100 | 2.15849800 |
| N | -2.43648200 | -0.30288300 | -0.44998100 |
| C | -0.28156700 | -1.35880800 | -0.48372700 |
| H | -0.85046700 | 0.42902900 | 0.64883900 |
| C | -3.24833700 | 0.75961000 | -0.24861700 |
| N | 1.03197700 | -1.13474100 | -0.33823000 |
| O | -0.72291500 | -2.21675300 | -1.24543400 |
| O | -3.03542200 | 1.58450900 | 0.63356600 |
| C | 2.01225800 | -1.90565600 | -1.12036800 |
| C | 1.65798700 | -0.12036300 | 0.49933200 |
| C | 3.32683000 | -1.16832100 | -0.86467400 |
| H | 2.04034300 | -2.93482500 | -0.75196800 |
| H | 1.71965600 | -1.92317100 | -2.17022900 |
| C | 3.12889600 | -0.56947200 | 0.53039000 |
| C | 1.50636100 | 1.26663400 | -0.12014400 |
| H | 1.22135600 | -0.10166000 | 1.49860200 |
| H | 4.18825000 | -1.83172400 | -0.92192700 |
| H | 3.23954200 | -1.33960000 | 1.29627900 |
| H | 3.80340500 | 0.25310900 | 0.76070400 |
| O | 0.93653200 | 1.50603100 | -1.15403800 |
| C | 2.00153600 | 3.54275200 | 0.17608600 |
| H | 2.52887600 | 4.14014900 | 0.91349200 |
| H | 2.47789200 | 3.62785000 | -0.79997700 |
| H | 0.95880000 | 3.85106900 | 0.10631400 |
| H | -2.63644500 | -0.98136000 | -1.17223700 |
| C | -4.43035500 | 0.87571100 | -1.17928100 |
| H | -5.34166200 | 0.89193300 | -0.58048200 |
| H | -4.35888500 | 1.82672800 | -1.70858500 |

| | | | |
|---|---|---|---|
| H | -4.48583100 | 0.06289800 | -1.90165100 |
| H | 3.45735800 | -0.37157200 | -1.60116900 |
| H | -0.68694700 | -1.93847100 | 3.21136200 |
| H | -2.06008200 | -2.28602900 | 1.32475800 |
| O | 2.08168200 | 2.18814700 | 0.64581600 |

1 2 1.0 34 1.0
2 3 1.0 4 1.0 35 1.0
3 5 1.0 6 1.0 7 1.0
4
5 8 1.5 28 1.0
6 9 1.5 10 2.0
7
8 11 2.0 29 1.0
9 12 1.0 13 1.0
10
11
12 14 1.0 15 1.0 16 1.0
13 17 1.0 18 1.0 19 1.0
14 17 1.0 20 1.0 33 1.0
15
16
17 21 1.0 22 1.0
18 23 2.0 36 1.5
19
20
21
22
23
24 25 1.0 26 1.0 27 1.0 36 1.0
25
26
27
28
29 30 1.0 31 1.0 32 1.0
30
31
32
33
34
35
36

**Geometry-optimized structure of Ac-Ser-*cis*-Pro-OMe in a type VIb (BcisP) β-turn with a C–H/O interaction**
optimized M06-2X/6-311++G(d,p)/H2O

0 1

| | | | |
|---|---|---|---|
| O | -1.42440900 | 0.92831400 | 2.30694000 |
| C | -1.37529800 | -0.36754000 | 1.75353800 |
| C | -1.19146300 | -0.33315700 | 0.21152400 |
| H | -2.28513700 | -0.93672500 | 1.97253200 |
| N | -2.43561800 | -0.50799200 | -0.50827400 |
| C | -0.23769100 | -1.43366700 | -0.25200000 |
| H | -0.77394000 | 0.64691300 | -0.04238800 |
| C | -3.45936700 | 0.35186500 | -0.37346400 |
| N | 1.06229800 | -1.26157300 | 0.04339300 |
| O | -0.65888000 | -2.43271200 | -0.82447700 |
| O | -3.39489200 | 1.31956000 | 0.38980800 |
| C | 2.05932800 | -2.27623700 | -0.33779000 |
| C | 1.67107200 | -0.06048700 | 0.59700800 |
| C | 3.39608300 | -1.57254000 | -0.09319100 |
| H | 1.93572900 | -3.15868800 | 0.29526300 |
| H | 1.90966400 | -2.57199000 | -1.37633300 |
| C | 3.07082800 | -0.55367900 | 1.00278300 |
| C | 1.76688000 | 1.04112300 | -0.45643100 |
| H | 1.11881800 | 0.33657800 | 1.45091700 |
| H | 4.17871200 | -2.26997300 | 0.20041900 |
| H | 2.99493700 | -1.04385800 | 1.97521800 |
| H | 3.78769200 | 0.26229000 | 1.07723700 |
| O | 1.49478900 | 0.90807800 | -1.62183100 |
| C | 2.35956200 | 3.28713800 | -0.80158600 |
| H | 2.69482100 | 4.11243900 | -0.18101400 |
| H | 3.10219900 | 3.05228500 | -1.56336600 |
| H | 1.40680200 | 3.52345400 | -1.27401900 |
| H | -2.51097800 | -1.33131000 | -1.09197500 |
| C | -4.68451200 | 0.08767000 | -1.20744600 |
| H | -5.55727100 | 0.08751900 | -0.55456400 |
| H | -4.79689400 | 0.90650400 | -1.91994700 |
| H | -4.63102100 | -0.85454000 | -1.74999700 |
| H | 3.71893500 | -1.05724300 | -1.00072300 |
| H | -2.13832900 | 1.38520200 | 1.83362300 |
| H | -0.53749800 | -0.88168200 | 2.22766800 |
| O | 2.20132400 | 2.17136700 | 0.08930200 |

1 2 1.0 34 1.0
2 3 1.0 4 1.0 35 1.0
3 5 1.0 6 1.0 7 1.0
4
5 8 1.5 28 1.0
6 9 1.5 10 2.0
7
8 11 2.0 29 1.0
9 12 1.0 13 1.0

10
11
12 14 1.0 15 1.0 16 1.0
13 17 1.0 18 1.0 19 1.0
14 17 1.0 20 1.0 33 1.0
15
16
17 21 1.0 22 1.0
18 23 2.0 36 1.5
19
20
21
22
23
24 25 1.0 26 1.0 27 1.0 36 1.0
25
26
27
28
29 30 1.0 31 1.0 32 1.0
30
31
32
33
34
35
36

**Geometry-optimized structures of Ac-Ser-Pro-OMe upon rotation of Ser $\chi_1$ with hydrogen bonds not observed in the crystal structures of 3 or 5**

**Geometry-optimized structure of Ac-Ser-*cis*-Pro-OMe (BcisP) with a *g*⁻ Ser $\chi_1$ rotamer and a *g*⁺ Ser $\chi_2$ rotamer containing a Ser $O_\gamma$–H/O=$C_{i\text{-}1}$ hydrogen bond**

optimized M06-2X/6-311++G(d,p)/H2O

0 1

| | | | |
|---|---|---|---|
| O | -2.01094900 | 1.48762500 | 1.93739200 |
| C | -1.34918700 | 0.35641800 | 1.41796200 |
| C | -1.44790800 | 0.28061900 | -0.13046500 |
| H | -1.75615100 | -0.57583300 | 1.82528900 |
| N | -2.51874000 | -0.58116800 | -0.58958300 |
| C | -0.14429600 | -0.26585600 | -0.69802700 |
| H | -1.61740300 | 1.29584800 | -0.50274800 |
| C | -3.80819300 | -0.32844100 | -0.30768000 |
| N | 0.88815000 | 0.59020100 | -0.74754000 |
| O | -0.04274300 | -1.44070800 | -1.04138000 |
| O | -4.14327500 | 0.64321400 | 0.37549100 |
| C | 0.89878200 | 2.00858200 | -0.34757600 |
| C | 2.20573400 | 0.09774500 | -1.12554400 |
| C | 2.38421600 | 2.27803400 | -0.10930000 |
| H | 0.50673100 | 2.62444000 | -1.16238400 |
| H | 0.29270500 | 2.17683300 | 0.54225000 |
| C | 3.07201000 | 1.37179600 | -1.13477500 |
| C | 2.74355100 | -0.93319700 | -0.14161700 |
| H | 2.17886800 | -0.39317200 | -2.09875600 |
| H | 2.63442300 | 3.32960100 | -0.24092300 |
| H | 3.02202600 | 1.81804000 | -2.12965200 |
| H | 4.11580900 | 1.16063400 | -0.90448900 |
| O | 3.57553900 | -1.75222900 | -0.44180000 |
| C | 2.72297900 | -1.71798900 | 2.06921000 |
| H | 2.19456900 | -1.46953900 | 2.98477100 |
| H | 2.50066600 | -2.73978200 | 1.76314600 |
| H | 3.79791100 | -1.59736300 | 2.20048700 |
| H | -2.24589900 | -1.42048200 | -1.08480900 |
| C | -4.82872900 | -1.27849900 | -0.87565200 |
| H | -5.45785900 | -1.64008800 | -0.06221100 |
| H | -5.46224600 | -0.72389300 | -1.56961100 |
| H | -4.37853100 | -2.12238400 | -1.39535100 |
| H | 2.65503700 | 1.97861800 | 0.90519100 |
| H | -2.91100800 | 1.45380200 | 1.57548800 |
| H | -0.30400700 | 0.42752100 | 1.72654100 |
| O | 2.24076900 | -0.79556600 | 1.08134800 |

1 2 1.0 34 1.0

2 3 1.0 4 1.0 35 1.0
3 5 1.0 6 1.0 7 1.0
4
5 8 1.5 28 1.0
6 9 1.5 10 2.0
7
8 11 2.0 29 1.0
9 12 1.0 13 1.0
10
11
12 14 1.0 15 1.0 16 1.0
13 17 1.0 18 1.0 19 1.0
14 17 1.0 20 1.0 33 1.0
15
16
17 21 1.0 22 1.0
18 23 2.0 36 1.5
19
20
21
22
23
24 25 1.0 26 1.0 27 1.0 36 1.0
25
26
27
28
29 30 1.0 31 1.0 32 1.0
30
31
32
33
34
35
36

**Geometry-optimized structure of Ac-Ser-*cis*-Pro-OMe (BcisP) with a $g^+$ Ser $\chi_1$ rotamer and a $g^+$ Ser $\chi_2$ rotamer containing an intraresidue Ser $O_\gamma$–H/O=C hydrogen bond**
optimized M06-2X/6-311++G(d,p)/H2O

0 1

| | | | |
|---|---|---|---|
| O | -3.50318600 | -1.79877700 | -0.31113000 |
| C | -2.76096000 | -1.57162600 | 0.87249000 |
| C | -1.53324200 | -0.69264400 | 0.61366000 |
| H | -2.44004700 | -2.51435600 | 1.32785900 |
| N | -1.94332900 | 0.51467100 | -0.07937000 |

| | | | |
|---|---|---|---|
| C | -0.51517900 | -1.47133100 | -0.23117200 |
| H | -1.10615500 | -0.39357400 | 1.56992100 |
| C | -1.58725800 | 1.75726200 | 0.32345600 |
| N | 0.79709800 | -1.28821300 | -0.02747500 |
| O | -0.91975100 | -2.27002700 | -1.08218400 |
| O | -0.79413400 | 1.94652800 | 1.23902300 |
| C | 1.76957400 | -2.06387700 | -0.82035600 |
| C | 1.46428800 | -0.36354100 | 0.88103200 |
| C | 3.13171200 | -1.58254700 | -0.30879900 |
| H | 1.60489100 | -3.12703600 | -0.63226900 |
| H | 1.61796200 | -1.87762800 | -1.88434700 |
| C | 2.82429900 | -1.04345500 | 1.09153600 |
| C | 1.67303100 | 1.02960100 | 0.28885100 |
| H | 0.91742500 | -0.22802300 | 1.81326200 |
| H | 3.87052800 | -2.38212800 | -0.30381800 |
| H | 2.70006900 | -1.86037800 | 1.80460400 |
| H | 3.57044400 | -0.34866500 | 1.47245800 |
| O | 2.31248600 | 1.88039000 | 0.85385800 |
| C | 1.22137500 | 2.49579400 | -1.47447600 |
| H | 0.64118800 | 2.46878600 | -2.39255500 |
| H | 0.81270700 | 3.22779100 | -0.77768800 |
| H | 2.26438800 | 2.73124600 | -1.68507600 |
| H | -2.64160000 | 0.40546300 | -0.80276900 |
| C | -2.25413800 | 2.89245600 | -0.41468300 |
| H | -3.18774400 | 3.13611300 | 0.09776500 |
| H | -1.60771400 | 3.76728800 | -0.38539000 |
| H | -2.48448100 | 2.63281700 | -1.44791100 |
| H | 3.50863300 | -0.77647400 | -0.94343000 |
| H | -2.89195700 | -2.23855800 | -0.91996200 |
| H | -3.42047000 | -1.05233400 | 1.56880300 |
| O | 1.11640200 | 1.18287300 | -0.90716300 |

1 2 1.0 34 1.0
2 3 1.0 4 1.0 35 1.0
3 5 1.0 6 1.0 7 1.0
4
5 8 1.5 28 1.0
6 9 1.5 10 2.0
7
8 11 2.0 29 1.0
9 12 1.0 13 1.0
10
11
12 14 1.0 15 1.0 16 1.0
13 17 1.0 18 1.0 19 1.0
14 17 1.0 20 1.0 33 1.0
15

16
17 21 1.0 22 1.0
18 23 2.0 36 1.5
19
20
21
22
23
24 25 1.0 26 1.0 27 1.0 36 1.0
25
26
27
28
29 30 1.0 31 1.0 32 1.0
30
31
32
33
34
35
36

**Geometry-optimized structure of Ac-Ser-*cis*-Pro-OMe (PcisP) with a $g^+$ Ser $\chi_1$ rotamer and a *t* Ser $\chi_2$ rotamer containing an intraresidue Ser $O_\gamma$/H–N hydrogen bond**
optimized M06-2X/6-311++G(d,p)/H2O

0 1

| | | | |
|---|---|---|---|
| O | -2.32981200 | -2.11537200 | 1.62761400 |
| C | -1.40245900 | -1.09362400 | 1.95575100 |
| C | -1.28843100 | -0.17784100 | 0.73779000 |
| H | -0.42805700 | -1.52248700 | 2.20506100 |
| N | -2.60889700 | 0.29257200 | 0.39978200 |
| C | -0.65547500 | -0.92598100 | -0.45231000 |
| H | -0.67186200 | 0.68481000 | 1.00101400 |
| C | -2.78500000 | 1.31531500 | -0.47261700 |
| N | 0.68480400 | -1.09393900 | -0.42738700 |
| O | -1.34613000 | -1.38705900 | -1.34919700 |
| O | -1.84528100 | 1.99267000 | -0.86934600 |
| C | 1.35053400 | -1.87378600 | -1.48470800 |
| C | 1.63468300 | -0.49196900 | 0.49570900 |
| C | 2.84270400 | -1.69662900 | -1.18979200 |
| H | 1.03368000 | -2.91723400 | -1.41108800 |
| H | 1.06098900 | -1.49665300 | -2.46605800 |
| C | 2.88413100 | -1.37266400 | 0.30663200 |
| C | 1.96501000 | 0.94424300 | 0.10393200 |
| H | 1.29261800 | -0.50370300 | 1.52910900 |

| | | | |
|---|---|---|---|
| H | 3.41696300 | -2.58737400 | -1.43961500 |
| H | 2.75651000 | -2.27797400 | 0.90327900 |
| H | 3.79693500 | -0.87005700 | 0.62494300 |
| O | 1.80637200 | 1.41409300 | -0.99313100 |
| C | 2.94666100 | 2.93848000 | 0.85957300 |
| H | 3.34904800 | 3.30923100 | 1.79739200 |
| H | 3.71601000 | 2.92891900 | 0.08803400 |
| H | 2.10471200 | 3.54869700 | 0.53476800 |
| H | -3.36817800 | -0.35651900 | 0.55488200 |
| C | -4.20329900 | 1.56234500 | -0.92186300 |
| H | -4.36881400 | 2.63563000 | -1.00106800 |
| H | -4.32725500 | 1.12348400 | -1.91456400 |
| H | -4.93796000 | 1.12196000 | -0.24892100 |
| H | 3.24118400 | -0.85939600 | -1.76552200 |
| H | -2.50532000 | -2.63857400 | 2.41475300 |
| H | -1.75731900 | -0.50080100 | 2.80373900 |
| O | 2.50615200 | 1.59820100 | 1.12837700 |

1 2 1.0 34 1.0
2 3 1.0 4 1.0 35 1.0
3 5 1.0 6 1.0 7 1.0
4
5 8 1.5 28 1.0
6 9 1.5 10 2.0
7
8 11 2.0 29 1.0
9 12 1.0 13 1.0
10
11
12 14 1.0 15 1.0 16 1.0
13 17 1.0 18 1.0 19 1.0
14 17 1.0 20 1.0 33 1.0
15
16
17 21 1.0 22 1.0
18 23 2.0 36 1.5
19
20
21
22
23
24 25 1.0 26 1.0 27 1.0 36 1.0
25
26
27
28
29 30 1.0 31 1.0 32 1.0

30
31
32
33
34
35
36

**Geometry-optimized structure of Ac-Ser-*cis*-Pro-OMe (BcisD) with a $g^+$ Ser $\chi_1$ rotamer and a $g^-$ Ser $\chi_2$ rotamer containing an intraresidue Ser $O_\gamma$–H/N hydrogen bond**
optimized M06-2X/6-311++G(d,p)/H2O

0 1

| | | | |
|---|---|---|---|
| O | -3.50318600 | -1.79877700 | -0.31113000 |
| C | -2.76096000 | -1.57162600 | 0.87249000 |
| C | -1.53324200 | -0.69264400 | 0.61366000 |
| H | -2.44004700 | -2.51435600 | 1.32785900 |
| N | -1.94332900 | 0.51467100 | -0.07937000 |
| C | -0.51517900 | -1.47133100 | -0.23117200 |
| H | -1.10615500 | -0.39357400 | 1.56992100 |
| C | -1.58725800 | 1.75726200 | 0.32345600 |
| N | 0.79709800 | -1.28821300 | -0.02747500 |
| O | -0.91975100 | -2.27002700 | -1.08218400 |
| O | -0.79413400 | 1.94652800 | 1.23902300 |
| C | 1.76957400 | -2.06387700 | -0.82035600 |
| C | 1.46428800 | -0.36354100 | 0.88103200 |
| C | 3.13171200 | -1.58254700 | -0.30879900 |
| H | 1.60489100 | -3.12703600 | -0.63226900 |
| H | 1.61796200 | -1.87762800 | -1.88434700 |
| C | 2.82429900 | -1.04345500 | 1.09153600 |
| C | 1.67303100 | 1.02960100 | 0.28885100 |
| H | 0.91742500 | -0.22802300 | 1.81326200 |
| H | 3.87052800 | -2.38212800 | -0.30381800 |
| H | 2.70006900 | -1.86037800 | 1.80460400 |
| H | 3.57044400 | -0.34866500 | 1.47245800 |
| O | 2.31248600 | 1.88039000 | 0.85385800 |
| C | 1.22137500 | 2.49579400 | -1.47447600 |
| H | 0.64118800 | 2.46878600 | -2.39255500 |
| H | 0.81270700 | 3.22779100 | -0.77768800 |
| H | 2.26438800 | 2.73124600 | -1.68507600 |
| H | -2.64160000 | 0.40546300 | -0.80276900 |
| C | -2.25413800 | 2.89245600 | -0.41468300 |
| H | -3.18774400 | 3.13611300 | 0.09776500 |
| H | -1.60771400 | 3.76728800 | -0.38539000 |
| H | -2.48448100 | 2.63281700 | -1.44791100 |
| H | 3.50863300 | -0.77647400 | -0.94343000 |

H -2.89195700 -2.23855800 -0.91996200
H -3.42047000 -1.05233400 1.56880300
O 1.11640200 1.18287300 -0.90716300

1 2 1.0 34 1.0
2 3 1.0 4 1.0 35 1.0
3 5 1.0 6 1.0 7 1.0
4
5 8 1.5 28 1.0
6 9 1.5 10 2.0
7
8 11 2.0 29 1.0
9 12 1.0 13 1.0
10
11
12 14 1.0 15 1.0 16 1.0
13 17 1.0 18 1.0 19 1.0
14 17 1.0 20 1.0 33 1.0
15
16
17 21 1.0 22 1.0
18 23 2.0 36 1.5
19
20
21
22
23
24 25 1.0 26 1.0 27 1.0 36 1.0
25
26
27
28
29 30 1.0 31 1.0 32 1.0
30
31
32
33
34
35
36

**Geometry-optimized structure of Ac-Ser-*trans*-Pro-OMe (BtransP) with a $g^{-}$ Ser $\chi_1$ rotamer and a $g^{+}$ Ser $\chi_2$ rotamer containing a Ser $O_\gamma$–H/O=$C_{i-1}$ hydrogen bond**

optimized M06-2X/6-311++G(d,p)/H2O

0 1
O -2.01094900 1.48762500 1.93739200

| | | | |
|---|---|---|---|
| C | -1.34918700 | 0.35641800 | 1.41796200 |
| C | -1.44790800 | 0.28061900 | -0.13046500 |
| H | -1.75615100 | -0.57583300 | 1.82528900 |
| N | -2.51874000 | -0.58116800 | -0.58958300 |
| C | -0.14429600 | -0.26585600 | -0.69802700 |
| H | -1.61740300 | 1.29584800 | -0.50274800 |
| C | -3.80819300 | -0.32844100 | -0.30768000 |
| N | 0.88815000 | 0.59020100 | -0.74754000 |
| O | -0.04274300 | -1.44070800 | -1.04138000 |
| O | -4.14327500 | 0.64321400 | 0.37549100 |
| C | 0.89878200 | 2.00858200 | -0.34757600 |
| C | 2.20573400 | 0.09774500 | -1.12554400 |
| C | 2.38421600 | 2.27803400 | -0.10930000 |
| H | 0.50673100 | 2.62444000 | -1.16238400 |
| H | 0.29270500 | 2.17683300 | 0.54225000 |
| C | 3.07201000 | 1.37179600 | -1.13477500 |
| C | 2.74355100 | -0.93319700 | -0.14161700 |
| H | 2.17886800 | -0.39317200 | -2.09875600 |
| H | 2.63442300 | 3.32960100 | -0.24092300 |
| H | 3.02202600 | 1.81804000 | -2.12965200 |
| H | 4.11580900 | 1.16063400 | -0.90448900 |
| O | 3.57553900 | -1.75222900 | -0.44180000 |
| C | 2.72297900 | -1.71798900 | 2.06921000 |
| H | 2.19456900 | -1.46953900 | 2.98477100 |
| H | 2.50066600 | -2.73978200 | 1.76314600 |
| H | 3.79791100 | -1.59736300 | 2.20048700 |
| H | -2.24589900 | -1.42048200 | -1.08480900 |
| C | -4.82872900 | -1.27849900 | -0.87565200 |
| H | -5.45785900 | -1.64008800 | -0.06221100 |
| H | -5.46224600 | -0.72389300 | -1.56961100 |
| H | -4.37853100 | -2.12238400 | -1.39535100 |
| H | 2.65503700 | 1.97861800 | 0.90519100 |
| H | -2.91100800 | 1.45380200 | 1.57548800 |
| H | -0.30400700 | 0.42752100 | 1.72654100 |
| O | 2.24076900 | -0.79556600 | 1.08134800 |

1 2 1.0 34 1.0
2 3 1.0 4 1.0 35 1.0
3 5 1.0 6 1.0 7 1.0
4
5 8 1.5 28 1.0
6 9 1.5 10 2.0
7
8 11 2.0 29 1.0
9 12 1.0 13 1.0
10
11
12 14 1.0 15 1.0 16 1.0
13 17 1.0 18 1.0 19 1.0
14 17 1.0 20 1.0 33 1.0
15

16
17 21 1.0 22 1.0
18 23 2.0 36 1.5
19
20
21
22
23
24 25 1.0 26 1.0 27 1.0 36 1.0
25
26
27
28
29 30 1.0 31 1.0 32 1.0
30
31
32
33
34
35
36

**Geometry-optimized structure of Ac-Ser-*trans*-Pro-OMe (PtransP) with a $g^+$ Ser $\chi_1$ rotamer and a $g^+$ Ser $\chi_2$ rotamer containing intraresidue Ser $O_\gamma$–H/O=C and Ser $O_\gamma$–H/N hydrogen bonds**
optimized M06-2X/6-311++G(d,p)/H2O

0 1

| | | | |
|---|---|---|---|
| O | -2.67386400 | 1.60494500 | -0.58576200 |
| O | -0.08338600 | -1.10815500 | -0.79707900 |
| O | 2.56611600 | -0.97424500 | 0.86464500 |
| O | -1.54073600 | -2.89903100 | 0.79697100 |
| N | -2.82521600 | -0.55022800 | 0.03241800 |
| H | -3.18748900 | -1.47182900 | -0.16973400 |
| N | 0.40611700 | 0.77661700 | 0.32482300 |
| C | -3.22755900 | 0.51584800 | -0.69696900 |
| C | -1.65828500 | -0.46254600 | 0.88457300 |
| H | -1.78863300 | 0.35066700 | 1.59861000 |
| C | -1.50549800 | -1.77796000 | 1.65854500 |
| H | -0.57353600 | -1.74261100 | 2.23344200 |
| C | -0.39058500 | -0.26295100 | 0.05085000 |
| C | 0.11541000 | 1.94785700 | 1.17086900 |
| H | 0.32170700 | 1.71550200 | 2.21969900 |
| H | -0.92545300 | 2.24308700 | 1.05090400 |
| C | 1.09294000 | 2.99053800 | 0.62994800 |
| H | 1.32464200 | 3.75886800 | 1.36613600 |
| H | 0.66579800 | 3.46972900 | -0.25412800 |
| C | 2.31292500 | 2.15392700 | 0.23680700 |
| H | 2.87798300 | 1.87627300 | 1.13069300 |
| H | 2.98244100 | 2.65234600 | -0.46185800 |
| C | 1.67778600 | 0.89558200 | -0.38061700 |

| | | | |
|---|---|---|---|
| H | 1.50928600 | 1.01311400 | -1.45377700 |
| C | 2.54050800 | -0.33249900 | -0.15399900 |
| C | -4.37830800 | 0.29074900 | -1.64392400 |
| H | -5.13076200 | 1.05961300 | -1.46880600 |
| H | -4.00875600 | 0.40716100 | -2.66413800 |
| H | -4.82982100 | -0.69419400 | -1.53566000 |
| C | 4.23432700 | -1.66634900 | -1.07751200 |
| H | 3.68725800 | -2.59110000 | -0.89695600 |
| H | 4.76854700 | -1.71591800 | -2.02155500 |
| H | 4.92486500 | -1.48036400 | -0.25546600 |
| H | -2.33960000 | -1.88265700 | 2.35359900 |
| H | -0.90595500 | -2.72740000 | 0.08506200 |
| O | 3.31677000 | -0.57078900 | -1.20729700 |

1 8 2.0
2 13 2.0
3 25 2.0
4 11 1.0 35 1.0
5 6 1.0 8 1.5 9 1.0
6
7 13 1.5 14 1.0 23 1.0
8 26 1.0
9 10 1.0 11 1.0 13 1.0
10
11 12 1.0 34 1.0
12
13
14 15 1.0 16 1.0 17 1.0
15
16
17 18 1.0 19 1.0 20 1.0
18
19
20 21 1.0 22 1.0 23 1.0
21
22
23 24 1.0 25 1.0
24
25 36 1.5
26 27 1.0 28 1.0 29 1.0
27
28
29
30 31 1.0 32 1.0 33 1.0 36 1.0
31
32
33
34
35
36

**Geometry-optimized structure of Ac-Ser-*trans*-Pro-OMe (PtransP) with a $g^+$ Ser $\chi_1$ rotamer and a *t* Ser $\chi_2$ rotamer containing an intraresidue Ser $O_\gamma$/H–N hydrogen bond**
optimized M06-2X/6-311++G(d,p)/H2O

0 1

| | | | |
|---|---|---|---|
| O | 3.03626800 | 0.64332500 | 1.73139700 |
| O | 0.37108800 | -1.04225400 | 0.58251000 |
| O | -2.31128800 | -0.96087800 | -1.13339000 |
| O | 1.34734400 | -0.91805400 | -2.51852600 |
| N | 2.84905600 | -0.31818600 | -0.29840600 |
| H | 2.98585500 | -1.16196100 | -0.83793300 |
| N | -0.57971500 | 0.92732000 | 0.07614100 |
| C | 3.37221000 | -0.23842900 | 0.94997800 |
| C | 1.73144700 | 0.51298100 | -0.67074600 |
| H | 1.96309300 | 1.54233700 | -0.39334800 |
| C | 1.53973200 | 0.44644400 | -2.18107700 |
| H | 0.67708800 | 1.04998400 | -2.47568500 |
| C | 0.45669600 | 0.06287200 | 0.06156700 |
| C | -0.63072500 | 2.32300400 | -0.38380800 |
| H | -0.87673900 | 2.36761500 | -1.44923300 |
| H | 0.31840900 | 2.82978800 | -0.21627700 |
| C | -1.76781700 | 2.90093300 | 0.45660900 |
| H | -2.20722000 | 3.78831300 | 0.00323600 |
| H | -1.39701100 | 3.15983400 | 1.45087500 |
| C | -2.74904800 | 1.72962000 | 0.54451200 |
| H | -3.30072300 | 1.63919600 | -0.39511800 |
| H | -3.46217900 | 1.81315900 | 1.36250500 |
| C | -1.82174200 | 0.51228400 | 0.71813100 |
| H | -1.64573700 | 0.28227700 | 1.77137700 |
| C | -2.40030800 | -0.71746300 | 0.04265400 |
| C | 4.39161600 | -1.28811400 | 1.31370100 |
| H | 5.23632200 | -0.80414600 | 1.80282300 |
| H | 3.93420900 | -1.97483500 | 2.02890700 |
| H | 4.74011500 | -1.85408900 | 0.45100900 |
| C | -3.74832200 | -2.61013500 | 0.36410300 |
| H | -3.01927100 | -3.28696900 | -0.08025600 |
| H | -4.24943500 | -3.08537400 | 1.20199400 |
| H | -4.47135200 | -2.30015800 | -0.38986900 |
| H | 2.43510400 | 0.84721900 | -2.66500400 |
| H | 1.32807300 | -1.00339100 | -3.47586300 |
| O | -3.08296600 | -1.46172800 | 0.90900500 |

1 8 2.0
2 13 2.0
3 25 2.0
4 11 1.0 35 1.0
5 6 1.0 8 1.5 9 1.0
6
7 13 1.5 14 1.0 23 1.0
8 26 1.0

9 10 1.0 11 1.0 13 1.0
10
11 12 1.0 34 1.0
12
13
14 15 1.0 16 1.0 17 1.0
15
16
17 18 1.0 19 1.0 20 1.0
18
19
20 21 1.0 22 1.0 23 1.0
21
22
23 24 1.0 25 1.0
24
25 36 1.5
26 27 1.0 28 1.0 29 1.0
27
28
29
30 31 1.0 32 1.0 33 1.0 36 1.0
31
32
33
34
35
36

**Geometry-optimized structure of Ac-Ser-*trans*-Pro-OMe (BtransP) with a *t* Ser $\chi_1$ rotamer and a $g^{-}$ Ser $\chi_2$ rotamer containing an intraresidue Ser $O_\gamma$/O=$C_{i+1}$ hydrogen bond**
optimized M06-2X/6-311++G(d,p)/H2O

0 1

| | | | |
|---|---|---|---|
| O | -0.35974800 | 0.22129600 | 2.30189500 |
| C | -1.44594400 | -0.38250400 | 1.63528600 |
| C | -1.65890200 | 0.14990000 | 0.19627300 |
| H | -2.34909300 | -0.16475500 | 2.20658300 |
| N | -2.75521700 | -0.55741600 | -0.42574600 |
| C | -0.39025100 | -0.14516000 | -0.58607500 |
| H | -1.89525600 | 1.21315400 | 0.22105600 |
| C | -4.04328100 | -0.18250900 | -0.25626600 |
| N | 0.57810000 | 0.78707500 | -0.58309000 |
| O | -0.23268100 | -1.25109200 | -1.09955600 |
| O | -4.35373800 | 0.79109400 | 0.42188400 |
| C | 0.52534100 | 2.16218200 | -0.04509300 |
| C | 1.91284500 | 0.41050900 | -1.02318100 |
| C | 2.00165000 | 2.49854400 | 0.16041400 |
| H | 0.06333000 | 2.82446000 | -0.78231900 |
| H | -0.03930200 | 2.19572700 | 0.88311000 |

| | | | |
|---|---|---|---|
| C | 2.69815600 | 1.73282300 | -0.96770600 |
| C | 2.49237400 | -0.61087100 | -0.05091300 |
| H | 1.90242800 | -0.03315500 | -2.01867200 |
| H | 2.18647600 | 3.57085000 | 0.12196000 |
| H | 2.57193100 | 2.25217200 | -1.91929700 |
| H | 3.76258800 | 1.57290400 | -0.80007600 |
| O | 2.06726000 | -0.82623400 | 1.06135500 |
| C | 4.23554200 | -2.14516400 | 0.29186200 |
| H | 5.08282700 | -2.50271100 | -0.28500200 |
| H | 4.57041200 | -1.64538100 | 1.19980900 |
| H | 3.56687800 | -2.96683900 | 0.54491300 |
| H | -2.51981100 | -1.38118800 | -0.96277900 |
| C | -5.07826000 | -1.02712400 | -0.95782800 |
| H | -5.76171700 | -1.42951400 | -0.20914700 |
| H | -5.65319800 | -0.38408600 | -1.62526000 |
| H | -4.64482000 | -1.84650200 | -1.52918200 |
| H | 2.33254700 | 2.12410900 | 1.13251400 |
| H | 0.46942400 | -0.12171600 | 1.93225300 |
| H | -1.32364600 | -1.46997100 | 1.57254300 |
| O | 3.55668100 | -1.20743100 | -0.56080100 |

1 2 1.0 34 1.0
2 3 1.0 4 1.0 35 1.0
3 5 1.0 6 1.0 7 1.0
4
5 8 1.5 28 1.0
6 9 1.5 10 2.0
7
8 11 2.0 29 1.0
9 12 1.0 13 1.0
10
11
12 14 1.0 15 1.0 16 1.0
13 17 1.0 18 1.0 19 1.0
14 17 1.0 20 1.0 33 1.0
15
16
17 21 1.0 22 1.0
18 23 2.0 36 1.5
19
20
21
22
23
24 25 1.0 26 1.0 27 1.0 36 1.0
25
26
27
28
29 30 1.0 31 1.0 32 1.0
30

### Geometry-optimized structures of Ac-Ser-*trans*-Pro-Ala-NHMe type I β-turns

#### Geometry-optimized structure of an Ac-Ser-*trans*-Pro-Ala-NHMe type I β-turn with Ser in the PPII conformation and a *trans* $\chi_1$ rotamer

optimized M06-2X/6-311++G(d,p)/H2O

0 1

| | | | |
|---|---|---|---|
| N | 1.80360000 | 0.81010000 | 0.29020000 |
| C | 3.12400000 | 0.23880000 | 0.47690000 |
| C | 3.32850000 | -1.09110000 | -0.25990000 |
| O | 4.46380000 | -1.54760000 | -0.37460000 |
| C | 3.39650000 | 0.02970000 | 1.96430000 |
| H | 3.86070000 | 0.92630000 | 0.05630000 |
| H | 3.30850000 | 0.97610000 | 2.49930000 |
| H | 4.40260000 | -0.36400000 | 2.10210000 |
| N | 2.23570000 | -1.72640000 | -0.70250000 |
| C | 2.34160000 | -3.00230000 | -1.38710000 |
| H | 2.84560000 | -3.73440000 | -0.75430000 |
| H | 1.32620000 | -1.27930000 | -0.65230000 |
| C | -4.68270000 | -2.35640000 | -0.97940000 |
| C | -3.84650000 | -1.25360000 | -0.38460000 |
| O | -4.02200000 | -0.07230000 | -0.65980000 |
| H | -4.46740000 | -3.33220000 | -0.54700000 |
| N | -2.87530000 | -1.62530000 | 0.48510000 |
| C | -1.98340000 | -0.62150000 | 1.01610000 |
| C | -1.12770000 | -0.00710000 | -0.09620000 |
| O | -0.55980000 | -0.74280000 | -0.90570000 |
| C | -1.02770000 | -1.25670000 | 2.02240000 |
| H | -2.57020000 | 0.14290000 | 1.52370000 |
| H | -0.48330000 | -2.07800000 | 1.54230000 |
| H | -2.64970000 | -2.60310000 | 0.59470000 |
| N | -0.92110000 | 1.31720000 | -0.07020000 |
| C | 0.03700000 | 1.92510000 | -0.99540000 |
| C | 1.44760000 | 1.35980000 | -0.88960000 |
| O | 2.21930000 | 1.46640000 | -1.83430000 |
| C | 0.00390000 | 3.41490000 | -0.61170000 |
| H | 0.72420000 | 3.60120000 | 0.18990000 |
| H | -0.28400000 | 1.76950000 | -2.02870000 |
| H | -1.60060000 | -1.64570000 | 2.86740000 |
| H | 0.25300000 | 4.05170000 | -1.45840000 |
| H | 1.11060000 | 0.67470000 | 1.02230000 |
| H | 2.68370000 | -0.68370000 | 2.38510000 |
| H | -5.73460000 | -2.11470000 | -0.82790000 |
| H | -4.49520000 | -2.38840000 | -2.05380000 |
| H | 2.90590000 | -2.90280000 | -2.31700000 |
| H | 1.33760000 | -3.35520000 | -1.61420000 |

| | | | |
|---|---|---|---|
| C | -1.68310000 | 2.32300000 | 0.69540000 |
| C | -1.42110000 | 3.60690000 | -0.09100000 |
| H | -1.53170000 | 4.49670000 | 0.52720000 |
| H | -2.12000000 | 3.67370000 | -0.92800000 |
| H | -1.29770000 | 2.38640000 | 1.71670000 |
| H | -2.73880000 | 2.05700000 | 0.71680000 |
| O | -0.12980000 | -0.23520000 | 2.43540000 |
| H | 0.30200000 | -0.50790000 | 3.25050000 |

1 2 1.0 27 1.5 34 1.0
2 3 1.0 5 1.0 6 1.0
3 4 2.0 9 1.5
4
5 7 1.0 8 1.0 35 1.0
6
7
8
9 10 1.0 12 1.0
10 11 1.0 38 1.0 39 1.0
11
12
13 14 1.0 16 1.0 36 1.0 37 1.0
14 15 2.0 17 1.5
15
16
17 18 1.0 24 1.0
18 19 1.0 21 1.0 22 1.0
19 20 2.0 25 1.5
20
21 23 1.0 32 1.0 46 1.0
22
23
24
25 26 1.0 40 1.0
26 27 1.0 29 1.0 31 1.0
27 28 2.0
28
29 30 1.0 33 1.0 41 1.0
40 41 1.0 44 1.0 45 1.0
41 42 1.0 43 1.0
42
43
44
45
46 47 1.0
47

**Geometry-optimized structure of an Ac-Ser-*trans*-Pro-Ala-NHMe type I β-turn with Ser in the β conformation and a *trans* $\chi_1$ rotamer**
optimized M06-2X/6-311++G(d,p)/H2O

0 1

| | | | |
|---|---|---|---|
| N | 1.88050000 | 0.47150000 | 0.46170000 |
| C | 2.93560000 | -0.43120000 | 0.88020000 |
| C | 2.98820000 | -1.73080000 | 0.06700000 |
| O | 3.98160000 | -2.44980000 | 0.14560000 |
| C | 2.76120000 | -0.77700000 | 2.35700000 |
| H | 3.90090000 | 0.05470000 | 0.72110000 |
| H | 2.77970000 | 0.13150000 | 2.96080000 |
| H | 3.56850000 | -1.43310000 | 2.67950000 |
| N | 1.90850000 | -2.05120000 | -0.65850000 |
| C | 1.87380000 | -3.27390000 | -1.44100000 |
| H | 2.63550000 | -3.26000000 | -2.22390000 |
| H | 1.14500000 | -1.39170000 | -0.75870000 |
| C | -4.97140000 | -2.14710000 | -0.96020000 |
| C | -4.24430000 | -1.14170000 | -0.10260000 |
| O | -4.79910000 | -0.55070000 | 0.81650000 |
| H | -4.33180000 | -2.61140000 | -1.70900000 |
| N | -2.94500000 | -0.92250000 | -0.41390000 |
| C | -2.12890000 | 0.01290000 | 0.32450000 |
| C | -0.97540000 | 0.45580000 | -0.56390000 |
| O | -0.46600000 | -0.36030000 | -1.33830000 |
| C | -1.52260000 | -0.67900000 | 1.55560000 |
| H | -2.75020000 | 0.84550000 | 0.65060000 |
| H | -2.49140000 | -1.44240000 | -1.15290000 |
| N | -0.46820000 | 1.68210000 | -0.38590000 |
| C | 0.78380000 | 2.05240000 | -1.05370000 |
| C | 1.94840000 | 1.12730000 | -0.71590000 |
| O | 2.90400000 | 1.04020000 | -1.47500000 |
| C | 1.04610000 | 3.48630000 | -0.55860000 |
| H | 1.64580000 | 3.45580000 | 0.35540000 |
| H | 0.65740000 | 2.01570000 | -2.13810000 |
| H | -0.97740000 | -1.57090000 | 1.22510000 |
| H | -2.33830000 | -0.97830000 | 2.21720000 |

| | | | |
|---|---|---|---|
| H | 1.57990000 | 4.07370000 | -1.30320000 |
| H | 1.05870000 | 0.56550000 | 1.05420000 |
| H | 1.80860000 | -1.28830000 | 2.51800000 |
| H | -5.39000000 | -2.91640000 | -0.31070000 |
| H | -5.79940000 | -1.63980000 | -1.45730000 |
| H | 0.89070000 | -3.36220000 | -1.89910000 |
| H | 2.05180000 | -4.13980000 | -0.80180000 |
| C | -1.04660000 | 2.80410000 | 0.38020000 |
| C | -0.35500000 | 4.01380000 | -0.24460000 |
| H | -0.34490000 | 4.87150000 | 0.42610000 |
| H | -0.86800000 | 4.29580000 | -1.16670000 |
| H | -0.80090000 | 2.69470000 | 1.43900000 |
| H | -2.12810000 | 2.83350000 | 0.26240000 |
| O | -0.64420000 | 0.23890000 | 2.19390000 |
| H | -0.50000000 | -0.03970000 | 3.10320000 |

1 2 1.0 26 1.5 34 1.0
2 3 1.0 5 1.0 6 1.0
3 4 2.0 9 1.5
4
5 7 1.0 8 1.0 35 1.0
6
7
8
9 10 1.0 12 1.0
10 11 1.0 38 1.0 39 1.0
11
12
13 14 1.0 16 1.0 36 1.0 37 1.0
14 15 2.0 17 1.5
15
16
17 18 1.0 23 1.0
18 19 1.0 21 1.0 22 1.0
19 20 2.0 24 1.5
20
21 31 1.0 32 1.0 46 1.0
22
23
24 25 1.0 40 1.0
25 26 1.0 28 1.0 30 1.0
26 27 2.0
27
28 29 1.0 33 1.0 41 1.0
40 41 1.0 44 1.0 45 1.0
41 42 1.0 43 1.0
42
43
44
45
46 47 1.0
47

**Geometry-optimized structure of an Ac-Ser-*trans*-Pro-Ala-NHMe type I β-turn with Ser in the PPII conformation and a *trans* $g^+$ rotamer**
optimized M06-2X/6-311++G(d,p)/H2O

0 1

| | | | |
|---|---|---|---|
| N | 1.86370000 | 0.36730000 | 0.56440000 |
| C | 2.84350000 | -0.61290000 | 0.99260000 |
| C | 2.87750000 | -1.86500000 | 0.10690000 |
| O | 3.82990000 | -2.63690000 | 0.18840000 |
| C | 2.56790000 | -1.02950000 | 2.43490000 |
| H | 3.83770000 | -0.16830000 | 0.91420000 |
| H | 2.58620000 | -0.15630000 | 3.08850000 |
| H | 3.32790000 | -1.73600000 | 2.76600000 |
| N | 1.82230000 | -2.08510000 | -0.68840000 |
| C | 1.76500000 | -3.25790000 | -1.54270000 |
| H | 2.60630000 | -3.27180000 | -2.23860000 |
| H | 1.08500000 | -1.39300000 | -0.76080000 |
| C | -4.54320000 | -2.55510000 | -0.78720000 |
| C | -4.11830000 | -1.23930000 | -0.18460000 |
| O | -4.90420000 | -0.51690000 | 0.42060000 |
| H | -3.72900000 | -3.07530000 | -1.28960000 |
| N | -2.82010000 | -0.90140000 | -0.34440000 |
| C | -2.30320000 | 0.35700000 | 0.14200000 |
| C | -0.98350000 | 0.62010000 | -0.58560000 |
| O | -0.48540000 | -0.24660000 | -1.30520000 |
| C | -2.06550000 | 0.33360000 | 1.65660000 |
| H | -3.02760000 | 1.14590000 | -0.08060000 |
| H | -2.19210000 | -1.47340000 | -0.89240000 |
| N | -0.35400000 | 1.78130000 | -0.34570000 |
| C | 0.94310000 | 2.04000000 | -0.97950000 |

| | | | |
|---|---|---|---|
| C | 2.04650000 | 1.07320000 | -0.57020000 |
| O | 3.06400000 | 0.99800000 | -1.24680000 |
| C | 1.27280000 | 3.48300000 | -0.55690000 |
| H | 1.83760000 | 3.47260000 | 0.37960000 |
| H | 0.84540000 | 1.95450000 | -2.06470000 |
| H | -2.89460000 | -0.20170000 | 2.12290000 |
| H | -2.03950000 | 1.35010000 | 2.05490000 |
| H | 1.86490000 | 3.99380000 | -1.31350000 |
| H | 0.96750000 | 0.39140000 | 1.04390000 |
| H | 1.58950000 | -1.51010000 | 2.51200000 |
| H | -4.94230000 | -3.18760000 | 0.00650000 |
| H | -5.34610000 | -2.36470000 | -1.50060000 |
| H | 0.83480000 | -3.22930000 | -2.10620000 |
| H | 1.79540000 | -4.17110000 | -0.94580000 |
| C | -0.87060000 | 2.97220000 | 0.35250000 |
| C | -0.10600000 | 4.10490000 | -0.32740000 |
| H | -0.07170000 | 5.00310000 | 0.28730000 |
| H | -0.57770000 | 4.34770000 | -1.28230000 |
| H | -0.61730000 | 2.92220000 | 1.41560000 |
| H | -1.95110000 | 3.05280000 | 0.24200000 |
| O | -0.82570000 | -0.32740000 | 1.89340000 |
| H | -0.75890000 | -0.53940000 | 2.82950000 |

1 2 1.0 26 1.5 34 1.0
2 3 1.0 5 1.0 6 1.0
3 4 2.0 9 1.5
4
5 7 1.0 8 1.0 35 1.0
6
7
8
9 10 1.0 12 1.0
10 11 1.0 38 1.0 39 1.0
11
12
13 14 1.0 16 1.0 36 1.0 37 1.0
14 15 2.0 17 1.5
15
16
17 18 1.0 23 1.0
18 19 1.0 21 1.0 22 1.0
19 20 2.0 24 1.5
20
21 31 1.0 32 1.0 46 1.0
22
23
24 25 1.0 40 1.0

25 26 1.0 28 1.0 30 1.0
26 27 2.0
27
28 29 1.0 33 1.0 41 1.0
40 41 1.0 44 1.0 45 1.0
41 42 1.0 43 1.0
42
43
44
45
46 47 1.0
47

### Geometry-optimized structures of Ac-Ala$_5$-NHMe, Ac-Pro-Ala$_4$-NHMe, and Pro-Ala$_4$-NHMe with an Ac-Ser Ncap in $3_{10}$-helices

#### Geometry-optimized structure of Ac-Ala$_5$-NHMe in a $3_{10}$-helix with two $H_2O$ molecules at the N-terminus

optimized M06-2X/6-311++G(d,p)/H2O

0 1

| | | | |
|---|---|---|---|
| C | 4.39153000 | 1.77299100 | -1.13312700 |
| H | 5.30390200 | 1.22159300 | -0.91137700 |
| C | 3.18317900 | 0.88071000 | -1.23405300 |
| O | 2.05032200 | 1.33930200 | -1.39902200 |
| N | 3.39832100 | -0.44455900 | -1.12993400 |
| H | 4.33560300 | -0.78086400 | -0.93115400 |
| C | 2.31016200 | -1.39852500 | -1.22673100 |
| H | 1.81822100 | -1.29304400 | -2.19715300 |
| C | 2.85146300 | -2.81644100 | -1.06318500 |
| C | 1.21139900 | -1.14529400 | -0.19581300 |
| H | 3.31784900 | -2.93730800 | -0.08212900 |
| H | 2.03961200 | -3.53731500 | -1.15599800 |
| H | 3.59362000 | -3.02376400 | -1.83564300 |
| O | 0.05124500 | -1.47806300 | -0.44073100 |
| N | 1.57165200 | -0.57600800 | 0.96466400 |
| H | 2.54515500 | -0.32253700 | 1.14394900 |
| C | 0.58329100 | -0.30815500 | 1.99138300 |
| H | 0.06675900 | -1.23604500 | 2.25167600 |
| C | 1.25723600 | 0.27836600 | 3.22895800 |
| C | -0.52165000 | 0.63532100 | 1.51553900 |
| H | 1.75377600 | 1.22042300 | 2.98494200 |
| H | 0.50954100 | 0.46120300 | 4.00024400 |
| H | 2.00222300 | -0.41887700 | 3.61351400 |
| O | -1.61084800 | 0.63249200 | 2.08925000 |
| N | -0.25349700 | 1.45391000 | 0.48456400 |
| H | 0.62841700 | 1.38352100 | -0.02146200 |
| C | -1.27691000 | 2.34604600 | -0.03089200 |
| H | -1.66507400 | 2.95572000 | 0.78852800 |
| C | -0.68931200 | 3.24188000 | -1.11733100 |
| C | -2.49585200 | 1.59917100 | -0.58424100 |
| H | -0.29587500 | 2.63909100 | -1.93827000 |
| H | -1.46210200 | 3.90835900 | -1.49916200 |
| H | 0.12375400 | 3.84211900 | -0.70683400 |
| O | -3.56682100 | 2.18273600 | -0.69932300 |
| N | -2.31367700 | 0.31526000 | -0.95307900 |
| H | -1.41725600 | -0.14043000 | -0.79814200 |
| C | -3.41165700 | -0.49137100 | -1.44871400 |
| H | -4.02482400 | 0.12754300 | -2.10730900 |
| C | -2.86889200 | -1.69380500 | -2.21686300 |

| | | | |
|---|---|---|---|
| C | -4.35959900 | -0.97926900 | -0.34473000 |
| H | -2.22957200 | -2.30054900 | -1.57157200 |
| H | -3.69666100 | -2.30441200 | -2.57498500 |
| H | -2.27987000 | -1.35608400 | -3.07064000 |
| O | -5.42104600 | -1.51583100 | -0.65600100 |
| N | -3.96391100 | -0.82738800 | 0.92558900 |
| H | -3.10557000 | -0.33249600 | 1.14140000 |
| C | -4.80172300 | -1.27201700 | 2.02499400 |
| H | -4.99129500 | -2.34442200 | 1.95452900 |
| H | 4.21110100 | 2.51418500 | -0.35361100 |
| H | 4.51061800 | 2.30167600 | -2.08033400 |
| O | 6.04459700 | -1.18575900 | 0.04395000 |
| H | 6.19292600 | -2.11114100 | 0.26842300 |
| H | 6.90513000 | -0.84977900 | -0.22926500 |
| O | 4.27113000 | 0.08566100 | 1.86002200 |
| H | 5.01445400 | -0.25427600 | 1.33667600 |
| H | 4.54472900 | 0.94793200 | 2.18541800 |
| H | -4.28480900 | -1.05973800 | 2.95858300 |
| H | -5.76129600 | -0.75073800 | 2.01820800 |

1 2 1.0 3 1.0 49 1.0 50 1.0
2
3 4 2.0 5 1.5
4
5 6 1.0 7 1.0
6
7 8 1.0 9 1.0 10 1.0
8
9 11 1.0 12 1.0 13 1.0
10 14 2.0 15 1.5
11
12
13
14
15 16 1.0 17 1.0
16
17 18 1.0 19 1.0 20 1.0
18
19 21 1.0 22 1.0 23 1.0
20 24 2.0 25 1.5
21
22
23
24
25 26 1.0 27 1.0
26
27 28 1.0 29 1.0 30 1.0

28
29 31 1.0 32 1.0 33 1.0
30 34 2.0 35 1.5
31
32
33
34
35 36 1.0 37 1.0
36
37 38 1.0 39 1.0 40 1.0
38
39 41 1.0 42 1.0 43 1.0
40 44 2.0 45 1.5
41
42
43
44
45 46 1.0 47 1.0
46
47 48 1.0 57 1.0 58 1.0
48
49
50
51 52 1.0 53 1.0
52
53
54 55 1.0 56 1.0
55
56
57
58

**Geometry-optimized structure of Ac-Pro-Ala$_4$-NHMe in a $3_{10}$-helix with one $H_2O$ molecule at the N-terminus**

optimized M06-2X/6-311++G(d,p)/H2O

0 1

| | | | |
|---|---|---|---|
| C | -4.40521000 | -1.25139000 | -1.74978700 |
| H | -5.07793600 | -1.41692400 | -0.90793400 |
| C | -3.22255200 | -0.41466500 | -1.33748900 |
| O | -2.06580700 | -0.71263100 | -1.65745300 |
| N | -3.47674100 | 0.68426300 | -0.60141100 |
| C | -2.40575900 | 1.60882700 | -0.23113700 |
| H | -1.98040500 | 2.07009600 | -1.12705300 |
| C | -3.12480200 | 2.64742400 | 0.64754700 |
| C | -1.24329500 | 0.96165100 | 0.50244300 |
| O | -0.11295300 | 1.44062600 | 0.40975100 |

| | | | |
|---|---|---|---|
| N | -1.51324600 | -0.10813500 | 1.26897600 |
| H | -2.44955000 | -0.51341000 | 1.28359000 |
| C | -0.45013000 | -0.76217200 | 2.00881400 |
| H | 0.04135700 | -0.03122400 | 2.65618100 |
| C | -1.01011200 | -1.90609400 | 2.84930700 |
| C | 0.65846200 | -1.29272000 | 1.09970100 |
| H | -1.47926900 | -2.65808100 | 2.21142100 |
| H | -0.20232800 | -2.37431100 | 3.41127600 |
| H | -1.75425500 | -1.52664200 | 3.55087900 |
| O | 1.78553200 | -1.47999900 | 1.55822500 |
| N | 0.34531500 | -1.55631000 | -0.17950400 |
| H | -0.57438200 | -1.31549600 | -0.54834800 |
| C | 1.35460900 | -2.04148700 | -1.10318500 |
| H | 1.81755300 | -2.93987300 | -0.68759700 |
| C | 0.71642900 | -2.35873600 | -2.45254700 |
| C | 2.50932900 | -1.05459400 | -1.30063500 |
| H | 0.25178600 | -1.46531600 | -2.87447700 |
| H | 1.47743500 | -2.72745600 | -3.13959000 |
| H | -0.05146600 | -3.12404200 | -2.33112100 |
| O | 3.58472900 | -1.45514500 | -1.72992800 |
| N | 2.27281300 | 0.24127000 | -1.01297500 |
| H | 1.37969100 | 0.52964100 | -0.62009600 |
| C | 3.32444800 | 1.23331300 | -1.12462800 |
| H | 3.85400800 | 1.07676500 | -2.06698800 |
| C | 2.72076900 | 2.63508100 | -1.09180500 |
| C | 4.39179200 | 1.11648500 | -0.02910200 |
| H | 2.18802800 | 2.79844400 | -0.15241100 |
| H | 3.51154300 | 3.37813800 | -1.18707800 |
| H | 2.01695500 | 2.75787500 | -1.91622800 |
| O | 5.44810600 | 1.73389600 | -0.14895900 |
| N | 4.10017300 | 0.37090500 | 1.04502400 |
| H | 3.24154000 | -0.16788500 | 1.07993800 |
| C | 5.05421800 | 0.22275100 | 2.12947500 |
| H | 5.25407100 | 1.18562600 | 2.60332000 |
| H | -4.04257900 | -2.20180500 | -2.13422500 |
| H | -4.96174300 | -0.73680400 | -2.53746200 |
| O | -4.07368100 | -1.56278800 | 1.50323400 |
| H | -4.46796300 | -1.51867400 | 2.38051700 |
| H | -4.03945200 | -2.50211400 | 1.29434100 |
| H | 4.63402600 | -0.45872300 | 2.86642300 |
| H | 5.99855600 | -0.18325800 | 1.76125700 |
| C | -4.80726300 | 1.17697100 | -0.21054200 |
| C | -4.54640300 | 2.65382600 | 0.08156000 |
| H | -2.63102300 | 3.61688300 | 0.61021300 |
| H | -3.13816000 | 2.30014800 | 1.68462700 |
| H | -4.57704400 | 3.22670800 | -0.84804300 |
| H | -5.27558200 | 3.07258500 | 0.77372400 |

| | | | |
|---|---|---|---|
| H | -5.14991900 | 0.63942000 | 0.67812300 |
| H | -5.52628500 | 1.02725800 | -1.01485900 |

1 2 1.0 3 1.0 45 1.0 46 1.0
2
3 4 2.0 5 1.5
4
5 6 1.0 52 1.0
6 7 1.0 8 1.0 9 1.0
7
8 53 1.0 54 1.0 55 1.0
9 10 2.0 11 1.5
10
11 12 1.0 13 1.0
12
13 14 1.0 15 1.0 16 1.0
14
15 17 1.0 18 1.0 19 1.0
16 20 2.0 21 1.5
17
18
19
20
21 22 1.0 23 1.0
22
23 24 1.0 25 1.0 26 1.0
24
25 27 1.0 28 1.0 29 1.0
26 30 2.0 31 1.5
27
28
29
30
31 32 1.0 33 1.0
32
33 34 1.0 35 1.0 36 1.0
34
35 37 1.0 38 1.0 39 1.0
36 40 2.0 41 1.5
37
38
39
40
41 42 1.0 43 1.0
42
43 44 1.0 50 1.0 51 1.0
44

45
46
47 48 1.0 49 1.0
48
49
50
51
52 53 1.0 58 1.0 59 1.0
53 56 1.0 57 1.0
54
55
56
57
58
59

**Geometry-optimized structure of Pro-Ala$_4$-NHMe in a 310-helix with an Ac-Ser Ncap in the PPII conformation**
optimized M06-2X/6-311++G(d,p)/H2O

0 1

| | | | |
|---|---|---|---|
| C | 3.88447000 | 0.40788900 | 0.60706600 |
| C | 2.66641300 | 0.02780000 | -0.23919500 |
| O | 1.70091700 | 0.79143000 | -0.30294800 |
| N | 2.63554000 | -1.20038900 | -0.77535900 |
| C | 1.40040800 | -1.69618500 | -1.38467800 |
| H | 1.14772800 | -1.10923600 | -2.27107900 |
| C | 1.74127300 | -3.15324700 | -1.74508300 |
| C | 0.19977700 | -1.61373600 | -0.45294500 |
| O | -0.93574400 | -1.50698800 | -0.91216300 |
| N | 0.43842900 | -1.70126800 | 0.86850200 |
| H | 1.39403400 | -1.70714900 | 1.22030400 |
| C | -0.65470100 | -1.64492800 | 1.82144200 |
| H | -1.36934400 | -2.44111400 | 1.59906600 |
| C | -0.11875800 | -1.80300100 | 3.24170400 |
| C | -1.45820100 | -0.34749800 | 1.72289300 |
| H | 0.57205700 | -0.99127700 | 3.48187000 |
| H | -0.94585500 | -1.78195000 | 3.95063700 |
| H | 0.40424600 | -2.75542500 | 3.34070700 |
| O | -2.62702800 | -0.32154100 | 2.10621600 |
| N | -0.83293600 | 0.73641400 | 1.23424700 |
| H | 0.10471000 | 0.66048000 | 0.84421600 |
| C | -1.53659500 | 1.99834000 | 1.09372900 |
| H | -1.97053200 | 2.27425600 | 2.05787600 |
| C | -0.56906300 | 3.08170100 | 0.62557200 |
| C | -2.72612200 | 1.91727400 | 0.13115100 |
| H | -0.12180000 | 2.80622200 | -0.33172500 |

| | | | |
|---|---|---|---|
| H | -1.10147400 | 4.02589400 | 0.51560300 |
| H | 0.22940000 | 3.20983000 | 1.35817100 |
| O | -3.61661500 | 2.75589500 | 0.19623000 |
| N | -2.71961200 | 0.92293000 | -0.77934600 |
| H | -1.98608100 | 0.21818800 | -0.77180000 |
| C | -3.82576900 | 0.74329800 | -1.69991900 |
| H | -4.12691400 | 1.72279400 | -2.07751100 |
| C | -3.39417600 | -0.15378200 | -2.85734700 |
| C | -5.07996000 | 0.15585900 | -1.03949000 |
| H | -3.06764500 | -1.12662200 | -2.48322700 |
| H | -4.22683600 | -0.29527000 | -3.54503300 |
| H | -2.56424800 | 0.30771200 | -3.39447200 |
| O | -6.14348900 | 0.16056700 | -1.65593200 |
| N | -4.94576700 | -0.37918300 | 0.18109100 |
| H | -4.06386600 | -0.31188400 | 0.67689800 |
| C | -6.08432800 | -0.96149200 | 0.86865500 |
| H | -6.50288600 | -1.78630700 | 0.28947900 |
| H | 4.74300700 | -0.21311700 | 0.35317900 |
| H | -5.74864100 | -1.33497600 | 1.83395300 |
| H | -6.86729100 | -0.21616800 | 1.02356300 |
| C | 3.75349200 | -2.15411200 | -0.90417200 |
| C | 3.25113200 | -3.10661200 | -1.98884200 |
| H | 1.16848200 | -3.49742800 | -2.60420200 |
| H | 1.51950000 | -3.80231600 | -0.89347400 |
| H | 3.45852600 | -2.68443100 | -2.97472700 |
| H | 3.72097100 | -4.08677800 | -1.92287400 |
| H | 3.91694000 | -2.66647500 | 0.04754600 |
| H | 4.66340800 | -1.63286300 | -1.19830800 |
| C | 4.62565200 | 2.22036300 | -0.80835400 |
| C | 4.87616600 | 3.69748500 | -0.96476300 |
| H | 4.81353900 | 4.24012200 | -0.02283300 |
| H | 5.86477000 | 3.83778800 | -1.40199200 |
| H | 4.13911500 | 4.09894400 | -1.66219400 |
| O | 4.76780400 | 1.42672100 | -1.73084500 |
| N | 4.23035600 | 1.79583200 | 0.41723100 |
| H | 3.98061800 | 2.47440900 | 1.12166400 |
| C | 3.51269300 | 0.16795600 | 2.06992900 |
| H | 4.37671300 | 0.38868700 | 2.70054700 |
| H | 2.68170800 | 0.82762800 | 2.34848500 |
| O | 3.12585400 | -1.19525800 | 2.18378700 |
| H | 3.16545900 | -1.45934000 | 3.10826500 |

1 2 1.0 44 1.0 61 1.0 63 1.0
2 3 2.0 4 1.5
3
4 5 1.0 47 1.0
5 6 1.0 7 1.0 8 1.0

6
7 48 1.0 49 1.0 50 1.0
8 9 2.0 10 1.5
9
10 11 1.0 12 1.0
11
12 13 1.0 14 1.0 15 1.0
13
14 16 1.0 17 1.0 18 1.0
15 19 2.0 20 1.5
16
17
18
19
20 21 1.0 22 1.0
21
22 23 1.0 24 1.0 25 1.0
23
24 26 1.0 27 1.0 28 1.0
25 29 2.0 30 1.5
26
27
28
29
30 31 1.0 32 1.0
31
32 33 1.0 34 1.0 35 1.0
33
34 36 1.0 37 1.0 38 1.0
35 39 2.0 40 1.5
36
37
38
39
40 41 1.0 42 1.0
41
42 43 1.0 45 1.0 46 1.0
43
44
45
46
47 48 1.0 53 1.0 54 1.0
48 51 1.0 52 1.0
49
50
51
52

53
54
55 56 1.0 60 2.0 61 1.5
56 57 1.0 58 1.0 59 1.0
57
58
59
60
61 62 1.0
62
63 64 1.0 65 1.0 66 1.0
64
65
66 67 1.0
67

**Geometry-optimized structure of Pro-Ala$_4$-NHMe in a $3_{10}$-helix with an Ac-Ser Ncap in the β conformation**

optimized M06-2X/6-311++G(d,p)/H2O

0 1

| | | | |
|---|---|---|---|
| C | -3.82516400 | -0.30506900 | 0.01603100 |
| C | -2.55625800 | 0.24837100 | -0.61374900 |
| O | -1.62042100 | -0.52121900 | -0.85531200 |
| N | -2.44424700 | 1.57348800 | -0.76747300 |
| C | -1.14834800 | 2.14728600 | -1.14011600 |
| H | -0.84796000 | 1.79756200 | -2.13036500 |
| C | -1.40761200 | 3.66502500 | -1.12306100 |
| C | -0.03133600 | 1.75796300 | -0.18197600 |
| O | 1.13550200 | 1.73050800 | -0.56624600 |
| N | -0.38207100 | 1.48508000 | 1.08879700 |
| H | -1.36280700 | 1.47288000 | 1.36381500 |
| C | 0.61988400 | 1.11641100 | 2.07144700 |
| H | 1.37580900 | 1.90332500 | 2.13280300 |
| C | -0.03735500 | 0.91310900 | 3.43415100 |
| C | 1.38833100 | -0.14746600 | 1.68364500 |
| H | -0.77496900 | 0.10835700 | 3.38649100 |
| H | 0.72053300 | 0.65295700 | 4.17245700 |
| H | -0.53496900 | 1.83124800 | 3.74994700 |
| O | 2.51033000 | -0.34578800 | 2.14711800 |
| N | 0.78394800 | -1.01456100 | 0.85405000 |
| H | -0.11162100 | -0.78310200 | 0.42987200 |
| C | 1.45431700 | -2.22988400 | 0.42838500 |
| H | 1.77816900 | -2.78673700 | 1.31099700 |
| C | 0.50328900 | -3.08151700 | -0.40817200 |
| C | 2.73634000 | -1.95718200 | -0.36462900 |
| H | 0.18036000 | -2.53412900 | -1.29606200 |

| | | | |
|---|---|---|---|
| H | 1.00733800 | -3.99725500 | -0.71550800 |
| H | -0.37849800 | -3.34322100 | 0.17914900 |
| O | 3.58690000 | -2.83262800 | -0.46608800 |
| N | 2.85573600 | -0.74798100 | -0.94961000 |
| H | 2.14460400 | -0.03505400 | -0.80885400 |
| C | 4.05495300 | -0.38460600 | -1.68002300 |
| H | 4.34874400 | -1.22787200 | -2.30875600 |
| C | 3.78303800 | 0.84411000 | -2.54414300 |
| C | 5.26030400 | -0.10892800 | -0.77151500 |
| H | 3.47434000 | 1.68654200 | -1.92104000 |
| H | 4.68499000 | 1.11617500 | -3.09091400 |
| H | 2.98640800 | 0.62960100 | -3.25790300 |
| O | 6.38104500 | -0.01725700 | -1.26803500 |
| N | 5.02253100 | 0.06530600 | 0.53502500 |
| H | 4.09160800 | -0.07981200 | 0.90962500 |
| C | 6.10601400 | 0.33844800 | 1.46217200 |
| H | 6.63133900 | 1.25255400 | 1.18094900 |
| H | -4.69407400 | 0.32410200 | -0.17107300 |
| H | 5.68372200 | 0.45963500 | 2.45764400 |
| H | 6.82495700 | -0.48358200 | 1.47411500 |
| C | -3.49926800 | 2.59982100 | -0.64799800 |
| C | -2.90332800 | 3.76915100 | -1.42886700 |
| H | -0.77583300 | 4.18633800 | -1.83949100 |
| H | -1.20320400 | 4.06152300 | -0.12467700 |
| H | -3.08050500 | 3.62855800 | -2.49734400 |
| H | -3.33088900 | 4.72350800 | -1.12527300 |
| H | -3.65537500 | 2.84757000 | 0.40445200 |
| H | -4.43338700 | 2.24145500 | -1.07670100 |
| C | -5.30594300 | -2.17628100 | -0.52816400 |
| C | -5.41172000 | -3.54946900 | -1.14337900 |
| H | -4.44197300 | -4.01848600 | -1.30340700 |
| H | -6.02211700 | -4.17692600 | -0.49487000 |
| H | -5.92380800 | -3.45541800 | -2.10296600 |
| O | -6.28169300 | -1.58588400 | -0.08024400 |
| N | -4.07017500 | -1.62233500 | -0.52370100 |
| H | -3.26688200 | -2.13709600 | -0.85896300 |
| C | -3.57950600 | -0.39932700 | 1.53076500 |
| H | -4.49570000 | -0.75571200 | 2.00602400 |
| H | -2.77389000 | -1.12128800 | 1.70890400 |
| O | -3.21438500 | 0.89027000 | 2.00599700 |
| H | -3.36652200 | 0.93500500 | 2.95495800 |

1 2 1.0 44 1.0 61 1.0 63 1.0
2 3 2.0 4 1.5
3
4 5 1.0 47 1.0
5 6 1.0 7 1.0 8 1.0

6
7 48 1.0 49 1.0 50 1.0
8 9 2.0 10 1.5
9
10 11 1.0 12 1.0
11
12 13 1.0 14 1.0 15 1.0
13
14 16 1.0 17 1.0 18 1.0
15 19 2.0 20 1.5
16
17
18
19
20 21 1.0 22 1.0
21
22 23 1.0 24 1.0 25 1.0
23
24 26 1.0 27 1.0 28 1.0
25 29 2.0 30 1.5
26
27
28
29
30 31 1.0 32 1.0
31
32 33 1.0 34 1.0 35 1.0
33
34 36 1.0 37 1.0 38 1.0
35 39 2.0 40 1.5
36
37
38
39
40 41 1.0 42 1.0
41
42 43 1.0 45 1.0 46 1.0
43
44
45
46
47 48 1.0 53 1.0 54 1.0
48 51 1.0 52 1.0
49
50
51
52

53
54
55 56 1.0 60 2.0 61 1.5
56 57 1.0 58 1.0 59 1.0
57
58
59
60
61 62 1.0
62
63 64 1.0 65 1.0 66 1.0
64
65
66 67 1.0
67

### Geometry-optimized structures of Pro-Ala$_6$-NHMe in $\alpha$-helices with an Ac-Ser Ncap

**Geometry-optimized structure of Pro-Ala$_6$-NHMe in an $\alpha$-helix with an Ac-Ser Ncap in the PPII conformation; one $H_2O$ molecule at the N-terminus and two $H_2O$ molecules at the C-terminus**
optimized M06-2X/6-311++G(d,p)/H2O

0 1

| | | | |
|---|---|---|---|
| C | 4.70895300 | 0.52482800 | -0.41444000 |
| O | 3.52848600 | 0.85741400 | -0.48458600 |
| N | 5.13929800 | -0.64324700 | -0.92888600 |
| C | 4.16060800 | -1.51441400 | -1.58536900 |
| H | 3.70028900 | -0.99428600 | -2.42975500 |
| C | 5.00665500 | -2.70883800 | -2.05190800 |
| C | 3.02010300 | -1.90306300 | -0.65472200 |
| O | 1.87853300 | -2.03767500 | -1.08501400 |
| N | 3.32739800 | -2.12529200 | 0.63745500 |
| H | 4.26358200 | -1.93826500 | 0.97243300 |
| C | 2.28492900 | -2.46985700 | 1.59025100 |
| H | 1.71296300 | -3.31160400 | 1.19486700 |
| C | 2.89774500 | -2.83031200 | 2.93856900 |
| C | 1.27691200 | -1.32742200 | 1.74161000 |
| H | 3.48632700 | -1.99730600 | 3.32928800 |
| H | 2.10806000 | -3.06272700 | 3.65330900 |
| H | 3.54406000 | -3.70261900 | 2.83481400 |
| O | 0.07086400 | -1.55735300 | 1.81115500 |
| N | 1.78304400 | -0.08698900 | 1.81699200 |
| H | 2.78375400 | 0.06925800 | 1.72067700 |
| C | 0.89965300 | 1.06343600 | 1.88992500 |
| H | 0.22586400 | 0.95176200 | 2.74226900 |
| C | 1.72054800 | 2.34351200 | 2.01384800 |
| C | -0.00643800 | 1.13121000 | 0.65884600 |
| H | 2.38384800 | 2.44016400 | 1.15080600 |
| H | 1.05845500 | 3.20886000 | 2.05510600 |
| H | 2.32071300 | 2.31553100 | 2.92512300 |
| O | -1.19803100 | 1.41648100 | 0.76590200 |
| N | 0.57569900 | 0.88947200 | -0.52575500 |
| H | 1.57103400 | 0.67849400 | -0.57033400 |
| C | -0.21135500 | 0.91127300 | -1.74630200 |
| H | -0.72796400 | 1.87057800 | -1.82959700 |
| C | 0.69529800 | 0.68332700 | -2.95173700 |
| C | -1.31880200 | -0.14456000 | -1.69495300 |
| H | 1.21210000 | -0.27478700 | -2.85848800 |
| H | 0.10321500 | 0.68126400 | -3.86741800 |
| H | 1.43581000 | 1.48237200 | -3.01637300 |
| O | -2.45816200 | 0.10743100 | -2.08425400 |
| N | -0.96624200 | -1.35015500 | -1.22351100 |
| H | -0.00601800 | -1.52683900 | -0.93445500 |
| C | -1.94812900 | -2.41278500 | -1.11907400 |
| H | -2.41962200 | -2.57220400 | -2.09155600 |
| C | -1.27718800 | -3.69352000 | -0.63150900 |

| | | | |
|---|---|---|---|
| C | -3.08834000 | -2.01379200 | -0.18065100 |
| H | -0.79498700 | -3.52176900 | 0.33367900 |
| H | -2.01955800 | -4.48516700 | -0.52494500 |
| H | -0.52274900 | -4.01427900 | -1.35143600 |
| O | -4.25429600 | -2.28743900 | -0.46204200 |
| N | -2.73622700 | -1.38514900 | 0.95099300 |
| H | -1.75102600 | -1.21742500 | 1.15092300 |
| C | -3.73505000 | -0.94552600 | 1.90728900 |
| H | -4.40197900 | -1.77758800 | 2.14797600 |
| C | -3.04757100 | -0.42939400 | 3.16893700 |
| C | -4.63766000 | 0.14326300 | 1.32291200 |
| H | -2.36942500 | 0.38872500 | 2.91459900 |
| H | -3.79292100 | -0.07012200 | 3.87892000 |
| H | -2.47494000 | -1.23281800 | 3.63412000 |
| O | -5.83662300 | 0.18568300 | 1.62158500 |
| N | -4.06093500 | 1.03549500 | 0.51503200 |
| H | -3.05514800 | 0.99511500 | 0.35465000 |
| C | -4.84192900 | 2.06832900 | -0.14284300 |
| H | -5.44963700 | 2.59160700 | 0.59903100 |
| C | -3.90681900 | 3.04756900 | -0.84702900 |
| C | -5.83220000 | 1.47108300 | -1.15180700 |
| H | -3.28922900 | 2.51535700 | -1.57434700 |
| H | -4.48954500 | 3.81234000 | -1.36106200 |
| H | -3.25562200 | 3.53070300 | -0.11714300 |
| O | -6.93615700 | 1.99660300 | -1.32572200 |
| N | -5.41326800 | 0.40706900 | -1.83503500 |
| H | -4.47137300 | 0.05870000 | -1.68516000 |
| C | -6.25053200 | -0.23814000 | -2.83002400 |
| H | -5.74003700 | -1.13316400 | -3.17925000 |
| H | -6.43782600 | 0.42563800 | -3.67670300 |
| H | -7.20860900 | -0.52045500 | -2.39062700 |
| O | -6.90130000 | -2.25111200 | 0.54642400 |
| H | -6.82414400 | -1.42426700 | 1.04064700 |
| H | -6.01714800 | -2.33839300 | 0.15700000 |
| O | -8.00092600 | 2.04323600 | 1.33098500 |
| H | -7.83484400 | 2.20281600 | 0.38997300 |
| H | -7.30055700 | 1.41439700 | 1.56095900 |
| C | 6.52891500 | -1.08841900 | -1.15757000 |
| H | 7.17125400 | -0.24427100 | -1.40360500 |
| H | 6.92243700 | -1.59217000 | -0.27066100 |
| C | 6.37228200 | -2.07024400 | -2.31905100 |
| H | 6.35279100 | -1.52314900 | -3.26403700 |
| H | 7.18239700 | -2.79690200 | -2.35198700 |
| H | 4.57132200 | -3.19148800 | -2.92479300 |
| H | 5.08755500 | -3.44531100 | -1.24768100 |
| C | 5.71866700 | 1.36269300 | 0.38310000 |
| H | 6.65724200 | 1.44572300 | -0.16733100 |
| N | 5.19481600 | 2.68247600 | 0.62801300 |
| H | 4.68252100 | 2.82043100 | 1.48848500 |
| C | 4.98276300 | 3.53326100 | -0.40923300 |
| O | 5.39998700 | 3.28240800 | -1.53203800 |

| | | | |
|---|---|---|---|
| C | 4.20924200 | 4.78817200 | -0.09414900 |
| H | 4.12995800 | 4.97867500 | 0.97551000 |
| H | 4.69041500 | 5.63315000 | -0.58511400 |
| H | 3.20515900 | 4.68020000 | -0.51114200 |
| C | 5.98162000 | 0.66688200 | 1.71358600 |
| H | 6.48738800 | -0.28743100 | 1.55006900 |
| H | 6.62006800 | 1.30399800 | 2.32945200 |
| O | 4.71470600 | 0.46443700 | 2.33541200 |
| H | 4.84530800 | 0.28295500 | 3.27176600 |

1 2 2.0 3 1.5 89 1.0
2
3 4 1.0 81 1.0
4 5 1.0 6 1.0 7 1.0
5
6 84 1.0 87 1.0 88 1.0
7 8 2.0 9 1.5
8
9 10 1.0 11 1.0
10
11 12 1.0 13 1.0 14 1.0
12
13 15 1.0 16 1.0 17 1.0
14 18 2.0 19 1.5
15
16
17
18
19 20 1.0 21 1.0
20
21 22 1.0 23 1.0 24 1.0
22
23 25 1.0 26 1.0 27 1.0
24 28 2.0 29 1.5
25
26
27
28
29 30 1.0 31 1.0
30
31 32 1.0 33 1.0 34 1.0
32
33 35 1.0 36 1.0 37 1.0
34 38 2.0 39 1.5
35
36
37
38
39 40 1.0 41 1.0
40
41 42 1.0 43 1.0 44 1.0

42
43 45 1.0 46 1.0 47 1.0
44 48 2.0 49 1.5
45
46
47
48
49 50 1.0 51 1.0
50
51 52 1.0 53 1.0 54 1.0
52
53 55 1.0 56 1.0 57 1.0
54 58 2.0 59 1.5
55
56
57
58
59 60 1.0 61 1.0
60
61 62 1.0 63 1.0 64 1.0
62
63 65 1.0 66 1.0 67 1.0
64 68 2.0 69 1.5
65
66
67
68
69 70 1.0 71 1.0
70
71 72 1.0 73 1.0 74 1.0
72
73
74
75 76 1.0 77 1.0
76
77
78 79 1.0 80 1.0
79
80
81 82 1.0 83 1.0 84 1.0
82
83
84 85 1.0 86 1.0
85
86
87
88
89 90 1.0 91 1.0 99 1.0
90
91 92 1.0 93 1.5
92

93 94 2.0 95 1.0
94
95 96 1.0 97 1.0 98 1.0
96
97
98
99 100 1.0 101 1.0 102 1.0
100
101
102 103 1.0
103

**Geometry-optimized structure of Pro-Ala$_6$-NHMe in an $\alpha$-helix with an Ac-Ser Ncap in the $\beta$ conformation; one $H_2O$ molecule at the N-terminus and two $H_2O$ molecules at the C-terminus**
optimized M06-2X/6-311++G(d,p)/H2O

0 1

| | | | |
|---|---|---|---|
| C | 4.48190000 | 0.50110000 | -0.79220000 |
| O | 3.27900000 | 0.80060000 | -0.83550000 |
| N | 4.91060000 | -0.71690000 | -1.15080000 |
| C | 3.93750000 | -1.66110000 | -1.71740000 |
| H | 3.47950000 | -1.21960000 | -2.60930000 |
| C | 4.79580000 | -2.88710000 | -2.06870000 |
| C | 2.79030000 | -1.96840000 | -0.76420000 |
| O | 1.62630000 | -2.01320000 | -1.17600000 |
| N | 3.10590000 | -2.21410000 | 0.51930000 |
| H | 4.07520000 | -2.18770000 | 0.84690000 |
| C | 2.04630000 | -2.54490000 | 1.45550000 |
| H | 1.43680000 | -3.35130000 | 1.03490000 |
| C | 2.61650000 | -2.96820000 | 2.80680000 |
| C | 1.08960000 | -1.36730000 | 1.63250000 |
| H | 3.18530000 | -2.15510000 | 3.26650000 |
| H | 1.79870000 | -3.24260000 | 3.47640000 |
| H | 3.27180000 | -3.83360000 | 2.68280000 |
| O | -0.12120000 | -1.55470000 | 1.78850000 |
| N | 1.63290000 | -0.13620000 | 1.65120000 |
| H | 2.62070000 | -0.01000000 | 1.43760000 |
| C | 0.77010000 | 1.02620000 | 1.77090000 |
| H | 0.11300000 | 0.89170000 | 2.63540000 |
| C | 1.60480000 | 2.29250000 | 1.92910000 |
| C | -0.17110000 | 1.12950000 | 0.56750000 |
| H | 2.31720000 | 2.38260000 | 1.10370000 |
| H | 0.95780000 | 3.17200000 | 1.93390000 |
| H | 2.15290000 | 2.26240000 | 2.87400000 |
| O | -1.35470000 | 1.45270000 | 0.71870000 |
| N | 0.35800000 | 0.86860000 | -0.63760000 |
| H | 1.33810000 | 0.60180000 | -0.71950000 |
| C | -0.47470000 | 0.89330000 | -1.82670000 |
| H | -0.98700000 | 1.85860000 | -1.89310000 |
| C | 0.38460000 | 0.65310000 | -3.06550000 |

| | | | |
|---|---|---|---|
| C | -1.58700000 | -0.15320000 | -1.72300000 |
| H | 0.90530000 | -0.30640000 | -2.98490000 |
| H | -0.24270000 | 0.64580000 | -3.95930000 |
| H | 1.12510000 | 1.45070000 | -3.16420000 |
| O | -2.74170000 | 0.10220000 | -2.08250000 |
| N | -1.23240000 | -1.35410000 | -1.23820000 |
| H | -0.26310000 | -1.53740000 | -0.97910000 |
| C | -2.22320000 | -2.40320000 | -1.09040000 |
| H | -2.72140000 | -2.57540000 | -2.04980000 |
| C | -1.55470000 | -3.68290000 | -0.59390000 |
| C | -3.33420000 | -1.96580000 | -0.13330000 |
| H | -1.05250000 | -3.50140000 | 0.36160000 |
| H | -2.30260000 | -4.46770000 | -0.46140000 |
| H | -0.81560000 | -4.02330000 | -1.32350000 |
| O | -4.51700000 | -2.22380000 | -0.38330000 |
| N | -2.94540000 | -1.31990000 | 0.97760000 |
| H | -1.95160000 | -1.16440000 | 1.14860000 |
| C | -3.91230000 | -0.84560000 | 1.95000000 |
| H | -4.58960000 | -1.66290000 | 2.22060000 |
| C | -3.18380000 | -0.32300000 | 3.18690000 |
| C | -4.80820000 | 0.24900000 | 1.36560000 |
| H | -2.49540000 | 0.47980000 | 2.90410000 |
| H | -3.90480000 | 0.06110000 | 3.91130000 |
| H | -2.61390000 | -1.13150000 | 3.65080000 |
| O | -6.00210000 | 0.32510000 | 1.69960000 |
| N | -4.23920000 | 1.11030000 | 0.51790000 |
| H | -3.23900000 | 1.04450000 | 0.32700000 |
| C | -5.02010000 | 2.14410000 | -0.13790000 |
| H | -5.59590000 | 2.69420000 | 0.61340000 |
| C | -4.08860000 | 3.09300000 | -0.88870000 |
| C | -6.05250000 | 1.54240000 | -1.10060000 |
| H | -3.50260000 | 2.53680000 | -1.62710000 |
| H | -4.67300000 | 3.86170000 | -1.39830000 |
| H | -3.40520000 | 3.57680000 | -0.18650000 |
| O | -7.15000000 | 2.09790000 | -1.26630000 |
| N | -5.68580000 | 0.44210000 | -1.75780000 |
| H | -4.75060000 | 0.06840000 | -1.61800000 |
| C | -6.56930000 | -0.20780000 | -2.70810000 |
| H | -6.09240000 | -1.12690000 | -3.04720000 |
| H | -6.76540000 | 0.43830000 | -3.56860000 |
| H | -7.52260000 | -0.45170000 | -2.23220000 |
| O | 5.65340000 | -1.96640000 | 1.87090000 |
| H | 5.89680000 | -2.61950000 | 2.53750000 |
| H | 5.39240000 | -1.16030000 | 2.34770000 |
| O | -7.13100000 | -2.13030000 | 0.73810000 |
| H | -7.03580000 | -1.28680000 | 1.20610000 |
| H | -6.26180000 | -2.23760000 | 0.31550000 |
| O | -8.14550000 | 2.22770000 | 1.41540000 |
| H | -7.99990000 | 2.36530000 | 0.46450000 |
| H | -7.45280000 | 1.58840000 | 1.65040000 |
| C | 6.30040000 | -1.19230000 | -1.30300000 |

| | | | |
|---|---|---|---|
| H | 6.95990000 | -0.37620000 | -1.60200000 |
| H | 6.64520000 | -1.61750000 | -0.35490000 |
| C | 6.16400000 | -2.27260000 | -2.37610000 |
| H | 6.15930000 | -1.81480000 | -3.37010000 |
| H | 6.97590000 | -2.99970000 | -2.32890000 |
| H | 4.36900000 | -3.44750000 | -2.90130000 |
| H | 4.86870000 | -3.55300000 | -1.20170000 |
| C | 5.49580000 | 1.50390000 | -0.25040000 |
| H | 6.37480000 | 1.56930000 | -0.90200000 |
| N | 4.84940000 | 2.79580000 | -0.19280000 |
| H | 3.86110000 | 2.82240000 | -0.41030000 |
| C | 5.56900000 | 3.93650000 | -0.07300000 |
| O | 6.79320000 | 3.91450000 | 0.06690000 |
| C | 4.79060000 | 5.22840000 | -0.11850000 |
| H | 3.71090000 | 5.07460000 | -0.13950000 |
| H | 5.05620000 | 5.82590000 | 0.75560000 |
| H | 5.09180000 | 5.78320000 | -1.01050000 |
| C | 5.95770000 | 1.03850000 | 1.13560000 |
| H | 6.59590000 | 0.15650000 | 1.03900000 |
| H | 6.53180000 | 1.83890000 | 1.60720000 |
| O | 4.83800000 | 0.66220000 | 1.93870000 |
| H | 4.58740000 | 1.39900000 | 2.50950000 |

1 2 2.0 3 1.5 92 1.0
2
3 4 1.0 84 1.0
4 5 1.0 6 1.0 7 1.0
5
6 87 1.0 90 1.0 91 1.0
7 8 2.0 9 1.5
8
9 10 1.0 11 1.0
10
11 12 1.0 13 1.0 14 1.0
12
13 15 1.0 16 1.0 17 1.0
14 18 2.0 19 1.5
15
16
17
18
19 20 1.0 21 1.0
20
21 22 1.0 23 1.0 24 1.0
22
23 25 1.0 26 1.0 27 1.0
24 28 2.0 29 1.5
25
26
27
28

29 30 1.0 31 1.0
30
31 32 1.0 33 1.0 34 1.0
32
33 35 1.0 36 1.0 37 1.0
34 38 2.0 39 1.5
35
36
37
38
39 40 1.0 41 1.0
40
41 42 1.0 43 1.0 44 1.0
42
43 45 1.0 46 1.0 47 1.0
44 48 2.0 49 1.5
45
46
47
48
49 50 1.0 51 1.0
50
51 52 1.0 53 1.0 54 1.0
52
53 55 1.0 56 1.0 57 1.0
54 58 2.0 59 1.5
55
56
57
58
59 60 1.0 61 1.0
60
61 62 1.0 63 1.0 64 1.0
62
63 65 1.0 66 1.0 67 1.0
64 68 2.0 69 1.5
65
66
67
68
69 70 1.0 71 1.0
70
71 72 1.0 73 1.0 74 1.0
72
73
74
75 76 1.0 77 1.0
76
77
78 79 1.0 80 1.0
79

80
81 82 1.0 83 1.0
82
83
84 85 1.0 86 1.0 87 1.0
85
86
87 88 1.0 89 1.0
88
89
90
91
92 93 1.0 94 1.0 102 1.0
93
94 95 1.0 96 1.5
95
96 97 2.0 98 1.0
97
98 99 1.0 100 1.0 101 1.0
99
100
101
102 103 1.0 104 1.0 105 1.0
103
104
105 106 1.0
106

### Geometry-optimized structures of Ac-Ser-*trans*-Pro-Ser-NHMe and Ac-Ser-trans-Pro-Thr-NHMe type I β-turns

**Ac-Ser-*trans*-Pro-Ser-NHMe in a type I β-turn with the N-terminal Ser in the PPII conformation**
optimized M06-2X/6-311++G(d,p)/H2O

0 1

| | | | |
|---|---|---|---|
| C | -4.69083400 | -2.60606400 | -0.63705300 |
| C | -3.91087300 | -1.37511800 | -0.25480700 |
| O | -4.12343000 | -0.27881300 | -0.75945900 |
| N | -2.94940200 | -1.53014500 | 0.68839800 |
| C | -2.10619700 | -0.40961500 | 1.02694700 |
| C | -1.26079900 | 0.02487400 | -0.17743300 |
| O | -0.72826100 | -0.82514300 | -0.89165400 |
| C | -1.13985800 | -0.80633200 | 2.14444600 |
| O | -0.32340600 | 0.32604200 | 2.39109400 |
| H | -2.69116700 | -2.45888500 | 0.98786200 |
| H | 0.37113100 | 0.12681500 | 3.03212100 |
| N | -1.02077100 | 1.33761800 | -0.32442800 |
| C | -0.01223200 | 1.80744900 | -1.27795000 |
| C | 1.37116200 | 1.20328400 | -1.06473200 |
| O | 2.19175600 | 1.21262200 | -1.97305600 |
| C | 0.01121800 | 3.33039800 | -1.05242100 |
| C | -1.40555500 | 3.63500200 | -0.56581100 |
| C | -1.72218200 | 2.45016500 | 0.34454300 |
| N | 1.63461800 | 0.70374400 | 0.15947900 |
| C | 2.88792600 | 0.04232700 | 0.44646500 |
| C | 3.18007200 | -1.14008900 | -0.48694800 |
| O | 4.34032200 | -1.48909600 | -0.68219500 |
| C | 2.84578200 | -0.52714000 | 1.86987400 |
| O | 2.46389000 | 0.43275700 | 2.83780500 |
| H | 3.18141000 | 1.05884400 | 2.97820000 |
| H | 0.93043100 | 0.74984900 | 0.89348500 |
| N | 2.11704500 | -1.79460400 | -0.97197400 |
| C | 2.28717200 | -2.94200500 | -1.84589400 |
| H | 1.18390500 | -1.41549500 | -0.83933500 |
| H | -4.43866400 | -3.47431700 | -0.03032200 |
| H | -5.75422800 | -2.38810300 | -0.53734100 |
| H | -4.48914200 | -2.82820300 | -1.68613500 |
| H | -2.73038800 | 0.41159200 | 1.37780300 |
| H | -0.53747800 | -1.66127900 | 1.81352700 |
| H | -1.71003100 | -1.09035800 | 3.03301000 |
| H | 2.81840000 | -3.74345600 | -1.32983800 |
| H | -0.30650800 | 1.55687600 | -2.30028500 |
| H | 3.72586000 | 0.73727400 | 0.33999900 |
| H | 2.08961200 | -1.31591100 | 1.91430500 |

| | | | |
|---|---|---|---|
| H | 0.73451800 | 3.57123800 | -0.26794500 |
| H | 0.29089700 | 3.86561700 | -1.95781600 |
| H | -1.47909500 | 4.58660800 | -0.04132700 |
| H | -2.09850300 | 3.64651300 | -1.41026000 |
| H | -2.79014200 | 2.24489900 | 0.40140600 |
| H | -1.32210000 | 2.59123700 | 1.35220000 |
| H | 2.85475300 | -2.66918300 | -2.73822400 |
| H | 1.30318600 | -3.29843900 | -2.14287800 |
| H | 3.81713800 | -0.96979300 | 2.10181600 |

1 2 1.0 30 1.0 31 1.0 32 1.0
2 3 2.0 4 1.5
3
4 5 1.0 10 1.0
5 6 1.0 8 1.0 33 1.0
6 7 2.0 12 1.5
7
8 9 1.0 34 1.0 35 1.0
9 11 1.0
10
11
12 13 1.0 18 1.0
13 14 1.0 16 1.0 37 1.0
14 15 2.0 19 1.5
15
16 17 1.0 40 1.0 41 1.0
17 18 1.0 42 1.0 43 1.0
18 44 1.0 45 1.0
19 20 1.0 26 1.0
20 21 1.0 23 1.0 38 1.0
21 22 2.0 27 1.5
22
23 24 1.0 39 1.0 48 1.0
24 25 1.0
25
26
27 28 1.0 29 1.0
28 36 1.0 46 1.0 47 1.0

**Ac-Ser-*trans*-Pro-Ser-NHMe in a type I β-turn with the N-terminal Ser in the β conformation**
optimized M06-2X/6-311++G(d,p)/H2O

0 1

| | | | |
|---|---|---|---|
| C | -5.03518400 | -2.28835200 | -0.92127300 |
| C | -4.31880200 | -1.24082700 | -0.10599700 |
| O | -4.87575600 | -0.63129200 | 0.79971200 |
| N | -3.02778500 | -1.00614600 | -0.43959700 |
| C | -2.22263600 | -0.02957600 | 0.25582400 |
| C | -1.09315000 | 0.41797100 | -0.66408700 |
| O | -0.57285400 | -0.40576600 | -1.42203000 |
| C | -1.57752900 | -0.66697600 | 1.49831200 |
| O | -0.73309200 | 0.30488300 | 2.09327500 |
| H | -2.57283600 | -1.54023500 | -1.16751000 |
| H | -0.23769200 | -0.06793100 | 2.83404400 |
| N | -0.61903700 | 1.66321600 | -0.52250200 |
| C | 0.63691400 | 2.05062900 | -1.17349800 |
| C | 1.81904700 | 1.16562600 | -0.79043200 |
| O | 2.81228700 | 1.12245100 | -1.50345700 |
| C | 0.85508500 | 3.50166400 | -0.70586900 |
| C | -0.56223400 | 3.99906300 | -0.41862900 |
| C | -1.23118600 | 2.78369900 | 0.21915800 |
| N | 1.70928300 | 0.47651500 | 0.36388200 |
| C | 2.74468500 | -0.43556500 | 0.79522100 |
| C | 3.03016500 | -1.55546100 | -0.21426400 |
| O | 4.11992600 | -2.11936100 | -0.20188100 |
| C | 2.30517400 | -1.12107000 | 2.09521600 |
| O | 1.88488600 | -0.20240600 | 3.08690700 |
| H | 2.64753800 | 0.26497700 | 3.44237600 |
| H | 0.89032100 | 0.59731700 | 0.95756000 |
| N | 2.01433200 | -1.92282800 | -1.00658600 |
| C | 2.17446300 | -2.99714100 | -1.97065700 |
| H | 1.16254400 | -1.37098700 | -1.02870600 |
| H | -4.39534600 | -2.76330300 | -1.66312800 |
| H | -5.43001800 | -3.04449100 | -0.24208900 |

| | | | |
|---|---|---|---|
| H | -5.87951900 | -1.81378400 | -1.42321100 |
| H | -2.86026300 | 0.79638800 | 0.56753200 |
| H | -1.00642100 | -1.54618300 | 1.17687100 |
| H | -2.37256600 | -0.98342800 | 2.17849600 |
| H | 2.47617600 | -3.91694600 | -1.46773200 |
| H | 0.53669000 | 1.98662500 | -2.25952700 |
| H | 3.68975800 | 0.09413700 | 0.94620400 |
| H | 1.44021400 | -1.75649500 | 1.88666500 |
| H | 1.44455900 | 3.50578000 | 0.21527000 |
| H | 1.38314200 | 4.08691500 | -1.45615000 |
| H | -0.58144200 | 4.86842900 | 0.23687200 |
| H | -1.07054500 | 4.25216000 | -1.35172400 |
| H | -2.31171000 | 2.78565600 | 0.08871300 |
| H | -0.99442200 | 2.69789800 | 1.28232600 |
| H | 2.93213700 | -2.74462400 | -2.71598300 |
| H | 1.22097600 | -3.15731800 | -2.46953900 |
| H | 3.11914200 | -1.75603900 | 2.45164900 |

1 2 1.0 30 1.0 31 1.0 32 1.0
2 3 2.0 4 1.5
3
4 5 1.0 10 1.0
5 6 1.0 8 1.0 33 1.0
6 7 2.0 12 1.5
7
8 9 1.0 34 1.0 35 1.0
9 11 1.0
10
11
12 13 1.0 18 1.0
13 14 1.0 16 1.0 37 1.0
14 15 2.0 19 1.5
15
16 17 1.0 40 1.0 41 1.0
17 18 1.0 42 1.0 43 1.0
18 44 1.0 45 1.0
19 20 1.0 26 1.0
20 21 1.0 23 1.0 38 1.0
21 22 2.0 27 1.5
22
23 24 1.0 39 1.0 48 1.0
24 25 1.0
25
26
27 28 1.0 29 1.0
28 36 1.0 46 1.0 47 1.0

**Ac-Ser-*trans*-Pro-Thr-NHMe in a type I β-turn with Ser in the PPII conformation**
optimized M06-2X/6-311++G(d,p)/H2O

0 1

| | | | |
|---|---|---|---|
| C | -4.70940000 | -2.81050000 | -0.62300000 |
| C | -4.00690000 | -1.55300000 | -0.18150000 |
| O | -4.32940000 | -0.44260000 | -0.58690000 |
| N | -2.98350000 | -1.70290000 | 0.69490000 |
| C | -2.19520000 | -0.55200000 | 1.06310000 |
| C | -1.45290000 | 0.01770000 | -0.15220000 |
| O | -0.92230000 | -0.74580000 | -0.95970000 |
| C | -1.13790000 | -0.95240000 | 2.09380000 |
| O | -0.40380000 | 0.22120000 | 2.39420000 |
| H | -2.64190000 | -2.62950000 | 0.90430000 |
| H | 0.39670000 | 0.01870000 | 2.89760000 |
| N | -1.29380000 | 1.34920000 | -0.21130000 |
| C | -0.37070000 | 1.94770000 | -1.17870000 |
| C | 1.04780000 | 1.39330000 | -1.11210000 |
| O | 1.79160000 | 1.49800000 | -2.07900000 |
| C | -0.39730000 | 3.44350000 | -0.81620000 |
| C | -1.79930000 | 3.63770000 | -0.23790000 |
| C | -2.01830000 | 2.36460000 | 0.57760000 |
| N | 1.43190000 | 0.82860000 | 0.05040000 |
| C | 2.73160000 | 0.20200000 | 0.18650000 |
| C | 2.99950000 | -0.81940000 | -0.92970000 |
| O | 4.13360000 | -0.98320000 | -1.36550000 |
| C | 2.79730000 | -0.52390000 | 1.54560000 |
| C | 4.17200000 | -1.09960000 | 1.83350000 |
| O | 2.36670000 | 0.33890000 | 2.59370000 |

| | | | |
|---|---|---|---|
| H | 3.05580000 | 0.98410000 | 2.78910000 |
| H | 0.79280000 | 0.80480000 | 0.84220000 |
| N | 1.94540000 | -1.55630000 | -1.31280000 |
| C | 2.07660000 | -2.53220000 | -2.38050000 |
| H | 1.01050000 | -1.27420000 | -1.03180000 |
| H | -4.42030000 | -3.68420000 | -0.04080000 |
| H | -5.78500000 | -2.65580000 | -0.54270000 |
| H | -4.47070000 | -2.98540000 | -1.67370000 |
| H | -2.84900000 | 0.19890000 | 1.50570000 |
| H | -0.48940000 | -1.72490000 | 1.66200000 |
| H | -1.63150000 | -1.35570000 | 2.98260000 |
| H | 2.81530000 | -3.28940000 | -2.11480000 |
| H | -0.72140000 | 1.77650000 | -2.19970000 |
| H | 3.53250000 | 0.94600000 | 0.12250000 |
| H | 2.05320000 | -1.32740000 | 1.52730000 |
| H | 4.92300000 | -0.30430000 | 1.84040000 |
| H | 4.16280000 | -1.58520000 | 2.80970000 |
| H | 4.45840000 | -1.83070000 | 1.07740000 |
| H | 0.35450000 | 3.64510000 | -0.04760000 |
| H | -0.18560000 | 4.06940000 | -1.68110000 |
| H | -1.88550000 | 4.53550000 | 0.37250000 |
| H | -2.53410000 | 3.69120000 | -1.04460000 |
| H | -3.07190000 | 2.10000000 | 0.65740000 |
| H | -1.58350000 | 2.43950000 | 1.57770000 |
| H | 2.38990000 | -2.05380000 | -3.31180000 |
| H | 1.11140000 | -3.01150000 | -2.53060000 |

1 2 1.0 31 1.0 32 1.0 33 1.0
2 3 2.0 4 1.5
3
4 5 1.0 10 1.0
5 6 1.0 8 1.0 34 1.0
6 7 2.0 12 1.5
7
8 9 1.0 35 1.0 36 1.0
9 11 1.0
10
11
12 13 1.0 18 1.0
13 14 1.0 16 1.0 38 1.0
14 15 2.0 19 1.5
15
16 17 1.0 44 1.0 45 1.0
17 18 1.0 46 1.0 47 1.0
18 48 1.0 49 1.0
19 20 1.0 27 1.0
20 21 1.0 23 1.0 39 1.0
21 22 2.0 28 1.5
22
23 24 1.0 25 1.0 40 1.0
24 41 1.0 42 1.0 43 1.0

25 26 1.0
26
27
28 29 1.0 30 1.0
29 37 1.0 50 1.0 51 1.0

**Ac-Ser-*trans*-Pro-Thr-NHMe in a type I β-turn with Ser in the β conformation**
optimized M06-2X/6-311++G(d,p)/H2O

0 1

| | | | |
|---|---|---|---|
| C | -5.00040000 | -2.54800000 | -1.07610000 |
| C | -4.35440000 | -1.53580000 | -0.16310000 |
| O | -4.92750000 | -1.09580000 | 0.82690000 |
| N | -3.10700000 | -1.13730000 | -0.50510000 |
| C | -2.36560000 | -0.17880000 | 0.28120000 |
| C | -1.31970000 | 0.47870000 | -0.61030000 |
| O | -0.76830000 | -0.19210000 | -1.48750000 |
| C | -1.61750000 | -0.89530000 | 1.41880000 |
| O | -0.84720000 | 0.07190000 | 2.11130000 |
| H | -2.63730000 | -1.53080000 | -1.30920000 |
| H | -0.19470000 | -0.35020000 | 2.68730000 |
| N | -0.94590000 | 1.73360000 | -0.32400000 |
| C | 0.23630000 | 2.31420000 | -0.96730000 |
| C | 1.50710000 | 1.49580000 | -0.76790000 |
| O | 2.45180000 | 1.63370000 | -1.53330000 |
| C | 0.36010000 | 3.70230000 | -0.31170000 |
| C | -1.07840000 | 4.03240000 | 0.08940000 |
| C | -1.61690000 | 2.69000000 | 0.57900000 |

| | | | |
|---|---|---|---|
| N | 1.53280000 | 0.65610000 | 0.28760000 |
| C | 2.66510000 | -0.21670000 | 0.52360000 |
| C | 3.02100000 | -1.04540000 | -0.71990000 |
| O | 4.18670000 | -1.33700000 | -0.96150000 |
| C | 2.32190000 | -1.17980000 | 1.67810000 |
| C | 3.49630000 | -2.05400000 | 2.07880000 |
| O | 1.80890000 | -0.45770000 | 2.79360000 |
| H | 2.52540000 | 0.01510000 | 3.23290000 |
| H | 0.74720000 | 0.62370000 | 0.93370000 |
| N | 1.98170000 | -1.48670000 | -1.44550000 |
| C | 2.20260000 | -2.25960000 | -2.65540000 |
| H | 1.06020000 | -1.08970000 | -1.28890000 |
| H | -4.36690000 | -2.83300000 | -1.91450000 |
| H | -5.24710000 | -3.43410000 | -0.49000000 |
| H | -5.93260000 | -2.12590000 | -1.45370000 |
| H | -3.06230000 | 0.54120000 | 0.70860000 |
| H | -0.98120000 | -1.67110000 | 0.97670000 |
| H | -2.35500000 | -1.36870000 | 2.07270000 |
| H | 2.78190000 | -3.15530000 | -2.42920000 |
| H | 0.08250000 | 2.39070000 | -2.04640000 |
| H | 3.55900000 | 0.36330000 | 0.77600000 |
| H | 1.48650000 | -1.80670000 | 1.34950000 |
| H | 4.34120000 | -1.43590000 | 2.39560000 |
| H | 3.19970000 | -2.69720000 | 2.90780000 |
| H | 3.82400000 | -2.67730000 | 1.24690000 |
| H | 0.98890000 | 3.63230000 | 0.58040000 |
| H | 0.80420000 | 4.42640000 | -0.99200000 |
| H | -1.13940000 | 4.80230000 | 0.85720000 |
| H | -1.64700000 | 4.36150000 | -0.78330000 |
| H | -2.69860000 | 2.61480000 | 0.48190000 |
| H | -1.33090000 | 2.48570000 | 1.61330000 |
| H | 2.74530000 | -1.67450000 | -3.40200000 |
| H | 1.23630000 | -2.55140000 | -3.06130000 |

1 2 1.0 31 1.0 32 1.0 33 1.0
2 3 2.0 4 1.5
3
4 5 1.0 10 1.0
5 6 1.0 8 1.0 34 1.0
6 7 2.0 12 1.5
7
8 9 1.0 35 1.0 36 1.0
9 11 1.0
10
11
12 13 1.0 18 1.0
13 14 1.0 16 1.0 38 1.0
14 15 2.0 19 1.5
15
16 17 1.0 44 1.0 45 1.0
17 18 1.0 46 1.0 47 1.0

18 48 1.0 49 1.0
19 20 1.0 27 1.0
20 21 1.0 23 1.0 39 1.0
21 22 2.0 28 1.5
22
23 24 1.0 25 1.0 40 1.0
24 41 1.0 42 1.0 43 1.0
25 26 1.0
26
27
28 29 1.0 30 1.0
29 37 1.0 50 1.0 51 1.0

### Geometry-optimized structure of Ac-Ser-*cis*-Arg-OMe modified from XGPRT (pdb 6kp5)

**Ac-Ser-cis-Arg-OMe-SO4**-
optimized M06-2X/6-311++G(2d,2p)/H2O

-1 1

| | | | |
|---|---|---|---|
| N | -4.03146500 | 1.00325600 | -0.34800600 |
| C | -1.64460500 | 0.78668900 | -0.36589600 |
| O | -1.56931100 | 2.01172800 | -0.32002600 |
| O | -2.02222000 | -0.83887000 | -2.84911100 |
| H | -3.79338500 | 1.96384800 | -0.15478600 |
| H | -2.40156300 | -1.71972600 | -2.89554500 |
| N | -0.60055400 | -0.00815900 | -0.09486600 |
| O | -1.53848700 | -3.10632100 | 1.52288500 |
| N | 3.63218400 | -1.01702800 | -0.37651600 |
| C | 4.92689900 | -0.96777900 | -0.12434000 |
| N | 5.74213700 | -1.93457500 | -0.57389000 |
| N | 5.42017900 | 0.05755900 | 0.57140300 |
| H | 3.03524100 | -0.26112700 | 0.01491000 |
| H | 6.70358500 | -1.94139800 | -0.27943300 |
| H | 5.35712300 | -2.81294000 | -0.87559900 |
| H | 0.26467900 | 0.46192400 | 0.19786100 |
| H | 4.85752200 | 0.92522800 | 0.63422300 |
| H | 6.41808400 | 0.13012700 | 0.67290000 |
| O | 3.92930800 | 2.36891200 | 0.59088700 |
| O | 1.75078600 | 3.27008300 | 1.20059400 |
| O | 2.16826700 | 2.52777300 | -1.08420400 |
| O | 2.00642300 | 0.89816800 | 0.71661200 |
| S | 2.45472800 | 2.29674100 | 0.34643800 |
| C | -5.29184000 | 0.54866500 | -0.19363000 |
| O | -5.59568200 | -0.61499900 | -0.43670900 |
| C | -6.30546400 | 1.54833500 | 0.30149900 |
| H | -5.88156700 | 2.53553900 | 0.46452400 |
| H | -7.11019600 | 1.61502800 | -0.42811800 |
| H | -6.72722500 | 1.17659400 | 1.23356300 |
| C | -1.37994400 | -1.93499500 | 1.28645900 |
| O | -1.84232200 | -0.95531200 | 2.05330100 |
| C | -2.57051100 | -1.37242600 | 3.21701300 |
| H | -2.85817000 | -0.45996500 | 3.72582700 |
| H | -3.44856800 | -1.94088200 | 2.92038300 |
| H | -1.93548700 | -1.98383400 | 3.85274600 |
| C | 1.50725300 | -1.78427800 | -1.26491100 |
| H | 1.36198500 | -0.76161800 | -1.61810000 |
| H | 1.07367000 | -2.44906800 | -2.01231300 |
| C | 0.79891900 | -1.99097900 | 0.07206700 |
| H | 0.75757300 | -3.05309000 | 0.30884300 |
| H | 1.34246000 | -1.49241200 | 0.87718300 |
| C | 3.00092600 | -2.06003700 | -1.17563500 |
| H | 3.16785200 | -3.03896800 | -0.71648000 |
| H | 3.44167700 | -2.07343700 | -2.17399700 |

| | | | |
|---|---|---|---|
| C | -0.63309100 | -1.44820600 | 0.04643100 |
| H | -1.14831400 | -1.89550200 | -0.80447800 |
| C | -2.96047100 | 0.14921700 | -0.80773800 |
| H | -3.10871700 | -0.84268600 | -0.38465400 |
| C | -2.99591500 | 0.05435600 | -2.34278700 |
| H | -2.78541500 | 1.04155000 | -2.75482300 |
| H | -4.00289300 | -0.23907500 | -2.63952200 |

1 5 1.0 24 1.5 47 1.0
2 3 2.0 7 1.5 47 1.0
3
4 6 1.0 49 1.0
5
6
7 16 1.0 45 1.0
8 30 2.0
9 10 2.0 13 1.0 42 1.0
10 11 1.5 12 1.5
11 14 1.0 15 1.0
12 17 1.0 18 1.0
13
14
15
16
17
18
19 23 2.0
20 23 2.0
21 23 2.0
22 23 1.5
23
24 25 2.0 26 1.0
25
26 27 1.0 28 1.0 29 1.0
27
28
29
30 31 1.5 45 1.0
31 32 1.0
32 33 1.0 34 1.0 35 1.0
33
34
35
36 37 1.0 38 1.0 39 1.0 42 1.0
37
38
39 40 1.0 41 1.0 45 1.0
40
41
42 43 1.0 44 1.0
43

44
45 46 1.0
46
47 48 1.0 49 1.0
48
49 50 1.0 51 1.0
50
51

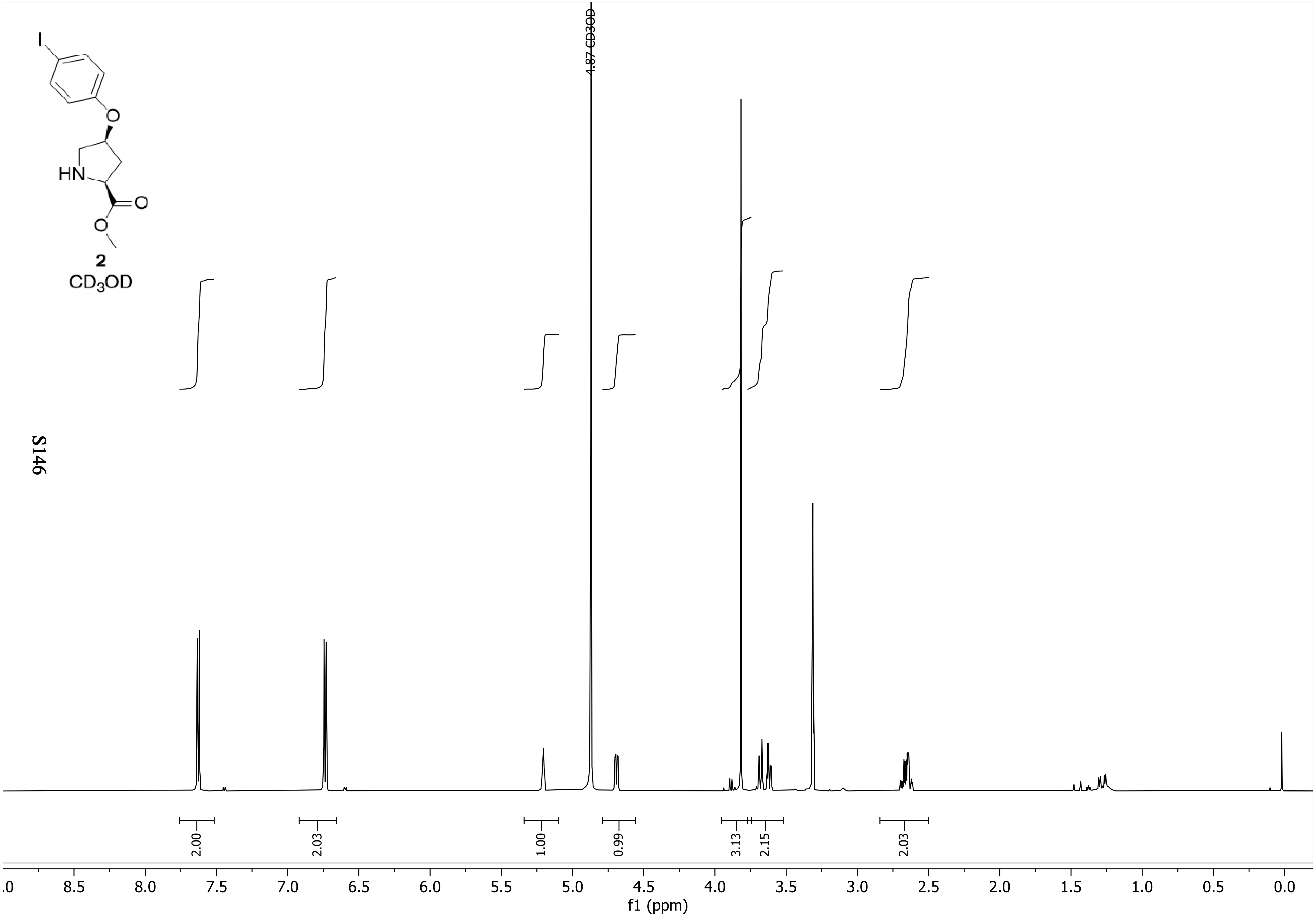

I
O
HN
O
O
2
$CD_3OD$
4.87 CD3OD
2.00
2.03
1.00
0.99
3.13
2.15
2.03
.0
8.5
8.0
7.5
7.0
6.5
6.0
5.5
5.0
4.5
4.0
3.5
3.0
2.5
2.0
1.5
1.0
0.5
0.0
f1 (ppm)

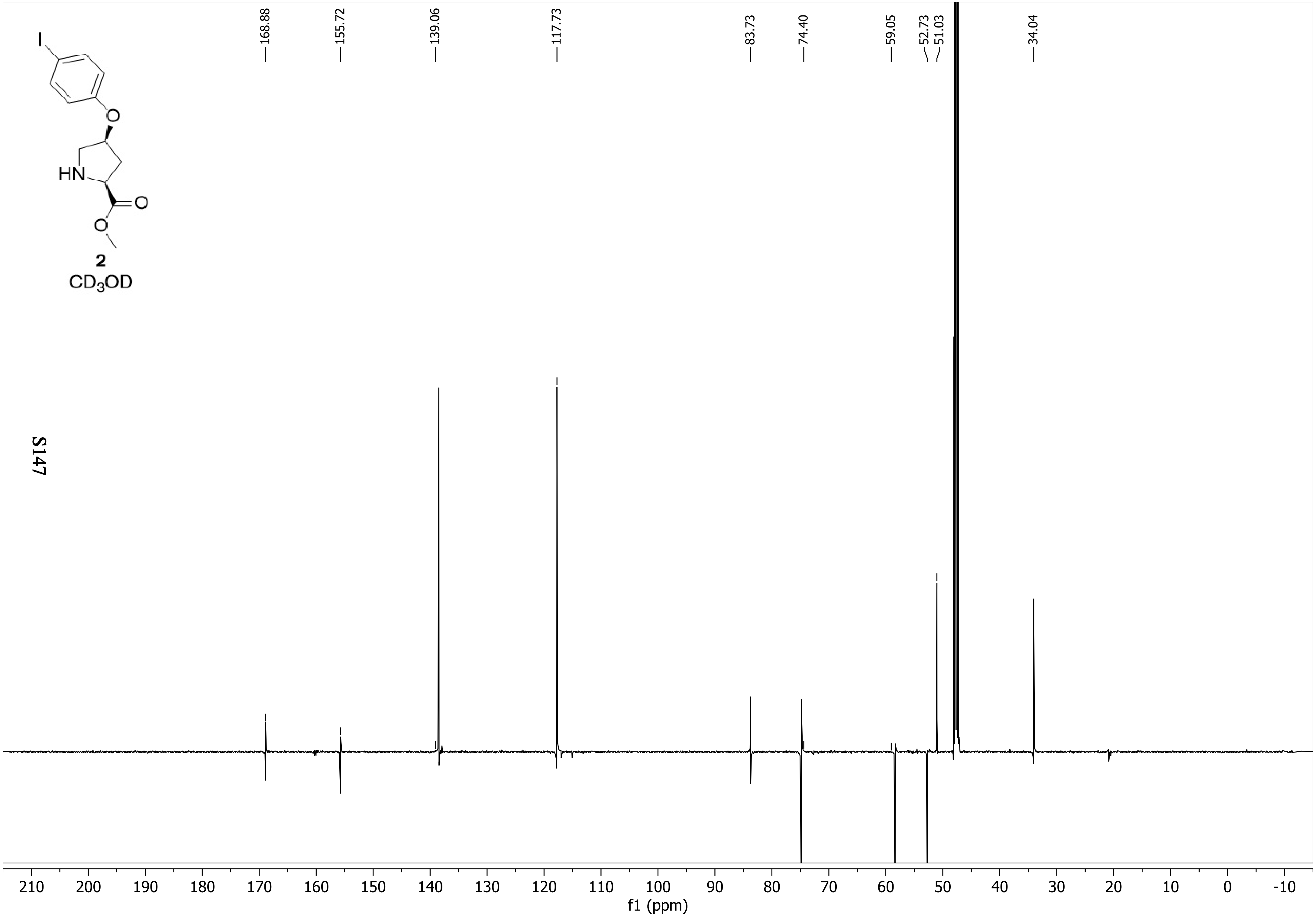
168.88
155.72
139.06
117.73
83.73
74.40
59.05
52.73
51.03
34.04
I
O
HN
O
O
2
CD3OD
210 200 190 180 170 160 150 140 130 120 110 100 90 80 70 60 50 40 30 20 10 0 -10
f1 (ppm)

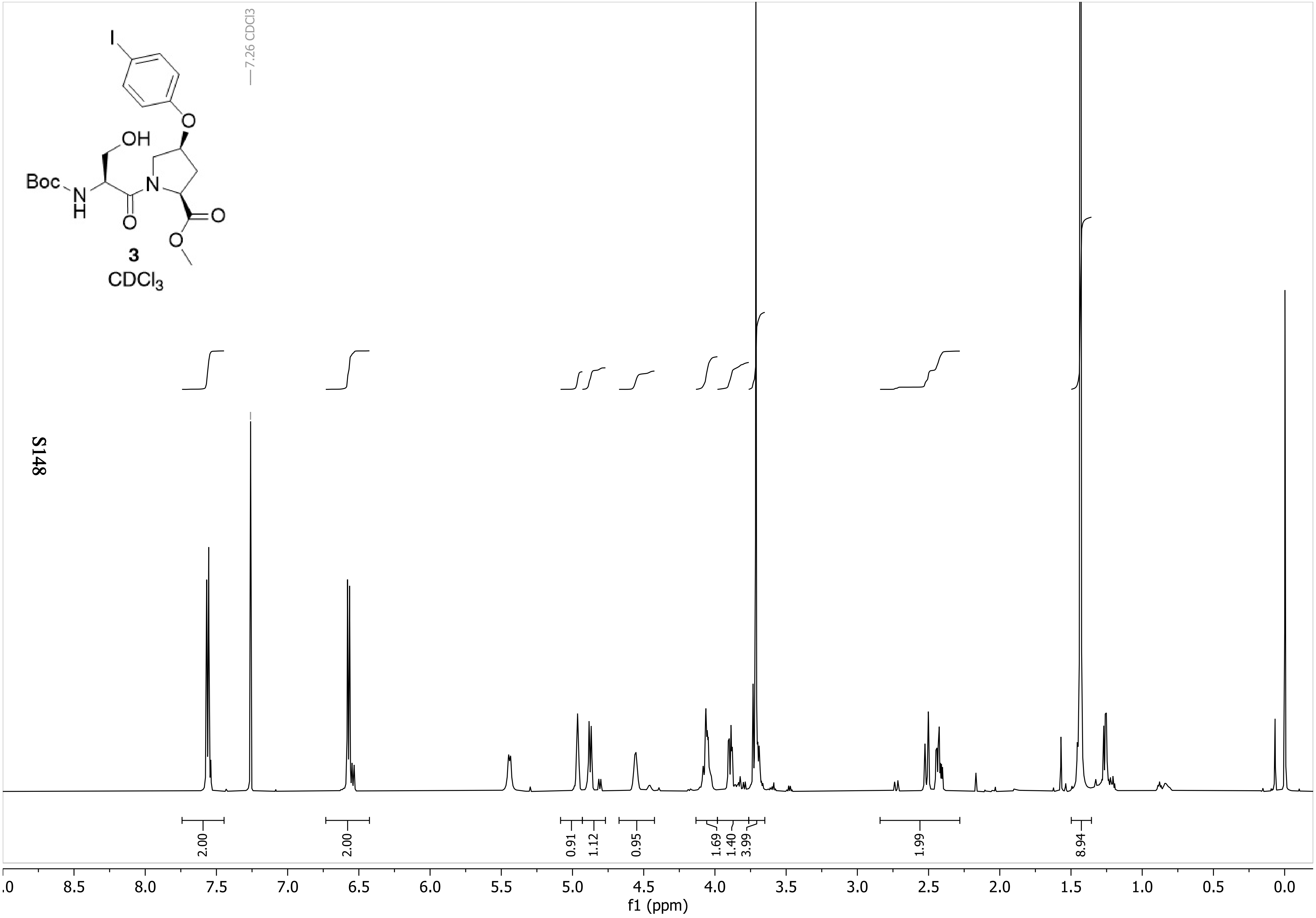

7.26 CDCl3
I
O
OH
Boc
N
H
O
N
O
O
3
CDCl3
2.00
2.00
0.91
1.12
0.95
1.69
1.40
3.99
1.99
8.94
8.5
8.0
7.5
7.0
6.5
6.0
5.5
5.0
4.5
4.0
3.5
3.0
2.5
2.0
1.5
1.0
0.5
0.0
f1 (ppm)

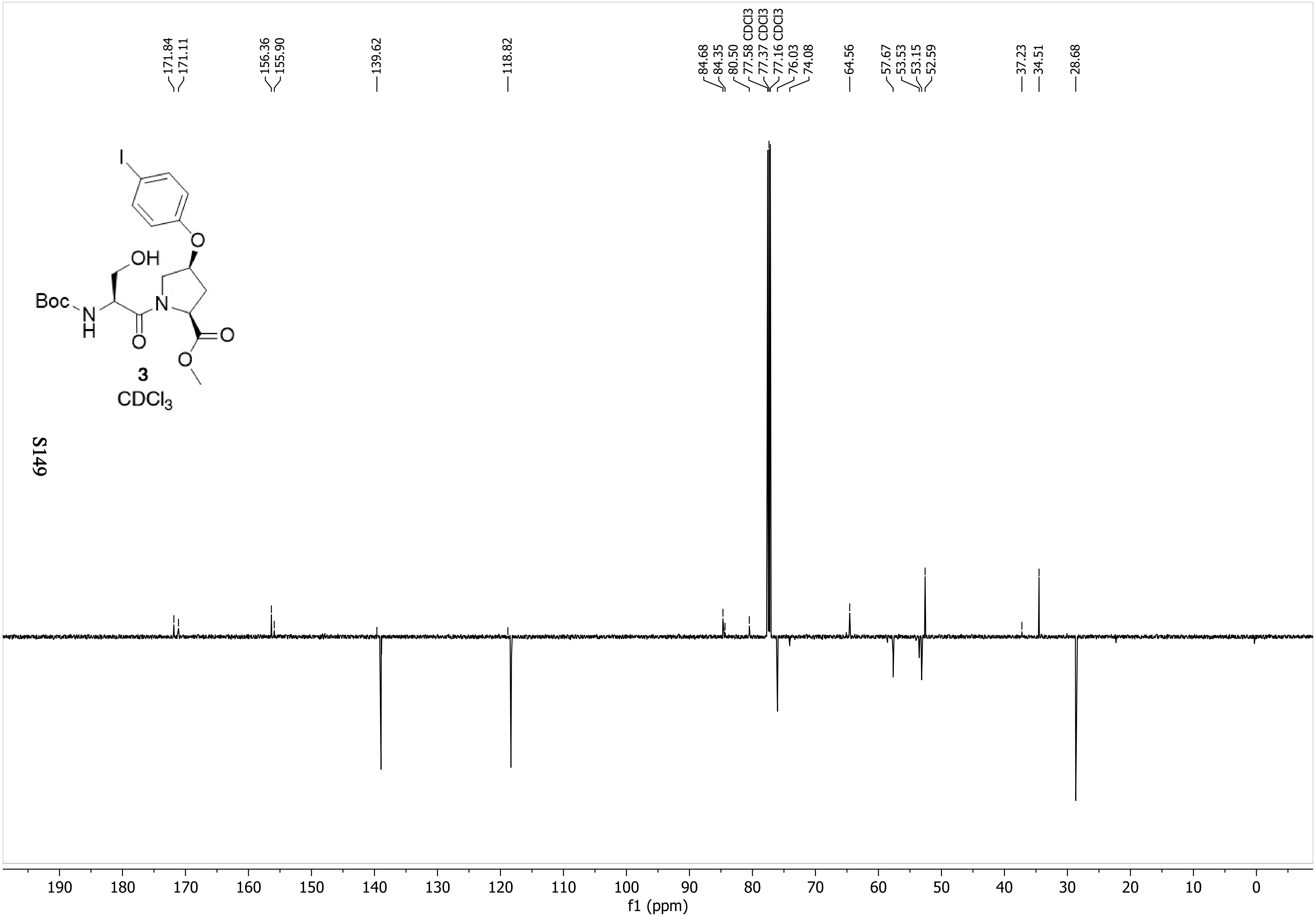
171.84
171.11
156.36
155.90
139.62
118.82
84.68
84.35
80.50
77.58 CDCl3
77.37 CDCl3
77.16 CDCl3
76.03
74.08
64.56
57.67
53.53
53.15
52.59
37.23
34.51
28.68
I
O
OH
Boc
N
H
N
O
O
O
3
CDCl3
190
180
170
160
150
140
130
120
110
100
90
80
70
60
50
40
30
20
10
0
f1 (ppm)

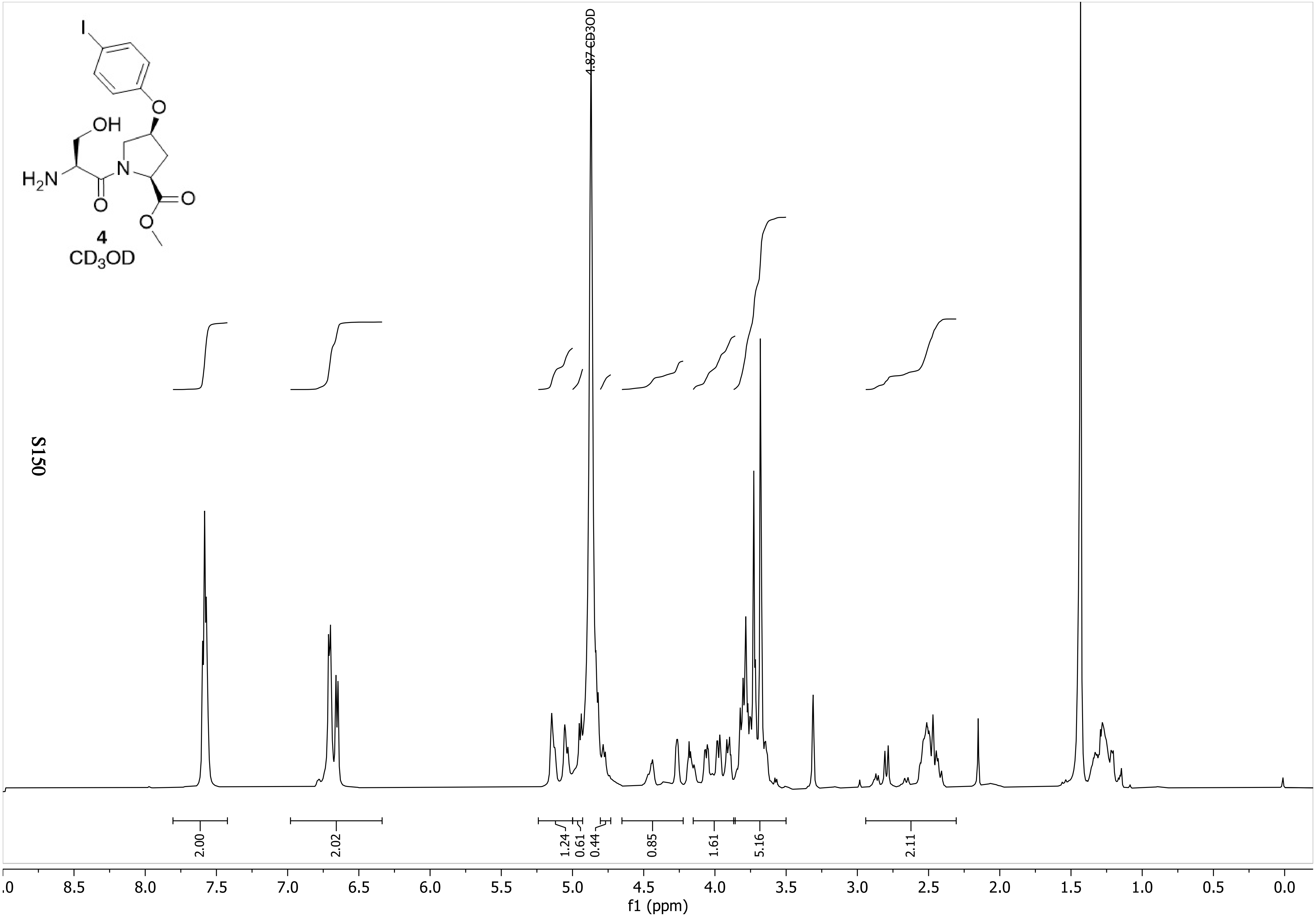

4
CD3OD
4.87 CD3OD
2.00
2.02
1.24
0.61
0.44
0.85
1.61
5.16
2.11
8.5 8.0 7.5 7.0 6.5 6.0 5.5 5.0 4.5 4.0 3.5 3.0 2.5 2.0 1.5 1.0 0.5 0.0
f1 (ppm)

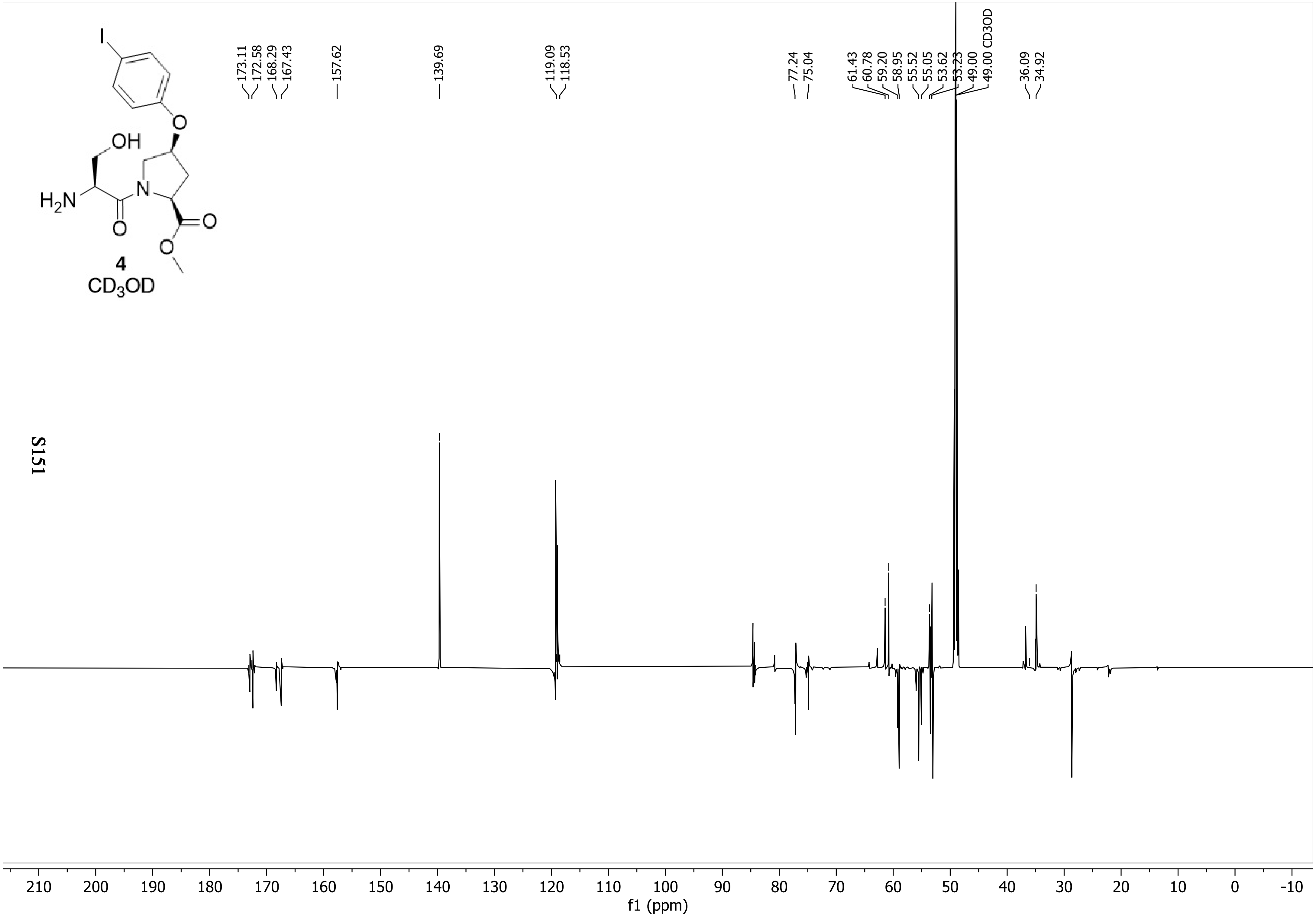

4
$CD_3OD$
173.11
172.58
168.29
167.43
157.62
139.69
119.09
118.53
77.24
75.04
61.43
60.78
59.20
58.95
55.52
55.05
53.62
53.23
49.00
49.00 CD3OD
36.09
34.92
210 200 190 180 170 160 150 140 130 120 110 100 90 80 70 60 50 40 30 20 10 0 -10
f1 (ppm)

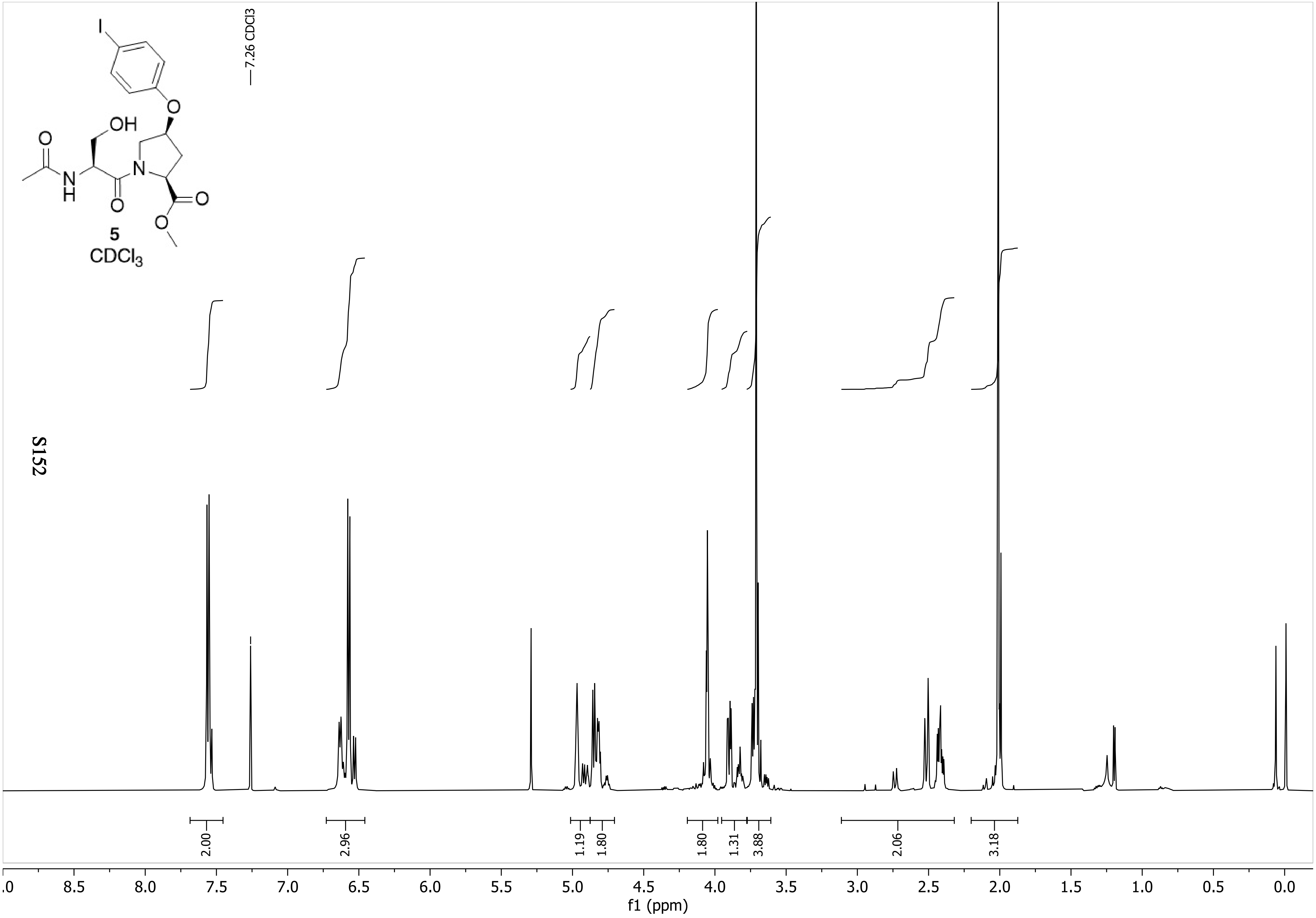

7.26 CDCl3
5
$CDCl_3$
2.00
2.96
1.19
1.80
1.80
1.31
3.88
2.06
3.18
8.5
8.0
7.5
7.0
6.5
6.0
5.5
5.0
4.5
4.0
3.5
3.0
2.5
2.0
1.5
1.0
0.5
0.0
f1 (ppm)

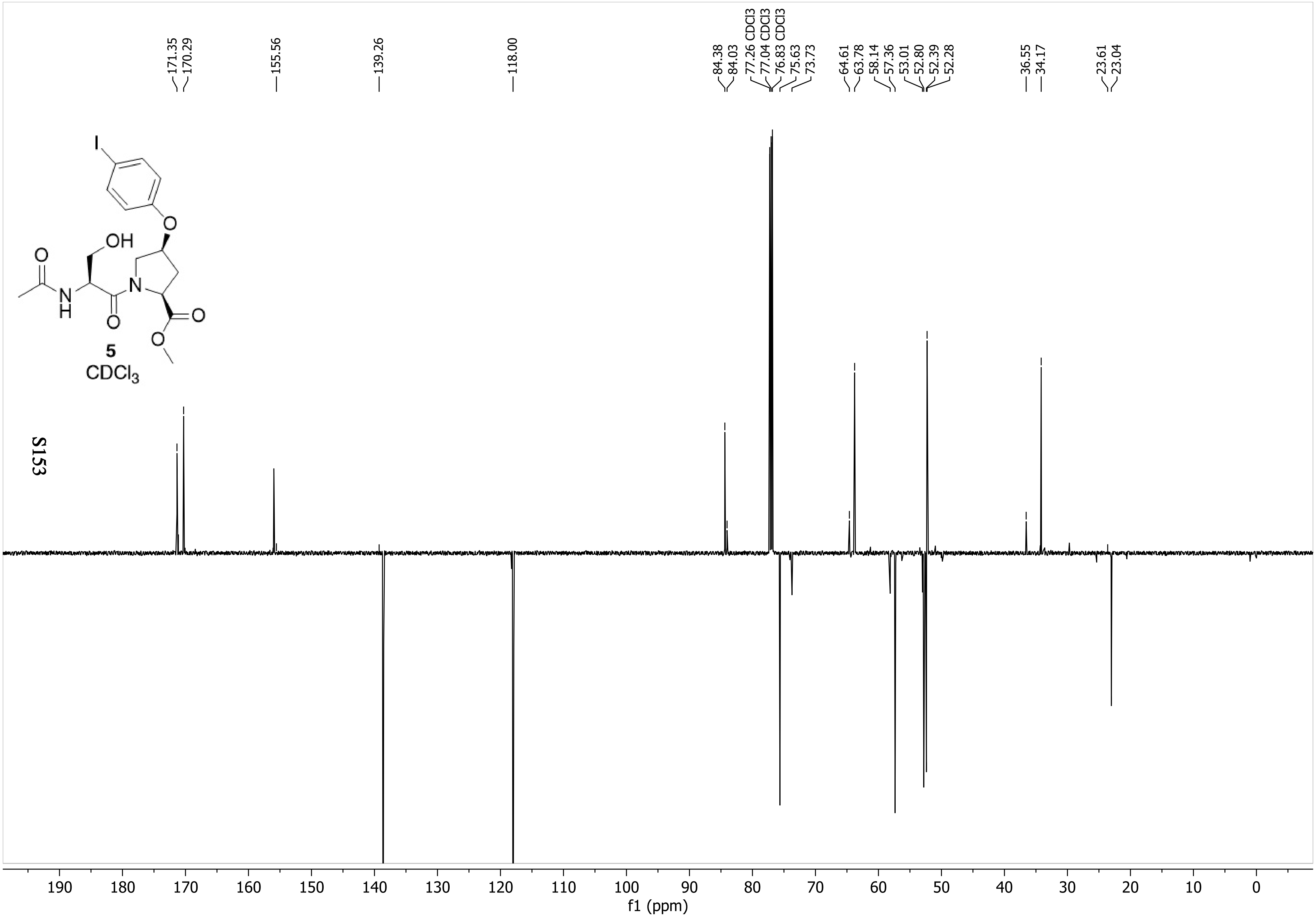
171.35
170.29
155.56
139.26
118.00
84.38
84.03
77.26 CDCl3
77.04 CDCl3
76.83 CDCl3
75.63
73.73
64.61
63.78
58.14
57.36
53.01
52.80
52.39
52.28
36.55
34.17
23.61
23.04
I
OH
O
N
H
O
O
O
O
5
$CDCl_3$
190 180 170 160 150 140 130 120 110 100 90 80 70 60 50 40 30 20 10 0
f1 (ppm)

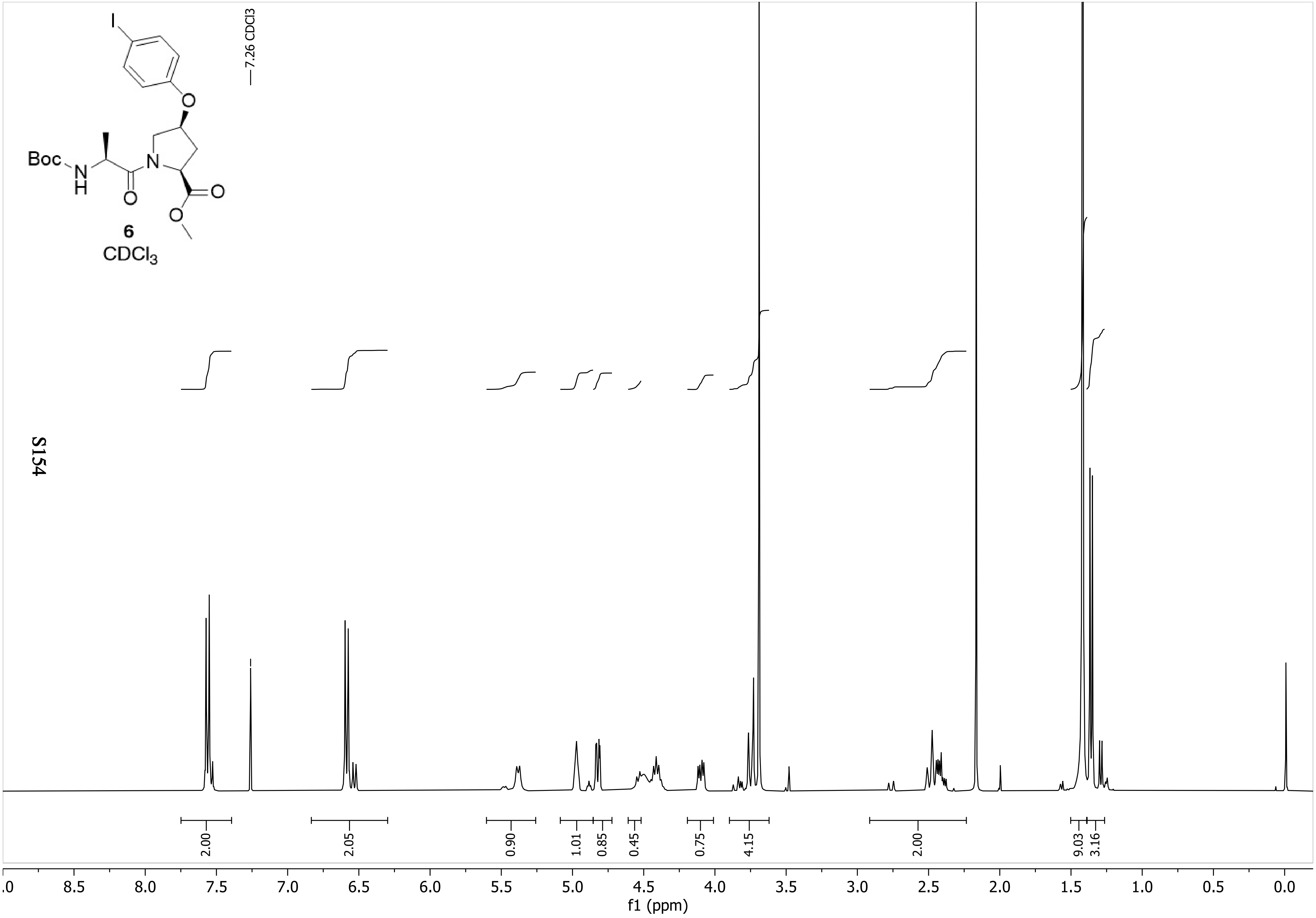

I
O
Boc
N
H
N
O
O
O
6
CDCl3
7.26 CDCl3
2.00
2.05
0.90
1.01
0.85
0.45
0.75
4.15
2.00
9.03
3.16
.0
8.5
8.0
7.5
7.0
6.5
6.0
5.5
5.0
4.5
4.0
3.5
3.0
2.5
2.0
1.5
1.0
0.5
0.0
f1 (ppm)

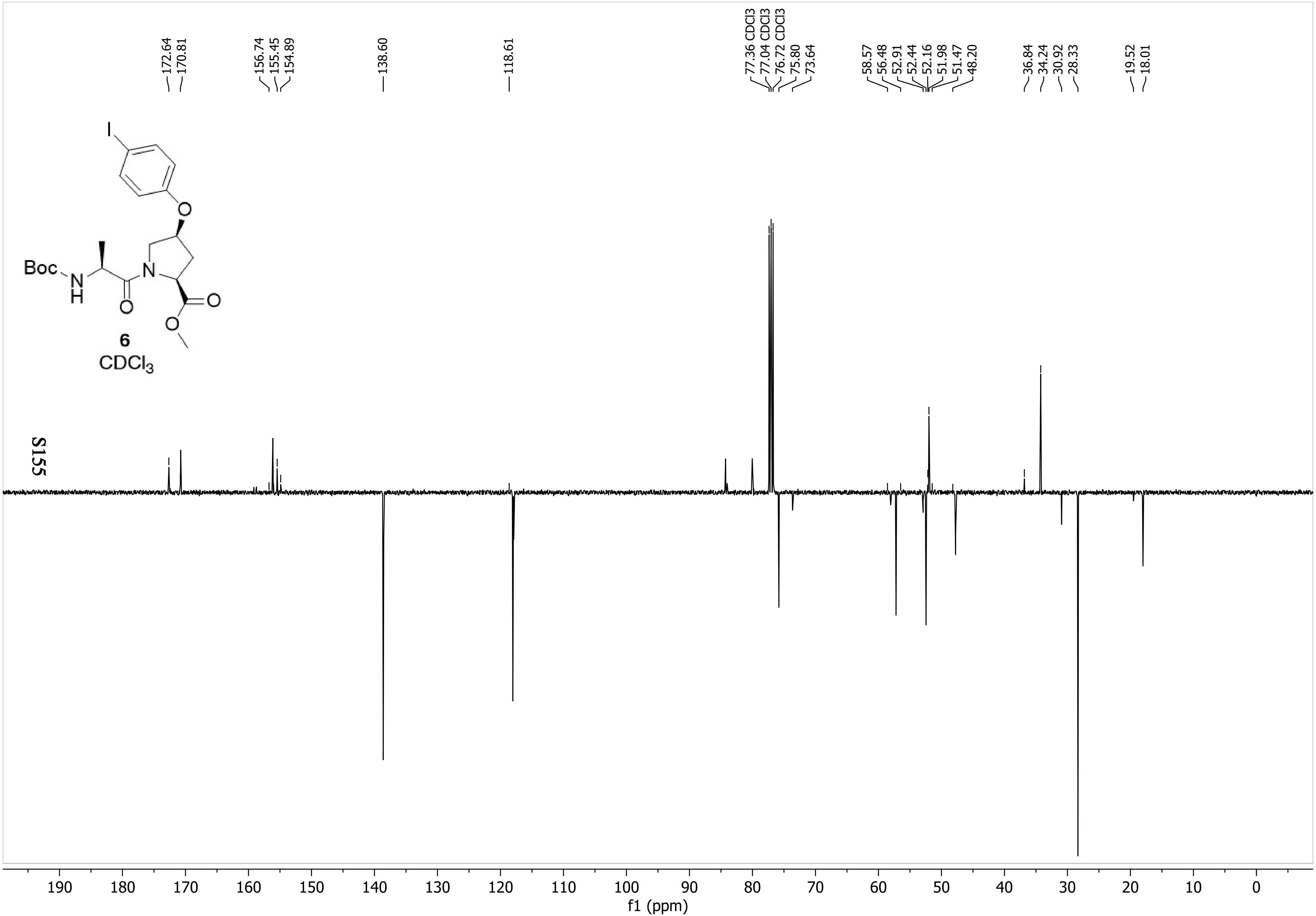
172.64
170.81
156.74
155.45
154.89
138.60
118.61
77.36 CDCl3
77.04 CDCl3
76.72 CDCl3
75.80
73.64
58.57
56.48
52.91
52.44
52.16
51.98
51.47
48.20
36.84
34.24
30.92
28.33
19.52
18.01
I
O
Boc
N
H
N
O
O
O
6
CDCl3
190 180 170 160 150 140 130 120 110 100 90 80 70 60 50 40 30 20 10 0
f1 (ppm)

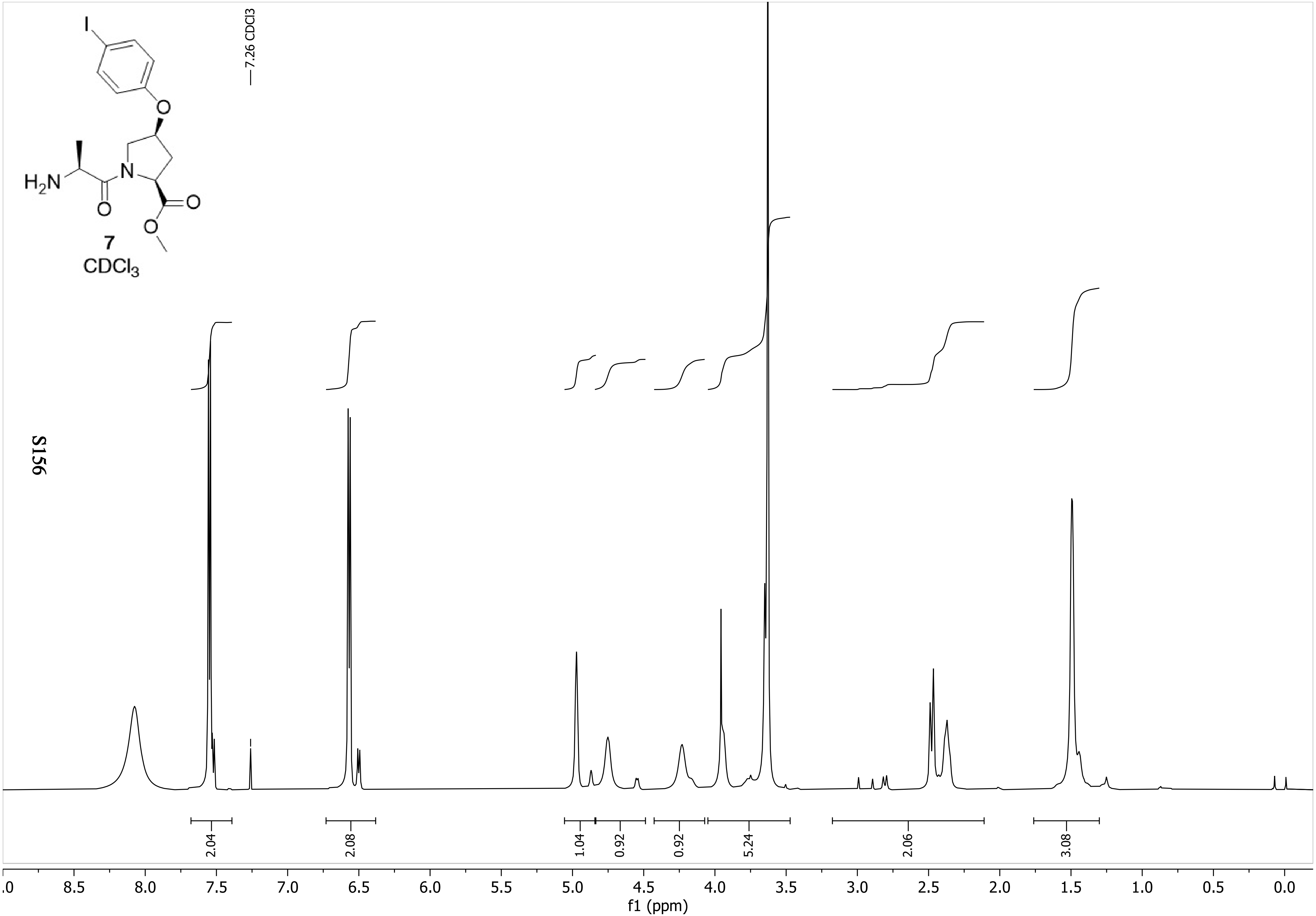
I
O
H2N
N
O
O
O
7
CDCl3
7.26 CDCl3
2.04
2.08
1.04
0.92
0.92
5.24
2.06
3.08
.0
8.5
8.0
7.5
7.0
6.5
6.0
5.5
5.0
4.5
4.0
3.5
3.0
2.5
2.0
1.5
1.0
0.5
0.0
f1 (ppm)

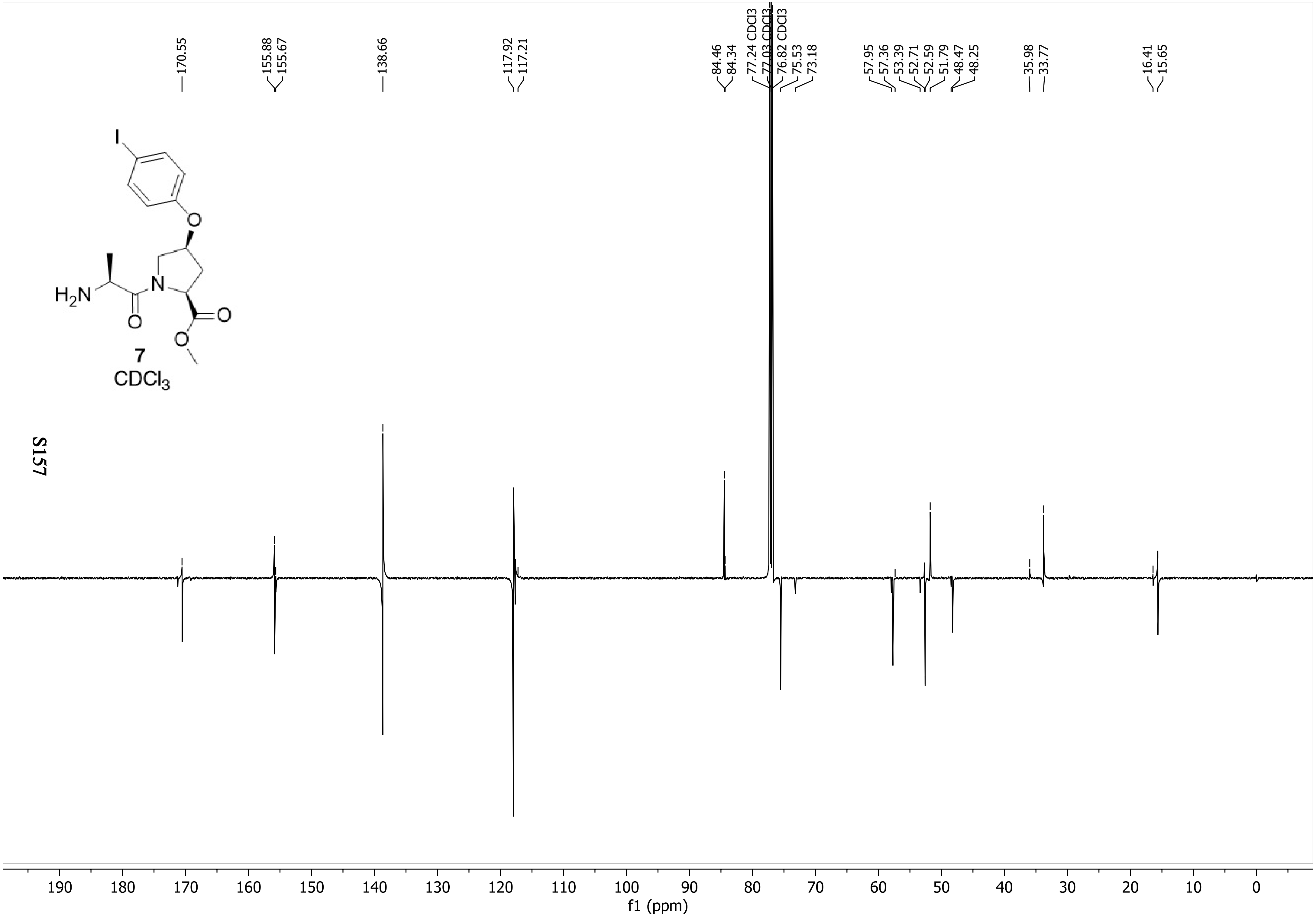
170.55
155.88
155.67
138.66
117.92
117.21
84.46
84.34
77.24 CDCl3
77.03 CDCl3
76.82 CDCl3
75.53
73.18
57.95
57.36
53.39
52.71
52.59
51.79
48.47
48.25
35.98
33.77
16.41
15.65
I
O
$H_2N$
N
O
O
O
7
$CDCl_3$
190 180 170 160 150 140 130 120 110 100 90 80 70 60 50 40 30 20 10 0
f1 (ppm)

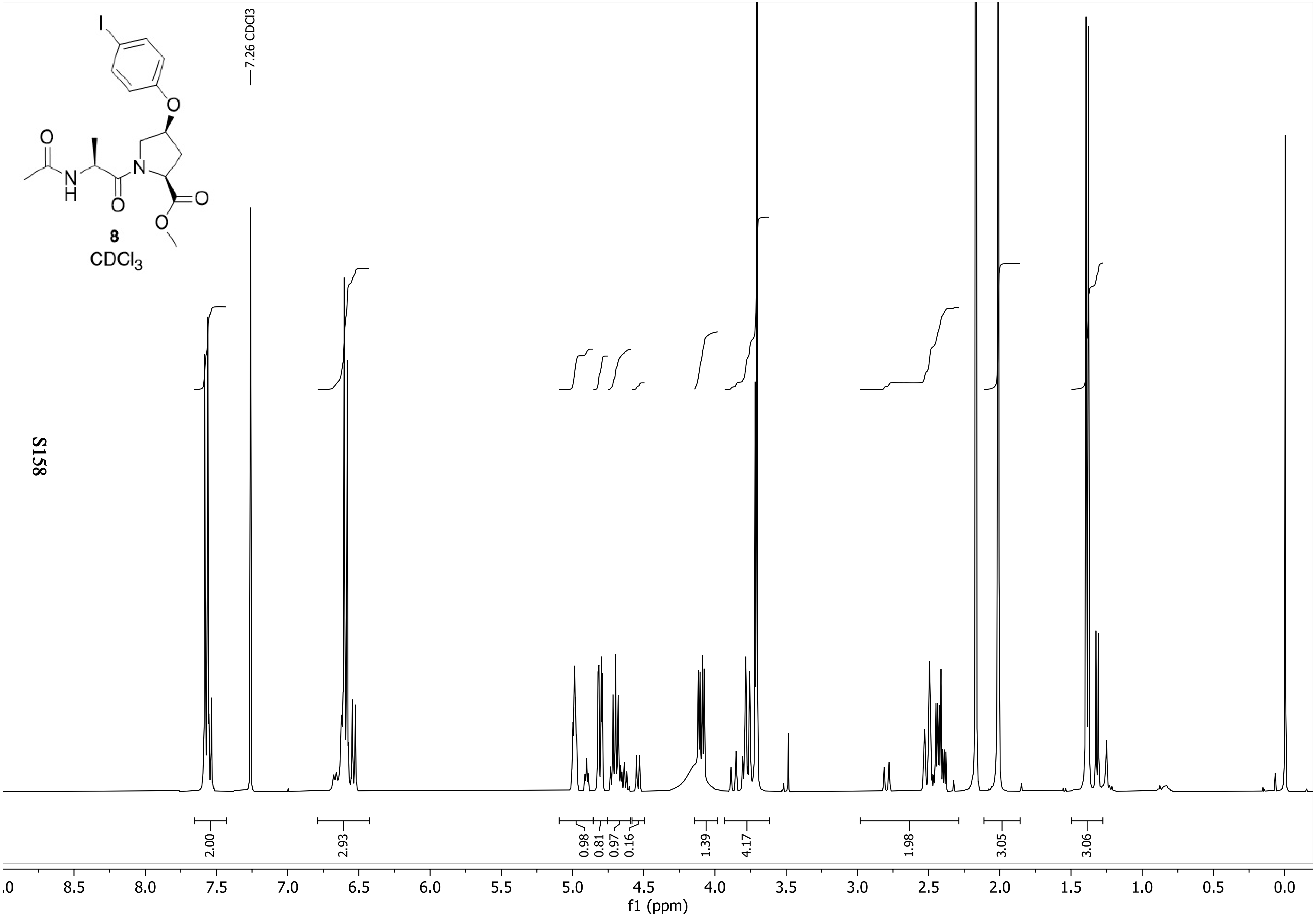
7.26 CDCl3
8
CDCl3
2.00
2.93
0.98
0.81
0.97
0.16
1.39
4.17
1.98
3.05
3.06
.0
8.5
8.0
7.5
7.0
6.5
6.0
5.5
5.0
4.5
4.0
3.5
3.0
2.5
2.0
1.5
1.0
0.5
0.0
f1 (ppm)

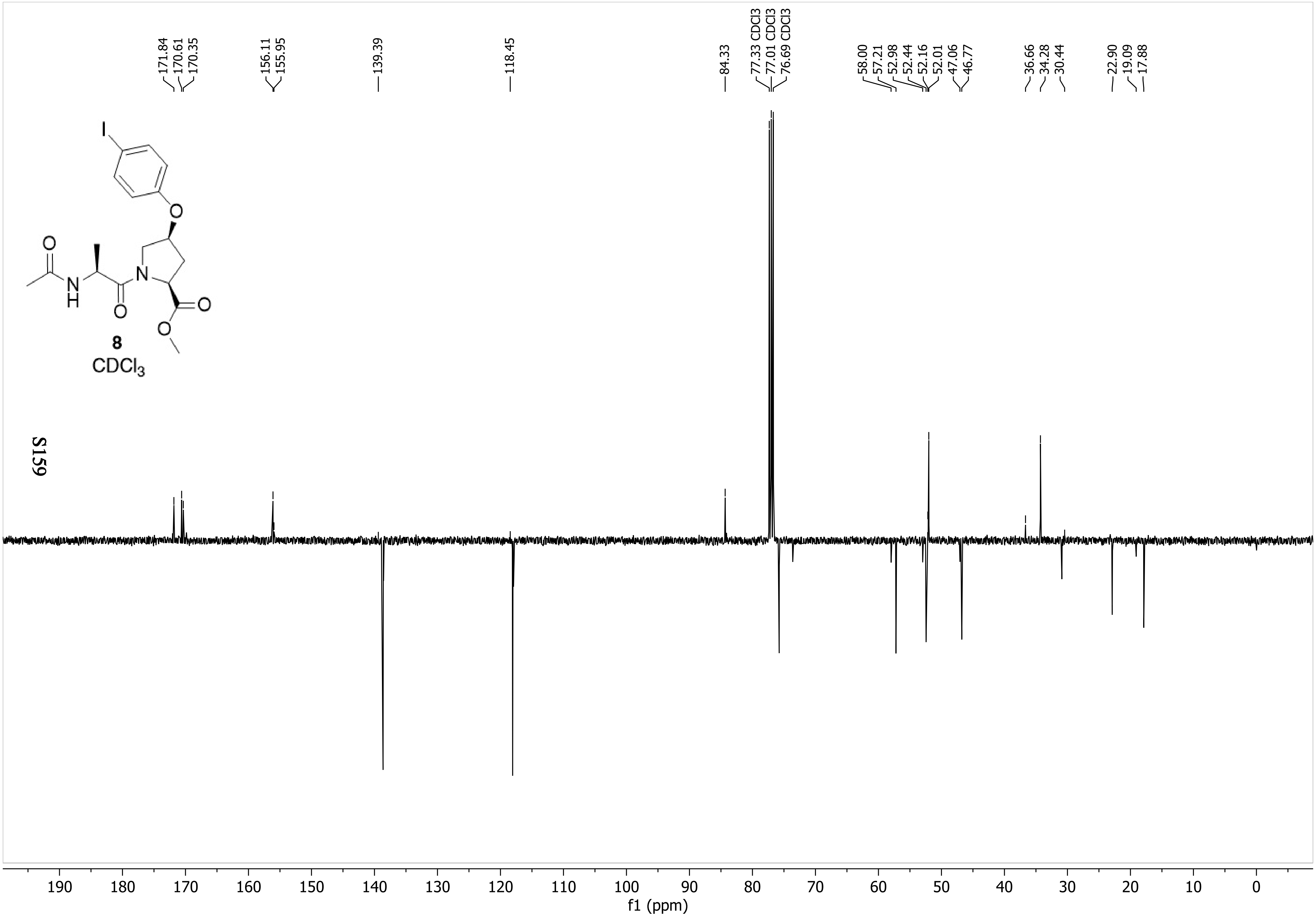
171.84
170.61
170.35
156.11
155.95
139.39
118.45
84.33
77.33 CDCl3
77.01 CDCl3
76.69 CDCl3
58.00
57.21
52.98
52.44
52.16
52.01
47.06
46.77
36.66
34.28
30.44
22.90
19.09
17.88
I
O
N
H
8
CDCl3
190 180 170 160 150 140 130 120 110 100 90 80 70 60 50 40 30 20 10 0
f1 (ppm)
